# Genome-wide association analyses of risk tolerance and risky behaviors in over 1 million individuals identify hundreds of loci and shared genetic influences^1^

**DOI:** 10.1101/261081

**Authors:** Richard Karlsson Linnér, Pietro Biroli, Edward Kong, S Fleur W Meddens, Robbee Wedow, Mark Alan Fontana, Maël Lebreton, Abdel Abdellaoui, Anke R Hammerschlag, Michel G Nivard, Aysu Okbay, Cornelius A Rietveld, Pascal N Timshel, Stephen P Tino, Maciej Trzaskowski, Ronald de Vlaming, Christian L Zünd, Yanchun Bao, Laura Buzdugan, Ann H Caplin, Chia-Yen Chen, Peter Eibich, Pierre Fontanillas, Juan R Gonzalez, Peter K Joshi, Ville Karhunen, Aaron Kleinman, Remy Z Levin, Christina M Lill, Gerardus A Meddens, Gerard Muntané, Sandra Sanchez-Roige, Frank J van Rooij, Erdogan Taskesen, Yang Wu, Futao Zhang, 23andMe Research Team, eQTLgen Consortium, International Cannabis Consortium, Psychiatric Genomics Consortium, Social Science Genetic Association Consortium,, Adam Auton, Jason D Boardman, David W Clark, Andrew Conlin, Conor C Dolan, Urs Fischbacher, Patrick JF Groenen, Kathleen Mullan Harris, Gregor Hasler, Albert Hofman, Mohammad A Ikram, Sonia Jain, Robert Karlsson, Ronald C Kessler, Maarten Kooyman, James MacKillop, Minna Männikkö, Carlos Morcillo-Suarez, Matthew B McQueen, Klaus M Schmidt, Melissa C Smart, Matthias Sutter, A Roy Thurik, Andre G Uitterlinden, Jon White, Harriet de Wit, Jian Yang, Lars Bertram, Dorret Boomsma, Tõnu Esko, Ernst Fehr, David A Hinds, Magnus Johannesson, Meena Kumari, David Laibson, Patrik KE Magnusson, Michelle N Meyer, Arcadi Navarro, Abraham A Palmer, Tune H Pers, Danielle Posthuma, Daniel Schunk, Murray B Stein, Rauli Svento, Henning Tiemeier, Paul RHJ Timmers, Patrick Turley, Robert J Ursano, Gert G Wagner, James F Wilson, Jacob Gratten, James J Lee, David Cesarini, Daniel J Benjamin, Philipp D Koellinger, Jonathan P Beauchamp

## Abstract

Humans vary substantially in their willingness to take risks. In a combined sample of over one million individuals, we conducted genome-wide association studies (GWAS) of general risk tolerance, adventurousness, and risky behaviors in the driving, drinking, smoking, and sexual domains. We identified 611 approximately independent genetic loci associated with at least one of our phenotypes, including 124 with general risk tolerance. We report evidence of substantial shared genetic influences across general risk tolerance and risky behaviors: 72 of the 124 general risk tolerance loci contain a lead SNP for at least one of our other GWAS, and general risk tolerance is moderately to strongly genetically correlated (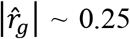 to 0.50) with a range of risky behaviors. Bioinformatics analyses imply that genes near general-risk-tolerance-associated SNPs are highly expressed in brain tissues and point to a role for glutamatergic and GABAergic neurotransmission. We find no evidence of enrichment for genes previously hypothesized to relate to risk tolerance.

## Main Text

Choices in important domains of life, including health, fertility, finance, employment, and social relationships, rarely have consequences that can be anticipated perfectly. The degree of variability in possible outcomes is called risk. Risk tolerance—defined as the willingness to take risks, typically to obtain some reward—varies substantially across humans and has been actively studied in the behavioral and social sciences. An individual’s risk tolerance may vary across domains, but survey-based measures of *general* risk tolerance (e.g., “Would you describe yourself as someone who takes risks?”) have been found to be good all-around predictors of risky behaviors such as portfolio allocation, occupational choice, smoking, drinking alcohol, and starting one’s own business^1–3^.

Twin studies have established that various measures of risk tolerance are moderately heritable (*h*^2^~30%, although estimates in the literature vary^3–5^). Discovery of specific genetic variants associated with general risk tolerance could advance our understanding of how genetic influences are amplified and dampened by environmental factors; provide insights into underlying biological pathways; enable the construction of polygenic scores (indexes of many genetic variants) that can be used as overall measures of genetic influences on individuals; and help distinguish genetic variation associated with general versus domain-specific risk tolerance.

Although risk tolerance has been one of the most studied phenotypes in social science genetics, most claims of positive findings have been based on small-sample candidate gene studies (**Supplementary Table 11.1**), whose limitations are now appreciated^6^. To date, only two loci associated with risk tolerance have been identified in genome-wide association studies (GWAS)^7,8^.

Here, we report results from large-scale GWAS of self-reported general risk tolerance (our primary phenotype) and six supplementary phenotypes: “adventurousness” (defined as the self-reported tendency to be adventurous vs. cautious); four risky behaviors: “automobile speeding propensity” (the tendency to drive faster than the speed limit), “drinks per week” (the average number of alcoholic drinks consumed per week), “ever smoker” (whether one has ever been a smoker), and “number of sexual partners” (the lifetime number of sexual partners); and the first principal component (PC) of these four risky behaviors, which we interpret as capturing the general tendency to take risks across domains. All seven phenotypes are coded such that higher phenotype values are associated with higher risk tolerance or risk taking. **Table 1** lists, for each GWAS, the datasets we analyzed and the GWAS sample size.

**Table 1.**
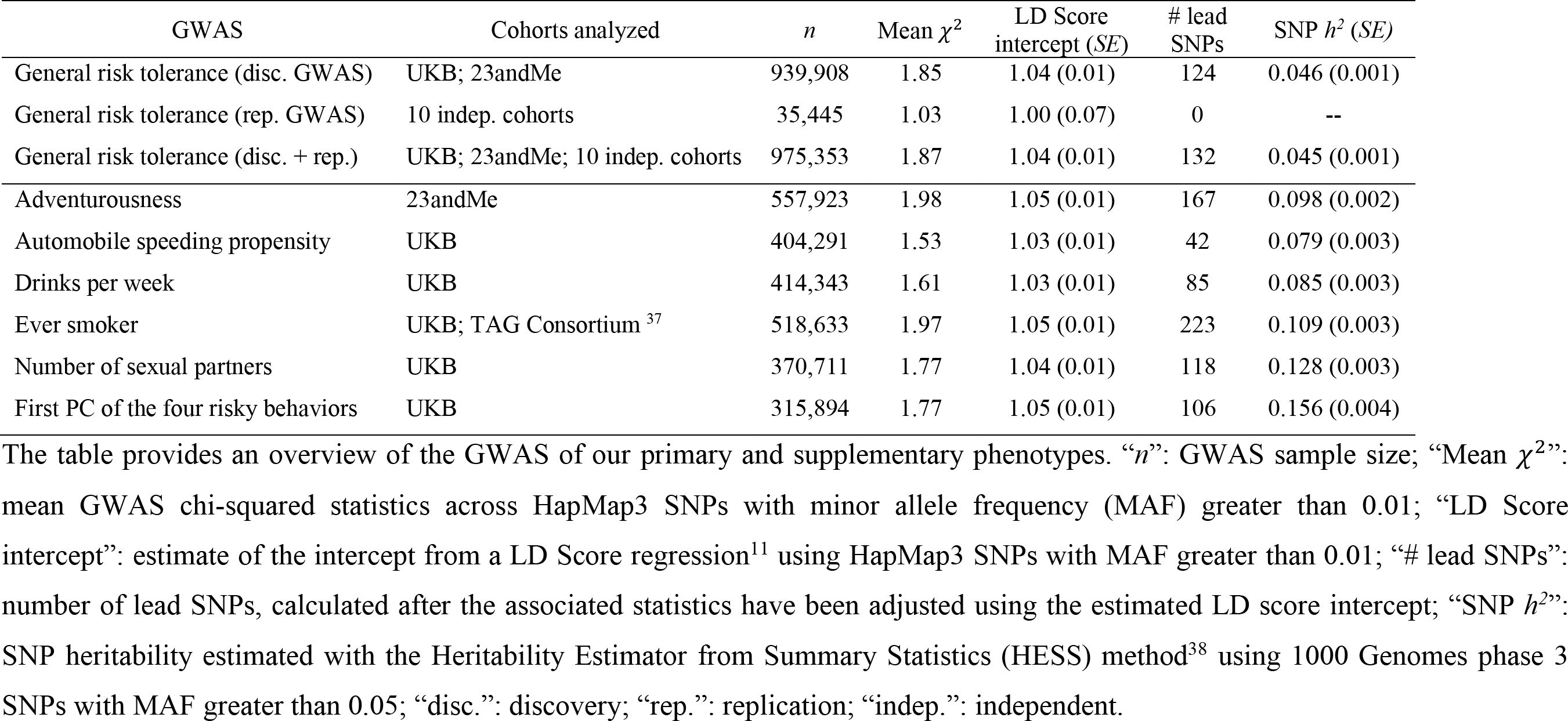
GWAS results.

### Association analyses

All seven GWAS were performed in European-ancestry subjects, following procedures described in a pre-specified analysis plan (https://osf.io/cjx9m/) and in **Supplementary Information section 2**.

In the discovery phase of our GWAS of general risk tolerance (*n* = 939,908), we performed a sample-size-weighted meta-analysis of results from the UK Biobank (UKB, *n* = 431,126) and a sample of research participants from 23andMe (*n* = 508,782). The UKB measure of general risk tolerance is based on the question: “Would you describe yourself as someone who takes risks? Yes / No.” The 23andMe measure is based on a question about overall comfort taking risks, with five response options ranging from “very comfortable” to “very uncomfortable.” The genetic correlation^9^ between the UKB and 23andMe cohorts (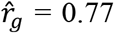, *SE* = 0.02) is smaller than one but high enough to justify our approach of pooling the two cohorts^10^.

The Q-Q plot (**Extended Data Fig. 3.2a**) from the discovery GWAS exhibits substantial inflation (*λ*_*GC*_ = 1.41). According to the estimated intercept from a linkage disequilibrium (LD) Score regression^11^, only a small share of this inflation (~5%) in test statistics is due to bias. To account for this bias, we inflated GWAS standard errors by the square root of the LD Score regression intercept.

We identified 124 approximately independent SNPs (pairwise *r*^*2*^ < 0.1) that attained genome-wide significance (*P* < 5×10^−8^). These 124 “lead SNPs” are listed in **Supplementary Table 3.1** and shown in **Fig. 1a**. All have coefficients of determination (*R*^*2*^’s) below 0.02%, and the SNP with the largest per-allele effect is estimated to increase general risk tolerance by ~0.026 standard deviations in our discovery sample (**Extended Data Fig. 3.3**).

**Figure 1.**
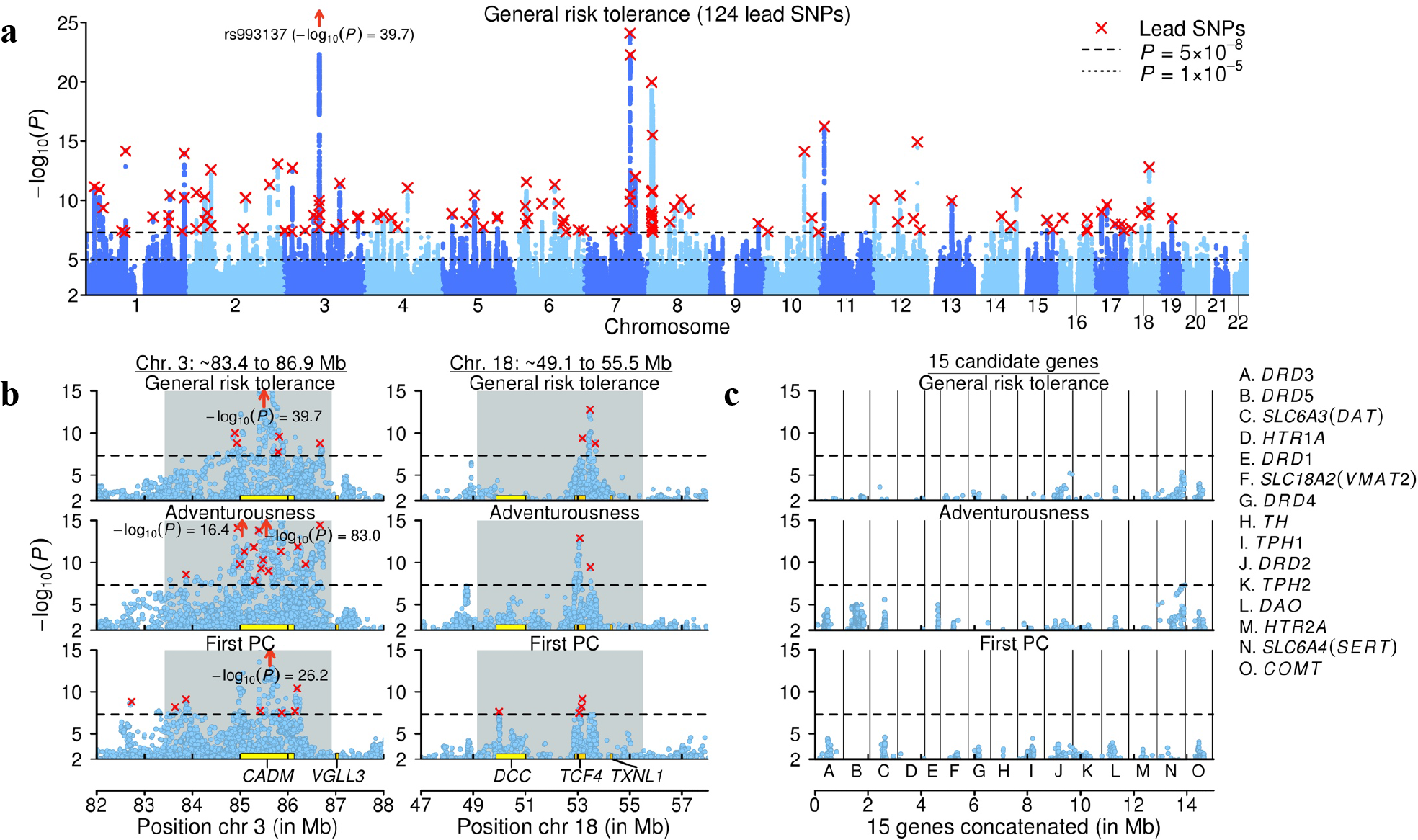
Manhattan plots. In all panels, the *x*-axis is chromosomal position; the *y*-axis is the significance on a −log_10_ scale; the horizontal dashed line marks the threshold for genome-wide significance (*P* = 5×10^−8^); and each approximately independent (pairwise *r*^2^ < 0.1) genome-wide significant association (“lead SNP”) is marked by a red ×. **a**, Manhattan plots for the discovery GWAS of general risk tolerance. **b**, Local Manhattan plots of two genomic regions that contain lead SNPs for all seven of our GWAS. The gray background marks the locations of long-range LD or candidate inversion regions. **c**, Local Manhattan plots of the loci around the 15 most commonly tested candidate genes in the prior literature on the genetics of risk tolerance. Each locus comprises all SNPs within 500 kb of the gene’s borders that are in LD (*r*^2^ > 0.1) with a SNP in the gene. The 15 plots are concatenated and shown together in the panel, divided by the black vertical lines. The 15 genes are not particularly strongly associated with general risk tolerance or the risky behaviors, as can be seen by comparing the results within each row across panels **b** and **c** (the three rows correspond to the GWAS of general risk tolerance, adventurousness, and the first PC of the four risky behaviors).

In the replication phase of our GWAS of general risk tolerance (combined *n* = 35,445), we meta-analyzed summary statistics from ten smaller cohorts. Additional details on cohort-level phenotype measures are provided in **Supplementary Table 1.2**. The questions differ in terms of their exact wording and number of response categories, but all questions ask subjects about their overall or general attitudes toward risk. The genetic correlation^9^ between the discovery and replication GWAS is 0.83 (*SE* = 0.13). 123 of the 124 lead SNPs were available or well proxied by an available SNP in the replication GWAS results. Out of the 123 SNPs, 94 have a concordant sign (*P* = 1.7×10^−9^) and 23 are significant at the 5% level in one-tailed *t* tests (*P* = 4.5×10^−8^) (**Extended Data Fig. 5.1**). This empirical replication record matches theoretical projections that take into account sampling variation and the winner’s curse (**Supplementary Information section 5**).

Our six supplementary GWAS—of adventurousness, four risky behaviors, and their principal component (*n* = 315,894 to 557,923; **Supplementary Tables 1.1-1.2**)—were conducted using methods comparable to those in the primary GWAS, but without a replication phase. **Extended Data Fig. 3.2** (**c** to **h**) shows Q-Q plots and **Extended Data Fig. 3.1** (**a** to **f**) shows Manhattan plots.

**Table 1** provides a summary overview of the seven GWAS. We identified a total of 865 lead SNPs across the seven GWAS. The lead SNPs are located in 611 approximately independent loci, where a locus is defined as the set of all SNPs in weak LD (pairwise *r*^*2*^ > 0.1) with a lead SNP. The SNP heritabilities of the seven phenotypes range from ~0.05 (for general risk tolerance) to ~0.16 (for the first PC of the four risky behaviors).

### Genetic overlap

There is substantial overlap across the results of our GWAS. For example, 72 of the 124 general-risk-tolerance lead SNPs are in loci that also contain lead SNPs for at least one of the other GWAS, including 45 for adventurousness and 49 for at least one of the four risky behaviors or their first PC. To empirically assess if this overlap could be attributed to chance, we conducted a resampling exercise under the null hypothesis that the lead SNPs of our supplementary GWAS are distributed independently of the 124 general-risk-tolerance lead loci. We strongly rejected this null hypothesis (**P** < 0.0001; **Supplementary Information section 3.3.3**).

Several regions of the genome stand out for being associated both with general risk tolerance and with all or most of the supplementary phenotypes. We tested whether the signs of the lead SNPs located in these regions tend to be concordant across our primary and supplementary GWAS. We strongly rejected the null hypothesis of no concordance (*P* < 3×10^−30^; **Supplementary Information section 3.2.3**), suggesting that these regions represent shared genetic influences, rather than colocalization of causal SNPs. **Fig. 1b** and **Extended Data Fig. 3.4** show local Manhattan plots for some of these regions. The long-range LD region^12^ on chromosome 3 (~83.4 to 86.9 Mb) contains lead SNPs from all seven GWAS as well as the most significant lead SNP from the general risk tolerance GWAS, rs993137 (*P* = 2.14×10^−40^), which is located in the gene *CADM2*. Another long-range LD region, on chromosome 6 (~25.3 to 33.4 Mb), covers the HLA-complex and contains lead SNPs from all GWAS except drinks per week. Three candidate inversions (i.e., genomic regions that are highly prone to inversion polymorphisms; **Supplementary Information section 2.9.2**) on chromosomes 7 (~124.6 to 132.7 Mb), 8 (~7.89 to 11.8 Mb), and 18 (~49.1 to 55.5 Mb) contain lead SNPs from six, five, and all seven of our GWAS, respectively. Finally, four other LD blocks^13^ that do not overlap known long-range LD or candidate inversion regions each contain lead SNPs from five of our GWAS (including general risk tolerance). The two long-range LD regions and the three candidate inversions have previously been found to be associated with numerous phenotypes, including many cognitive and neuropsychiatric phenotypes^14^.

To investigate genetic overlap at the genome-wide level, we estimated genetic correlations with self-reported general risk tolerance using bivariate LD Score regression^9^. (For this and all subsequent analyses involving general risk tolerance, we used the summary statistics from the combined meta-analysis of our discovery and replication GWAS.) The estimated genetic correlations with our six supplementary phenotypes are all positive, larger than ~0.25, and highly significant (*P* < 2.3×10^−30^; **Fig. 2**), indicating that SNPs associated with higher general risk tolerance also tend to be associated with riskier behavior. The largest estimated genetic correlations are with adventurousness (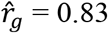, *SE* = 0.01), number of sexual partners (0.52, *SE* = 0.02), automobile speeding propensity (0.45, *SE* = 0.02), and the first PC of the four risky behaviors (0.50, *SE* = 0.02).

**Figure 2.**
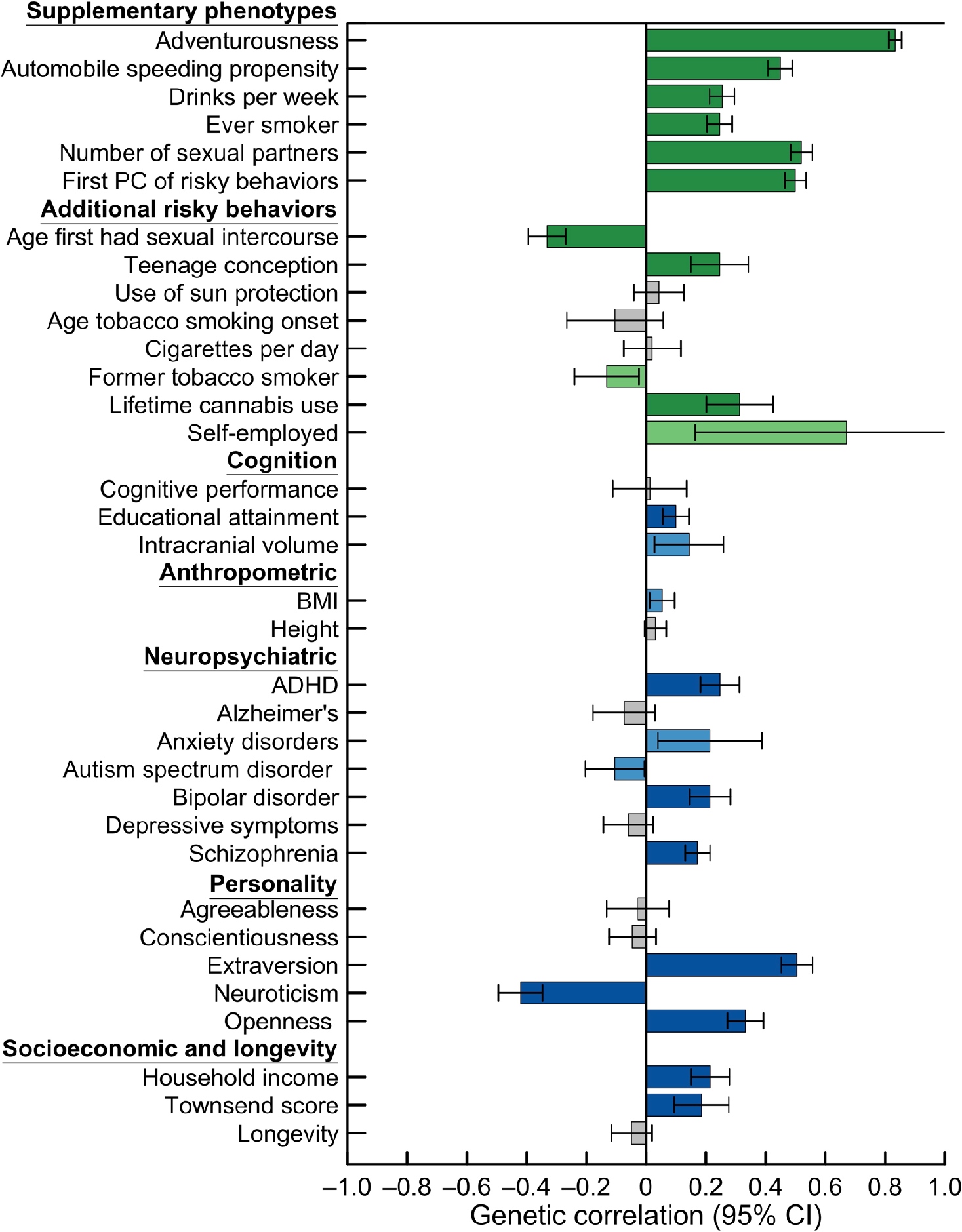
Genetic correlations with general risk tolerance. The genetic correlations were estimated using bivariate LD Score (LDSC) regression^9^. Error bars show 95% confidence intervals. For the supplementary phenotypes and the additional risky behaviors, green bars represent significant estimates with the expected signs, where higher risk tolerance is associated with riskier behavior. For the other phenotypes, blue bars represent significant estimates. Light green and light blue bars represent genetic correlations that are statistically significant at the 5% level, and dark green and dark blue bars represent correlations that are statistically significant after Bonferroni correction for 34 tests (the total number of phenotypes tested). Grey bars represent correlations that are not statistically significant at the 5% level.

Our estimates of the genetic correlations between general risk tolerance and the supplementary risky behaviors are substantially higher than the corresponding phenotypic correlations (**Supplementary Tables 1.3** and **7.1**). Although measurement error partly accounts for the low phenotypic correlations, the genetic correlations remain considerably higher even after adjustment of the phenotypic correlations for measurement error. The comparatively large genetic correlations support the view that a general factor of risk tolerance partly accounts for cross-domain variation in risky behavior^15,16^ and imply that this factor is genetically influenced. The lower phenotypic correlations suggest that environmental factors are more important contributors to domain-specific risky behavior^17,18^.

To increase the precision of our estimates of the SNPs’ effects on general risk tolerance, we leveraged the high degree of genetic overlap across our phenotypes by conducting Multi-Trait Analysis of GWAS (MTAG)^19^. We used as inputs the summary statistics of our GWAS of general risk tolerance, of our first five supplementary GWAS (i.e., not including the first PC of the four risky behaviors), and of a previously published GWAS on lifetime cannabis use^20^ (**Supplementary Information section 9**). MTAG increased the number of general-risk-tolerance lead SNPs from 124 to 312 (**Extended Data Fig. 9.1**, **Supplementary Table 9.1**).

We also estimated genetic correlations between general risk tolerance and 28 additional phenotypes (**Fig. 2** and in **Supplementary Table 7.1**). These included phenotypes for which we could obtain summary statistics from previous GWAS, as well as five phenotypes for which we conducted new GWAS. The estimated genetic correlations for the personality traits extraversion (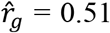, *SE* = 0.03), neuroticism (−0.42, *SE* = 0.04), and openness to experience (0.33, *SE* = 0.03) are substantially larger in magnitude than previously reported phenotypic correlations^21^, pointing to substantial shared genetic influences among general risk tolerance and these traits. After Bonferroni correction, we also find significant positive genetic correlations with the neuropsychiatric phenotypes ADHD, bipolar disorder, and schizophrenia. Viewed in light of the genetic correlations we find with risky behaviors classified as externalizing (e.g., substance use, elevated sexual behavior, and fast driving), these results suggest the hypothesis that the overlap with the neuropsychiatric phenotypes is driven by their externalizing component.

### Biological annotation

To gain insights into the biological mechanisms through which genetic variation influences general risk tolerance, we conducted a number of analyses. First, we systematically reviewed the literature that aimed to link risk tolerance to biological pathways (**Supplementary Information section 11**). Our review covered studies based on candidate genes (i.e., specific genetic variants used as proxies for biological pathways), pharmacological manipulations, biochemical assays, genetic manipulations in rodents, as well as other research designs. Our review identified 132 articles that matched our search criteria (**Supplementary Table 11.1**).

Previous work has focused on five main biological pathways: the steroid hormone cortisol, the monoamines dopamine and serotonin, and the steroid sex hormones estrogen and testosterone. Using a MAGMA^22^ competitive gene-set analysis, we found no evidence that SNPs within genes associated with these five pathways tend to be more associated with general risk tolerance than SNPs in other genes (**Supplementary Table 11.3**). Further, none of the other bioinformatics analyses we report below point to these pathways.

We also examined the 15 most commonly tested autosomal genes within the dopamine and serotonin pathways, which were the focus of most of the 34 candidate-gene studies identified by our literature review. We verified that the SNPs available in our GWAS results tag most of the genetic variants typically used to test the 15 genes. Across one SNP-based test and two gene-based tests, we found no evidence of non-negligible associations between those genes and general risk tolerance (**Fig. 1c** and **Supplementary Table 11.4**). (We note, however, that some brain regions identified in analyses we report below are areas where dopamine and serotonin play important roles.)

Second, we performed a MAGMA^22^ gene analysis to test each of ~18,000 protein-coding genes for association with general risk tolerance (**Supplementary Information section 12.2**). After Bonferroni correction, 285 genes were significant (**Extended Data Fig. 12.1** and **Supplementary Table 12.3**). To gain insight into the functions and expression patterns of these 285 genes, we looked up these genes in the Gene Network^23^ co-expression database. Third, to identify relevant biological pathways and identify tissues in which genes near general-risk-tolerance-associated SNPs are expressed, we applied the software tool DEPICT^24^ to the SNPs with *P* values less than 10^−5^ in our GWAS of general risk tolerance (**Supplementary Information section 12.4**).

Both the Gene Network and the DEPICT analyses separately point to a role for glutamate and GABA neurotransmitters, which are the main excitatory and inhibitory neurotransmitters in the brain, respectively^25^ (**Fig. 3a** and **Supplementary Tables 12.4** and **12.8**). To our knowledge, no published large-scale GWAS of cognition, personality, or neuropsychiatric phenotypes has pointed to clear roles both for glutamate and GABA (although glutamatergic neurotransmission has been implicated in recent GWAS of schizophrenia^26^ and major depression^27^). Our results suggest that the balance between excitatory and inhibitory neurotransmission may contribute to variation in general risk tolerance across individuals.

**Figure 3.**
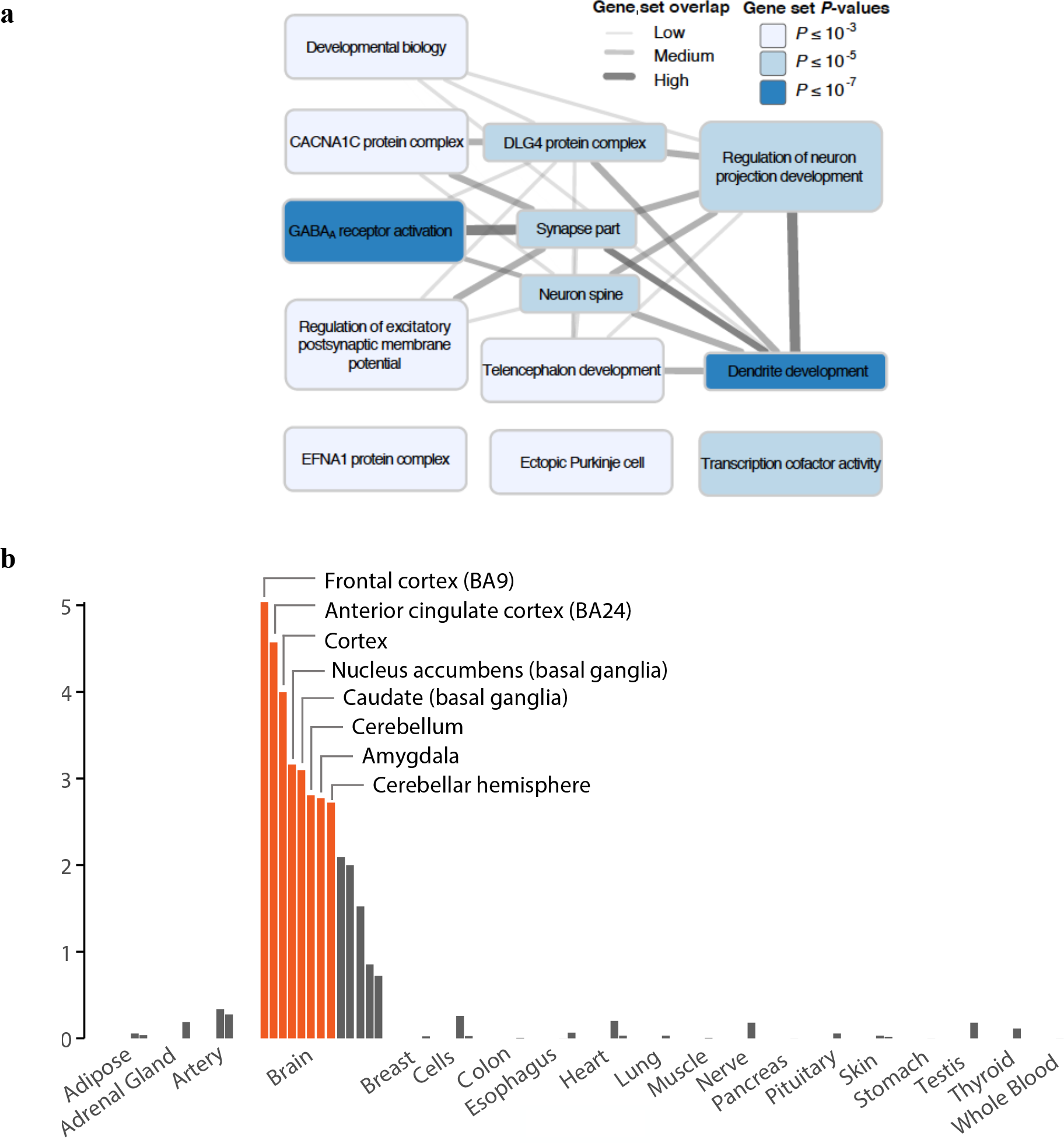
Results from selected biological analyses. **a**, DEPICT gene-set enrichment diagram. We identified 93 reconstituted gene sets that are significantly enriched (FDR < 0.01) for genes overlapping loci associated with general risk tolerance; using the Affinity Propagation method^36^, these were grouped into the 13 clusters displayed in the graph. Each cluster was named after the most significant gene set it contained, and each cluster’s color represents the permutation *P* value of its most significant gene set. The “synapse part” cluster includes the gene set “glutamate receptor activity,” and several members of the “GABA_A_ receptor activation” cluster are defined clusters is represented by an edge. Edge width represents the Pearson correlation *ρ* between the two respective vectors of gene membership scores (*ρ* < 0.3, no edge; 0.3 ≤ *ρ* < 0.5, thin edge; 0.5 ≤ *ρ* < 0.7, intermediate edge; *ρ* ≥ 0.7, thick edge). **b**, Results of DEPICT tissue enrichment analysis using GTEx data. The panel shows whether the genes overlapping loci associated with general risk tolerance are significantly overexpressed (relative to genes in random sets of loci matched by gene density) in various tissues. Tissues are grouped by organ or tissue type. The orange bars correspond to tissues with significant overexpression (FDR < 0.01). The *y*-axis is the significance on a −log_10_ scale.

The Gene Network and the DEPICT tissue enrichment analyses also both separately point to enrichment of the prefrontal cortex and the basal ganglia (**Fig. 3b** and **Supplementary Tables 12.4**, **12.6**, and **12.7**). The cortical and subcortical regions highlighted by DEPICT include some of the major components of the cortical-basal ganglia circuit, which is known as the reward system in human and non-human primates and is critically involved in learning, motivation, and decision-making, notably under risk and uncertainty^28,29^. We caution, however, that our results do not point exclusively to the reward system.

Lastly, we used stratified LD Score regression^30^ to test for the enrichment of SNPs associated with histone marks in 10 tissue or cell types (**Supplementary Information section 12.1**). Central nervous system tissues are the most enriched, accounting for 44% (*SE* = 3%) of the heritability while comprising only 15% of the SNPs (**Extended Data Fig. 12.3a** and **Supplementary Table 12.2**). Immune/hematopoietic tissues are also significantly enriched. While a role for the immune system in modulating risk tolerance is plausible given prior evidence of its involvement in several neuropsychiatric disorders^26,27^, future work is needed to confirm this result and to uncover specific pathways that might be involved.

### Polygenic prediction

We constructed polygenic scores of general risk tolerance to gauge their potential usefulness in empirical research (**Supplementary Information section 10**). We used the Add Health, HRS, NTR, STR, UKB-siblings, and Zurich cohorts as validation cohorts (**Supplementary Table 1.1** provides an overview of these cohorts; the UKB-siblings cohort comprised individuals with at least one full sibling in the UKB). For each validation cohort, we constructed the score using summary statistics from a meta-analysis of our discovery and replication GWAS that excluded the cohort. Our measure of predictive power is the incremental *R*^*2*^ (or pseudo-*R*^*2*^) from adding the score to a regression of the phenotype on sex, birth year, and the top ten principal components of the genetic relatedness matrix.

Our preferred score was constructed with LDpred^31^. In the UKB-siblings cohort, which is our largest validation cohort (*n* ~ 35,000), the score’s predictive power is 1.6% for general risk tolerance, 1.0% for the first PC of the four risky behaviors, 0.8% for number of sexual partners, 0. 6% for automobile speeding propensity, and ~0.15% for drinks per week and ever smoker. Across our validation cohorts, the score is also predictive of several personality phenotypes and a suite of real-world measures of risky behaviors in the health, financial, career, and other domains (**Extended Data Figs. 10.1-10.2** and **Supplementary Tables 10.1-10.3**). The incremental *R*^*2*^ we observe for general risk tolerance is consistent with the theoretical prediction, given the SNP heritability of general risk tolerance (**Table 1**) and the imperfect genetic correlations across the GWAS and validation cohorts^32,33^ (**Supplementary Information section 10.4**).

## Discussion

Our results provide insights into biological mechanisms that influence general risk tolerance. Our bioinformatics analyses point to the role of gene expression in brain regions that have been identified by neuroscientific studies on decision-making, notably the prefrontal cortex, basal ganglia, and midbrain, thereby providing convergent evidence with that from neuroscience^28,29^. Yet our analyses failed to find evidence for the main biological pathways that had been previously hypothesized to influence risk tolerance. Instead, our analyses implicate genes involved in glutamatergic and GABAergic neurotransmission, which were heretofore not generally believed to play a role in risk tolerance.

Although our focus has been on the genetics of general risk tolerance and risky behaviors, environmental and demographic factors account for a substantial share of these phenotypes’ variation. We observe sizeable effects of sex and age on general risk tolerance in the UKB data (**Extended Data Fig. 1.1**), and life experiences have been shown to affect both measured risk tolerance and risky behaviors (e.g., refs. 34,35). The data we have generated will allow researchers to construct and use polygenic scores of general risk tolerance to measure how environmental, demographic, and genetic factors interact with one another.

For the behavioral sciences, our results bear on the ongoing debate about the extent to which risk tolerance is a “domain-general” as opposed to a “domain-specific” trait. Low phenotypic correlations in risk tolerance across decision-making domains have been interpreted as supporting the domain-specific view^17,18^. Across the risky behaviors we study, we find that the genetic correlations are considerably higher than the phenotypic correlations (even after the latter are corrected for measurement error) and that many lead SNPs are shared across our phenotypes. These observations suggest that the low phenotypic correlations across domains are due to environmental factors that dilute the effects of a genetically-influenced domain-general factor of risk tolerance.

## Supporting information

Extended Data Items

## Acknowledgments

This research was carried out under the auspices of the Social Science Genetic Association Consortium (SSGAC). The research has also been conducted using the UK Biobank Resource under Application Number 11425. The study was supported by funding from the Ragnar Söderberg Foundation (E9/11 and E42/15), the Swedish Research Council (421-2013-1061), The Jan Wallander and Tom Hedelius Foundation, an ERC Consolidator Grant to Philipp Koellinger (647648 EdGe), the Pershing Square Fund of the Foundations of Human Behavior, and the NIA/NIH through grants P01-AG005842, P01-AG005842-20S2, P30-AG012810, and T32-AG000186-23 to NBER, and R01-AG042568-02 to the University of Southern California. We thank the International Cannabis Consortium, the Psychiatric Genomics Consortium, and the eQTLgen Consortium for sharing summary statistics from the GWAS of lifetime cannabis use, summary statistics from the GWAS of ADHD, and eQTL summary statistics, respectively. A full list of acknowledgments is provided in **Supplementary Information section 13**.

## Author Contributions

A full list of author contributions is included in **Supplementary Information section 13**.

## Author Information

Adam Auton, Pierre Fontanillas, David A Hinds, and Aaron Kleinman are employees of 23andMe, and Ronald Kessler has had ties to various companies in the past three years; the authors declare no other competing financial interests. Further details are provided in **Supplementary Information section 13**. Correspondence and requests for materials should be addressed to J.P.B, R.K.L.

## Data Availability

Upon publication, results can be downloaded from the SSGAC website (http://thessgac.org/data).

## Footnotes

1 Previous title: Genome-wide study identifies 611 loci associated with risk tolerance and risky behaviors.

## References

1. Dohmen, T. et al. Individual risk attitudes: Measurement, determinants, and behavioral consequences. J. Eur. Econ. Assoc. 9, 522–550 (2011).

2. Falk, A., Dohmen, T., Falk, A. & Huffman, D. The nature and predictive power of preferences: Global evidence. IZA Discussion Papers (2015).

3. Beauchamp, J. P., Cesarini, D. & Johannesson, M. The psychometric and empirical properties of measures of risk preferences. J. Risk Uncertain. 54, 203–237 (2017).

4. Cesarini, D., Dawes, C. T., Johannesson, M., Lichtenstein, P. & Wallace, B. Genetic variation in preferences for giving and risk taking. Q. J. Econ. 124, 809–842 (2009).

5. Harden, K. P. et al. Beyond dual systems: A genetically-informed, latent factor model of behavioral and self-report measures related to adolescent risk-taking. Dev. Cogn. Neurosci. 25, 221–234 (2017).

6. Hewitt, J. K. Editorial policy on candidate gene association and candidate gene-by-environment interaction studies of complex traits. Behav. Genet. 42, 1–2 (2012).

7. Day, F. R. et al. Physical and neurobehavioral determinants of reproductive onset and success. Nat. Genet. 48, 617–623 (2016).

8. Strawbridge, R. J. et al. Genome-wide analysis of risk-taking behaviour and cross-disorder genetic correlations in 116 255 individuals from the UK Biobank cohort. bioRxiv (2017). doi:http://dx.doi.org/10.1101/177014

9. Bulik-Sullivan, B. K. et al. An atlas of genetic correlations across human diseases and traits. Nat. Genet. 47, 1236–1241 (2015).

10. Okbay, A. et al. Genetic variants associated with subjective well-being, depressive symptoms, and neuroticism identified through genome-wide analyses. Nat. Genet. 48, 624–633 (2016).

11. Bulik-Sullivan, B. K. et al. LD Score regression distinguishes confounding from polygenicity in genome-wide association studies. Nat. Genet. 47, 291–295 (2015).

12. Price, A. L. et al. Long-range LD can confound genome scans in admixed populations. Am. J. Hum. Genet. 83, 132–139 (2008).

13. Berisa, T. & Pickrell, J. K. Approximately independent linkage disequilibrium blocks in human populations. Bioinformatics 32, 283–285 (2016).

14. Welter, D. et al. The NHGRI GWAS Catalog, a curated resource of SNP-trait associations. Nucleic Acids Res. 42, D1001–1006 (2014).

15. Einav, B. L., Finkelstein, A., Pascu, I. & Cullen, M. R. How general are risk preferences? Choices under uncertainty in different domains. Am. Econ. Rev. 102, 2606–2638 (2016).

16. Frey, R., Pedroni, A., Mata, R., Rieskamp, J. & Hertwig, R. Risk preference shares the psychometric structure of major psychological traits. Sci. Adv. 3, e1701381 (2017).

17. Weber, E. U., Blais, A. E. & Betz, N. E. A domain-specific risk-attitude scale: Measuring risk perceptions and risk behaviors. J. Behav. Decis. Mak. J. Behav. Dec. Mak. 15, 263–290 (2002).

18. Hanoch, Y., Johnson, J. G. & Wilke, A. Domain specificity in experimental measures and participant recruitment: an application to risk-taking behavior. Psychol. Sci. 17, 300–304 (2006).

19. Turley, P. et al. MTAG: Multi-Trait Analysis of GWAS. bioRxiv (2017). doi:https://doi.org/10.1101/118810

20. Stringer, S. et al. Genome-wide association study of lifetime cannabis use based on a large meta-analytic sample of 32 330 subjects from the International Cannabis Consortium. Transl. Psychiatry 6, e769 (2016).

21. Becker, A., Deckers, T., Dohmen, T., Falk, A. & Kosse, F. The relationship between economic preferences and psychological personality measures. Annu. Rev. Econom. 4, 453–478 (2012).

22. de Leeuw, C. A., Mooij, J. M., Heskes, T. & Posthuma, D. MAGMA: Generalized gene-set analysis of GWAS data. PLoS Comput. Biol. 11, 1–19 (2015).

23. Fehrmann, R. S. N. et al. Gene expression analysis identifies global gene dosage sensitivity in cancer. Nat. Genet. 47, 115–125 (2015).

24. Pers, T. H. et al. Biological interpretation of genome-wide association studies using predicted gene functions. Nat. Commun. 6, 5890 (2015).

25. Petroff, O. A. C. GABA and glutamate in the human brain. Neurosci. 8, 562–573 (2002).

26. Ripke, S. et al. Biological insights from 108 schizophrenia-associated genetic loci. Nature 511, 421–427 (2014).

27. Wray, N. R. et al. Genome-wide association analyses identify 44 risk variants and refine the genetic architecture of major depression. bioRxiv (2017). doi:https://doi.org/10.1101/167577

28. Haber, S. N. & Knutson, B. The reward circuit: linking primate anatomy and human imaging. Neuropsychopharmacology 35, 4–26 (2010).

29. Tobler, P. & Weber, E. U. Valuation for Risky and Uncertain Choices. in Neuroeconomics 149–172 (Elsevier, 2014). doi:10.1016/B978-0-12-416008-8.00009-7

30. Finucane, H. K. et al. Partitioning heritability by functional annotation using genome-wide association summary statistics. Nat. Genet. 47, 1228–1235 (2015).

31. Vilhjálmsson, B. J. et al. Modeling linkage disequilibrium increases accuracy of polygenic risk scores. Am. J. Hum. Genet. 97, 576–592 (2015).

32. Daetwyler, H. D., Villanueva, B. & Woolliams, J. A. Accuracy of predicting the genetic risk of disease using a genome-wide approach. PLoS One 3, e3395 (2008).

33. de Vlaming, R. et al. Meta-GWAS Accuracy and Power (MetaGAP) calculator shows that hiding heritability is partially due to imperfect genetic correlations across studies. PLoS Genet. 13, e1006495 (2017).

34. Sahm, C. R. How much does risk tolerance change? Q. J. Financ. 2, 1250020 (2012).

35. Malmendier, U. & Nagel, S. Depression babies: Do macroeconomic experiences affect risk taking? Q. J. Econ. 126, 373–416 (2011).

36. Frey, B. J. & Dueck, D. Clustering by passing messages between data points. Science 315, (2007).

37. Furberg, H. et al. Genome-wide meta-analyses identify multiple loci associated with smoking behavior. Nat. Genet. 42, 441–447 (2010).

38. Shi, H., Kichaev, G. & Pasaniuc, B. Contrasting the genetic architecture of 30 complex traits from summary association data. Am. J. Hum. Genet. 99, 139–153 (2016).

