## Supplementary Information for "Genome-wide association analyses of risk tolerance and risky behaviors in over 1 million individuals identify hundreds of loci and shared genetic influences^1^"

[a] Previous title: Genome-wide study identifies 611 loci associated with risk tolerance and risky behaviors.

7.4 OTHER RESULTS ... 62
7.5 COMPARISON OF THE GENETIC AND PHENOTYPIC CORRELATIONS ... 62
7.6 APPENDIX: GWAS OF OTHER RISKY BEHAVIORS AND OF SOCIOECONOMIC PHENOTYPES USING THE FIRST RELEASE OF UKB DATA ... 64

**8 PROXY-PHENOTYPE ANALYSES ... 66**

8.1 INTRODUCTION ... 66
8.2 METHODOLOGY ... 66
8.3 RESULTS FROM PROXY-PHENOTYPE ENRICHMENT ANALYSES ... 68

**9 MULTI-TRAIT ANALYSIS OF GWAS (MTAG) ... 72**

9.1 INTRODUCTION ... 72
9.2 METHODS ... 72
9.3 RESULTS ... 73

**10 PREDICTIVE POWER OF GENERAL-RISK-TOLERANCE POLYGENIC SCORE 75**

10.1 PHENOTYPE DEFINITION ... 76
10.2 METHODS ... 86
10.3 RESULTS ... 90
10.4 EXPECTED PREDICTIVE POWER OF GENERAL-RISK-TOLERANCE POLYGENIC SCORE ... 94
10.5 APPENDIX: DEFINITION OF THE INCOME RISK GAMBLE VARIABLE IN THE HRS COHORT AND MERGER OF THE STR1 AND STR2 COHORTS ... 96

**11 TESTING HYPOTHESES ABOUT SPECIFIC GENES AND GENE SETS ... 99**

11.1 LITERATURE REVIEW ... 99
11.2 GENE ANALYSIS AND COMPETITIVE GENE-SET ANALYSES WITH MAGMA ... 101
11.3 DISCUSSION ... 106

**12 BIOLOGICAL ANNOTATION ... 107**

12.1 FUNCTIONAL PARTITIONING OF HERITABILITY WITH STRATIFIED LD SCORE REGRESSION 108
12.2 IDENTIFYING GENES ASSOCIATED WITH GENERAL RISK TOLERANCE WITH MAGMA ... 112
12.3 IDENTIFYING GENES ASSOCIATED WITH GENERAL RISK TOLERANCE WITH SMR ... 115
12.4 PRIORITIZATION OF TISSUES, GENE SETS, AND GENES USING DEPICT ... 119
12.5 COMPETITIVE GENE-SET ANALYSIS WITH MAGMA: TESTING GENE SETS RELATED TO GABA AND GLUTAMATE NEUROTRANSMITTERS ... 122
12.6 LOOKUP OF PROTEIN-ALTERING VARIANTS AND *CIS*-EQTLS ... 123
12.7 SUMMARY OF MAIN FINDINGS FROM THE BIOLOGICAL ANNOTATION ANALYSES ... 126

**13 ADDITIONAL INFORMATION ... 131**

13.1 AUTHOR CONTRIBUTIONS ... 131
13.2 COHORT-LEVEL CONTRIBUTIONS ... 132
13.3 ADDITIONAL ACKNOWLEDGMENTS ... 134

**14 REFERENCES ... 138**

### 1 Study overview

#### 1.1 Study motivation

Risk tolerance, or the willingness to take risks to obtain rewards, is a fundamental parameter in economics, finance, and behavioral decision theory. Different measures of risk preferences have previously been linked to real-world behaviors such as portfolio allocation and occupational choice, as well as health behaviors such as smoking, exercising, and alcohol and drug use[1–5]. Further, recent work has demonstrated the existence of a reliable general factor of risk preference that is generalizable both to specific and real-world risky behaviors[6].

Elucidating the determinants of individual differences in general risk tolerance is an active field of research[3,4]. General risk tolerance has been found to be moderately heritable in twin studies ($h^2 \sim 30\%$), although heritability estimates in the literature vary, ranging from 20% to 60%[2,7–9]; an earlier study based on molecular genetic data had confirmed the heritability of the trait[10]. Some previous studies have attempted to identify specific genetic variants that are associated with general risk tolerance. However, most of these attempts have been conducted in relatively small samples with a few hundred to at most a few thousand individuals[11–14]; see **Supplementary Table 11.1**. Given that the effect of any specific single nucleotide polymorphism (SNP) on a genetically complex trait like general risk tolerance is likely to be extremely small[10,15–18], these earlier studies were most likely underpowered. Low statistical power not only implies a low probability to detect true effects, but also a high chance of finding false positives, a strong tendency to overestimate the effects of statistically significant variables, and a high likelihood that significant findings will have the wrong sign[19]. Accordingly, the replication record of these underpowered studies has been disappointing[20,21].

The purpose of this study is to investigate the molecular genetic architecture of general risk tolerance, adventurousness, and of a number of risky behaviors and to identify specific genetic variants associated with the phenotypes in well-powered genome-wide association studies (GWAS). Our findings could help elucidate the genetic and biological mechanisms that underlie individual variation in the willingness to avoid or engage in risky behavior.

#### 1.2 Phenotype definitions

##### *1.2.1 General risk tolerance*

Our main phenotype is self-reported "general risk tolerance," defined as one's tendency or willingness to take risks in general. For our discovery stage, we combine data from the UK Biobank and from the 23andMe cohort.

We use the following survey question from the UK Biobank ($n = 431{,}126$):

*"Would you describe yourself as someone who takes risks? Yes / No."*

Throughout the study all risk-related phenotypes are coded so that a higher risk tolerance is associated with a higher phenotype value. For example, in the UKB "yes" is coded as 1 and "no" as 0. The majority of the 431,126 respondents in the UKB were only assessed once, while a subset of 18,102 individuals answered the survey question a second time. 15,618 of these re-assessed individuals gave consistent answers in both waves, while 2,484 changed their responses. When the answer between the two assessments changed, we took the average response (i.e., 0.5) as the individual's measure of risk tolerance. Consistent with prior research[22,23], a much higher fraction of males (33%) than females (18%) in the UKB cohort described themselves as risk tolerant on the general-risk-tolerance measure ($P < 1 \times 10^{-100}$, *t*-test; **Extended Data Fig. 1.1**).

In the 23andMe cohort, our main phenotype is again self-reported "general risk tolerance." We use the survey question (*n* = 508,782):

*"In general, people often face risks when making financial, career, or other life decisions. Overall, do you feel comfortable or uncomfortable taking risks? [1] Very comfortable / [2] Somewhat comfortable / [3] Neither comfortable nor uncomfortable / [4] Somewhat uncomfortable / [5] Very uncomfortable."*

We reverse this coding for our GWAS, so that "1" is coded as the least risk-related response category and "5" as the most risk-related response category.

In total, then, our full general risk tolerance discovery meta-analysis includes 939,908 people from the UK Biobank and from the 23andMe cohort.

For our replication stage (*n* = 35,445), we combined 10 independent cohorts from seven studies with survey questions on general risk tolerance, all of which included a question similar to the one asked in the UKB or in Dohmen *et al.*[4]:

*"How do you see yourself: are you generally a person who is fully prepared to take risks or do you try to avoid taking risks? Please tick a box on the scale, where the value 0 means: 'not at all willing to take risks' and the value 10 means: 'very willing to take risks'."*

In **Supplementary Table 1.1** we report an overview of the cohorts included in our study. **Supplementary Table 1.2** lists the detailed general-risk-tolerance survey questions available in each cohort. The UKB is the only cohort that asks the general-risk-tolerance question in a binary fashion; the 23andMe cohort asked this question on a 5-point Likert scale, and all seven replication cohorts asked their participants this question on a 10- or 11-point Likert scale. All of the replication cohorts are cross-sectional, and only the UKB includes more than one measurement over time for some individuals.

As we further describe in **Supplementary Information sections 3.3** and **7.4**, we used bivariate LD Score Regression[24] to estimate the genetic correlation between: (1) the UK Biobank risk-tolerance GWAS and the 23andMe risk-tolerance GWAS; (2) the UK Biobank risk-tolerance GWAS and the replication GWAS; (3) the 23andMe risk-tolerance GWAS and the replication GWAS; and (4) the full discovery GWAS of general risk tolerance (UK Biobank + 23andMe) and the replication GWAS. For the last three correlations, we see significant, moderately high, and positive genetic correlations between 0.75 and 0.83 that are indistinguishable from unity. However, for (1), we find a genetic correlation that *is* distinguishable from unity ($\hat{r}_g$= 0.767, *SE* = 0.021). Though this genetic correlation is lower than expected, it is high enough to justify the meta-analysis of the UK Biobank and 23andMe summary statistics for general risk tolerance that we perform as our discovery GWAS[18]. However, the imperfect correlation points to some degree of heterogeneity across the cohorts, which may attenuate the genetic signals we can observe in our study.

We note that there are various alternative ways to measure risk tolerance, including behavioral experiments with real stakes[25], hypothetical choices[1], and survey questions[25]. The phenotypic correlation between these measures is typically moderate[2,4,26], although correcting for measurement error substantially increases these correlations[2,26].

Our approach of using survey measures of general risk tolerance has several advantages. First, these measures have been shown to correlate with a wide range of risky behaviors such as investments in stocks, active sports, self-employment, and smoking, even after controlling for age, sex, education, wealth, and income[1,2,4,26]. Thus, these survey measures capture an important dimension of individual differences. There is evidence that measures of general risk tolerance are good all-around predictors of risky behavior and perform better in this respect than more specific survey measures of risk tolerance[4]. Second, survey questions of general risk tolerance are cheap and easy to collect and have been added to numerous questionnaires, including the UKB, thus allowing us to reach a large enough sample size to conduct a well-powered GWAS.

Although general risk tolerance may be closely related to some psychological measures of personality such as extraversion, novelty seeking, or sensation seeking, these constructs are not identical. For this reason, personality measures were not included in our main GWAS of general risk tolerance, but below we study the predictive power of a polygenic score of general risk tolerance for several of these psychological measures of personality, as well as the genetic correlation between general risk tolerance and some of these measures.

We also performed supplementary GWAS for adventurousness, four risky behaviors in the UK Biobank (automobile speeding propensity, drinks per week, ever smoker, and number of sexual partners), and the first principal component of these four risky behaviors. Moving forward, we refer to the main GWAS of general risk tolerance as our "primary GWAS" and the GWAS of these additional six phenotypes as our "supplementary GWAS."

##### *1.2.2 Adventurousness*

We also performed a GWAS of adventurousness, since this phenotype is known to be related to risk-taking behavior[27]. For this GWAS, we use only summary statistics from the 23andMe cohort, and we use the following survey question ($n$ = 557,923):

*"If forced to choose, would you consider yourself to be more cautious or more adventurous? [1] Very cautious / [2] Somewhat cautious / [3] Neither / [4] Somewhat adventurous / [5] Very adventurous."*

We maintain this coding, where "1" is coded as the least risk-related response category and "5" as the most risk-related response category.

##### *1.2.3 Other supplementary UKB Risky Behaviors*

We also conducted GWAS studies for other risky behaviors in the UKB. Our selection strategy for these phenotypes was twofold. First, we chose a set of potential GWAS phenotypes by searching the UKB database for risky behaviors. The risky behaviors we originally considered included automobile speeding propensity, use of sun protection, age of first sexual intercourse, number of lifetime sexual partners, teenage conception (females only), as well as whether an individual was ever a tobacco smoker, whether an individual is a former tobacco smoker, age of tobacco smoking onset, number of cigarettes per day. The inclusion of these risky behaviors as additional GWAS phenotypes builds on previous economics studies that demonstrate the predictive power of risk tolerance for various behaviors such as smoking, drinking, portfolio allocation, and occupational choice[1–5,28].

Finally, we considered whether an individual was ever an alcohol drinker, whether an individual is a current alcohol drinker, whether an individual is an excessive alcohol drinker, and number of drinks per day. Ultimately, we decided not to retain the use of sun protection in our main analyses, because there was no significant genetic correlation between this phenotype and the preliminary results from our GWAS of general risk tolerance using the first release of UKB data ($\hat{r}_g = 0.025$, $P$ value = 0.701, as estimated using bivariate LD Score regression).

Second, from the remaining phenotypes, we selected just one phenotype in each domain of risky behavior (i.e., driving behavior, drinking behavior, smoking behavior, and sexual behavior). To do so, we prioritized: 1) phenotypes available in the entire UKB sample, since our general-risk-tolerance phenotype is defined for everyone in the UKB sample; 2) phenotypes that had been previously explored in other published GWAS; and 3) phenotypes which showed a high phenotypic correlation with our main general-risk-tolerance phenotype in the first release of the UKB data (this is the data we had access to when deciding which phenotypes to select). If not reported in the text below, these correlations from the first release of the UKB are available upon request, although the differences in the correlations between the first and full release are small. For a list of the phenotypic correlations between the selected phenotypes in the full release of the UKB, see **Supplementary Table 1.3**. Below we highlight how the phenotypes in each domain of risky behavior were selected and are defined.

*Automobile speeding propensity*: Automobile speeding propensity is the only phenotype available that measures risky driving behavior in the UKB; its phenotypic correlation with general risk tolerance in the first release of the UKB is 0.164. Respondents were asked, "How often do you drive faster than the speed limit on the motorway?" Response options include: 1) Never/rarely; 2) Sometimes; 3) Often; 4) Most of the time; and 5) Do not drive on the motorway. We first dropped all participants who reported not driving on the motorway, and then we normalized[a] our categorical variable for males and females separately. In total, this GWAS includes 404,291 individuals in the UK Biobank.

*Drinks per week*: There are several phenotype options that measure drinking behavior in the UKB. After considering only phenotypes that cover the entire UKB sample, we were left with two: drinks per week and excessive alcohol drinking. Our drinks per week measure is constructed from responses to a sequence of questions. First, respondents were asked how often they drink alcohol, and response options include 1) daily or almost daily; 2) three or four times per week; 3) once or twice per week; 4) one to three times per month; 5) special occasions only; and 6) never. Respondents who reported drinking once per week or more were asked how many glasses of various types of alcoholic beverages they consume per week. We used the sum of all alcoholic drinks per week as our drinks per week phenotype for these respondents. Respondents who reported drinking less than once per week (one to three times per month or on special occasions only) were asked how many glasses of various types of alcoholic beverages they consume per *month*. For these respondents, we added the total number of drinks per month and

[a] The normalized variable is the inverse normal cumulative distribution function (CDF) of the observations' percentile ranks.

divided by 4 to arrive at an approximated number of drinks per week. Respondents who reported never drinking were coded as 0.

For the excessive alcohol drinking phenotype, we use current UK Chief Medical Officer drinking guidelines[b] to code respondents who drink 14 or fewer drinks per week as 0 and more than 14 drinks per week as 1. Our drinks per week phenotype had a higher phenotypic correlation with general risk tolerance in the first release of the UKB (0.139) than did our excessive alcohol drinking phenotype (0.074); further, drinks per week is a consistent phenotype studied in the alcohol GWAS literature. We therefore use this drinks per week phenotype as our final drinking behavior GWAS phenotype. In total, this GWAS includes 414,343 individuals in the UK Biobank.

*Ever smoker*: There are several potential measures of smoking behavior in the UKB. These include: 1) ever-tobacco smoker status; 2) former tobacco smoker status (among ever-tobacco smokers); 3) age of tobacco smoking onset (among ever-tobacco smokers); and 4) number of cigarettes per day. Because former tobacco smoker status and age of tobacco smoking onset were measured only for individuals who had ever been tobacco smokers, they are not defined for the entire UKB sample, and we thus did not select these phenotypes.

For our remaining possibilities, we coded ever-tobacco smoker status as 1 if a respondent reported that they were a current or previous smoker and 0 if they reported never smoking or only smoking once or twice. We coded cigarettes per day as 0 if ever-smoking status was also 0; otherwise, we used the maximum number of reported past or current cigarettes (or pipes/cigars) consumed per day, normalized separately for males and females. Our cigarettes per day phenotype had a slightly higher phenotypic correlation with general risk tolerance in the first release of the UKB (0.098) than ever-smoker status (0.092). However, we concluded that the consistency of the ever smoker phenotype with previous GWAS literature overrides this slightly higher phenotypic correlation, and we therefore use this ever smoker phenotype as our final smoking behavior GWAS phenotype.

For our GWAS of ever smoker, we meta-analyzed the summary statistics from the UKB GWAS with those from the Tobacco, Alcohol and Genetics (TAG) Consortium[29] (the TAG consortium refers to the ever smoker phenotype as "smoking initiation"). In total, this meta-analysis includes 518,633 people from the meta-analyzed UK Biobank (444,598) and TAG (74,035) summary statistics.

*Number of sexual partners*: The potential phenotypes for assessing risky sexual behavior in the UKB include teenage conception (which we coded ourselves from available phenotypes related to age and pregnancy), age of first sexual intercourse, and lifetime number of sexual partners. Because we only defined teenage conception for females, we did not pursue this phenotype. Lifetime number of sexual partners has a much higher phenotypic correlation with general risk tolerance in the first release of the UKB (0.207) than does age of first sexual intercourse. We therefore use lifetime number of sexual partners (hereafter referred to as simply "number of sexual partners") as our final risky sexual behavior GWAS phenotype. For this phenotype, respondents were asked, "About how many sexual partners have you had in your lifetime?" If respondents reported more than 99 lifetime sexual partners, they were asked to confirm their responses. We assigned a value of 0 to participants who reported having never had sex, and we

[b] https://www.drinkaware.co.uk/alcohol-facts/alcoholic-drinks-units/latest-uk-alcohol-unit-guidance/.

again normalized this measure separately for males and females. In total, this GWAS includes 370,711 individuals in the UK Biobank.

*First PC of risky behaviors:* We performed a principal component analysis (PCA) with our four selected risky behaviors above and obtained the first principal component (PC) (see **Supplementary Table 1.4**). The first PC is the linear combination of the four risky behaviors that has the largest possible variance (among all possible linear combinations where the squares of the weights on the four risky behaviors sum to one). It can be interpreted as a general factor of risky behavior. As we describe below, we performed a GWAS of this first PC of risky behaviors, and we also examined the genetic correlation between this PC and general risk tolerance (see **Supplementary Table 7.1**). In total, this GWAS includes 315, 894 people in the UK Biobank.

### 2 GWAS, quality control and meta-analysis

#### 2.1 Overview of primary GWAS

All analyses were performed at the cohort level according to a pre-specified and publicly archived analysis plan[30]. The original analysis plan was archived on February 4, 2016. For self-reported general risk tolerance, the analysis plan specified that the discovery GWAS would be conducted in the UKB and that the replication would be carried out in a meta-analysis of all other cohorts.

We updated the analysis plan on November 9, 2016 to include the analysis of the four risky behaviors and their first PC. For these phenotypes, the analysis plan specified that the GWAS would be conducted in the UKB. We did not attempt replication for these phenotypes. We updated the analysis plan a second time on July 10, 2017 to add the 23andMe cohort to the discovery GWAS of self-reported general risk tolerance. Two minor updates were made on August 7 and September 14, 2017 to specify that follow-up analyses (such as polygenic prediction) would be performed, whenever possible, on a meta-analysis combining the discovery and replication GWAS of general risk tolerance, and that we would add a GWAS of adventurousness alongside the primary GWAS of general risk tolerance and the other supplementary GWAS.

Cohorts other than the UKB could join this study by first supplying descriptive statistics and thereafter GWAS summary statistics from GWAS of self-reported general risk tolerance in November 2015 and December 2015, respectively. Two additional cohorts, Army STARRS and VHSS, joined the study later and provided descriptive and GWAS summary statistics in late 2016. Summary statistics were uploaded to a central, secure server and subsequently meta-analyzed. The lead PI of each cohort affirmed that the results contributed to the study were based on analyses approved by the local Research Ethics Committee and/or Institutional Review Board responsible for overseeing research. All participants provided written informed consent. We also obtained the descriptive and GWAS summary statistics from GWAS of self-reported general risk tolerance and adventurousness from 23andMe in late July 2017. An overview of the participating cohorts is reported in **Supplementary Table 1.1**.

The analysis plan instructed all cohorts to limit the analysis to individuals of European ancestry, to exclude individuals with missing covariates, to remove samples that displayed a SNP call rate of less than 95%, and to apply cohort-specific standard quality control filters before imputation. The cohort-specific standard quality control filters are reported in **Supplementary Table 2.1**. GWAS were limited to the 22 autosomes. The cohorts were required to provide unfiltered GWAS summary statistics including the following information for each SNP: chromosome and base-pair position, rsID, effect-coded allele, other allele, sample size per SNP, coefficient estimate (beta), standard error of the coefficient estimate, *P* value of the association uncorrected for genomic control, effect-coded allele frequency (EAF), imputation status, imputation quality, and Hardy-Weinberg equilibrium exact test *P* value for directly genotyped markers.

The analysis plan included power calculations assuming that 100,000 individuals in the UKB answered “Yes” (“cases”) to the general-risk-tolerance question, and 270,000 individuals answered “No” (“controls”). Under this assumption, our study would have 73% power to detect single nucleotide polymorphisms (SNPs) with a minor allele frequency (MAF) of 0.3 and an odds ratio of 1.05 with a genome-wide significance threshold of $P = 5\times10^{-8}$.

For general risk tolerance, the final sample size for the "discovery GWAS" meta-analysis of the UKB and 23andMe cohorts was 939,908 individuals. Replication was performed in a meta-analysis of 10 independent cohorts from seven studies totaling 35,445 individuals. We will henceforth refer to this meta-analysis of the 10 replication cohorts as the "replication GWAS." The follow-up analyses we describe in the following **Supplementary Information sections** were performed with GWAS summary statistics from a meta-analysis combining the discovery and replication GWAS ($n$ = 975,353), except where otherwise noted.

For adventurousness, GWAS summary statistics from 23andMe were analyzed ($n$ = 557,923).

For three of the four risky behaviors, namely automobile speeding propensity ($n$ = 404,291), drinks per week ($n$ = 414,343), and number of sexual partners ($n$ = 370,711), and for the first PC of the four risky behaviors ($n$ = 315,894), GWAS were conducted in the UKB only, as specified in the updated analysis plan. For the remaining risky behavior, ever smoker, we meta-analyzed the summary statistics from the UKB GWAS ($n$ = 444,598) with those from the Tobacco, Alcohol and Genetics (TAG) Consortium[29] ($n$ = 74,035), leading to a total sample size of 518,633 (the TAG consortium refers to the ever smoker phenotype as "smoking initiation").

#### 2.2 Genotyping and imputation

Genotyping[b] was performed using a range of common, commercially available genotyping arrays. An overview of the genotyping and imputation procedure is provided in **Supplementary Table 2.1**. The participating cohorts were encouraged to use their standard quality control protocols before imputation, as long as the applied filters satisfied the minimum requirements specified in the analysis plan (SNP call rate > 95%, HWE exact test $P$ value > $10^{-6}$, MAF > 1%). For the UKB, different filters were used, following ref.[31].

The cohorts, except for 23andMe, Army STARRS, BASE-II, UKB, and VHSS imputed markers using the 1000 Genomes phase 1 reference panel (March 2012 release version 3). 23andMe used the 1000 Genomes phase 1 (September 2013 haplotype release). Army STARSS used the 1000 Genomes phase 1 (August 2012 haplotype release). BASE-II used the more recent reference panel 1000 Genomes phase 3 (October 2014 haplotype release version 5). UKB used a customized reference panel based on the Haplotype Reference Consortium release 1.1 combined with the UK10K haplotype reference panel[32]. VHSS used the Haplotype Reference Consortium release 1.1[33].

All genetic positions reported in this study are denoted with those of the Genome Reference Consortium's human assembly 37 (GRCh37, sometimes referred to as the National Center for Biotechnology Information hg19).

#### 2.3 Association analyses

Cohorts were encouraged to exclude individuals with SNP call rates less than 95%, with excessive autosomal heterozygosity, and with sex mismatch. Family-based cohorts were informed to control for family structure either with mixed linear modeling or with a procedure

[b] The UKB genotype data was handled with QCtool, available at http://www.well.ox.ac.uk/~gav/qctool/#overview

selecting only one individual in each pair that displayed relatedness greater than 5% in a genetic relatedness matrix.

The genome-wide association analysis performed in each cohort estimated the following regression for each SNP, as in Okbay *et al.* 2016[16]:

(1)
$$Y_i = \beta_0 + \beta_1 SNP_i + \boldsymbol{PC_i}\boldsymbol{\gamma} + \boldsymbol{X_i}\boldsymbol{\alpha} + \boldsymbol{C_i}\boldsymbol{\theta} + \epsilon_i,$$

where $Y_i$ is the phenotype for individual $i$, $SNP_i$ is the number of effect-coded alleles of the SNP[c], $\boldsymbol{PC_i}$ is a vector of principal components of the genetic relatedness matrix after application of the pre-imputation filters described above, and $\boldsymbol{X_i}$ is a vector of control variables. The control variables include sex, birth year, birth year squared, birth year cubed, and the interactions between sex and the three birth-year variables. $\boldsymbol{C_i}$ is a vector containing cohort-specific controls and technical covariates (such as dummy variables for genotyping array and genotyping batches) that were recommended in the analysis plan. All associations were performed with males and females pooled. A summary of the GWAS association models and control variables for each cohort is reported in **Supplementary Table 2.2**.

In practice, the phenotype was often residualized by first regressing the phenotype on the control variables, and the residualized phenotype was then regressed on the genotypes. This approach leads to almost identical results as estimating the full regression model directly while drastically reducing the computation time needed for the GWAS.

##### *2.3.1 Linear mixed models in the UKB*

The association analyses in the UKB were performed with linear mixed models (LMM) with the BOLT-LMM v2.2 software[34]. The benefit of LMM is that the method accounts for cryptic relatedness and population structure, which allows the inclusion of related individuals in the sample, thereby yielding a larger sample size and greater statistical power. Using LMM is computationally intensive, and the BOLT algorithm is a new method that makes LMM analysis of hundreds of thousands of individuals computationally feasible. The method requires a set of SNPs to be included in the genetic variance component, and we included 483,680 directly genotyped bi-allelic autosomal SNPs with MAF > 0.005 and HWE *P* value > $10^{-6}$. We included individuals based on self-reported ancestry, specifically those who self-reported to be of "white" ancestry (i.e., self-reported white, British, Irish, or any other white background). In addition, we limited the GWAS to individuals for whom the value of the first principal component of the genetic relatedness matrix was less than "0," which identifies the cluster of individuals of European ancestry. We dropped individuals whose reported sex did not match their genetic sex, individuals with putative sex chromosome aneuploidy, individuals that did not pass the UKB internal genotype quality control, and individuals with missing values.

#### 2.4 Main reference panel

The full release of the UK Biobank genetic data was imputed with haplotypes from the Haplotype Reference Consortium v1.1 (HRC) and the UK10K haplotype reference panel[32]. A recommendation was communicated soon after the release of the genotype data in July 2017. It

---

[c] For imputed SNPs we used best-guess data for samples that were imputed with IMPUTE2[42], and dosage data for samples that were imputed with MaCH/Minimac[243].

was recommended that only SNPs available in the HRC be used for analysis, because a subset of variants imputed from the UK10K reference panel have wrongfully imputed genomic positions, while none of the HRC SNPs are affected. We therefore used the HRC v.1.1 as the reference panel for quality control of the GWAS summary statistics, and to determine the independence of significant loci. The following section describes our quality control of the HRC whole-genome sequence data (WGS) when constructing the reference panel. We will hereafter refer to the resulting reference panel as the "main reference panel."

##### *2.4.1 Quality-control of the main reference panel*

The HRC haplotypes were downloaded from the European Genome-phenome Archive (EGA) on August 1, 2017. Strict internal quality control had already been applied to the WGS data, as described in-depth elsewhere[33], and we restricted the main reference panel to variants that passed all pre-applied genotype call filters (i.e., variants whose VCF FILTER status is "PASS"; these pre-applied filters included, among other filters, a filter to remove variants with minor allele count (MAC) $\leq$ 5.) Before our internal QC the WGS data contained 40,405,506 autosomal and X-chromosome SNPs. The following protocol was restricted to the 39,131,579 autosomal SNPs because the pre-registered analysis plan restricts our analyses to the autosomes. The HRC reference panel does not include any structural variants, such as INDELs[33].

We performed a series of best-practice alignments of the WGS data for consistent and unique identification of variants[35] with the open-source software BCFtools[d] created by the Wellcome Trust Sanger Institute. Because PLINK cannot properly handle truly multi-allelic variants, we split multi-allelic variants into multiple bi-allelic variants. We then confirmed that all reference alleles and genomic positions matched the Genome Reference Consortium Human genome build 37 (GRCh37)[36]. To avoid issues with chromosomal positions mapping to multiple NCBI marker IDs (rsIDs), all rsIDs were removed, and all variants were given a unique identifier (henceforth referred to the as the "unique ID") in the form of chromosome, base-pair position, reference allele, and alternative allele, separated by colons (e.g., 1:123456:C:T). Using this format for variant IDs, together with the alignment to the reference genome GRCh37, ensures a unique representation of all SNPs and a lack of duplicate variants with switched reference alleles (e.g., 1:123456:C:T and 1:123456:T:C).

To avoid possible issues with inconsistencies with the UK10K haplotype reference in future work that may use that reference, we investigated possible strand and allele issues across the reference panels. By comparing the reference alleles and allele frequencies we found 24,394 loci with inconsistent alleles, and we decided to drop these from the main reference panel so that they would be removed during QC of the GWAS summary statistics.

We converted the VCF data to PLINK binary format with PLINK v.1.9b3.46[37], and we removed all multi-allelic loci (without retaining any of the multi-allelic variants coded as bi-allelic SNPs). Monomorphic SNPs (i.e., SNPs with MAF = 0) were kept in the reference panel as recommended[33]. We thereafter excluded one member of each pair of individuals with genomic relatedness greater than 0.025 from the sample, which removed 4,917 individuals of the 22,691 individuals for whom data were available for all autosomes in the VCF data.

[d] BCFtools can be downloaded here: http://samtools.github.io/bcftools/bcftools.html.

In summary, the main reference panel consists of 17,774 individuals and includes 38,889,224 bi-allelic autosomal SNPs that passed QC.

###### *2.4.2 rsID mapping*

Since rsIDs were removed from the SNPs to ensure unique identification, we created a map file so that each SNP could be assigned an rsID after meta-analysis, which is performed on the unique ID format described above (e.g., 1:123456:C:T). The map file contains the rsIDs from the HRC v1.1 sites list, and because we removed all multi-allelic variants there are no duplicate rsIDs.

#### 2.5 Quality control of allele-frequency differences between the UK Biobank genotyping arrays

The participants of the UKB were genotyped with two different but similar genotyping arrays: the UK BiLEVE Axiom array ($n$ ~ 50,000) and the *UK Biobank Axiom array* ($n$ ~ 450,000)[32]. It was communicated that the first release of UKB data contained a small set of 65 autosomal SNPs that appeared to have flipped reference alleles contingent on the array. A subset of these had unfortunately been used during the imputation procedure. We already control for genotype array and batch during GWAS analyses, but as an additional QC step beyond excluding the 65 previously reported flipped SNPs, we investigated the allele frequencies across the arrays to be sure that our results were unaffected by artifacts from the genotyping procedure. It should be noted that the participants genotyped with the UK BiLEVE array were chosen based on lung function and smoking behavior[38], but the sample is in all other respects comparable to the rest of the UK Biobank[39].

We restricted the imputed genotype data to unrelated individuals of British ancestry to ensure that allele-frequency differences across the genotyping arrays would not be caused by differences in the proportion of ancestries or be affected by dependent observations. With PLINK[37], we calculated the allele frequencies contingent on the genotyping array for both the directly genotyped and imputed SNPs. Because our quality-control protocol, described in the next section, restricts the GWAS to SNPs with MAF $\geq 0.001$, we chose not to investigate SNPs with MAF < 0.001 (in the imputed genotype data) for allele-frequency differences between the UKB genotyping arrays. SNPs available on only one of the genotyping arrays and SNPs that were not available in our main reference panel were not considered in this investigation of allele-frequency differences.

For each SNP included in the investigation, three quantities measuring differences in allele frequencies were examined: the absolute value of the difference between the two arrays and the absolute value of the difference between the main reference panel and each of the arrays. We flagged a SNP as problematic if it fulfilled the following two conditions: (i) if the absolute value of the difference between the two arrays was greater than 0.25; and (ii) if the absolute value of the difference between the main reference panel and at least one of the genotyping arrays was greater than 0.25. The comparison resulted in 600 flagged autosomal SNPs (including the 65 SNPs that were already reported as problematic) that were removed from the UKB summary statistics during QC in **Supplementary Information section 2.6.2.**

#### 2.6 Description of major steps in quality-control (QC) analyses

For each cohort, we applied a stringent quality-control protocol based on the EasyQC software (version 9.2) developed by the GIANT consortium[40], as well as additional steps developed by the Social Science Genetic Association Consortium[16,18]. All issues raised during implementation of the protocol described below were resolved through iterations between the meta-analyst and the cohort analysts before any GWAS summary statistics were forwarded for meta-analysis.

##### *2.6.1 Pre-QC verification and harmonization of GWAS summary statistics*

All cohorts were asked to supply descriptive statistics and phenotype definitions according to the pre-specified analysis plan[30]. The completeness of these documents was assessed as the first step of the quality control, together with examination of the uploaded GWAS summary statistics. All GWAS summary statistics were harmonized to ensure that the SNP identifier was in an admissible format (i.e. either an rsID, or in a format containing the chromosome, base pair (bp), and the two alleles of the SNP), that the missing string operator was set to "NA," and that all files had the same column delimiter.

##### *2.6.2 Filters applied before EasyQC protocol*

Following recommendations provided by the UK Biobank, we removed the 65 autosomal SNPs from the UKB that had been flagged as having incorrect annotation, together with the additional 535 SNPs that we flagged in section 1.5, before applying the EasyQC protocol described below.

Also, for cohorts imputed with the September or December 2013 haplotype release of the 1000 Genomes imputation reference panel, we removed 737 SNPs with incorrect strand alignment[e].

##### *2.6.3 EasyQC protocol*

The filters applied in the EasyQC software are explained below in chronological order of implementation. Note that the order of the filters does not influence the outcome of the cleaned GWAS summary statistics (although it affects at which specific filter a SNP is removed). The number of SNPs filtered at each step of the EasyQC protocol is reported in **Supplementary Table 2.3**.

Step 1 in the EasyQC protocol filtered out SNPs for which either the effect-coded allele or the other allele has values different from "A," "C," "G," or "T." This step removed all structural variants such as INDELs.

Step 2 filtered out SNPs with missing values for one or more of the following variables: *P* value, an estimated effect size (beta) or its standard error, frequency of the reference allele, imputation status, and imputation accuracy (conditional on the SNP being imputed). This filter also removed SNPs with nonsense values outside of permissible ranges such as negative or infinite standard errors, nonsensical *P* values, allele frequencies greater than 1 or below 0, as well as imputation status not equal to 1 or 0.

---

[e] The announcements are available on the webpage of IMPUTE2 https://mathgen.stats.ox.ac.uk/impute/data_download_1000G_1_integrated_SHAPEIT2_16-06-14.html and https://mathgen.stats.ox.ac.uk/impute/data_download_1000G_phase1_integrated_SHAPEIT2_9-12-13.html.

The thresholds chosen for the filters applied in steps 3 to 5 are summarized in **Supplementary Table 2.4**. Step 3 filtered out SNPs with a MAF below 0.1% for the UKB and 23andMe cohorts and below 1% for all other cohorts; this effectively removed any SNPs that were monomorphic in the summary statistics. Step 4 excluded SNPs based on imputation accuracy with a threshold contingent on the cohort-specific imputation software (0.6 for MACH, 0.7 for IMPUTE, and 0.8 for PLINK). Step 5 filtered out directly genotyped SNPs with a Hardy-Weinberg equilibrium exact test *P* value below a threshold contingent on the cohort sample size. The applied thresholds were $10^{-3}$ if $n < 1{,}000$, $10^{-4}$ if $1{,}000 \leq n < 2{,}000$, and $10^{-5}$ if $2{,}000 \leq n < 10{,}000$.

Two additional filters were applied to ensure that only high-quality SNPs were being forwarded to the meta-analysis; step 6 removed SNP $j$ if

$$\text{(2)} \qquad \widehat{SE}_j > 1.4 \frac{\hat{\sigma}_Y}{\sqrt{2 \cdot n_j \cdot MAF_j \cdot \left(1 - MAF_j\right)}},$$

where $\hat{\sigma}_Y$ is the standard deviation of the phenotype, $n_j$ is the sample size, $SE_j$ is the standard error of the coefficient estimate for SNP $j$, and $MAF_j$ is the minor allele frequency of SNP $j$. This filter eliminates SNPs whose coefficient estimates have standard errors that are more than ~40% larger than what would be expected given the sample size, the MAF of the SNP, and the standard deviation of the phenotype. The second additional filter, step 7, removes SNPs with coefficient estimates larger than what would correspond to an $R^2$ greater than 5%. We adapted the filter from Okbay *et al.*[16] using an approximation to the $R^2$: SNP $j$ is dropped if

$$\text{(3)} \qquad \left|\hat{\beta}_j\right| > \frac{\sqrt{0.05} \cdot \hat{\sigma}_Y}{\sqrt{2 \cdot MAF_j \cdot \left(1 - MAF_j\right)}}.$$

Step 8 filtered out duplicate SNPs (SNPs with identical NCBI build 37 (UCSC hg19) chromosome and base-pair positions). This was implemented after the chromosome and base-pair positions of the SNPs had been harmonized with the main reference panel described above.

Step 9 aligned the SNPs to the main reference panel to ensure that the effect-coded allele was the same for all SNPs across the cohorts. This step removed SNPs that were not available in the main reference panel as well as SNPs that displayed an allele mismatch when compared to the reference (e.g., a SNP with the alleles A and T in the GWAS summary statistics would be removed if the alleles according to the reference panel were A and G).

Step 10 removed SNPs that deviated from the main reference panel in terms of allele frequency. A SNP was removed if the absolute difference between its allele frequencies in the cohort's data and in the main reference panel was greater than 0.2. Step 10 was applied to all cohorts including the UKB (for the UKB, this filter was thus applied in addition to the filter described in **Supplementary Information section 2.5** and **Supplementary Information section 2.6.2**, the purpose of which was to avoid potential strand issues caused by the two different UKB genotyping arrays).

The output from the quality control was examined to see if any filters removed an unusual or unexpected number of SNPs. Some cohorts required iterations with the analysts to ensure that all possible errors were resolved. The number of SNPs filtered at each step of the final quality control iteration is reported in **Supplementary Table 2.3** together with the estimated genomic inflation factor ($\lambda_{GC}$).

##### *2.6.4 Visual inspection of diagnostic plots*

Once low-quality SNPs were filtered out, the remaining SNPs were used to produce several diagnostic plots for each cohort, most of which are the standard output of the EasyQC software. Visual inspection of these plots enabled the identification of possible issues or errors in the GWAS summary statistics of each cohort; for a more thorough discussion we refer the interested reader to Winkler *et al.*[40]. For any potential issues observed in these plots, we contacted the cohort-specific analyst and ensured that the observed issues were completely resolved. The following plots were examined:

*Allele Frequency Plots (AF Plots):* The AF plot contrasts the *observed* allele frequencies with the *expected* allele frequencies calculated according to the main reference panel, and the plot was created before step 10 of the EasyQC protocol. If the sample closely resembles the reference panel in terms of allele frequencies, then the SNPs should align in a diagonal with positive slope. This plot enables the analyst to detect deviations in ancestry from the reference as well as issues related to the alignment of the effect-coded allele. If the wrong effect-coded allele has been specified, then the AF plot shows a diagonal with negative slope.

*P-Z Plots:* Inspection of this plot shows if the reported *P* values are consistent with the reported coefficient estimates and their standard errors. One common problem observable with the P-Z plot is an erroneous column header in the GWAS summary statistics, such that the wrong column is used for either the beta estimates, standard errors or *P* values in the analysis.

*Q-Q Plots:* Inspection of Q-Q plots enables visualization of unaccounted-for stratification in the cohorts. No cohort displayed premature lift-off in the Q-Q plot associated with unaccounted-for stratification. The genomic inflation factors $\lambda_{GC}$ are displayed in **Supplementary Table 2.3**.

*SE Plots:* We plotted the observed standard errors ($SE_j$) of the coefficient estimates versus the standard errors expected given the $MAF_j$ and the sample size $n_j$ of a given SNP *j,* and the standard deviation of the phenotype $\hat{\sigma}_Y$ (which is equal to 1 if the phenotype has been standardized). This enables visual inspection to identify groups of outlier SNPs with regard to the observed standard error. The expected standard error was calculated according to the following formula:

$$\widehat{SE}_j \approx \frac{\hat{\sigma}_Y}{\sqrt{2 \cdot n_j \cdot MAF_j \cdot \left(1 - MAF_j\right)}} \quad (4)$$

All cohorts had to pass visual inspection as well as inspection of the number of excluded SNPs at each of the exclusion filters described in the previous subsection before being passed on for the meta-analysis.

#### 2.7 Meta-analysis and adjustment of the standard errors

Sample-size weighted meta-analysis of the cleaned cohort-level GWAS summary statistics were carried out using the METAL software[41]. We conducted four main meta-analyses: (1) we meta-analyzed the discovery GWAS combining the UKB and 23andMe cohorts; (2) we meta-analyzed the results of the 10 replication cohorts without the UKB and 23andMe discovery cohorts, to obtain our replication GWAS; (3) we meta-analyzed the results of the 10 replication cohorts together with those of the UKB and 23andMe discovery cohorts for the follow-up analyses that use GWAS summary statistics; and (4) we meta-analyzed the results from our UKB GWAS of ever smoker with those from the TAG Consortium[29]. No meta-analyses were conducted for the five other supplementary GWAS, because data for these GWAS each came from only one cohort (either the UKB or the 23andMe cohort).

All meta-analyses were restricted to SNPs with a sample size greater than half of the maximum SNP sample size and were performed with the "ChrPosID" as the identifier of each SNP (e.g. 1:123456)[18] .

Instead of applying genomic control with the over-conservative $\lambda_{GC}$, we inflated the standard errors by the square root of the estimated intercept from an LD Score regression. This procedure allows us to correct only for inflation of test statistics caused by population stratification and other confounding factors rather than polygenicity[42].

For the discovery and replication GWAS of general risk tolerance and for the GWAS of ever smoker—all of which involved meta-analyses of cohort-level data—we only inflated the meta-level standard errors (i.e., we did not inflate the cohort-level standard errors before the meta-analysis). Likewise, for the meta-analysis of the discovery and replication GWAS for the follow-up analyses, we only inflated the meta-level standard errors. We also inflated the standard errors of the other supplementary GWAS.

In practice, for a given meta-analysis, the METAL software[41] outputs the SNPs' meta-analyzed $z$-statistics, deflated by the square root of the estimated intercept from an LD Score regression. We use SNP $j$'s GWAS sample size $n_j$ and minor allele frequency $MAF_j$, as well as the phenotype's standard deviation $\hat{\sigma}_y$, to approximate the inflated standard error of our estimate of SNP $j$'s effect size:

$$\widehat{SE}_j \approx \sqrt{Intercept} \cdot \frac{\hat{\sigma}_y}{\sqrt{2 \cdot n_j \cdot MAF_j(1 - MAF_j)}},$$

where $\sqrt{Intercept}$ is the square-root of the LD Score intercept used to deflate the $z$-statistic in the meta-analysis.

We then used SNP $j$'s deflated $z$-statistics $\hat{z}_j$ to approximate SNP $j$'s effect size as $\hat{\beta}_j \approx \hat{z}_j \cdot \widehat{SE}_j$ [f]. Since the general-risk-tolerance phenotype is not measured in natural units, and since its standard deviation differs across cohorts, we normalize it to have a standard deviation of one when we estimate the SNPs' effect sizes and standard errors. For consistency, we make the same assumption when approximating the SNPs' effect sizes and standard errors for the risky

[f] Since $\hat{z}_j$ is deflated and $\widehat{SE}$ is inflated by the square root of the intercept from the LD score regression, $\hat{\beta}_j$ is neither deflated nor inflated.

behaviors and their first PC. Hence, our estimated effect sizes (the $\hat{\beta}_j$'s) are expressed in standard-deviation units of the phenotype per effect-coded allele; we hereafter refer to this as a "standardized beta," although it is only standardized in terms of standard deviation units of the phenotype (and not with respect to the genotype). The standard deviations of the phenotypes, as originally measured, are reported in **Supplementary Table 1.2**.

The coefficient of determination of SNP $j$ is approximated as[43]:

(5)
$$R_j^2 \approx \frac{2 \cdot MAF_j(1 - MAF_j) \cdot \hat{\beta}_j^2}{\hat{\sigma}_y^2}.$$

#### 2.8 Independence of significant loci and definition of locus

To identify approximately independent genome-wide significant loci, we used PLINK[37] to apply a "clumping algorithm" to the GWAS results. Our clumping algorithm identifies lead SNPs and defines independent loci that include SNPs with an $r^2$ greater than 0.1 with each lead SNP. (A lead SNP is the most significant of the genome-wide significant SNPs in an independent locus.) Specifically, the clumping algorithm uses four parameters: a primary $P$ value threshold ($5\times10^{-8}$), a secondary $P$ value threshold ($1\times10^{-4}$), an $r^2$ threshold (0.1), and a SNP window defined in kilobases (1,000,000 kb). First, the SNP with the lowest $P$ value (less than the primary $P$ value threshold) is taken as the lead SNP in the first clump, and the first clump is formed by all SNPs with a $P$ value smaller than the secondary $P$ value threshold[g], with an $r^2$ greater than 0.1 with the clump's lead SNP, and within a distance less than the SNP window from the lead SNP[h]. Next, the SNP with the second lowest $P$ value (less than the primary $P$ value threshold) outside the first clump becomes the lead SNP of the second clump, and the second clump is created analogously but using only the SNPs outside of the first clump. This process continues until all SNPs with a $P$ value less than the primary $P$ value threshold are either defined as the lead SNP of a clump or clumped with another lead SNP. The $r^2$ was calculated with the main reference panel.

Thus, our definition of a locus is the set of all SNPs that are in weak or stronger LD (pairwise $r^2 > 0.1$) with a lead SNP (and that have not been clumped to another, more significant, lead SNP).

We note that, because we consider approximately independent (pairwise $r^2 < 0.1$) lead SNPs (rather than fully independent lead SNPs), some of our lead SNPs could in principle be secondary associations that are driven by their LD with extremely strong primary associations. However, based on the results we present in **Supplementary Information section 3.3** and report in **Supplementary Tables 3.1-3.2**, we believe this is unlikely to be the case for the vast majority of our lead SNPs, because they are not located near other associations that could be strong enough to be driving secondary associations. One exception is a very strong association in the gene *ADH1B* with drinks per week (rs1229984, $P = 7.80\times10^{-202}$), which could potentially be responsible for the genome-wide significance of some nearby lead SNPs in our GWAS of drinks per week.

[g] The secondary $P$ value threshold lowers the computational effort by allowing the algorithm to ignore SNPs with large $P$ values.

[h] We used a very wide SNP window of 1,000,000 kb, which effectively makes the $r^2$ and $P$ value thresholds the only binding parameters for the PLINK clumping algorithm.

To see how sensitive the number of lead SNPs we report is to the $r^2$ threshold used in the clumping algorithm, for our discovery GWAS of general risk tolerance we also applied the clumping algorithm with the more stringent $r^2$ threshold of 0.01 (instead of 0.1) (**Supplementary Information section 3.3**).

#### 2.9 Check for long-range LD regions, candidate inversions, and 1000 Genomes structural variants

##### *2.9.1 Check for long-range LD regions*

We investigated if the lead SNPs were located in larger structural variation in the form of long-range LD regions, by using a set of long-range LD regions from Price *et al.*[44]. Price *et al.* identified 24 long-range LD regions from 327 European-ancestry individuals, which replicated in two independent samples comprising 1593 European-ancestry Americans and 3004 British individuals. We lifted the genomic positions of the long-range LD regions to build 37 (GRCh37) with the UCSC genome-annotation lift-over tool[i], and there were nine long-range LD regions that could not be lifted due to non-overlapping genome sequences or ambiguous mapping across the builds. Hence, the combined map contains 15 long-range LD regions that have non-ambiguous genomic positions available in build 37. These range from ~2.5 to ~8 Mb in genomic size.

We checked if each lead SNP from the GWAS was within a long-range LD region, or within 250 bp from the breakpoints of such a region. The results are reported in **Supplementary Table 3.1** and **Supplementary Table 3.2**.

##### *2.9.2 Check for candidate inversions*

We also investigated if the lead SNPs were located in larger structural variation in the form of candidate inversions, by using a list of genomic segments highly prone to inversion polymorphisms from an unpublished resource by Gonzalez, J.R. & Esko, T., (2017, unpublished). The genomic segments highly prone to inversion polymorphisms were identified based on the knowledge that submicroscopic human inversions are typically flanked by highly homologous flanking repeats[45], which predisposes their occurrence by non-allelic homologous recombination. Therefore, a set of segments was selected that may be prone to submicroscopic inversions, consisting of all single copy segments in the Genome Reference Consortium's human reference sequence build 36 (GRCh36) between 0.1 and 8 Mb in length, and flanked by segmental duplications with 90% identity (across the flanking duplications). In total, there were 173 segments that met these criteria and that were thus considered as genomic segments highly prone to inversions. As detectable traces of inversions in SNPs depend on many factors—such as being frequent, ancient and nonrecurring—we tested whether the segments showed inversion patterns in any of two different SNP datasets. First, 69 (40%) of the segments overlapped with the inversions that Caceres *et al.*[46] obtained in the phased genotypes of CEU individuals from the HapMap III project. Second, inversion-like haplotypes[47] were inferred in a subsample of 882 Estonians for which gene expression data was available in peripheral blood. In this case 65 (38%) of the 173 segments were significantly associated with the expression of single copy

[i] The lift-over tool is available here: https://genome.ucsc.edu/cgi-bin/hgLiftOver

genes within the segment. In total 104 (60%) of the 173 segments showed an inversion signal, indicating their predisposition for inversion occurrence.

We lifted the genomic positions of the genomic segments highly prone to inversion polymorphisms to build 37 (GRCh37) with the UCSC genome-annotation lift-over tool[j], and there were 19 genomic segments that could not be lifted due to non-overlapping genome sequences or ambiguous mapping across the builds. Hence, the combined map contains 154 segments prone to inversion polymorphisms (hereafter referred to as "the 154 candidate inversions"), that have non-ambiguous genomic positions available in build 37. These range from ~500 kb to ~8 Mb in genomic size.

We checked if each lead SNP from the GWAS was within such a candidate inversion, or within 250 bp from the breakpoints of such a candidate inversion. The results are reported in **Supplementary Table 3.1** and **Supplementary Table 3.2**.

##### *2.9.3 Check for 1000 Genomes structural variants*

Sudmant *et al*. (2015)[48] called and classified a large number of structural variants (SVs) with the final version of the 1000 Genomes Project phase 3 reference panel. They have released an integrated map of 37,250 smaller structural variants, together with enhanced resolution of the size and breakpoint compared to previous publications, and we hereafter refer to these as the "1000G structural variants." The structural variants range from 1 bp to ~445 kb in genomic size. The smallest variants are generally insertions of 1 bp (~6,900 variants), and the larger variants are generally deletions larger than 50 bp. The majority of these structural variants are in LD with proximate SNPs, and we therefore checked if any of the lead SNPs from our main GWAS, and SNPs in strong LD with those lead SNPs, were located within the start and end positions of any of the 37,250 structural variants. We defined strong LD as an $r^2$ greater than 0.8, which is the definition used by the 1000 Genomes Project Consortium[48]. The SNPs in LD were extracted using PLINK[37] and the main reference panel. We report the results in **Supplementary Table 3.1** and **Supplementary Table 3.2**.

#### 2.10 Investigation of the novelty of our GWAS associations

To investigate the novelty of our GWAS associations, we performed lookups of our lead SNPs (and the SNPs in LD with the lead SNPs, $r^2 > 0.1$) in the NHGRI-EBI GWAS Catalog database (revision 2017-08-15)[49] of genome-wide significant associations from previous GWAS. We also looked up our lead SNPs (and the SNPs in LD, $r^2 > 0.1$) in some recent GWAS articles that have not been catalogued in the NHGRI-EBI GWAS Catalog database. The NHGRI-EBI GWAS Catalog is a resource that aims to catalogue all associations reported in published GWAS.

For general risk tolerance, we performed a search with the term "risk" in the index of phenotypes, and we did not find any previous studies on general risk tolerance in the Catalog. We know of one previous study that identified one independent genome-wide significant association with general risk tolerance, and of one concurrent study that identified a second genome-wide significant association, both using the first UKB data release[50,51]; the authors of ref.[50] referred to the phenotype as "risk-taking propensity," and the authors of ref.[51] referred to it as "risk-taking behavior." The first genome-wide significant association is replicated in an online

[j] The lift-over tool is available here: https://genome.ucsc.edu/cgi-bin/hgLiftOver

publication published in advance[52]. We added the first of these two studies (i.e., Day *et al.*[50]) to our investigation of the novelty of our general-risk-tolerance lead loci, and the second we consider concurrent. We also note that, in **Supplementary Information section 11.1**, we report the results of a literature search of association studies of risk tolerance; that literature search identified no previously reported genome-wide significant associations.

The phenotypes drinks per week, ever smoker, and number of sexual partners (or related phenotypes such as alcoholism and age at first sex), were available in the NHGRI-EBI GWAS Catalog database (revision 2017-08-15). Since the GWAS Catalog is not always up-to-date, we additionally performed a literature search for genome-wide significant findings that might not yet have been included in the Catalog. We searched the Pubmed literature database on March 6 2017 and September 13 2017, for the term "genome-wide association study" together with each of the terms "alcohol," "sexual," and "smoking" individually. We screened the abstracts and compared the resulting articles with the article list of the GWAS catalog. In addition to what was already reported in the GWAS Catalog (revision 2017-08-15), we found three additional studies with genome-wide significant findings on alcohol consumption, two additional studies on smoking, and no additional studies on sexual behaviors.

To our knowledge, this is the first GWAS of adventurousness, automobile speeding propensity, and of the first PC of the four risky behaviors; unsurprisingly, we could not find previous GWAS on any of these phenotypes in the NHGRI-EBI GWAS Catalog database (revision 2017-08-15).

#### 2.11 GWAS catalog lookup

We investigated whether the lead SNPs from our discovery GWAS of general risk tolerance and from our supplementary GWAS have previously been associated at genome-wide significance with any phenotypes in the NHGRI-EBI GWAS Catalog database[49] (revision 2017-08-15). We query the GWAS Catalog with our list of lead SNPs, and the SNPs in LD with a lead SNP ($r^2 > 0.6$).

Since the NGRI-EBI GWAS Catalog is not always up-to-date with results from the most recent publications (especially in non-peer reviewed outlets such as BioRxiv), we also queried the summary statistics of the most recently published GWAS on attention deficit hyperactivity disorder[53], autism spectrum disorder[54], and anorexia nervosa[k]. We also queried the genome-wide significant findings from the additional GWAS on alcohol intake and smoking, which we found in addition to the GWAS Catalog, as detailed in the previous section. We queried the additional GWAS on alcohol intake and smoking because they contained at least one genome-wide significant result; because their results were publicly available; and because the meta-analysis that produced them did not include the UK Biobank (that comprise our discovery sample together with 23andMe).

We perform these lookups because the existence of SNPs and genes associated with both one of our studied phenotypes and another phenotype can point to a common genetic etiology. However, it is important to note that two phenotypes that share a genetic locus do not necessarily have to share the same causal *variant* at that locus due to the widespread LD that characterizes

[k] The Psychiatric Genomics Consortium's GWAS summary statistics for attention deficit hyperactivity disorder, autism spectrum disorder, and anorexia nervosa (referred to as "ED," i.e. eating disorder) are publicly available and can be downloaded here: https://www.med.unc.edu/pgc/results-and-downloads.

the human genome. Moreover, even if two phenotypes do share a single causal variant, they do not have to share general underlying genetic etiologies. For instance, recent work[24] has shown that two types of autoimmune diseases (rheumatoid arthritis and ulcerative colitis/Crohn's disease) that are known to share risk loci are not genetically correlated at a genome-wide level. Here, the reason was the lack of an overall directional trend: some risk alleles for one disease were also risk alleles for the other disease, but some alleles that were *protective* for the one disease were risk alleles for the other. This resulted in a near-zero correlation at the genome-wide level. Thus, we emphasize that the current lookup does not make it possible to determine *etiological* overlap, but only hints at overlapping loci, between general risk tolerance or the supplementary GWAS phenotypes, and the phenotypes reported in the GWAS Catalog.

#### 2.12 Cross-lookup of GWAS results

We performed cross-lookups of the lead SNPs across our discovery GWAS of our primary and supplementary phenotypes. Specifically, for each lead SNP in each of the GWAS, we checked if the SNP is in LD (with an $r^2$ greater than 0.1) with lead SNPs in the other GWAS. LD was calculated using PLINK[37] and the main reference panel. The results of the cross-lookups are reported in **Supplementary Table 3.1** and **Supplementary Table 3.2**. As we describe above, we also investigated if there were any long-range LD regions or candidate inversions that contained lead SNPs for multiple GWAS.

#### 2.13 Gene annotation

We annotated the lead SNPs with gene information using the National Institute of Health (NIH) National Center for Biotechnology Information (NCBI) gene ontology database (version 2016-05-25)[1]. As the general rule, a SNP was annotated to its most proximate gene. If a SNP was located between two genes, then we compared the distance to the end coordinate of the gene upstream with the distance to the start coordinate of the gene downstream to find the most proximate gene. If a SNP was located within multiple overlapping genes, then the SNP was annotated to the gene with the most proximate start coordinate. This means that all lead SNPs were annotated to a single gene. The approach roughly partitions the SNPs throughout the genome into separate genomic segments. The annotations are displayed in **Supplementary Table 3.1** and **Supplementary Table 3.2**, where we also indicate if a lead SNP is located within or outside the gene's start and end coordinates. We also checked if there were genes to which lead SNPs from multiple GWAS were annotated (the results are reported in **Supplementary Information section 3.2**).

---

[1] The NCBI gene ontology database is available here:
ftp://ftp.ncbi.nlm.nih.gov/gene/DATA/GENE_INFO/Mammalia/.

### 3 GWAS results

In this section, we report, compare, and discuss the results of our seven main GWAS (of our primary phenotype—self-reported general risk tolerance—and of our six supplementary phenotypes—adventurousness, the four risky behaviors, and their first PC). In **Supplementary Information sections 3.1** and **3.2,** we present a summary of our results, and we also describe several notable genomic regions that contain lead SNPs for all or most of our main GWAS. **Supplementary Information sections 3.3** and **3.4** contain a more detailed description of the results of our discovery GWAS of general risk tolerance and the results of our six other main GWAS, respectively. Throughout, we place particular emphasis on long-range LD regions and candidate inversions (defined in **Supplementary Information section 2.9**) that contain lead SNPs for all or most of our GWAS. (We focus on these because, as we explain below, very few genomic blocks outside of these regions or candidate inversions contain lead SNPs shared across most of our GWAS.)

#### 3.1 Summary of the GWAS results

We identified 124 lead SNPs associated with general risk tolerance, 167 with adventurousness, 42 with automobile speeding propensity, 85 with drinks per week, 223 with ever smoker, 118 with number of sexual partners, and 106 with the first PC of the four risky behaviors[m]. To the best of our knowledge, 853 of these 865 associations are novel. We were able to replicate the only previously published[50] genome-wide significant association with general risk tolerance, located within *CADM2* on chromosome 3, that was also replicated in ref.[52] and in a concurrent study[51]. We replicated another genome-wide significant association with general risk tolerance from the concurrent study[51], located in proximity to the HLA-complex on chromosome 6. We also replicated the TAG Consortium's previous association with ever smoker in the gene *NCAM1*[29].

The NHGRI-EBI GWAS Catalog database[49] lookup of general-risk-tolerance lead SNPs resulted in 61 overlapping associations distributed across 27 lead SNPs, and the lookup of our six other main GWAS resulted in 939 overlaps distributed across 130 lead SNPs. Notably, for all our main phenotypes we find overlaps with schizophrenia, cognitive performance, and information processing speed.

The detailed results of the cross-lookup of our GWAS results (the investigation of whether the lead SNPs of each of our GWAS are in LD, defined as a $r^2$ greater than 0.1, with the lead SNPs of our other GWAS (**Supplementary Information section 2.12)**) are reported in **Supplementary Tables 3.1** and **3.2**. In total, we identified 865 lead SNPs across the GWAS of our primary and supplementary phenotypes. Of these 865, 34 exact lead SNPs are counted twice by being shared by two phenotypes (there are no exact lead SNPs shared across more than two

[m] It should be noted that for each of our seven main GWAS phenotypes, there are sets of lead SNPs located within the same long-range LD region[44] or candidate inversion. Taking this into account in our clumping algorithm might result in lower estimates of the total number of loci we identified. However, estimating the independence of GWAS associations in these regions requires conditional GWAS analyses, and we therefore chose to keep our definition of independent loci in these regions and to indicate which SNPs are located in these regions (**Supplementary Table 3.1** and **3.2**).

phenotypes; thus, in total there are 831 unique lead SNPs identified across the phenotypes). 441 of the 865 lead SNPs are in loci that also contain a lead SNP for at least one of the other GWAS.

To find the number of independent loci across all our phenotypes, we used the clumping algorithm described in **Supplementary Information section 2.8** on the 865 lead SNPs. We first removed the duplicate lead SNPs by keeping the strongest associated SNP as one record, and the clumping algorithm was thereafter applied to the 831 unique lead SNPs that had the strongest associations across the phenotypes. The clumping algorithm resulted in 611 independent loci across the phenotypes.

To find how many loci from the six supplementary GWAS are independent from the 124 loci identified in our GWAS of general risk tolerance (our primary GWAS), we removed the 124 general-risk-tolerance lead SNPs, as well as the SNPs in LD ($r^2 > 0.1$) with them, from the 865 lead SNPs. After removing duplicate lead SNPs by keeping the strongest associated SNP as one record, there were 599 remaining unique lead SNPs that are approximately independent from the 124 general-risk-tolerance lead SNPs (but not necessarily independent from one another). Finally, we re-ran the clumping algorithm and we identified 483 approximately independent lead SNPs across our six supplementary GWAS.

In sum, our seven GWAS identified 865 lead SNPs located across 611 approximately independent loci. Of the 865 lead SNPs, 68 involve 34 exact lead SNPs that were each identified in two separate GWAS, and 441 are in loci that also contain a lead SNP for at least one of the other GWAS. Removing the loci around our 124 general-risk-tolerance lead SNPs left us with 483 approximately independent loci that contain a lead SNP for at least one of our other six GWAS phenotypes.

#### 3.2 Summary of genetic overlap across our main GWAS

##### *3.2.1 Summary and discussion of notable genomic regions*

Five genomic regions stand out because they contain lead SNPs for all or most of our seven phenotypes. Two of these regions are among the 15 long-range LD regions identified by Price *et al.*[44], and the other three are among the 154 genomic segments deemed highly prone to inversion polymorphisms (i.e., the 154 "candidate inversions"; both the 15 long-range LD regions and the 154 candidate inversions are described in **Supplementary Information section 2.9**). One long-range LD region (chromosome 3, ~83.4 to 86.9 Mb, **Extended Data Fig. 3.4**) and one candidate inversion (chromosome 18, ~49.1 to 55.5 Mb, **Extended Data Fig. 3.4**) each contain lead SNPs for all of our seven GWAS phenotypes; the other three regions each contain lead SNPs for general risk tolerance and for four or five of our six supplementary GWAS.

The first notable region is the long-range LD region on chromosome 3 (~83.4 to 86.9 Mb, **Extended Data Fig. 3.4**), which contains lead SNPs for all our GWAS. The only gene within the long-range LD region is *CADM2*, and there are few other genes in proximity (only *VGLL3* is within 250 kb of the breakpoints, ~69.8 kb downstream). The relatively large *CADM2* gene (~85.0 to 86.2 Mb) covers ~1.2 Mb of the ~3.5 Mb long-range LD region and contains our strongest association with general risk tolerance (rs993137, $P = 2.14\times10^{-40}$). The Bonferroni-corrected $P$-value of *CADM2* in the MAGMA gene analysis is $P = 1.09\times10^{-50}$ (**Supplementary Table 12.3**), consistent with the GWAS results. A recent study[52] reports that *CADM2* contains a replicated association with general risk tolerance, and that study also reports suggestive associations between SNPs in *CADM2* and different measures of personality. As we discuss in **Supplementary Information section 12**, *CADM2* “encodes a member of the synaptic cell adhesion molecule 1 (SynCAM) family which belongs to the immunoglobulin (Ig) superfamily”[55], and it is related to synapse formation[56] and brain plasticity[57]. *CADM2* is overexpressed in the brain, and in particular in the frontal cortex, according to GTEx[58]. The GWAS Catalog database[49] reports genome-wide significant associations in the long-range LD region with many phenotypes, including age at menarche, BMI, educational attainment, and information processing speed. Most of the GWAS Catalog database[49] associations within the long-range LD region (~83.4 to 86.9 Mb on chromosome 3) are annotated to the gene *CADM2*.

The second notable region is a candidate inversion located on chromosome 18 (~49.1 to 55.5 Mb, **Extended Data Fig. 3.4**). Within the candidate inversion there are previous genome-wide associations in the GWAS Catalog database[49] with traits such as autism spectrum disorder, ADHD, depression, educational attainment, schizophrenia, and subcortical brain region volumes. The candidate inversion contains ~20 genes, and the MAGMA gene analysis resulted in one significant gene after Bonferroni correction—*TCF4* (Bonferroni-corrected $P = 5.51\times10^{-9}$, **Supplementary Table 12.3**). *TCF4* is interesting because it is known to play an important role in nervous system development[55]. *De novo* mutations in *TCF4*[59] are known to cause the rare Pitt-Hopkins syndrome[60], with few described cases in the medical literature[61]. The syndrome is characterized by distinct facial features, intellectual disability, delayed motor skills, and epilepsy, among many other symptoms[59–61]. The GWAS Catalog database[49] reports genome-wide significant associations mapped to *TCF4* with schizophrenia, and *TCF4* has been hypothesized to be involved in more neuropsychiatric phenotypes[62]. This observation is consistent with the non-zero genetic correlations that we estimated with bivariate LD Score regression between general risk tolerance and many neuropsychiatric disorders (**Fig. 2, Supplementary Information section 7**). As we describe below, all phenotypes have lead SNPs annotated to *TCF4* except ever smoker, for which there are three lead SNPs within the candidate inversion annotated to the proximate genes *DCC* and *TXNL1*. We annotated two lead SNPs for general risk tolerance, two for adventurousness, two for automobile speeding propensity, one for drinks per week, six for number of sexual partners, and three for the first PC of the risky behaviors, to *TCF4*.

The third notable genomic region is the long-range LD region located on chromosome 6 (~25.3 to 33.4 Mb, **Extended Data Fig. 3.4**). The region covers the HLA-complex[55] (~29.6 to 33.1 Mb on chromosome 6[63]), and contains lead SNPs for all phenotypes except drinks per week (for which we identified a suggestive association, with a $P = 3.83\times10^{-7}$). There are at least 250 genes in the region, and the MAGMA gene analysis resulted in ~30 significant genes after Bonferroni-correction (however, none of the actual HLA genes were significant; **Supplementary Table 12.3**). The GWAS Catalog database[49] reports more than a thousand genome-wide significant associations in the region. The associations relate to hundreds of traits, including alcohol consumption, Alzheimer’s disease, autism spectrum disorder, educational attainment, and schizophrenia. The HLA-complex contains thousands of SNP associations in the GWAS Catalog database[49], and those relate to hundreds of traits, including alcohol consumption, Alzheimer’s disease, autism spectrum disorder, and educational attainment. The HLA genes encode the major histocompatibility complex (MHC) proteins, whose main function is to alert cells of the immune system about infection from pathogens[64]. The region is known to be highly polygenic, and its high rates of recombination lead to a large number of possible haplotypes in the population. For certain classes of MHC proteins there are more than 1,000 alleles in the human population, and most individuals are therefore heterozygous. However, there is also a small subset of proteins encoded for which the coding alleles are practically monomorphic, and this is probably caused by certain constraints on the viability of the variability for these specific protein chains. The HLA genes have been found to be under strong selection[n]. The constant struggle to counter pathogens results in selection pressures that favor polymorphisms that are able to provide protection.

The fourth notable genomic region is the candidate inversion on chromosome 7 (~124.6 to 132.7 Mb; **Extended Data Fig. 3.4**), which was identified in all our GWAS except automobile speeding propensity (for which it contains a suggestive association: rs141450, $P = 7.88\times10^{-8}$). The candidate inversion contains ~50 genes, and 5 of those were significant after Bonferroni correction in the MAGMA gene analysis (**Supplementary Table 12.3**). Among them are *SND1* (Bonferroni-corrected $P = 5.08\times10^{-10}$) and *PAX4* (Bonferroni-corrected $P = 1.43\times10^{-5}$). *SND1* is implicated as an important factor for normal cell growth[55], and *PAX4* is critical to normal fetal development[55]. The candidate inversion contains genome-wide significant associations in the GWAS Catalog database[49] with alcohol dependence, educational attainment, and schizophrenia, among other phenotypes.

The fifth notable genomic region is the candidate inversion on chromosome 8 (~7.89 to 11.8 Mb), which contains lead SNPs for all of our GWAS except those of drinks per week and of the first PC of the risky behaviors (**Extended Data Fig. 3.4**). (The strongest associations with drinks per week and the first PC of the risky behaviors have $P$ values of $5.64\times10^{-4}$ and $1.27\times10^{-7}$, respectively.) There are ~20 genes within the candidate inversion, and interestingly, practically all are significant in the MAGMA gene analysis after Bonferroni-correction (**Supplementary Table 12.3**). Two notable examples are *MSRA* and *CTSB* (with Bonferroni-corrected $P$ values of $2.94\times10^{-24}$ and $4.37\times10^{-5}$, respectively). *MSRA* is known to be highly expressed in human nervous tissue[55], and *CTSB* has a known effect on the processing of an amyloid precursor protein (APP)[55]. Incomplete processing of APP has been suggested as one of the causes of Alzheimer's disease[65]. However, the GWAS Catalog database[49] does not report any previous associations with Alzheimer's disease within the candidate inversion, or within 500kb of its breakpoints. The GWAS Catalog database[49] reports genome-wide associations within the breakpoints of the candidate inversion with many other phenotypes, including extraversion, schizophrenia, and chronotype, and the candidate inversion has been analyzed in-depth in a study of neuroticism and depressive symptoms[18].

##### *3.2.2 Overlap within approximately independent LD blocks*

The observation that several genomic regions contain at least one lead SNP from all or most of our GWAS prompted further investigation. To begin with, we divided the genome into 1703 approximately independent LD blocks identified by Pickrell *et al.*[66]. We then counted the number of blocks that contain lead SNPs from exactly one, two, three, four, five, six, and seven of our main GWAS. Of the 1703 LD blocks, 234 contain at least one lead SNP from exactly one of our seven main GWAS; 93 contain at least one lead SNP from exactly two GWAS; 40 contain at least one lead SNP from exactly three GWAS; 20 contain at least one lead SNP from exactly four GWAS; 7 contain at least one lead SNP from exactly five GWAS; one block (~84.4 Mb to 85.6 Mb on chromosome 3) contains at least one lead SNP from exactly six GWAS; and one

[n] A good example of the strong selection is the differences in allele frequencies across populations for alleles that affect resistance to a lethal form of malaria (see, e.g., ref.[244]). The protective alleles are very common in areas where the disease is endemic compared to areas where the risk of infection is low or absent.

block (~51.6 Mb to 55.2 Mb on chromosome 18) contains at least one lead SNP from all seven of our main GWAS. These last two blocks on chromosomes 3 and 18 are respectively located within the first and the second notable genomic region highlighted above and displayed in **Extended Data Fig. 3.4**.

The nine blocks that contain at least one lead SNP from exactly five, six, or seven GWAS each contain at least one general-risk-tolerance lead SNP. Five of these nine blocks are located within the five notable regions described above, with one block in each of the five regions[o]. The striped areas in **Fig. 3.4** show these five blocks. The five blocks include the two blocks that contain lead SNPs from six and from all seven of our GWAS. The other four blocks are located on chromosome 2 (~44.3 to 46.5 Mb), chromosome 3 (~70.5 to 72.5 Mb), chromosome 6 (~97.8 to 100.6 Mb), and chromosome 7 (113.7 ~ 116.8 Mb). Thus, nine genomic regions contain lead SNPs for at least five of our seven GWAS: the five notable regions described in the previous subsection and the four LD blocks located on chromosomes 2, 3, 6, and 7.

We ran a simulation to benchmark those results and to assess the likelihood of observing such a high level of within-block overlap across the GWAS results, under the null hypothesis that the LD blocks containing the lead SNPs of each GWAS are distributed randomly across the genome and independently from those of the other GWAS. (This null hypothesis is not perfectly realistic: in practice, the phenotypes are phenotypically correlated and their GWAS samples overlap substantially, and some LD blocks are located within regions of the genome that are more likely to contain causal variants. Thus, although informative, this simulation exercise has limitations.)

We conducted this analysis with only the general risk tolerance, automobile speeding propensity, drinks per week, ever smoker, and number of sexual partners phenotypes. We excluded adventurousness and the first PC of risky behaviors, because adventurousness is strongly genetically correlated with general risk tolerance, and because the first PC of risky behaviors is constructed from the four risky behaviors. The simulation proceeded in the following way: Since larger LD blocks are more likely to contain lead SNPs for any of our phenotypes, we classified each LD block by length and randomly permuted the LD blocks within each length class independently for each phenotype[p]. We then counted the number of LD blocks that contain hits from exactly one, two, three, four, and five GWAS, and we averaged these numbers over 10,000 simulations.

Our results indicate that, under the null hypothesis, we would expect 357.064 LD blocks to contain at least one lead SNP from exactly one GWAS; 43.212 to contain at least one lead SNP from exactly two GWAS; 2.734 to contain at least one lead SNP from exactly three GWAS; 0.077 to contain at least one SNP from exactly four GWAS; and 0.001 to contain at least one lead SNP from all five GWAS. By contrast, the actual number of blocks that contain at least one lead SNP from exactly one, two, three, four, and five of these GWAS are 253, 61, 20, 3, and 1.

[o] There are 27 LD blocks that overlap with the five notable genomic regions, and these are shown, separated by the dotted vertical gray lines, in **Extended Data Fig. 3.4**. Of these 27 blocks, four blocks contain at least one lead SNP from exactly one GWAS; two contain at least one lead SNP from exactly two GWAS; four contain at least one lead SNP from exactly three GWAS; three contain at least one lead SNP from exactly four GWAS; three contain at least one lead SNP from exactly five GWAS; one contains at least one lead SNP from exactly six GWAS; and one contains at least one lead SNP from all seven GWAS (these last two blocks are the ones on chromosomes 3 and 18, located within the first and the second notable genomic regions).

[p] We sorted the LD blocks into 9 classes based on the following lengths: 0 to 0.5 Mb (36 blocks), 0.5 to 1 Mb (263 blocks), 1 to 1.5 Mb (526 blocks), 1.5 to 2 Mb (478 blocks), 2 to 2.5 Mb (256 blocks), 2.5 to 3 Mb (82 blocks), 3 to 3.5 Mb (29 blocks), 3.5 to 7.5 Mb (25 blocks), and greater than 7.5 Mb (8 blocks).

The expected overlap from our simulation thus differs markedly from the overlap that we actually observe. To investigate this more formally, we conducted a non-parametric Mann-Whitney test[67] to compare the actual and simulated distributions of within-block overlaps across GWAS. Our results strongly suggest that the distribution of overlap in the simulation is different from the distribution of overlap that we actually observe ($P = 0.0023$).

The simulation exercise thus clearly suggests that the high level of within-block overlap across the results of our seven GWAS are highly unlikely to be due to chance. However, we emphasize again that our null hypothesis is not perfectly realistic and that the simulation exercise has limitations, and that these results should therefore be interpreted with caution.

##### *3.2.3 Concordance of SNP effects across the nine regions associated with five or more of our phenotypes*

Above, we identified nine genomic regions that contain lead SNPs for at least five of our seven GWAS: the five notable regions on chromosomes 3, 6, 7, 8, and 18, and the four LD blocks located on chromosomes 2, 3, 6, and 7. We investigated whether the signs of the lead SNPs located in these regions tend to be concordant across our primary and supplementary GWAS (in the sense that general-risk-tolerance-increasing alleles are also associated with higher risk taking in the supplementary GWAS, and vice-versa).

These nine regions harbor a total of 37 general-risk-tolerance lead SNPs and 111 lead SNPs from our six supplementary GWAS[q]. 217 coefficients of the 37 general risk tolerance lead SNPs are available in the results of the six supplementary GWAS (one of the 37 SNPs is only available in the adventurousness GWAS). 205 of these coefficients are have concordant signs, 147 are significant at the 5% level in the supplementary GWAS, and 26 are genome-wide significant. Under the null hypothesis that the coefficients have an equal probability of being concordant or discordant across the general risk tolerance and supplementary GWAS, the probability of observing 205 or greater concordant coefficients is very small ($P \leq 9 \times 10^{-47}$). We only found 12 discordant coefficients (9 of which are for drinks per week), and the total number of discordant coefficients is reduced to 4 if for each of the nine regions we exclude coefficients of GWAS that do not contain any lead SNP in the region. None of the discordant coefficients are genome-wide significant, and only 1 is significant at the 5% level (for drinks per week).

As for the 111 lead SNPs from our six supplementary GWAS, 109 of these 111 SNPs have concordant coefficients in our GWAS of general risk tolerance. Under the null hypothesis that the coefficients have an equal probability of being concordant or discordant across the general risk tolerance and supplementary GWAS, the probability of observing 109 or greater concordant SNPs is very small ($P \leq 3 \times 10^{-30}$). 98 of the concordant coefficients are significant at the 5% level for general risk tolerance, and 31 of these are genome-wide significant; the two discordant coefficients are not significant at the 5% level for risk tolerance.

Thus, the signs of the lead SNPs located in these regions tend to be highly concordant across our primary and supplementary GWAS, which suggests that these regions represent shared genetic influences on our seven phenotypes (rather than colocalization of causal SNPs).

---

[q] There are 141 unique lead SNPs across our seven main GWAS in these nine regions. Note that these lead SNPs were obtained using a clumping algorithm with an $r^2$ threshold of 0.1. If we instead used a threshold of 0.01, there would likely be fewer lead SNPs. See **Supplementary Information sections 2.8** and **3.3** for more details.

##### *3.2.4 Gene overlap across our seven GWAS*

We annotated each lead SNP from our seven GWAS with its most proximate gene, as described in **Supplementary Information section 2.13**. This approach roughly partitions the SNPs throughout the genome into different genomic segments—each associated with a single gene—that can be compared across our main GWAS. When we compared the gene-associated segments identified across our seven GWAS, we found that the only gene segment that is identified in all of them is the one associated with *CADM2* on chromosome 3 (that is located within the long-range LD region on chromosome 3, ~83.4 to 86.9 Mb, that is shared across all our GWAS; **Extended Data Fig. 3.4**).

The second most shared gene segment is the one associated with *TCF4* (discussed in more detail above). *TCF4* is located within the candidate inversion on chromosome 18 (~49.1 to 55.5 Mb) that is shared across all our GWAS (**Extended Data Fig. 3.4**). Lead SNPs from six of our GWAS, but not ever smoker, are annotated to *TCF4*. Three lead SNPs for ever smoker are located within the candidate inversion but are instead annotated to the genes *DCC* and *TXNL1*. *DCC* ends ~1.5 Mb before *TCF4*, and *TXNL1* starts ~967 kb after *TCF4*. Four segments, which were associated with the genes *FOXP1, MDFIC, SIX3*, and *VGLL3* (which is located ~69.8 kb downstream of the notable long-range LD region spanning ~83.4 to 86.9 Mb on chromosome 3), were each identified in five GWAS.

We searched the GWAS Catalog database[49] for previous associations with the genes associated with these shared segments, and we found some notable associations: *FOXP1* has been associated with ADHD, autism spectrum disorder, chronotype, and schizophrenia; *MDFIC* with BMI and obesity; *SIX3* with fasting plasma glucose, metabolite levels, and myopia; and *VGLL3* with age at menarche and pubertal anthropometrics.

#### 3.3 Results of the discovery GWAS of general risk tolerance

The discovery GWAS of general risk tolerance identified 124 independent genome-wide significant SNPs (i.e., lead SNPs), distributed across chromosomes 1 to 19. The strongest associations were found in the gene *CADM2* on chromosome 3 (rs993137, $P = 2.14\times10^{-40}$), on chromosome 7 in the gene *FOXP2* (rs7783012, $P = 7.57\times10^{-25}$) and in proximity to the gene *MDFIC* (rs9641536, $P = 5.01\times10^{-23}$), and in the gene *MFHAS1* (rs2409095, $P = 1.01\times10^{-20}$) on chromosome 8. We report the association results for the 124 lead SNPs in **Supplementary Table 3.1**, together with estimated effect sizes (the $\hat{\beta}_j$'s) expressed in phenotype standard-deviation units per effect-coded allele. **Supplementary Information section 5** reports the results of our successful attempt to replicate these results in our replication GWAS.

As mentioned in **Supplementary Information section 2.8**, to see how sensitive the number of lead SNPs is to our definition of an approximately independent locus (i.e., the set of all SNPs in weak LD (pairwise $r^2 > 0.1$) with a lead SNP), we counted how many lead SNPs our clumping algorithm yields when the more stringent $r^2$ threshold of 0.01 (instead of 0.1) is used. 94 lead SNPs remained when we applied the clumping algorithm with the more stringent threshold. The column "Lead SNP with stringent locus definition ($r^2 > 0.01$)?" in **Supplementary Table 3.1** reports whether each lead SNP remains a lead SNP with the more stringent threshold.

We display the Manhattan plot of the discovery GWAS in **Fig. 1a** and the quantile-quantile (Q-Q) plot in **Extended Data Fig 3.2a**. The genomic inflation factor ($\lambda_{GC}$) was 1.405 before, and

1.378 after inflation of the standard errors with the square root of the estimated intercept from an LD Score regression ($\widehat{Intercept} = 1.040, SE = 0.011$). As we discuss in **Supplementary Information section 4,** an observed genomic inflation factor larger than 1 is consistent with the expectation that complex traits are polygenic[68], and an estimated LD Score regression intercept close to unity suggests that the inflation is mainly due to polygenicity rather than to confounding factors[42].

All 124 general-risk-tolerance lead SNPs have MAF larger than 1%; only two of the lead SNPs have MAF lower than 5%; four have MAF between 5% and 10%; 33 have MAF between 10% and 25%; and 85 have MAF between 25% and 50%. Thus the vast majority of our lead SNPs are very common SNPs.

To evaluate the magnitude of the effect-size estimates of the 124 lead SNPs, we compared them with the 124 top associations reported in recent GWAS of height and body mass index (BMI), with the 74 top associations reported in a recent GWAS of educational attainment, and with the 48 top associations reported in a recent GWAS of waist-to-hip ratio adjusted for BMI (WHR)[r]. We display the comparison in **Extended Data Fig. 3.3**. The effect-size estimates of the 124 lead SNPs are consistently smaller than the top effect-size estimates of height, BMI, educational attainment, and WHR, both in terms of standard deviation units of the phenotype per effect-increasing allele, as well as in terms of variance explained ($R^2$). The general-risk-tolerance effect sizes range from 0.008 to 0.026 in phenotype standard-deviation units per effect-increasing allele, compared to a range of 0.038 to 0.191 for height, 0.019 to 0.082 for BMI, 0.014 to 0.048 for educational attainment, and 0.012 to 0.043 for WHR. The variance explained ($R^2$) by the general-risk-tolerance lead SNPs ranges from 0.003% to 0.019%, compared to 0.036% to 0.283% for height, 0.013% to 0.325% for BMI, 0.010% to 0.036% for educational attainment, and 0.004% to 0.072% for WHR.

We then paired the top SNPs of general risk tolerance with the top SNPs of height, BMI, educational attainment, and WHR (after ranking the SNPs by $R^2$ for each phenotype), and we calculated the median of the ratio of the $R^2$ across the paired SNPs. The median ratio was 13.64 between general risk tolerance and height (i.e., for the median ratio, the variance explained was 13.64 times larger for height compared to general risk tolerance), 4.58 between general risk tolerance and BMI, 3.85 between general risk tolerance and WHR, and 2.73 between general risk tolerance and educational attainment. Thus, effect sizes and $R^2$ of the general-risk-tolerance lead SNPs are substantially lower than those for the comparison phenotypes.

##### *3.3.1 Genetic correlation between females and males*

We used bivariate LD Score regression[42] to calculate the genetic correlation between GWAS performed separately in the sample of females and in the sample of males in the first release of UKB data. Our estimate of the genetic correlation ($\hat{r}_g$ = 0.949, $SE$ = 0.135) is not significantly different from 1, thus validating our strategy of pooling females and males in our GWAS. (For further details, see **Supplementary Information section 7.4**)

---

[r] The data for height, BMI, and WHR are from the publicly available GWAS results of the GIANT consortium, with males and females pooled and restricted to European-ancestry individuals, and the data for educational attainment are from the largest previously published GWAS of educational attainment by Okbay *et al.*[16].

##### *3.3.2 Genetic correlations between the UKB, 23andMe, and the replication GWAS*

Using bivariate LD Score regression[42] we estimated the genetic correlation between the cohort-level GWAS summary statistics from the general-risk-tolerance GWAS in the 23andMe and UKB cohorts, as well as between these GWAS and our replication GWAS (which includes 10 cohorts). Our estimates of the genetic correlations between the 23andMe and UKB GWAS ($\hat{r}_g =$ 0.767, *SE* = 0.021), between the 23andMe and the replication GWAS ($\hat{r}_g = 0.759$, *SE* = 0.126), and between the UKB and the replication GWAS ($\hat{r}_g = 0.828$, *SE* = 0.135) are all smaller than unity (though only the first of these estimates is significantly different from unity at the 5% level), pointing to substantial cross-cohort heterogeneity. (For further details, see **Supplementary Information section 7.4**.)

We also used bivariate LD Score regression[42] to estimate the genetic correlation between the summary statistics from the discovery GWAS and those from the replication GWAS. As in the case of the genetic correlation between males and females, we could not reject the null hypothesis of a genetic correlation equal to 1 ($\hat{r}_g$= 0.834, *SE* = 0.129). (For further details, see **Supplementary Information section 7.4)**

##### *3.3.3 Results of the cross-lookup of the general-risk-tolerance lead SNPs in the supplementary GWAS*

The results of the cross-lookup (described in **Supplementary Information section 2.12**) suggest substantial genetic overlap between the lead SNPs of our primary GWAS and those of our supplementary GWAS. 72 of the 124 general-risk-tolerance lead SNPs are also lead SNPs, or in LD ($r^2 > 0.1$) with a lead SNP, for at least one of the other main phenotypes. The overlap is particularly large between general risk tolerance and adventurousness: 45 of the 124 general-risk-tolerance lead SNPs are also lead SNPs, or in LD with a lead SNP ($r^2 > 0.1$), for adventurousness. We also found that 49 of the 124 general-risk-tolerance lead SNPs are also lead SNPs, or in LD with a lead SNP ($r^2 > 0.1$), of one of the four risky behaviors or their first PC.

To benchmark the likelihood of observing such substantial genetic overlap between the lead SNPs of our primary and supplementary GWAS, we conducted a resampling exercise under the null hypothesis that the lead SNPs of our supplementary GWAS are distributed randomly across the genome and independently from those of the other GWAS. The resampling exercise involved 10,000 runs. In each run, for each lead SNP of the supplementary GWAS we randomly selected an autosomal SNP matched on MAF (with five-percentage-point MAF windows). We then counted how many of the 124 general risk tolerance lead loci contain at least one of the resampled lead SNP, and took the average count across the 10,000 runs. Our results imply that, under the null hypothesis, we would expect 10.3 of the 124 general-risk-tolerance loci to also contain a lead SNP for at least one of the supplementary phenotypes; 2.2 to also contain a lead SNP for adventurousness; and 8.1 to also contain a lead SNP for at least one of the four risky behaviors or their first PC. In all three cases, we strongly reject the null hypothesis ($P < 0.0001$ in all three cases). The overlap we observe between the 124 general risk tolerance loci and the lead SNPs from the other GWAS is thus much greater than what could be expected by chance.

##### 3.3.4 Novelty of the lead SNPs of the discovery GWAS of general risk tolerance

We assessed the novelty of the lead SNPs by performing a lookup in the NHGRI-EBI GWAS Catalog database[49] for each of the 124 general-risk-tolerance lead SNPs (and the SNPs in LD, $r^2 > 0.1$), searching specifically for previous associations with general risk tolerance (and similar phenotypes) (see **Supplementary Information section 2.10** for details). We found that the GWAS Catalog database contained no previous associations with general risk tolerance (or similar phenotypes).

However, we know of one previous study by Day *et al.*[50] that analyzed the same general-risk-tolerance phenotype measure (which they refer to as "risk-taking propensity") as the one we study, in the first release of the UKB data. Day *et. al* report one independent lead SNP (rs4856591) associated with general risk tolerance in the gene *CADM2*. This association was replicated by Boutwell *et al.*[52] using a proxy SNP available in an independent sample (rs1865251) located ~125 kb from the original lead SNP (i.e., rs4856591).

The original lead SNP from ref.[50] is not available in our main reference panel (and therefore not available in the summary statistics from our discovery GWAS), but the proxy SNP from ref.[52] is available. Our discovery GWAS also contains a proximate SNP (rs4856590) only 22 bp away from the original lead SNP from ref.[50]. Both rs1865251 and rs4856590 are genome-wide significant in our discovery GWAS. They are "clumped" with our top lead SNP rs993137 ($P = 2.14\times10^{-40}$), and the LD ($r^2$) between our lead SNP rs993137 and rs1865251 and rs4856590 is 0.996 and 0.949, respectively. Because its locus is associated with general risk tolerance in a previous peer-reviewed publication, we do not consider our lead SNP rs993137 to be a novel association.

We also know of one concurrent GWAS by Strawbridge *et al.*[51] on general risk tolerance (which Strawbridge *et al.* refers to as "risk-taking behavior"). Strawbridge *et al.* also analyzed the same phenotype measure as the one we and Day *et al.*[50] study, but used a somewhat different set of individuals than Day *et al.* in the first release of the UKB data. They identified one independent lead SNP on chromosome 6 (i.e., rs9379971) in addition to the previous association in *CADM2* from ref.[50] and ref.[52]. The SNP rs9379971 is genome-wide significant in our discovery GWAS, and it is "clumped" with a lead SNP of the discovery GWAS (rs1417998, $P = 2.92\times10^{-10}$).

Thus, to the best of our knowledge, all of the loci we identified through our GWAS of general risk tolerance are novel associations with general risk tolerance, except for our top lead SNP rs993137[s]. We therefore consider 123 of the 124 lead SNPs to be newly identified loci for general risk tolerance (and similar phenotypes). We report the novelty of our 124 lead SNPs in the column "New locus" in **Supplementary Table 3.1**.

##### 3.3.5 Results of the GWAS Catalog lookup of the lead SNPs of the primary GWAS of general risk tolerance

To investigate the potential overlap between general risk tolerance and other phenotypes at the loci identified in the discovery GWAS, we performed a lookup for each of our 124 lead SNPs (and the SNPs in LD, $r^2 > 0.6$) in the NHGRI-EBI GWAS Catalog database[49] (see **Supplementary Information section 2.11** for details). The results are reported in

[s] Since we consider the GWAS by Strawbridge *et al.*[51] to be concurrent, we consider our lead SNP rs1417998 to be a novel association with general risk tolerance.

**Supplementary Table 3.3**. In total, we found 61 overlaps between our lead SNPs (and the SNPs in LD, $r^2 > 0.6$) and the previous associations reported in the GWAS Catalog database. Some of our lead SNPs overlap with multiple previous associations: the 61 overlaps involve only 27 of our 124 lead SNPs.

Notably, we found overlaps at five schizophrenia loci, one bipolar disorder locus, four educational attainment loci, and two loci associated with cognitive function and information processing speed. The multiple overlaps with schizophrenia are consistent with the 16 second-stage hits we obtained for schizophrenia in the proxy-phenotype analysis (**Supplementary Table 8.1**), as well as with our finding of strong joint enrichment for association with schizophrenia ($P = 3.0\times10^{-9}$) among our general-risk-tolerance lead SNPs on a non-parametric Mann-Whitney test (**Supplementary Information section 8**). Further, 70% of the 122 general-risk-tolerance lead SNPs available in the schizophrenia summary statistics have concordant signs for the two phenotypes ($P = 8.3\times10^{-6}$). (We also note that general risk tolerance and schizophrenia are moderately and positively genetically correlated ($\hat{r}_g = 0.173$, $SE = 0.021$; **Supplementary Information section 7**)). However, we caution that it is possible that this overlap with the schizophrenia results is primarily attributable to the fact that the schizophrenia GWAS[69] was relatively well-powered, rather than to shared genetic etiology between risk tolerance and schizophrenia.

In addition, we found overlaps at one ADHD locus, one extraversion locus, and one brain volume (superior frontal gyrus grey matter volume) locus. Other potentially interesting overlaps were found at a resting heart rate locus and a motion sickness locus. Several traits associated with autoimmune disease also overlap with our lead SNPs: two loci associated with cholangitis, one locus associated with ulcerative colitis/atopic dermatitis, and one locus associated with type 1 diabetes (that locus is also associated with height, age at menarche, and male pattern baldness). We also found overlaps at one locus associated with age at menarche, at two loci associated with carcinoma, and at one locus associated with breast cancer.

The remaining overlaps are with seemingly unrelated traits, for example at loci associated with tooth development (one locus), gut microbiota diversity (one locus), blood protein levels (one locus), blood pressure and type 2 diabetes (one locus), and ear infection (one locus). The genetic overlap of general risk tolerance with a large number of traits underlines the possibility of widespread pleiotropy in the human genome.

We note that both the existence and absence of overlaps between any two phenotypes should be interpreted with care. The existence of overlaps is not necessarily evidence of a shared genetic architecture. For example, a shared tagging variant could be in LD with two different causal variants that each only cause one of the traits[70]. Further, the absence of overlaps may be the result of inadequate statistical power in the currently available GWAS for either phenotype.

##### *3.3.6 General-risk-tolerance lead SNPs in long-range LD regions*

8 of the 15 long-range LD regions identified by Price *et al.*[44] (**Supplementary Information section 2.9**) contain lead SNPs for at least one of our primary or supplementary GWAS (of general risk tolerance, adventurousness, the four risky behaviors, and their first PC). We focus this section on the two long-range LD regions that contain general-risk-tolerance lead SNPs and that are notable because they also contain lead SNPs for all or most of our other six main GWAS phenotypes: the long-range LD regions ~83.4 to 86.9 Mb on chromosome 3 (**Extended Data**

**Fig. 3.4**) and ~25.3 to 33.4 Mb on chromosome 6 (**Extended Data Fig. 3.4**). These two regions are both among the five notable genomic regions we highlighted in **Supplementary Information section 3.2**. Those two regions contain 10 of the 124 general-risk-tolerance lead SNPs. We also report a third long-range LD region (~135.5 to 138.8 Mb on chromosome 5) that contains two general-risk-tolerance lead SNPs (but no lead SNPs for the other phenotypes), and below in **Supplementary Information section 3.4** we present the five remaining long-range LD regions that we identified in our other six main GWAS. Only one long-range LD region (i.e., chromosome 3, ~83.4 to 86.9 Mb, **Extended Data Fig. 3.4**) contains lead SNPs for all seven of our phenotypes.

The first notable long-range LD region, spanning ~83.4 to 86.9 Mb on chromosome 3, is displayed in a local Manhattan plot in **Extended Data Fig. 3.4** and was discussed in **Supplementary Information section 3.2**. The region is noteworthy because it contains more than one lead SNP for all of our GWAS, and it contains six lead SNPs for general risk tolerance, of which three are located within the *CADM2* gene (~85.0 to 86.2 Mb), two are located ~76 kb and ~115 kb upstream of *CADM2*, and one is located ~547 kb downstream of *CADM2,* closer to the gene *VGLL3*. (The gene *VGLL3*, ~86.9 to 87.0 Mb on chromosome 3, is ~69.8 kb downstream of the long-range LD region.) As we discussed in **Supplementary Information section 3.2**, the long-range LD region contains only one gene—*CADM2*, which covers ~1.2 Mb of the ~3.5 Mb long-range LD region. *CADM2* was the most significantly associated gene in the MAGMA gene analysis (Bonferroni-corrected $P = 1.09\times10^{-50}$, **Supplementary Table 12.3**), and it contains our strongest association with general risk tolerance (rs993137, $P = 2.14\times10^{-40}$). The GWAS Catalog database[49] reports many genome-wide significant associations within the long-range LD region, including age at menarche, BMI, educational attainment, and information processing speed. Most of the previous associations within the long-range LD region are annotated to *CADM2*, which encodes a synaptic cell adhesion molecule[55], and is related to synapse formation[56] and brain plasticity[57].

The second long-range LD region (~25.3 to 33.4 Mb on chromosome 6) covers all the Human Leukocyte Antigen (HLA) genes[55]. It was also discussed in **Supplementary Information section 3.2**, and it is displayed in a local Manhattan plot in **Extended Data Fig. 3.4**. It contains four lead SNPs located in (or in close proximity to) four different genes: *HIST1H2AC*, *HIST1H2BD*, *TRIM27*, and *C4B*, of which the first three are significant in the MAGMA gene analysis after Bonferroni-correction (**Supplementary Table 12.3**). The region is notable because it contains lead SNPs for all our other main GWAS, except drinks per week for which we identified a suggestively associated SNP (rs6937318, $P = 3.83\times10^{-7}$). As we discussed in **Supplementary Information section 3.2**, the region contains ~250 genes, and the MAGMA gene analysis found ~30 significant genes in the region after Bonferroni-correction (however, none of the actual HLA genes are significant, **Supplementary Table 12.3**). The GWAS Catalog database[49] reports more than a thousand genome-wide significant associations across hundreds of traits within the long-range LD region (~25.3 to 33.4 Mb on chromosome 6), including Alzheimer's disease, autism spectrum disorder, educational attainment, and schizophrenia.

We also identified a third long-range LD region (~135.5 to 138.8 Mb on chromosome 5) that contains two lead SNPs associated with general risk tolerance within the genes *CTNNA1* and *ETF1*, but the region does not contain lead SNPs for any of our other main GWAS.

##### *3.3.7 General-risk-tolerance lead SNPs in candidate inversions*

Of the 154 genomic segments deemed highly prone to inversion polymorphisms (i.e., the 154 "candidate inversions," described in **Supplementary Information section 2.9**), we identified 44 that contain lead SNPs for at least one of our GWAS. 13 of these 44 candidate inversions contain a total of 30 general-risk-tolerance lead SNPs. This section focuses on four of the 13 candidate inversions that we find noteworthy because they contain lead SNPs for general risk tolerance as well as for most of our other main GWAS. Three of these four candidate inversions are among the five notable genomic regions we highlighted in **Supplementary Information section 3.2**. (Below, in **Supplementary Information section 3.4,** we discuss three other of the 44 candidate inversions of the 44, which are notable because they are shared across four of our GWAS, excluding general risk tolerance.) The four candidate inversions that we discuss in this section span ~124.6 to 132.7 Mb on chromosome 7 (**Extended Data Fig. 3.4**), ~7.89 to 11.8 Mb on chromosome 8 (**Extended Data Fig. 3.4**), ~70.1 to 74.4 Mb on chromosome 16, and ~49.1 to 55.5 Mb on chromosome 18 (**Extended Data Fig. 3.4**). Only one of these candidate inversions (i.e., ~49.1 to 55.5 Mb on chromosome 18, **Extended Data Fig. 3.4**) contains lead SNPs for all our GWAS.

The first candidate inversion is on chromosome 7 (~124.6 to 132.7 Mb). We discussed it in **Supplementary Information section 3.2** and we display it in a local Manhattan plot in **Extended Data Fig. 3.4**. This candidate inversion is notable because it contains lead SNPs for all our GWAS, except automobile speeding propensity (for which the strongest association is almost genome-wide significant (rs141450, $P = 7.88\times10^{-8}$)). The candidate inversion contains one lead SNP for general risk tolerance, two lead SNPs for adventurousness, one for drinks per week, one for ever smoker, one for number of sexual partners, and one for the first PC of the risky behaviors. As we discussed in **Supplementary Information section 3.2**, the candidate inversion contains ~50 genes, and five of those were significant after Bonferroni correction in the MAGMA gene analysis (**Supplementary Table 12.3**). The GWAS Catalog database[49] reports previous associations with traits such as alcohol dependence, educational attainment, and schizophrenia.

The second candidate inversion spans ~7.89 to 11.79 Mb on chromosome 8. We discussed it in **Supplementary Information section 3.2** and we display it in a local Manhattan plot in **Extended Data Fig. 3.4**. This candidate inversion is notable, because it contains associations for all our GWAS, except drinks per week and the first PC of the risky behaviors. (The strongest association with drinks per week within the candidate inversion is rs574968044, $P = 5.64\times10^{-4}$, and the strongest association with the first PC of the risky behaviors is rs2898249, $P = 1.27\times10^{-7}$.) The candidate inversion contains 15 lead SNPs for general risk tolerance, five for adventurousness, one for automobile speeding propensity, two for ever smoker, and four for number of sexual partners. As we discussed in **Supplementary Information section 3.2**, the candidate inversion contains ~20 genes, of which practically all were significant after Bonferroni correction in the MAGMA gene analysis (**Supplementary Table 12.3**). The GWAS Catalog database[49] reports genome-wide associations in the candidate inversion with many behavioral phenotypes, including extraversion, neuroticism, schizophrenia, and chronotype.

The third candidate inversion is on chromosome 18 (~49.1 to 55.5 Mb) and was also discussed in **Supplementary Information section 3.2**. It contains lead SNPs for all our GWAS, and it contains more than one lead SNP for all our GWAS except drinks per week, for which it contains only one. We display the region in a local Manhattan plot in **Extended Data Fig. 3.4**. The candidate inversion contains three lead SNPs for general risk tolerance, two for adventurousness, two for automobile speeding propensity, one for drinks per week, three for ever smoker, eight for number of sexual partners, and four for the first PC of the risky behaviors. As we discussed in **Supplementary Information section 3.2**, among the ~20 genes within the candidate inversion, we find only one gene—*TCF4* (Bonferroni-corrected $P$ = 5.51×10$^{-9}$)—significant after Bonferroni correction in the MAGMA gene analysis (**Supplementary Table 12.3**). *TCF4* plays an important role in nervous system development, and mutations in the gene are known to cause the rare Pitt-Hopkins syndrome[55]. The syndrome is characterized by distinct facial features, intellectual disability, delayed motor skills, and epilepsy, among many other symptoms[59–61]. The GWAS Catalog database[49] reports genome-wide significant associations mapped to *TCF4* with schizophrenia and reports previous genome-wide associations within the candidate inversion with traits such as autism spectrum disorder, ADHD, depression, educational attainment, schizophrenia, and subcortical brain region volumes.

A fourth candidate inversion is on chromosome 16 (~70.1 to 74.4 Mb) and contains more than one lead SNP for the GWAS of general risk tolerance, ever smoker, number of sexual partners, and the first PC of the risky behaviors. (The strongest association with adventurousness within the candidate inversion is rs9929242 ($P$ = 2.21×10$^{-6}$), the strongest association with automobile speeding propensity is rs2158268 ($P$ = 3.35×10$^{-6}$), and the strongest association with drinks per week is rs11648570 ($P$ = 1.50×10$^{-7}$).) The candidate inversion contains ~35 genes, of which 2 are significant after Bonferroni correction in the MAGMA gene analysis (**Supplementary Table 12.3**): *CHST4* (Bonferroni-corrected $P$ = 4.69×10$^{-3}$) and *CMTR2* (Bonferroni-corrected $P$ = 0.024). *CHST4* is involved in normal cell function via carbohydrate sulfotransferase[55], and *CMTR2* is involved in methyltransferase[55]. There is also one additional gene ~458 kb upstream of the candidate inversion—*NFAT5*—that is significant after Bonferroni correction in the MAGMA gene analysis (Bonferroni-corrected $P$ = 4.66×10$^{-3}$). It is notable because it plays a central role in gene transcription during immune response[55], consistent with the significant estimate of the category "Immune/Hematopoietic" in the partitioning of the SNP heritability into functional categories with LD Score regression (**Supplementary Information section 12**). The candidate inversion contains genome-wide significant associations in the GWAS Catalog database[49] with total cholesterol, prostate cancer, and stroke, among other phenotypes.

The remaining nine of the 13 candidate inversions that contain general-risk-tolerance lead SNPs each contain only one general-risk-tolerance lead SNP, and the overlap with the other GWAS is low.

###### *3.3.8 General-risk-tolerance lead SNPs in LD with 1000 Genomes structural variants*

In addition, 24 of the 124 lead SNPs are located within, or in strong LD with another variant within, 23 different 1000G structural variants. All candidate inversions that contain general-risk-tolerance lead SNPs, as well as these 1000G structural variants, are reported in **Supplementary Table 3.1**.

#### 3.4 Results of the supplementary GWAS

Our six supplementary GWAS—of adventurousness, the four risky behaviors, and of the first PC of the four risky behaviors—identified a total of 741 lead SNPs. We consider 730 of these 741 lead SNPs to be novel associations for these phenotypes. The association results are reported in **Supplementary Table 3.2**. We identified 167 lead SNPs for adventurousness, 42 for automobile speeding propensity, 85 for drinks per week, 223 for ever smoker, 118 for number of sexual partners, and 106 for the first PC of the risky behaviors.

The results are displayed in Manhattan plots in **Extended Data Fig. 3.1** and in Q-Q plots in **Extended Data Fig. 3.2**. The genomic inflation factors ($\lambda_{GC}$) across the phenotypes ranges from 1.254 to 1.470, and the estimated LD Score regression intercepts were in the range of 1.026 to 1.051, consistent with polygenicity and low levels of confounding from population stratification[42,68]. The estimated effect sizes (the $\hat{\beta}_j$'s) reported in **Supplementary Table 3.2** are expressed in phenotype standard deviation units per effect-coded allele. (The phenotype standard deviations are reported in **Supplementary Table 1.2**).

##### *3.4.1 GWAS of adventurousness*

We identified 167 lead SNPs associated with adventurousness. The strongest associations were found within, or in close proximity to, the genes *CADM2* (rs10433500, $P = 9.31\times10^{-84}$), *SATB1* (rs13090941, $P = 3.26\times10^{-21}$), and *FOXP2* (rs10228494, $P = 1.23\times10^{-19}$). 47 of the 167 lead SNPs are also lead SNPs, or in LD with a lead SNP ($r^2 > 0.1$), for general risk tolerance. 52 of the 167 lead SNPs are also lead SNPs, or in LD with a lead SNP ($r^2 > 0.1$), for the four risky behaviors, and their first PC.

To the best of our knowledge, there are no previous studies with genome-wide significant associations for adventurousness, and we consider all of our lead SNPs to be novel associations. We report the novelty of our adventurousness lead SNPs (and of the lead SNPs of the other supplementary GWAS) in the column "New locus" in **Supplementary Table 3.2**.

18 of the 167 lead SNPs are distributed across three of the 15 long-range LD regions identified by Price *et al.*[44], of which two are the long-range LD regions that we describe above in **Supplementary Information section 3.2** on chromosome 3 (~83.4 to 86.9 Mb) and chromosome 6 (~25.3 to 33.4 Mb). Remarkably, 16 of these 18 lead SNPs are located within the long-range LD region on chromosome 3. Most of these 16 SNPs are located within, or in proximity to, the *CADM2* gene, but three of them are closer to the gene *VGLL3* (which is located ~69.8 kb downstream of the long-range LD region). The third long-range LD region, on chromosome 8 (~111.9 to 114.9 Mb), does not contain lead SNPs for any of our other six main GWAS.

There is also a total of 24 lead SNPs located within 16 candidate inversions, as well as 29 lead SNPs located within, or in strong LD with another variant within, 29 different 1000G structural variants. All candidate inversions that contain adventurousness lead SNPs, as well as these 1000G structural variants, are reported in **Supplementary Table 3.2**.

##### *3.4.2 GWAS of automobile speeding propensity*

We identified 42 lead SNPs associated with automobile speeding propensity. The strongest association was found within the gene *CADM2* (rs17516256, $P = 9.9\times10^{-21}$). Seven of the 42 lead SNPs are also lead SNPs, or in LD with a lead SNP ($r^2 > 0.1$), for general risk tolerance. 17 of the 42 lead SNPs are also lead SNPs, or in LD with a lead SNP ($r^2 > 0.1$), for at least one of the other main phenotypes we analyze, excluding the first PC of the risky behaviors.

To the best of our knowledge, there are no previous studies with genome-wide significant associations for automobile speeding propensity, so we consider all of our lead SNPs to be novel associations (**Supplementary Table 3.2**).

Eight of the 42 lead SNPs are distributed across two of the long-range LD regions identified by Price *et al.*[44]. Those two regions are the two long-range LD regions we described above in **Supplementary Information section 3.2** and that are shared across general risk tolerance and all or most of our GWAS. Three of the eight lead SNPs are in the region on chromosomes 3 (~83.4 to 86.9 Mb), and the other five are in the region on chromosome 6 (~25.3 to 33.4 Mb). The three lead SNPs located in the long-range LD region on chromosome 3 (~83.4 to 86.9 Mb) are all within the *CADM2* gene.

There is also a total of 24 lead SNPs located within 16 candidate inversions, and seven lead SNPs are located within, or in strong LD with another variant within, 7 different 1000G structural variants. All candidate inversions that contain automobile speeding propensity lead SNPs, as well as these 1000G structural variants, are reported in **Supplementary Table 3.2**.

##### *3.4.3 GWAS of drinks per week*

We identified 85 lead SNPs associated with drinks per week. The strongest and most notable of these associations is located within the alcohol dehydrogenase 1B gene on chromosome 4 (*ADH1B*, rs1229984, $P = 7.8\times10^{-202}$). Multiple genome-wide significant associations were also found within, or in close proximity to, other alcohol dehydrogenase genes—*ADH1A* (rs62307263, $P = 3.42\times10^{-11}$), *ADH1C* (rs113659074, $P = 9.93\times10^{-23}$), and *ADH7* (rs114112910, $P = 1.08\times10^{-8}$). Four of the 85 lead SNPs are also lead SNPs, or in LD with a lead SNP ($r^2 > 0.1$), for general risk tolerance. 21 of the 85 lead SNPs are also lead SNPs, or in LD with a lead SNP ($r^2 > 0.1$), for at least one of the other main phenotypes we analyze, excluding the first PC of the risky behaviors.

Following the procedure described in **Supplementary Information section 2.10**, we found that the lead SNPs located within, or in close proximity to, the genes *ADH1A*, *ADH1B*, *ADH1C*, *ADH7*, *GCKR*, and *KLB,* have previously been associated with alcohol consumption (or similar phenotypes), and we therefore do not consider associations annotated to these genes to be novel associations with drinks per week (**Supplementary Table 3.2**).

Two of the 85 lead SNPs are located within the same long-range LD region on chromosome 3 (~83.4 to 86.9 Mb) that contains lead SNPs for all our GWAS phenotypes (**Supplementary Information section 3.2**), and both lead SNPs are located within *CADM2*. Unlike for the six other main phenotypes we analyze, there are no lead SNPs for drinks per week in the long-range LD region spanning ~25.3 to 33.4 Mb on chromosome 6, but the region contains a suggestive association at rs6937318 ($P$ value = $3.83\times10^{-7}$). However, as can be seen in **Extended Data Fig. 3.4**, the general level of association is much lower for drinks per week in that long-range LD region in comparison with the other main phenotypes we analyze.

15 lead SNPs are located across 15 candidate inversions. Of these, none is located within the candidate inversion spanning ~7.89 to 11.8 Mb on chromosome 8, unlike for general risk tolerance, adventurousness, ever smoker, and number of sexual partners. (The strongest association with drinks per week within that candidate inversion is rs574968044, ($P = 5.64\times10^{-4}$).) Just as for the long-range LD region on chromosome 6 (~25.3 to 33.4 Mb), the general level

of association is much lower for drinks per week in the candidate inversion on chromosome 8 in comparison with the other main phenotypes, as can be seen in **Extended Data Fig. 3.4**.

15 lead SNPs are located within, or in strong LD with another variant within, 20 different 1000G structural variants (some lead SNPs are in strong LD with SNPs in multiple structural variants). All candidate inversions that contain drinks per week lead SNPs, as well as these 1000G structural variants, are reported in **Supplementary Table 3.2**.

##### *3.4.4 GWAS of ever smoker*

We identified 223 lead SNPs associated with ever smoker. The strongest associations are located within, or in close proximity to, the genes *NCAM1* (rs7938812, $P = 7.1\times10^{-48}$), *ZEB2* (rs961414, $P = 6.99\times10^{-28}$), *REV3L* (rs240955, $P = 3.84\times10^{-23}$), *NT5C2* (rs7092200, $P = 7.44\times10^{-22}$), *CLU* (rs11783093, $P = 7.44\times10^{-22}$), *TMEM182* (rs1368550, $P = 1.08\times10^{-21}$), and *CADM2* (rs34495106, $P = 2.23\times10^{-20}$). There are three additional lead SNPs located within, or in close proximity to, *CADM2*. 22 of the 223 lead SNPs are also lead SNPs, or in LD with a lead SNP ($r^2 > 0.1$), for general risk tolerance. 59 of the 223 lead SNPs are also lead SNPs, or in LD with a lead SNP ($r^2 > 0.1$), for at least one of the other main phenotypes we analyze, excluding the first PC of the risky behaviors.

Following the procedure described in **Supplementary Information section 2.10**, we found that the two lead SNPs located within the genes *NCAM1* and *BDNF* have previously been associated with smoking behavior. We therefore do not consider those two loci to be novel associations with ever smoker (**Supplementary Table 3.2**).

Previous GWAS on nicotine dependence[71] and number of cigarettes per day (CPD)[29,71] report genes in the nicotine receptor gene family (i.e., the *CHRN* family[29]) as some of their strongest associations. In our results, none of the nicotine receptor genes (*CHRN*) contained any lead SNPs, consistent with the theory that the ever smoker phenotype is more strongly mediated by risk tolerance and social influences than by vulnerability to nicotine addiction[72].

14 lead SNPs are located in six of the long-range LD regions identified by Price *et al.*[44]. These include the long-range LD regions ~83.4 to 86.9 Mb on chromosome 3 and ~25.3 to 33.4 Mb on chromosome 6, which were also found to contain lead SNPs for general risk tolerance and most of our other GWAS phenotypes (**Supplementary Information section 3.2)**. The four remaining long-range LD regions are on chromosome 1 (~48.2 to 52.2 Mb), chromosome 2 (~134.7 to 138.2 Mb), chromosome 12 (~111.0 to 113.5 Mb), and chromosome 20 (~32.5 to 35.0 Mb).

24 lead SNPs are located within 16 candidate inversions, and there are 56 lead SNPs located within, or in strong LD with another variant within, 69 different 1000G structural variants (some lead SNPs are in strong LD with SNPs in multiple structural variants). All candidate inversions that contain ever smoker lead SNPs, as well as these 1000G structural variants, are reported in **Supplementary Table 3.2**.

We also performed a replication of the eight genome-wide significant SNPs from the GWAS of ever smoker by the TAG consortium[29] in our GWAS in the UKB only (and not in our meta-analyses of our UKB GWAS with the TAG summary statistics, as our replication sample must be independent of the TAG sample). All eight SNPs have concordant signs, and all are highly significant. Specifically, six of the eight SNPs are genome-wide significant ($P$ value $< 5\times10^{-8}$) in our GWAS of ever smoker in the UKB (rs6265, $P = 1.1\times10^{-14}$; rs4923457, $P = 2.3\times10^{-11}$;

rs4923460, $P = 1.8\times10^{-11}$; rs4074134, $P = 2.6\times10^{-11}$; rs6484320, $P = 2.1\times10^{-11}$; rs879048, $P = 1.2\times10^{-11}$), and the two remaining SNPs are suggestively significant (rs1013442, $P = 1.5\times10^{-7}$; rs1304100, $P = 1.2\times10^{-7}$). It should be noted that the eight SNPs reported in the TAG consortium results are not independent, and if we apply our locus definition from **Supplementary Information section 2.8,** then the SNP rs6265 would be the only lead SNP, and the other seven SNPs would be clumped to that SNP. rs6265 is one of the lead SNPs in our GWAS of ever smoker (which combines the UKB GWAS with the TAG GWAS; $n$ = 518,633), and its $P$ value in that GWAS is $3.92\times10^{-18}$.

##### *3.4.5 GWAS of number of sexual partners*

We identified 118 lead SNPs associated with number of sexual partners, and the strongest associations are located within, or in close proximity to, the genes *C14orf177* ($P = 4.61\times10^{-19}$) and *FURIN* ($P = 5.76\times10^{-17}$). Two lead SNPs are within, or in close proximity to, *CADM2* (rs2163971, $P = 4.6\times10^{-14}$; rs9856718, $P = 2.68\times10^{-8}$). 28 of the 118 lead SNPs are also lead SNPs, or in LD with a lead SNP ($r^2 > 0.1$), for general risk tolerance. 63 of the 118 lead SNPs are also lead SNPs, or in LD with a lead SNP ($r^2 > 0.1$), for at least one of the other main phenotypes we analyze, excluding the first PC of the risky behaviors.

Following the procedure described in **Supplementary Information section 2.10**, we determined that none of the loci we found to be associated with number of sexual partners have been previously associated with any similar phenotype, and we therefore consider all of our associations with number of sexual partners to be novel (**Supplementary Table 3.2**).

Six lead SNPs are located within the two long-range LD regions on chromosome 3 (~83.4 to 86.9 Mb) and chromosome 6 (~25.3 to 33.4 Mb) that are shared across general risk tolerance and all or almost all other GWAS (see **Supplementary Information section 3.2**).

25 lead SNPs are located within 14 candidate inversions, and 23 lead SNPs are located within, or in strong LD with another variant within, 33 different 1000G structural variants (some lead SNPs are in strong LD with SNPs in multiple structural variants). All candidate inversions that contain lead SNPs for number of sexual partners, as well as these 1000G structural variants, are reported in **Supplementary Table 3.2**.

##### *3.4.6 GWAS of the first PC of the four risky behaviors*

We identified 106 lead SNPs associated with the first PC of the four risky behaviors. The strongest association is located in the gene *MAPT* on chromosome 17 (rs62062288, $P = 1.02\times10^{-29}$), and that gene contains lead SNPs for the GWAS of automobile speeding propensity, drinks per week, and number of sexual partners (but not for general risk tolerance, adventurousness, or ever smoker). The second strongest association is in *CADM2* (rs6790699, $P = 5.99\times10^{-27}$), our top gene associated with general risk tolerance, which also contains lead SNPs for all our main GWAS. The three next strongest associations are in *NCAM1* (rs2155290, $P = 4.11\times10^{-24}$, top locus for ever smoker), *ADH1B* (rs1229984, $P = 3.70\times10^{-22}$, top locus for drinks per week), and *FOXP1* (rs4676964, $P = 7.80\times10^{-18}$). This latter gene (*FOXP1*) is also strongly associated with the phenotypes automobile speeding propensity and number of sexual partners.

18 of the 106 lead SNPs are also lead SNPs, or in LD with a lead SNP ($r^2 > 0.1$), for general risk tolerance. 89 of the 106 lead SNPs are also lead SNPs, or in LD with a lead SNP ($r^2 > 0.1$), for at

least one of the other main phenotypes we analyze. We note that a high level of overlap with the results of the GWAS of the four risky behaviors was to be expected, given that the first PC of the risky behaviors was obtained from a principal component analysis of the four risky behaviors.

We know of no previous GWAS of the first PC of the risky behaviors (or of similar phenotypes), and we consider all of our lead SNPs to be novel associations (**Supplementary Table 3.2**).

Seven of the 106 lead SNPs are located within the long-range LD region on chromosome 3 (~83.4 to 86.9 Mb), and three lead SNPs are located within the long-range LD region on chromosome 6 (~25.3 to 33.4 Mb), both shared with the general-risk-tolerance GWAS and all or most of our other main GWAS (see **Supplementary Information section 3.2**). One additional lead SNP is located within a long-range LD region on chromosome 2 (~134.7 to 138.2 Mb).

20 lead SNPs are located within 14 candidate inversions. Of these, none is located within the chromosome 8 candidate inversion spanning ~7.89 to 11.8 Mb. The strongest association with the first PC of the risky behaviors within that candidate inversion is rs2898249 ($P = 1.27\times10^{-7}$), which we consider suggestive. Also, as can be seen from **Extended Data Fig. 3.4**, the general level of association within the candidate inversion on chromosome 8 (~7.89 to 11.8 Mb) is more similar to the phenotypes for which there are lead SNPs in the candidate inversion than to drinks per week (for which the candidate inversion does not contain any lead SNPs).

20 lead SNPs are located within, or in strong LD with another variant within, 24 different 1000G structural variants (some lead SNPs are in strong LD with SNPs in multiple structural variants).

###### *3.4.7 Long-range LD regions and candidate inversions that contain lead SNPs for the supplementary GWAS phenotypes*

Our seven main GWAS identified lead SNPs located in eight long-range LD regions and 44 candidate inversions (defined in **Supplementary Information section 2.9**). In **Supplementary Information section 3.2**, we focused on the two long-range LD regions and on the three candidate inversions that contain lead SNPs for general risk tolerance and for all or most of our six other GWAS phenotypes, and above in this subsection (**Supplementary Information section 3.4**) we have briefly presented the long-range LD regions and candidate inversions that contain lead SNPs for each of the six supplementary phenotypes. We now focus on three candidate inversions that are notable because they contain lead SNPs for four of the six other GWAS (but not for general risk tolerance). The remaining 37 identified candidate inversions and the remaining six long-range LD regions have little overlap across the GWAS and are reported in **Supplementary Tables 3.1** and **3.2**.

The first of the three additional candidate inversions that are notable because they are shared across four supplementary GWAS is located on chromosome 11 (~112.5 to 113.5 Mb). It contains one lead SNP for drinks per week, four for ever smoker, one for number of sexual partners, and one for the first PC of the risky behaviors. The strongest association with general risk tolerance within the candidate inversion is rs78168664 ($P = 1.47\times10^{-4}$). The GWAS Catalog database [49] reports a few previous phenotypes with associations in the region, which include bone ultrasound measurement, cardiac muscle measurement, and gut microbiota.

The second additional candidate inversion is on chromosome 16 (~12.0 to 14.8 Mb) and contains one lead SNP for drinks per week, one lead SNP for ever smoker, one lead SNP for number of sexual partners, and one lead SNP for the first PC of the four risky behaviors. The strongest association with general risk tolerance within the candidate inversion is rs2866323 ($P = 1.36\times10^{-5}$). The GWAS Catalog database[49] reports notable previous associations with traits such as age at menarche, human standing height, and schizophrenia.

The third candidate inversion is on chromosome 17 (~43.6 to 44.3 Mb) and contains one lead SNP for automobile speeding propensity, one for drinks per week, one for number of sexual partners, and two lead SNPs for the first PC of the risky behaviors. The strongest association with general risk tolerance within the candidate inversion is rs2866323 ($P = 1.86\times10^{-4}$). The candidate inversion covers only four genes, including the gene *MAPT* (~43.9 to 44.1 Mb on chromosome 17). SNPs in and around *MAPT* have previously been associated with many neurodegenerative disorders, including Alzheimer's disease, Parkinson's disease and frontotemporal dementia[55]. However, most of those previous associations are located outside the candidate inversion on chromosome 17 (~43.6 to 44.3 Mb), and *MAPT* is not significant after Bonferroni correction in the MAGMA gene analysis (**Supplementary Table 12.3**).

##### *3.4.8 GWAS Catalog lookup of the lead SNPs from the supplementary GWAS*

To investigate the overlap of our six supplementary GWAS phenotypes with previous GWAS of other phenotypes, we performed a lookup for each of the lead SNPs (and the SNPs in LD, $r^2 > 0.6$) in the NHGRI-EBI GWAS Catalog database[49], as described in **Supplementary Information section 2.11**. We report the results in **Supplementary Table 3.3**. For the six GWAS, we find in total 939 overlaps distributed across 130 lead SNPs. Since the six phenotypes are genetically correlated, they tend to share lead SNPs (or loci where the lead SNPs for each phenotype are in LD, $r^2 > 0.1$), and as expected we find similar overlaps with other traits across the six phenotypes. Each of the six phenotypes have at least one overlap with all of the following phenotypes or phenotype categories: schizophrenia; educational attainment or cognitive performance (e.g. intelligence, information processing speed, cognitive function); BMI (body mass index) (or related anthropometric traits); circulating lipids (e.g. triglycerides, cholesterol, lipids); blood pressure or coronary artery disease (including arterial stiffness); and lung disease (e.g. interstitial lung disease, COPD, asthma).

Most of the six GWAS also have lead SNPs that overlap with the following phenotypes or phenotype categories: autoimmune diseases (e.g. Crohn's, rheumatoid arthritis, psoriasis, vitiligo); various cancers (e.g. breast cancer, lung cancer) or tumor formation; brain volume; bipolar disorder; age at menarche; Alzheimer's disease (driven by the *MAPT*-locus located in a candidate inversion on chromosome 17, ~43.6 to 44.3 Mb); height; Parkinson's disease; and attention hyperactivity deficit disorder.

The amount of overlap with schizophrenia seems particularly striking: in total, 19 distinct loci from our six additional GWAS overlap with loci associated with schizophrenia. We also observed many overlaps between general risk tolerance and schizophrenia (**Supplementary Information section 3.1**).

As discussed in **Supplementary Information section 3.3**, we note that both the existence and the absence of overlaps between any two phenotypes should be interpreted with care, and that the existence of overlaps is not necessarily evidence of shared genetic architecture[70].

### 4 Testing for population stratification

Population stratification can be an important source of bias in GWAS. As per our analysis plan[30], every cohort in our GWAS included the ten (or more) top principal components (PCs) of the genetic-relatedness matrix (GRM) in their analyses, and some also used mixed-linear models. These procedures should, in principle, control for population stratification. Nonetheless, some population stratification and confounding bias could still remain after these controls[73].

In this section, we report the results of two tests of population stratification that are based on two different sets of assumptions. The first test is the LD Score intercept test[42], which allows us to quantify the amount of stratification that is present in our estimates. The second is a sign test that compares the signs of GWAS estimates to the signs of the estimates of a within-family (WF) GWAS in an independent replication cohort. We applied the first test to the summary statistics of the discovery and replication GWAS of self-reported general risk tolerance and to the summary statistics of the supplementary GWAS; we applied the second test to the summary statistics of the discovery GWAS of self-reported general risk tolerance.

#### 4.1 LD Score intercept test

The LD Score intercept test uses GWAS summary statistics for all measured SNPs. Unlike the Genomic Control (GC) method, which assumes that confounding bias (e.g., due to population stratification and cryptic relatedness) is responsible for inflation in the GWAS $\chi^2$ statistics, the LD Score regression method can disentangle inflation that is due to true polygenic signal throughout the genome (which affects the slope of the LD Score regression) from inflation that is due to confounding biases such as cryptic relatedness and population stratification (which affects the intercept of the regression).

We used the LDSC software [42] to estimate the intercepts in LD Score regressions with the summary statistics of our discovery and replication GWAS of general risk tolerance. We also estimated LD Score regressions with the summary statistics from the supplementary GWAS.

For each phenotype, we used the "eur_w_ld_chr/" files of LD scores computed by Finucane *et al.*[74] and made available on https://github.com/bulik/ldsc/wiki/Genetic-Correlation, accessed on March 14, 2016. These LD scores were computed with genotypes from the European-ancestry samples in the 1000 Genomes Project; only HapMap3 SNPs with MAF > 0.01 were included in the LD Score regressions. Because GC will tend to bias the intercept of the LD Score regression downward, we did not apply GC to the summary statistics prior to estimating the LD Score regressions.

**Extended Data Fig. 4.1** shows LD Score regression plots for our discovery and replication GWAS and for the supplementary GWAS.

For our discovery GWAS, we estimated a LD Score intercept of 1.040 ($SE$ = 0.012); for the replication GWAS, we estimated a LD Score intercept of 1.002 ($SE$ = 0.069). The mean $\chi^2$ statistics for all the SNPs in the two LD Score regressions are 1.848 and 1.031, respectively. The mean $\chi^2$ statistics reflect the average strength of the GWAS associations between the SNPs and each phenotype. Under the null hypothesis that there is no confounding bias and that the SNPs have no causal effects on the phenotypes, the mean $\chi^2$ statistics would be 1. Thus, mean $\chi^2$

statistics greater than 1 indicate that some SNPs are associated with the phenotypes, either because they are in LD with causal SNPs or because of confounding bias.

One measure of stratification bias is given by the ratio $\frac{Intercept-1}{\overline{X^2}-1}$, which describes the share of the inflation in the mean $\chi^2$ statistic ($\overline{X^2}$) that is due to stratification. This ratio is 0.048 (*SE* = 0.014) for the discovery GWAS and 0.071 (*SE* = 0.221) for the replication GWAS. These estimates imply that only a small part of the observed inflation in the mean $\chi^2$ statistics from our discovery and replication GWAS is accounted for by confounding bias (due to population stratification, cryptic relatedness, or other confounds) rather than polygenic signal. This suggests that the bulk of the inflation in the $\chi^2$ statistics from the discovery and replication GWAS is attributable to true polygenic signal throughout the genome, and that population stratification is unlikely to be a major concern for the analyses we present in this paper.

**Supplementary Table 4.1** contains the full set of LD Score regression results for the discovery GWAS, the replication GWAS, the meta-analysis of the discovery and replication GWAS, and the supplementary GWAS. Although the LD Score intercepts from these regressions are often significantly larger than 1, the share of inflation in $\overline{X^2}$ that is due to stratification remains small (ranging from 0.047 to 0.071), which allows us to conclude that confounding bias is likely to account for no more than a small part of the inflation in these GWAS' mean $\chi^2$ statistics.

As mentioned in **Supplementary Information section 2.7,** rather than applying the usual GC correction, we followed the increasingly common practice of adjusting the standard errors of our GWAS estimates for the possible effects of population stratification by inflating them by the square root of our estimates of the intercepts from the LD Score regressions[42]. For the discovery and the replication GWAS of general risk tolerance, the meta-analysis of the discovery and replication GWAS of general risk tolerance, and the GWAS of ever smoker, each of which combines several cohorts, we inflated the standard errors at the meta-analysis level only.

#### 4.2 GWAS/WF GWAS sign test for the general-risk-tolerance GWAS

As a simple test of whether the results of our general-risk-tolerance GWAS are driven entirely by stratification or whether they capture some genetic signal, we performed a sign test that compares the signs of the estimates from our discovery GWAS of general risk tolerance (excluding all full siblings from the UKB cohort, as described below) to the signs of the estimates from within-family (WF) GWAS of general risk tolerance in the independent STR1, STR2, and UKB-siblings cohorts. The UKB-siblings cohort was defined in the same way as the full UKB cohort, but only includes individuals with at least one full sibling in the UKB. To avoid overfitting (i.e., to ensure that the two sets of signs originate from GWAS that were conducted using independent cohorts), we reran our discovery GWAS after excluding all individuals with at least one full sibling in the UKB.

If the discovery GWAS estimates are driven by stratification, then they should be independent of the signs of the WF GWAS estimates (which are immune to stratification) and therefore the two sets of signs should only have a concordance of roughly 50%. A significantly higher degree of sign concordance would suggest that at least some of the signal from the GWAS comes from true genetic effects.

##### *4.2.1 Background*

We followed the method outlined in Okbay *et al.* (2016)[18]. Here, we summarize the main assumptions and procedures that are described in more detail in that paper.

We first let $\hat{\beta}_j$ denote the estimate corresponding to SNP $j$ from the discovery GWAS excluding the full siblings from the UKB, and let $\hat{\beta}_{WF,j}$ denote the WF GWAS estimate corresponding to SNP $j$. The WF GWAS estimate is obtained in a sample of sibling pairs, from a regression of sibling differences in general risk tolerance on sibling differences in genotypes and in controls. Since WF regressions are not biased due to stratification, and under the assumption that the WF effect size of each SNP is equal to the true population effect size, we can decompose the GWAS and WF GWAS estimates as:

$$\hat{\beta}_j = \beta_j + s_j + U_j$$
$$\hat{\beta}_{WF,j} = \beta_{WF,j} + V_j,$$

where $\beta_j$ and $\beta_{WF,j}$ are the true underlying GWAS and WF GWAS parameters for SNP $j$, $s_j$ is the bias due to stratification (defined to be orthogonal to $\beta_j$ and $U_j$), and $U_j$ and $V_j$ are the error terms in the estimates due to sampling variation, with $\mathrm{E}(U_j) = \mathrm{E}(V_j) = 0$. Note that if $\hat{\beta}_j$ and $\hat{\beta}_{WF,j}$ are estimated in independent samples, then $U_j$ and $V_j$ will be independent.

Under the null hypothesis that the GWAS contains no genetic signal (i.e., $\beta_j = \beta_{WF,j} = 0$ for all $j$) the sign of $\hat{\beta}_{WF,j}$ will be random with equal odds of being positive or negative and will be independent of the sign of $\hat{\beta}_j$. This means that among a set of $M$ independent SNPs, the number of SNPs that have concordant signs, denoted $C$, follows a binomial distribution:

$$C \sim \text{Binomial}(M, 0.5).$$

We can thus measure the observed sign concordance and use this known distribution to formally test the null hypothesis. We tested this against the one-sided alternative hypothesis that $\beta_{WF,j}$ and $\beta_j$ are not equal to zero and that their estimates thus have concordant signs. We conducted one-sided tests, because there is no reason to suspect that the signs would be discordant.

##### *4.2.2 The GWAS and WF GWAS data*

For this analysis, WF GWAS estimates were obtained in the STR1 (STR-Twingene), STR2 (STR-SALTY), and UKB-siblings cohorts. These estimates were then combined using a sample-size weighted meta-analysis. The STR1, STR2, and UKB-siblings cohorts include 674, 680, and 16,330 sibling pairs, respectively. This gave us a total WF sample size of 17,684 sibling pairs (35,368 individuals).

The GWAS estimates are those from the discovery GWAS excluding all full siblings from the UKB, with a total sample size of 901,908.

Not every SNP from the GWAS results was available in the WF samples. To maximize power, we restricted our SNPs for each sign test to those that were available in both the GWAS results and in the three WF cohorts. Additionally, to ensure the quality of the WF GWAS estimates, we restricted our SNPs to those with MAF ≥ 0.05 in every WF cohort and with imputation quality

(INFO) above 95% in every WF cohort and above 99% in the discovery GWAS. Applying these filters left 2,005,496 SNPs in the intersection of the discovery and WF GWAS.

Then, we applied PLINK's[37] clumping algorithm to the GWAS results with the 1000 Genomes phase 3 EUR reference panel[t] to obtain a subset containing approximately independent SNPs. For this command, we used a threshold of $P = 1$ for lead SNPs and a LD threshold of $r^2 > 0.1$ in a 1,000,000 kb window[u]. We obtained 48,979 approximately independent SNPs with results available for both the GWAS and the WF GWAS.

##### *4.2.3 Bayesian estimation of the posterior distribution of the SNPs' true effect sizes*

Following Okbay *et al.* (2016)[18], we conducted simulations to benchmark the results of our sign tests. To do so, we first obtained estimates of the distribution of the SNPs' true effect sizes (the $\beta_j$'s).

We define $\beta_{j,\text{std}}$ as the coefficient from the regression where both the phenotype and the genotype have been standardized to have mean zero and unit variance. We further assume that the effect sizes are drawn from a mixture distribution of a Gaussian and a point mass at zero:

$$\beta_{j,\text{std}} \sim \begin{cases} N(0, \tau^2) & \text{with probability } \pi \\ 0 & \text{otherwise,} \end{cases}$$

where $\tau^2$ is the variance of non-null SNPs and $\pi$ is the fraction of non-null SNPs in our data. This distributional assumption implies that the variance of effect sizes is inversely proportional to the variance of the unstandardized genotypes[v]. By the Central Limit Theorem, we note that the estimation error of the GWAS estimate of $\beta_{j,\text{std}}$ is approximately normally distributed. We use $\sigma_j^2$ to denote the variance of this error and note that our assumptions imply that $\sigma_j^2 \approx 1/n$. This means the distribution of $\hat{\beta}_{j,\text{std}}$ is:

$$\hat{\beta}_{j,\text{std}} \sim \begin{cases} N\left(0, \sigma_j^2 + \tau^2\right) & \text{with probability } \pi \\ N(0, \sigma_j^2) & \text{otherwise.} \end{cases}$$

Because we have a closed-form distribution, we can use the discovery GWAS summary statistics to estimate its parameters: the probability of a SNP being null $\pi$ and the variance of the non-null effect sizes $\tau^2$ (as mentioned above, $\sigma_j^2 \approx 1/n$).

As shown in Okbay *et al.* (2016)[18], given these parameters, the posterior probability that SNP $j$ with estimated effect size $\hat{\beta}_{j,\text{std}}$ is non-null is:

[t] This smaller reference suffices for this analysis because we only used SNPs with MAF ≥ 0.05 in every WF cohort (and thus we do not use our main reference panel for this analysis).

[u] Note that the threshold of $P = 1$ means that this algorithm does not filter SNPs by $P$ value but rather continues to include SNPs until there is no SNP remaining that is sufficiently uncorrelated with the selected SNPs. We also highlight that this procedure does not choose an arbitrary set of SNPs but instead prioritizes those whose estimates are more significant.

[v] To see this, note that $\beta_{j,\text{std}} = \beta_j \cdot \text{Var}(SNP_j)$ (where $SNP_j$ is the unstandardized genotype at SNP $j$ and can take values 0, 1, or 2). Because we assume that the distribution of $\beta_{j,\text{std}}$ is the same for all SNPs, it follows that $\text{Var}(\beta_j)$ will be larger for SNPs whose unstandardized genotypes have lower variance.

$$p_{\hat{\beta},j} = \frac{\frac{\pi}{\sqrt{\sigma_j^2+\tau^2}}\phi\left(\frac{\hat{\beta}_j}{\sqrt{\sigma_j^2+\tau^2}}\right)}{\frac{1-\pi}{\sigma_j}\phi\left(\frac{\hat{\beta}_j}{\sigma_j}\right)+\frac{\pi}{\sqrt{\sigma_j^2+\tau^2}}\phi\left(\frac{\hat{\beta}_j}{\sqrt{\sigma_j^2+\tau^2}}\right)}.$$

And the posterior distribution of non-null SNPs is:

$$(\beta_j|\hat{\beta}_j, \beta_j \neq 0) \sim N\left(\frac{\tau^2}{\sigma_j^2+\tau^2}\hat{\beta}_j, \frac{\sigma_j^2\tau^2}{\sigma_j^2+\tau^2}\right).$$

Following Okbay *et al.*, we estimated the parameters $\pi$, $\tau^2$, and $p_{\hat{\beta},j}$ using the summary statistics from the discovery GWAS (including the full siblings in the UKB cohort). This generated a posterior distribution of the true effect size of each SNP given the estimated effect sizes.

##### *4.2.4 Simulations*

We used a simulation procedure whereby we drew, for each of the $M$ approximately independent SNPs, "true" effect sizes from the resulting posterior distributions as well as estimation errors, and we repeated the simulation 1,000 times.

We let $E(C)$ denote the expected number of concordant signs under the alternative hypothesis that the SNPs have effect sizes given by the true $\beta_j$'s. The quantity $E(C)$ provides a benchmark for the sign test. If the actual fraction of concordant SNPs matches the expected fraction $E(C)/M$ and is significantly larger then 50%, then we can be reasonably confident that most of the GWAS estimates are not driven by stratification.

We estimated $E(C)$ using the following method. For each of the 1,000 simulations, we generated discovery and WF GWAS estimates for each SNP $j$ by adding Gaussian noise to the "true" effect size $\beta_j$ drawn from the posterior distribution. We obtained the following quantities:

$$\hat{\beta}_{GWAS,j} = \beta_j + \varepsilon_j\,\hat{\sigma}_{GWAS,j}$$

$$\hat{\beta}_{WF,j} = \beta_j + \delta_j\hat{\sigma}_{WF,j},$$

where $\varepsilon_j$ and $\delta_j$ are independent draws from a standard normal distribution and $\hat{\sigma}_{GWAS,j}$ and $\hat{\sigma}_{WF,j}$ are the standard errors of the coefficients for SNP $j$ from the discovery and WF GWAS, respectively.

Let $\hat{C}_k$ be the number of SNPs with matching discovery and WF GWAS signs in simulation $k$. We obtained our estimate of $E(C)$ by averaging $\hat{C}_k$ across the 1,000 simulations:

$$\hat{E}(C) = \frac{1}{1000}\sum_{k=1}^{1000}\hat{C}_k.$$

In addition, we estimated the standard deviation of $C$ by using the formula for the sample standard deviation[w]:

$$\widehat{\mathrm{SD}}(C) = \sqrt{\frac{1}{999}\sum_{k=1}^{1000}\left(\hat{C}_k - \hat{E}(C)\right)^2}.$$

##### *4.2.5 Results of the sign test*

**Supplementary Table 4.2** reports the results of a range of sign tests we conducted. We conducted four sign tests, all of which compared the signs of the estimates from the discovery GWAS of risk tolerance (excluding all full siblings from the UKB) to the signs of the estimates from the meta-analysis of the WF GWAS in the UKB-siblings, STR1, and STR2 cohorts. Before each sign test, we pruned the list of independent clumped SNPs based on the $P$ values obtained in the discovery GWAS. Each sign tests corresponds to one of the following four $P$ value cutoffs: 1 (all SNPs included), 0.05, 0.005, and 0.0005.

As can be seen from **Supplementary Table 4.2**, we strongly reject the null hypothesis of no sign concordance for all of the sign tests. All four sign tests are significant at the $5 \times 10^{-10}$ level.

Note that all of the sign tests we conducted are well powered to reject the null hypothesis of no sign concordance. Based on these sign tests, we can be reasonably confident that at least some of the GWAS estimates are not driven by population stratification.

**Supplementary Table 4.2** also reports the expected number of concordant signs for each of the four sign tests that were conducted. For each test, the observed number of concordant signs is similar to, but nonetheless significantly smaller than, the expected number of concordant signs predicted by the simulation. This discrepancy might be the result of our assumptions about the SNPs' true effect sizes (the $\beta_j$'s). It is also possible that our estimate of the posterior distribution of $\beta_j$ is overly optimistic.

Another possibility is the following. Our simulation procedure assumes that the true effect sizes are the same for both the discovery and WF GWAS, and we use the discovery GWAS summary statistics to estimate the posterior distribution of $\beta_j$. However, it is possible that the true WF GWAS effect sizes are smaller than the true GWAS effect sizes. This could arise, for example, if genetic effects were mediated by the family environment (for instance, if parents successfully attempted to make their children more similar to each other in terms of their risk tolerance by partly counteracting their genetic propensity for risk tolerance). If this were the case, our simulation procedure would overestimate the expected number of concordant signs.

---

[w] This procedure should yield unbiased estimates of the expected value and the standard deviation of $C$, conditional on the estimated posterior distribution of the true effect sizes. Because we used an independent set of SNPs, both the expected value and the standard deviation of $C$ are additive functions of the SNP-level probabilities of sign concordance, and this holds even though the simulation does not take the LD between the SNPs into account.

#### 4.3 Discussion

We have presented the results of two tests of population stratification that rely on different sets of assumptions. The LD Score intercept test relies on stronger assumptions and allows us to quantify how much population stratification is present in our estimates. Our results from this test —for our discovery and replication GWAS of general risk tolerance and for our supplementary GWAS—imply that only a small part of the observed inflation in the mean $\chi^2$ statistics is likely to be accounted for by confounding bias rather than polygenic signal.

Our results from the second test, the sign test, allow us to strongly reject the null hypothesis that the coefficients from our discovery GWAS of general risk tolerance are driven by stratification. All four sign tests are significant at the $5 \times 10^{-10}$ level.

In sum, both the LD Score intercept tests and the sign tests allow us to conclude that our results are unlikely to be driven by population stratification. We can be reasonably confident that the bulk of the variation in the GWAS estimates is attributable to true polygenic signal.

### 5 Out-of-sample replication of the lead SNPs from our discovery GWAS of general risk tolerance

To assess the credibility of the results of our discovery GWAS of self-reported general risk tolerance, we attempted to replicate the associations of the lead SNPs from that GWAS in an independent replication GWAS of self-reported general risk tolerance.

As described in **Supplementary Information section 2.1,** the replication GWAS included 10 cohorts, with a combined sample size of 35,445. To assess the replicability of the lead SNPs from our discovery GWAS, we followed the procedure outlined in Supplementary Information section 1.8 of Okbay *et al.* (2016)[16] and conducted binomial tests to assess whether the associations of the lead SNPs from our discovery GWAS replicate in an independent replication GWAS. We conducted two binomial tests. First, we conducted a binomial sign test to assess whether the directions (i.e., the signs) of the effects of the lead SNPs are concordant across the discovery and the replication GWAS. Second, we conducted a binomial test to assess whether a significantly large fraction of the lead SNPs are significant at the 5% level on one-sided tests (i.e., with concordant signs and significant at the 10% level on two-sided tests) in the replication GWAS.

We benchmarked these results against a plausible alternative hypothesis, where we predicted the number of concordant signs and significant SNPs in the replication GWAS given a posterior estimate of the true distribution of effect sizes. This allowed us to determine whether the results of the two aforementioned binomial tests match what we would expect if the lead SNPs were all true positives.

###### *5.1.1 Constructing the set of lead and proxy-lead SNPs*

We used the 124 independent lead SNPs from our discovery GWAS of general risk tolerance to construct a set of lead and proxy-lead SNPs that are available in the replication GWAS summary statistics. We first identified the subset of the 124 lead SNPs that were directly available in the replication GWAS summary statistics and had a sample size of at least one-half the maximum sample size in that GWAS. We identified 122 such SNPs. Next, for each of the remaining two SNPs, we determined whether there exists a suitable "proxy-lead SNP" that satisfies three conditions: (1) the SNP is in high LD ($r^2 > 0.8$) with the original SNP (we used PLINK[37] and our main reference panel to compute LD); (2) the SNP is available in the summary statistics of both the discovery GWAS and replication GWAS; and (3) the SNP has a sample size of at least one-half the maximum sample size in the replication GWAS. If more than one proxy-lead SNP satisfied these three condition for one original SNP, we selected the one in highest LD with the original SNP as the proxy-lead SNP for the analyses. We identified one such proxy-lead SNP. We combined that proxy-lead SNP with those that were directly available in the replication GWAS to create the set of 123 lead and proxy-lead SNPs.

###### *5.1.2 Binomial replication tests*

We conducted two binomial tests of the null hypothesis that none of the lead SNPs are associated with risk tolerance. Rejecting this null provides evidence that our lead SNPs contain at least some truly associated SNPs. Under the null, we would expect that 50% of the lead SNPs have concordant signs and 5% are significant at the 5% level on the one-sided tests (i.e., significant at the 10% level, with concordant signs) in the replication GWAS. For both binomial tests, we used one-sided tests of the null hypothesis because we are specifically interested in testing for a larger share of concordant or significant SNPs relative to the null share.

Because the lead SNPs are located in independent loci, the number of lead SNPs that have concordant signs or that are significant in the replication GWAS under the null can be modeled as a series of coin flips, where the probability of a "success" is 0.5 for sign concordance and 0.05 for the one-sided tests at the 5% level of significance. It follows that under the null hypothesis, $k$ (which we define as the total number of concordant or significant SNPs) is distributed $k \sim \text{Binomial}(M, \pi)$, where $M$ is the total number of lead SNPs and $\pi$ is the probability of encountering a concordant or significant SNP (i.e., 0.5 for sign concordance or 0.05 for the one-sided tests at the 5% level of significance). Knowing the distribution of $k$ allows us to perform tests of the null hypothesis. The results of these tests are presented below.

###### *5.1.3 Expected replication record*

In addition to the test of the null hypothesis described above, we also benchmarked our replication record against an estimate of a plausible replication record.

As for the sign test in **Supplementary Information section 4.2**, we followed the procedure outlined in Okbay *et al.* (2016)[16] and conducted a Bayesian analysis to obtain estimates of the posterior distribution of the SNPs' true effect sizes (the $\beta_j$'s), given their GWAS estimates. That procedure is described in more detail in **Supplementary Information section 4.2** and in Okbay *et al.* (2016)[16].

To compute the expected sign concordance and its variance under the posterior distribution of the SNPs' true effect sizes, we proceeded as in **Supplementary Information section 4.2**, except that we used the set of lead and proxy-lead SNPs (instead of the ~50,000 approximately independent SNPs) and replaced the within-family GWAS by the replication GWAS.

To compute the expected replication record and its variance at the 5% level under the posterior distribution of the SNPs' true effect sizes, we proceeded as in **Supplementary Information section 4.2**, except that we replaced $\hat{C}_k$ with $\hat{R}_k$, where $\hat{R}_k$ is defined as the number of SNPs in simulation $k$ where the simulated replication effect is a significant replication of the simulated discovery effect.

###### *5.1.4 Replication results*

Out of the 123 lead or proxy-lead SNPs, 94 have concordant signs in the replication GWAS. Under the null hypothesis that the lead SNPs from our discovery GWAS are all null, we would expect 50% of the SNPs (i.e., 61.5 SNPs) to have concordant signs in the replication GWAS. Based on the results from our binomial test of sign concordance, we strongly reject the null hypothesis of 50% concordance ($P = 1.7\times10^{-9}$). Our expected replication record is that 95.9 (*SD* = 4.6) of the 123 SNPs would have matching signs, which matches our actual replication record very closely.

Out of the 123 lead or proxy-lead SNPs, 23 are significant at the 5% level on one-sided tests (i.e., significant at the 10%, with concordant signs) in the replication GWAS. Under the null hypothesis that the lead SNPs from our discovery GWAS are all null, we would expect 5% of the SNPs (i.e. 6.15 SNPs) to reach significance at the 5% level on the one-sided tests in the replication GWAS. We strongly reject this null hypothesis ($P = 4.5\times10^{-8}$). Consistent with this, our expected replication record is that 24.7 (*SD* = 4.4) SNPs would be significant at the 5% level, which again matches our actual replication record very closely.

The replication record of our lead and proxy-lead SNPs is shown in **Extended Data Fig. 5.1**, and the summary statistics for the 123 lead or proxy-lead SNPs are reported in **Supplementary Table 3.1**.

In sum, our actual replication record matches our expected replication record for both the sign and significance binomial tests. Moreover, both binomial tests strongly reject the null hypothesis that none of our lead SNPs are true associations. We also note that under our Bayesian model of true effect sizes, the posterior probability that a SNP is causal is above 99% for all of our 124 lead SNPs. Coupled with our empirical replication results, this suggests that most if not all of the lead SNPs we identified are true positives.

### 6 Estimation of genome-wide SNP heritability

Risk tolerance (both self-reported and experimentally elicited) has been found to be moderately heritable in twin studies, with heritability estimates ranging from 20% to 60%[2,7,8]. In this section, we employ three different methods to obtain estimates of the SNP heritability of our primary and supplementary GWAS phenotypes. A phenotype's SNP heritability is the fraction of the phenotype's variance that is accounted for by the additive genetic effects of a set of SNPs.

#### 6.1 Methods to estimate genome-wide SNP heritability

We used three methods, GCTA[75], LD Score regression[42], and Heritability Estimator from Summary Statistics (HESS)[76], to estimate the genome-wide SNP heritability ($h_G^2$). The GCTA method estimates the heritability of a phenotype directly from the individuals' genotypic data, while the LD Score and HESS methods use GWAS summary statistics as inputs.

For comparability across phenotypes and methods, for the LD Score and HESS methods, we used summary statistics from the UKB GWAS only for all phenotypes except adventurousness. For the adventurousness phenotype, we only report estimates that were obtained using the LD Score regression and HESS methods using the 23andMe summary statistics, because this phenotype is not available in the UKB and we did not have access to the individual-level genotypic data from the 23andMe cohort (and so could not obtain estimates with the GCTA method).

We emphasize that, for general risk tolerance and ever smoker, our main GWAS also analyzed data from cohorts other than the UKB, so the heritability estimates we report in this section are different from those that would have been obtained using the summary statistics from our main GWAS. (**Table 1** reports HESS h2 estimates obtained by using summary statistics from our main GWAS of the seven phenotypes.)

##### *6.1.1 The GCTA method*

The GCTA method is based on restricted maximum-likelihood estimation and uses the genetic relationship matrix (GRM) to estimate the SNP heritability. Under the assumptions discussed in Yang *et al.* (2011)[75], the method leads to unbiased estimates of the genome-wide SNP heritability. However, it is computationally intensive, and it is thus necessary to limit the number of SNPs and individuals included in the analysis in order to be computationally feasible. Therefore, we restricted the GCTA analysis to a random subset of 30,000 individuals out of the full sample from the discovery GWAS. We thereafter dropped one individual in each pair of individuals with a cryptic relatedness exceeding 0.025, to obtain a set of unrelated individuals. For comparability we used the same initial subset of 30,000 individuals for the GCTA estimation for all phenotypes, though the sample size varies slightly across phenotypes because of missing phenotypic observations. The final sample sizes for each phenotype are presented in **Supplementary Table 6.1**. In total 646,855 directly genotyped SNPs with MAF > 0.01 were included in the GCTA heritability estimation.

##### 6.1.2 *The LD Score regression method*

Under the assumptions discussed in Bulik-Sullivan *et al.* (2015)[42], a SNP's GWAS $\chi^2$ statistic is linearly related to its LD score, defined as the sum of the squared correlation coefficients between any single SNP and all the other SNPs. The slope of the LD Score regression (of the SNPs GWAS $\chi^2$ statistics on their LD scores and an intercept) can be rescaled to obtain an estimate of the heritability explained by the SNPs included in the LD Score analysis by dividing the slope by the sample size divided by the number of SNPs, i.e., by *n*/M. We used the "eur_w_ld_chr/" files of LD scores computed by Finucane *et al.*[74] and made available on https://github.com/bulik/ldsc/wiki/Genetic-Correlation, accessed on March 14, 2016. These LD scores were computed with genotypes from the European-ancestry samples in the 1000 Genomes Project. Only HapMap3 SNPs with MAF > 0.01 were included in the LD Score regression; for every phenotype, ~1.3 million SNPs were used for the LD Score heritability estimation. Since Genomic Control (GC) will tend to bias the intercept of the LD Score regression downward, we did not apply GC to the summary statistics prior to estimating the LD Score regressions.

##### 6.1.3 *The HESS method*

The HESS estimator can be described in brief as an analytical variance decomposition method that, unlike the GCTA and LD Score regression methods, assumes that the SNP effect sizes are fixed effects rather than random effects. The method assumes that the SNPs are randomly distributed in the population and requires a pre-specified SNP covariance matrix as input. The SNP covariance matrix can be estimated in the sample of interest if individual genotypic data is available, or with an external reference panel such as the 1000 Genomes[77]. As Shi *et al.*[76] show using simulations, heritability estimates from LD Score regression are sensitive to the true proportion of causal SNPs, and the HESS estimator yields more accurate heritability estimates than LD Score regression under a wider range of proportions of truly causal SNPs. We used the reference panel distributed with the HESS software for the calculation of the covariance matrix. That panel is the European subsample of the 1000 Genomes phase 3 version 5 reference panel, restricted to common variants (MAF > 0.05), which is the same as the reference panel used for the construction of the LD Scores[x]. For every phenotype, a total of ~4.9 million SNPs were used in the HESS heritability estimation. As with the LD Score regressions, we did not apply GC prior to estimating heritability with HESS.

#### 6.2 Results of genome-wide SNP heritability estimation

The results of the genome-wide SNP heritability estimations are reported in **Supplementary Table 6.1** and displayed in **Extended Data Fig. 6.1**. The estimated heritabilities of our primary and supplementary GWAS phenotypes range from 0.055 to 0.173.

For self-reported general risk tolerance, we obtained a GCTA heritability estimate of $\hat{h}^2_{GCTA}$ = 0.085 (*SE* = 0.018), a LD Score heritability estimate of $\hat{h}^2_{LD\ Score}$ = 0.055 (*SE* = 0.002), and a HESS heritability estimate of $\hat{h}^2_{HESS}$ = 0.063 (*SE* = 0.003). Of all estimated phenotype, the

[x] While the same reference panel was used for the construction of the LD scores, as indicated above HapMap3 SNPs with MAF > 0.01 were included in the LD score regression.

highest estimated heritability was for the first PC of the risky behaviors: $\hat{h}^2_{GCTA}$ = 0.173 ($SE$ = 0.025), $\hat{h}^2_{LD\ Score}$ = 0.114 ($SE$ = 0.004), and $\hat{h}^2_{HESS}$ = 0.156 ($SE$ = 0.004).

The methods yield broadly consistent results. For all phenotypes, the heritability estimates are similar across the three methods, although the GCTA heritability estimates are generally the highest and the LD Score regression estimates are generally the lowest.

### 7 Genetic correlations

To assess whether the genetic variants that influence general risk tolerance tend to be the same and to have similar effect sizes as those that influence plausibly related phenotypes, we use bivariate LD Score regression[24] to estimate pairwise genetic correlations for autosomal SNPs between self-reported general risk tolerance and a number of pre-selected phenotypes. We also verify that the genetic correlation for general risk tolerance between males and females is statistically indistinguishable from one (thereby justifying our approach of pooling the data across the sexes in the rest of our analyses) and that the genetic correlation between the discovery and replication GWAS of general risk tolerance is also statistically indistinguishable from one.

#### 7.1 Methodology

Under some assumptions, bivariate LD Score regression[24] produces unbiased estimates of genetic correlation, even in the presence of sample overlap. Under these assumptions the method merely requires GWAS summary statistics and an "LD score" (the amount of genetic variation tagged by a SNP) reference panel.

Bivariate LD Score regression utilizes the following moment condition:

(6)
$$E\left[z_{1j}z_{2j}\right] = Intercept + \frac{\sqrt{N_1 N_2}}{M} Cov_g \ell_j,$$

where $z_{kj}$ is the $z$-statistic of SNP $j$ from the GWAS of trait $k$ ($k$ = 1, 2), *Intercept* is the regression intercept, $N_k$ is the sample size of the GWAS of trait $k$, $M$ is the number of SNPs included in the GWAS, $Cov_g$ is the genetic covariance between traits *1* and *2*, and $\ell_j$ is the LD score of SNP $j$. The slope parameter from a regression of $\hat{z}_{1j}\hat{z}_{2j}$ on $\sqrt{N_1 N_2}\ell_j$ can therefore be used to estimate the genetic covariance between the two traits. From separate, univariate LD Score regressions of traits 1 and 2, we can also back out estimates of the respective heritabilities of the two traits, $h_{g1}^2$ and $h_{g2}^2$, and obtain an estimate of the genetic correlation as follows:

(7)
$$\hat{r}_g = \frac{\widehat{Cov}_g}{\sqrt{\hat{h}_{g1}^2 \hat{h}_{g2}^2}}.$$

We used the scores computed by Finucane *et al.*[74], which use genotypic data from the European-ancestry samples in the 1000 Genomes Project and only HapMap3 SNPs (eur_w_ld_chr, see https://github.com/bulik/ldsc/wiki/Genetic-Correlation, accessed on March 14, 2016). As is common in the literature, we restrict our analyses to SNPs with MAF > 0.01; this guarantees all analyses are performed using a set of SNPs that are imputed with reasonable accuracy across all contributing cohorts. The standard errors are estimated by the LDSC software using a block jackknife over the SNPs.

#### 7.2 Pre-selected phenotypes

For our bivariate LD Score analyses, we considered a wide range of phenotypes. First, we considered our supplementary GWAS phenotypes. These include adventurousness, our four main risky behaviors (automobile speeding propensity, drinks per week, ever smoker, and number of sexual partners) and their first PC (see **Supplementary Table 7.1A1**). We also analyzed additional risky behavior phenotypes for which we ran additional GWAS with only the *first* release of UKB data (for details regarding the methodology and phenotype definitions, see the Appendix at the end of this section); these include age first had sexual intercourse, teenage conception (females only), and use of sun protection (see **Supplementary Table 7.1A2**). Further, we analyzed additional risky behaviors for which we were able to obtain summary statistics from previously published well-powered GWAS. We include the risky behaviors age tobacco smoking onset (among ever-smokers)[29] cigarettes per day (among ever-smokers)[29], former tobacco smoker (among ever-smokers)[29] lifetime cannabis use[78], and self-employed[79] (**Supplementary Table 7.1A3**).

Second, we include the *cognition phenotypes* cognitive performance[80], educational attainment[16], and intracranial volume[81] (**Supplementary Table 7.1B**). Third, we selected the *anthropometric phenotypes* BMI[82] and height[83] (**Supplementary Table 7.1C**). Fourth, we analyzed the *neuropsychiatric phenotypes* ADHD[53], Alzheimer's disease[84], anxiety disorders[85,y], autism spectrum disorder[86], bipolar disorder[87], depressive symptoms[18], and schizophrenia[69] (**Supplementary Table 7.1D**).

Fifth, we analyzed the five *personality phenotypes* that make up the five-factor personality model (agreeableness, conscientiousness, extraversion, neuroticism, and openness to experience) using GWAS summary statistics provided by 23andMe and previously analyzed by Lo *et al.* (2016)[88] (**Supplementary Table 7.1E**). Also known as the Big Five, these traits constitute the most widely used taxonomy of personality traits in psychology. The Big Five have roots in Allport & Odbert's lexical hypothesis[89], which states that individual differences are encoded in language[90]. This is in contrast to economic preferences such as risk aversion, which are measures of individual heterogeneity that arise in a utility maximization framework.

Recent work in economics has highlighted the importance of personality for economic outcomes, particularly in education, crime, health, and the labor market[91]. However, little is known about the relationship between the Big Five and economic preferences. In the most comprehensive study of its kind to date, Becker *et al.* (2012)[3] find highly significant positive correlations for self-reported risk aversion with openness (0.28) and extraversion (0.26), and negative correlations with conscientiousness (-0.04), agreeableness (-0.14), and neuroticism (-0.09)[3].

We also ran GWAS using the first release of UKB data for the *socioeconomic phenotypes* household income and Townsend deprivation index (see **Supplementary Table 7.1E**; for details regarding the methodology and phenotype definitions, see again the Appendix at the end of this section). Finally, we evaluated the genetic correlation between risk and longevity[92].

---

[y] Otowa et al.[85] used two phenotype measures in their GWAS of anxiety disorders: one is based on a categorical case-control designation for having any anxiety disorder diagnosis, and the other is based on continuous factor scores derived by combining information across clinical phenotypes. Our analyses are based on the case-control measure.

#### 7.3 Results: Genetic correlation between general risk tolerance and pre-selected phenotypes

To estimate the genetic correlations between general risk tolerance and each of the above phenotypes, we used the summary statistics from the meta-analysis of the discovery and replication GWAS. The estimates of the genetic correlations are shown in **Supplementary Table 7.1** and in **Fig. 2**.

##### *7.3.1 Adventurousness and risky behaviors*

First, we examine the genetic correlation between adventurousness and general risk tolerance. Then, we return to our other supplementary GWAS phenotypes and their genetic correlations with general risk tolerance. These results are reported in **Supplementary Table 7.1A1**. As expected, the estimated genetic correlation with adventurousness is high and statistically significant ($\hat{r}_g$= 0.834, *SE* = 0.011). The correlation with automobile speeding propensity is moderately high and highly statistically significant ($\hat{r}_g$= 0.448, *SE* = 0.021), consistent with individuals with more risk-tolerance increasing alleles being more likely to exceed the automobile speed limit. For drinks per week the genetic correlation is positive and highly statistically significant ($\hat{r}_g$= 0.254, *SE* = 0.021), and the genetic correlation is comparable for ever smoker ($\hat{r}_g$= 0.246, *SE* = 0.022), implying that higher risk tolerance is genetically associated with more risky health behaviors. The genetic correlation estimates are highly significant for number of sexual partners ($\hat{r}_g$= 0.520, *SE* = 0.019). Finally, the genetic correlation for the first PC of these four risky behaviors, which can be interpreted as a general factor of risky behavior, is interestingly both moderately high in absolute value and highly statistically significant ($\hat{r}_g$= 0.500, *SE* = 0.018). Thus, all of our supplementary GWAS phentoypes are significantly genetically correlated in the expected direction with the primary general-risk-tolerance phenotype.

Next, let us turn to the three additional UKB risky behaviors (for which we ran GWAS using only the first release of UKB data) and their genetic correlations with general risk tolerance. The results are reported in **Supplementary Table 7.1A2**. The genetic correlation estimates are highly significant for age first had sexual intercourse ($\hat{r}_g$= -0.332, *SE* = 0.032) and teenage conception ($\hat{r}_g$= 0.246, *SE* = 0.049). Together with our results of number of sexual partners, this implies that that higher risk tolerance is genetically associated with more risky sexual behavior; this is also consistent with a finding from a recent GWAS of age at first sexual intercourse[50]. Use of sun protection has an insignificant genetic correlation with general risk tolerance, and we suspect that this phenotype may be more highly correlated with skin pigmentation and predisposition to skin cancer than to general risk tolerance.

All seven of our statistically significant genetic correlation estimates for the aforementioned risky behaviors have signs that are in the direction that would be expected based on the corresponding phenotypic correlations, and the absolute values of the estimates are higher than those of these phenotypic correlations.

Lastly, **Supplementary Table 7.1A3** reports the estimates for the risky behaviors for which we were able to obtain summary statistics from previously published, well-powered GWAS. One of the three cigarette-related phenotypes is moderately and significantly genetically correlated with general risk tolerance: former tobacco smoker (among ever-smokers) ($\hat{r}_g$= -0.131, $SE$ = 0.055). However, age of tobacco smoking onset (among ever-smokers) and cigarettes per day (among ever-smokers) are not significantly genetically correlated with general risk tolerance. These results suggest that, while higher risk tolerance may be genetically associated with some risk-related smoking behaviors such as smoking initiation (as suggested by the significant genetic correlation with ever smoker) and cessation, risk tolerance may not necessarily be genetically correlated with smoking addiction (as captured by cigarettes per day).

For lifetime cannabis use, the genetic correlation is positive and highly statistically significant ($\hat{r}_g$= 0.313, $SE$ = 0.057), implying that risk tolerance is genetically associated with risk-seeking cannabis use behavior. The genetic correlation estimate for self-employed is significant and large in magnitude ($\hat{r}_g$= 0.672, $SE$ = 0.259), implying that higher risk tolerance is genetically associated with a common proxy for entrepreneurship. The point estimate for this phenotype is the highest among any of the phenotypes tested. However, in interpreting this correlation it is important to note that the LD Score regression heritability of the self-employment phenotype is low and not significantly different from zero: 0.0126 ($SE$ = 0.0106). Indeed, the standard errors on the estimate are quite large ($SE$ = 0.259). This might be due to the fairly small sample size of the self-employment GWAS ($n$ = 50,627).

All 11 of our statistically significant genetic correlation estimates for the risky behaviors have signs that imply higher risk tolerance is associated with riskier behaviors.

###### *7.3.2 Cognition phenotypes*

Two of the three cognition phenotypes are significantly, though weakly, genetically correlated with general risk tolerance: educational attainment ($\hat{r}_g$= 0.099, $SE$ = 0.022) and intracranial volume ($\hat{r}_g$= 0.144, $SE$ = 0.059). These results are consistent with the well-established positive correlation between risk preferences and cognition and educational attainment in the literature[2,4,93].

###### *7.3.3 Anthropometric phenotypes*

Height is not significantly genetically correlated with general risk tolerance. BMI, on the other hand, is significantly, albeit weakly, positively correlated ($\hat{r}_g$= 0.053, $SE$ = 0.021).

###### *7.3.4 Neuropsychiatric phenotypes*

Among the neuropsychiatric traits, we find moderate (and highly significant) genetic correlations with ADHD ($\hat{r}_g$= 0.247, $SE$ = 0.033), anxiety disorders, ($\hat{r}_g$= 0.214, $SE$ = 0.089), autism spectrum disorder ($\hat{r}_g$= -0.105, $SE$ = 0.050), bipolar disorder ($\hat{r}_g$= 0.214, $SE$ = 0.035) and schizophrenia ($\hat{r}_g$= 0.173, $SE$ = 0.021). The positive and significant genetic correlation with ADHD is consistent with the significant effect of our polygenic score of general risk tolerance on ADHD in an independent sample (**Supplementary Information section 10**). We do not, however, find significant genetic correlations with either Alzheimer's disease or depressive symptoms.

##### *7.3.5 Personality phenotypes*

Agreeableness and conscientiousness are not significantly correlated with general risk tolerance. However, the genetic correlations between general risk tolerance and extraversion ($\hat{r}_g$= 0.505, *SE* = 0.027), neuroticism ($\hat{r}_g$= -0.420, *SE* = 0.038), and openness ($\hat{r}_g$= 0.332, *SE* = 0.031), are moderately high in absolute value and highly significant. The direction of the genetic correlation between general risk tolerance and all five personality traits is in line with the literature on the phenotypic correlations of these traits outlined above[3]. Our results for extraversion, neuroticism, and openness are also in line with the signs of the significant coefficients on the polygenic score for general risk tolerance in regressions of each of these personality phenotypes on the score, as highlighted in **Supplementary Information section 10** (although our predictive power for these traits is quite low).

##### *7.3.6 Socioeconomic phenotypes and longevity*

The genetic correlation estimates for both household income ($\hat{r}_g$= 0.215, *SE* = 0.033) and for Townsend score ($\hat{r}_g$= 0.185, *SE* = 0.047) are positive and significant, implying that higher risk tolerance is genetically associated with higher earnings and social deprivation. The genetic correlation with Townsend score is higher than even the phenotypic correlation. We do not find a significant genetic correlation with longevity.

##### *7.3.7 Summary of Findings*

In sum, general risk tolerance tends to be genetically correlated with adventurousness and with the risky behaviors involving automobile speeding propensity, substance use, sexual activity, and self-employment. Importantly, our estimates have signs that are consistent with higher self-reported risk tolerance being associated with riskier behavior. General risk tolerance is also genetically correlated with the neuropsychiatric phenotypes ADHD, anxiety disorders, autism spectrum disorder, bipolar disorder, and schizophrenia, but not with Alzheimer's disease or depressive symptoms, and with the personality phenotypes extraversion, neuroticism, and openness, but not with agreeableness or conscientiousness.

Our results also point toward distinctions in the genetic correlation between general risk tolerance and externalizing and internalizing behaviors and disorders. Externalizing behaviors and disorders are those in which individuals tend to express maladaptive thoughts and feelings toward others or their environment, while internalizing behaviors and disorders are those in which individuals tend to express thoughts and feelings inward. Overall, we find significant genetic correlations with behaviors and disorders typically classified as externalizing[94], such as substance abuse (smoking cigarettes, cannabis use, and drinking) and ADHD as well as thought disorders related to externalizing disorders such as bipolar disorder and schizophrenia. Conversely, we find little evidence of significant genetic correlation with internalizing behaviors or disorders, such as depression. We also find evidence confirming the well-documented phenotypic correlation between risk taking and cognitive performance and educational attainment (e.g., in the domain of financial risk taking[2,4,93]).

#### 7.4 Other results

##### *7.4.1 Genetic correlation between males and females*

Phenotypically, more than 33% of males in the UKB are categorized as risk tolerant (based on their answers to the general-risk-tolerance question), whereas only approximately 18% of females are categorized as risk tolerant. To test the extent to which the genetics of general risk tolerance differs between males and females, we conducted GWAS of general risk tolerance in the first release of UKB data, separately for males and females. We followed the same methodology and QC protocol for these two sex-specific GWAS as for our discovery GWAS of general risk tolerance and our GWAS of the four main risky behaviors and their first PC in the full release of UKB data, except that we only used the first release of UKB data and we only used SNPs with MAF > 0.005 instead of MAF > 0.001; the latter does not affect the analyses reported in this section, as we restrict these analyses to SNPs with MAF > 0.01. **Supplementary Table 7.2** reports the SNP-based heritabilities and other relevant summary statistics from the LD Score regressions. General risk tolerance is similarly heritable for males ($h^2$ = 0.066, *SE* = 0.009) and females ($h^2$ = 0.053, *SE* = 0.101), and the difference is not statistically significant. We then used bivariate LD Score regression to calculate the genetic correlation between these two samples, and obtained an estimate of $\hat{r}_g$ = 0.9489 (*SE* = 0.1351). We cannot reject perfect genetic correlation between the sexes, thereby justifying our approach of pooling males and females in our other analyses. The phenotypic differences between males and females can therefore either be the result of (1) factors (which could be biological or environmental) that act across gender independently of genotypes, or (2) the sex chromosomes (which we do not analyze here).

##### *7.4.2 Genetic correlation between the discovery and replication GWAS*

We estimated the genetic correlation between our discovery and our replication GWAS of general risk tolerance. We find a high genetic correlation that cannot be statistically distinguished from unity ($\hat{r}_g$= 0.8344, *SE* = 0.1289), suggesting the genetic underpinnings of general risk tolerance do not vary much between our discovery and replication samples.

As discussed in **Supplementary Information sections 1** and **3.3**, we also estimated the genetic correlation between: (1) the UK Biobank general-risk-tolerance GWAS and the 23andMe general-risk-tolerance GWAS; (2) the UK Biobank general-risk-tolerance GWAS and the replication GWAS; (3) the 23andMe general-risk-tolerance GWAS and the replication GWAS; and (4) the discovery and replication GWAS of general risk tolerance. For (1) we find a moderately high, positive genetic correlation ($\hat{r}_g$= 0.767, *SE* = 0.021) that *is* distinguishable from unity. For (2) we find a moderately high, positive genetic correlation ($\hat{r}_g$= 0.828, *SE* = 0.135); for (3) we find ($\hat{r}_g$= 0.759, *SE* = 0.126). Finally, for the final correlation (4), we find ($\hat{r}_g$= 0.834, *SE* = 0.129). The last three correlations are all indistinguishable from unity.

#### 7.5 Comparison of the genetic and phenotypic correlations

In **Supplementary Table 1.3** we show phenotypic correlations for our primary GWAS phenotype, general risk tolerance, and for our five supplementary phenotypes whose GWAS included data from the UKB: automobile speeding propensity, drinks per week, ever smoker, number of sexual partners, and the first PC of these four risky behaviors (the GWAS of our other supplementary phenotype, adventurousness, used 23andMe data only). Panel **A** shows values calculated naively assuming Pearson correlation coefficients for all of our variables using the “pwcorr” command in Stata. In the first column, we show test-retest correlations between the first and the second measurements if each phenotype in the UKB, to give a sense of the test-retest reliability of each phenotype. In the subsequent correlation matrix, coefficients below the diagonal are uncorrected Pearson coefficients. Above the diagonal, we adjust the correlation estimate for each pair of variables for measurement error by dividing it by the square root of the product of the two test-retest correlations for the two variables[z].

In Panel **B**, we recalculate Panel **A**, but this time we use the polychoric package in Stata to allow the correct estimation of tetrachoric (between two ordinal variables) or polyserial correlations (between an ordinal and a continuous variable). Again, uncorrected correlation coefficients are below the diagonal, while correlations corrected for measurement error are above the diagonal.

In summary, both correcting for measurement error (Panel **A** and **B**, above the diagonals) and specifying the correct type of correlation coefficient (Panel **B**) raises the value of the correlation estimates, so that our highest estimates are in Panel **B**, above the diagonal. While some correlations are quite high (especially between the first PC phenotype and the other phenotypes), the others remain relatively low, including for the correlations between general risk tolerance and the five supplementary phenotypes.

In **Supplementary Table 7.1**, Panel **A1**, we compare the *genetic* correlations with general risk tolerance to the *phenotypic* correlations with general risk tolerance. Even after specifying the correct type of correlation (Pearson or polyserial) and adjusting for measurement error, most genetic correlations remain considerably higher than the phenotypic correlations.

These results are relevant to the ongoing debate about the extent to which risk tolerance is a “domain-general” versus “domain-specific” trait. Low phenotypic correlations among measures of risky behaviors in various domains have led some researchers to conclude that risk tolerance is highly domain-specific[95,96]. The comparatively large genetic correlations we estimate support the view that a general factor of risk tolerance partly accounts for cross-domain variation in risky behavior[6,97] and imply that this factor is genetically influenced, while the lower phenotypic correlations suggest that environmental factors are more important contributors to domain-specific behavior.

[z] This is a standard method to correct for measurement error attenuation in the correlation estimates (see, e.g., https://en.wikipedia.org/wiki/Correction_for_attenuation).

#### 7.6 Appendix: GWAS of other risky behaviors and of socioeconomic phenotypes using the first release of UKB data

While we used existing published GWAS results for most phenotypes, we conducted our own GWAS using the full release of UKB data for our four main risky behaviors and their first PC. We describe the coding of our four main risky behaviors and their first PC in **Supplementary Information section 1.2** (for phenotypic correlations of these five behaviors see **Supplementary Table 1.3**). We also conducted our own GWAS using only the first release of UKB data for five phenotypes: age first had sexual intercourse[aa] (*n* = 98,956), teenage conception among females (*n* = 40,077), use of sun protection, household income (*n* = 97,059), and Townsend deprivation index[bb] score (*n* = 112,192). Throughout these analyses we dropped participants who answered "Do not know" or "Prefer not to answer" and averaged data across two assessment visits when possible.

The five remaining UKB phenotypes for which we ran GWAS in only the first release of UKB data were coded as follows:

- Age first had sexual intercourse: UKB respondents were asked "What was your age when you first had sexual intercourse? (Sexual intercourse includes vaginal, oral or anal intercourse)?" We dropped anyone reporting "Never had sex" or an age of first sexual encounter at less than 12 (given the high likelihood of associated abuse or misreporting). We then normalized the measure separately for males and females.
- Teenage conception among females: UKB females who bore at least one child were asked "How old were you when you had your FIRST child?" We recoded this variable into a case-control binary variable. Females reporting age of first live birth between 13 and 20 (inclusive) were coded as cases (*n* = 6,285), while females reporting higher ages of first live birth were coded as controls (*n* = 33,792). Childless females were dropped.
- Use of sun protection: UKB respondents were asked "Do you wear sun protection (e.g. sunscreen lotion, hat) when you spend time outdoors in the summer?"; eligible responses ranged from "1. Never/rarely" to "4. Always," and also included "5. Do not go out in sunshine." We dropped all participants who answered "5. Do not go out in sunshine" and normalized the resulting categorical variable separately for males and females.
- Household income: UKB respondents were asked "What is the average total income before tax received by your HOUSEHOLD?"; eligible responses were "1. Less than £18,000," "2. £18,000 to £30,999," "3. £52,000 to £100,000," "5. Greater than £100,000." We normalized the resulting categorical variable.
- Townsend deprivation index score: This score measures local social deprivation based on the preceding national census output areas; a higher score implies more social deprivation. Each participant was assigned a score corresponding to the output area in

[aa] Day *et al.*[50] report results from a GWAS of age of first sexual encounter. We conduct similar analyses ourselves here. We treat the phenotype slightly differently by separately normalizing phenotypes among males and females, and then conducting our GWAS on the combined sample.

[bb] Hill *et al.*[245] report results from GWAS for household income and the Townsend deprivation index in the UKB. We ran our own analyses because we could not find the summary statistics from their GWAS in the public domain.

which their postcode is located. The score is calculated by the UKB immediately prior to each participant joining the dataset. We normalized the scores for our analysis.

We followed the same methodology and QC protocol for these five additional GWAS as for our discovery GWAS of general risk tolerance and our GWAS of the four main risky behaviors and their first PC in the full release of UKB data (see **Supplementary Information section 2**), except that we only used the first release of UKB data and we only used SNPs with MAF > 0.005 instead of MAF > 0.001 (the latter does not affect the analyses reported in this section, as we restrict these analyses to SNPs with MAF > 0.01). **Supplementary Table 7.2B** reports the various statistics outputted by LD Score regressions for each of these five additional GWAS. The third column shows estimates of the SNP-based heritabilities. All five phenotypes have a higher SNP-based heritability than general risk tolerance, except Townsend score. The sexual activity phenotypes exhibit particularly high heritabilities. Age first had sexual intercourse has the highest SNP-based heritability: 0.167 (*SE* = 0.009). The only estimate of the LD Score regression intercept that is significantly different from one is for household income, for which the intercept is 1.035 (*SE* = 0.008). By comparison, the mean $\chi^2$ statistics for the SNPs in the LD Score regressions are larger than 1.10 for three of the five GWAS, including household income for which the mean $\chi^2$ statistic is 1.198 (the exceptions are teenage conception and Townsend score, for which the mean $\chi^2$ statistics are 1.074 and 1.094, respectively). These estimates imply that only a small part of the observed inflation in the mean $\chi^2$ statistics of the GWAS is likely to be accounted for by confounding bias (due to population stratification, cryptic relatedness, or other confounds), rather than by polygenic signals. Additional details for these five additional GWAS are available upon request.

### 8 Proxy-phenotype analyses

#### 8.1 Introduction

We conducted proxy-phenotype analyses[80] to search for additional SNPs that affect phenotypes that are plausibly related to general risk tolerance. These analyses allow us to test whether SNPs that are strongly associated with a "first-stage" phenotype are enriched for association with a related "second-stage" phenotype, and potentially to identify new SNPs associated with the second-stage phenotype. Proxy-phenotype analyses leverage the fact that, if the first- and second-stage phenotypes are genetically correlated, SNPs associated with the first-stage phenotype should have a higher probability of being associated with the second-stage phenotype, compared to what we would expect by chance.

In our study, the first-stage phenotype is always general risk tolerance, and the first-stage analysis was conducted in our discovery GWAS. We consider eight primary second-stage phenotypes: age of smoking onset, cigarettes per day (CPD), smoking cessation (which compares former smokers to current smokers), lifetime cannabis use, self-employment status (an indicator of whether an individual is self-employed or not), ADHD, bipolar disorder, and schizophrenia. We chose these second-stage phenotypes for the following two reasons. First, summary statistics from GWAS of these phenotypes that did not include the UKB are publicly available or could be obtained. Second, these phenotypes are plausibly related to general risk tolerance: the first five phenotypes are risky behaviors and all eight are at least moderately genetically correlated with general risk tolerance (**Supplementary Information section 7.3**). We also used height as a negative control.

#### 8.2 Methodology

##### *8.2.1 Data and Setup*

We used our discovery GWAS of general risk tolerance as our first-stage analysis. The second-stage lookups were all performed using summary statistics from previous GWAS. We obtained the summary statistics from the following sources: The Tobacco and Genetics Consortium (2010)[29] for the three tobacco smoking phenotypes, the International Cannabis Consortium lifetime for cannabis use (Stringer *et al.* 2016)[78], the Gentrepreneur Consortium for self-employment (van der Loos *et al.* 2013)[79], the Psychiatric Genomics Consortium for ADHD, bipolar disorder, and schizophrenia (Demontis *et al.* 2017, Sklar *et al.* 2011, and Ripke *et al.* 2014, respectively)[53,69,87], and the GIANT consortium for height (Wood *et al.* 2014)[83]. Our methodology follows Okbay *et al.* (2016)[18] and involves two main stages.

##### *8.2.2 Stage 1: Constructing the set of lead and proxy-lead SNPs*

The first stage involves constructing the set of lead SNPs from the first-stage analysis. Unlike Okbay *et al.*, we only used the lead SNPs (i.e., the independent SNPs with $P$ values $< 5 \times 10^{-8}$) from our discovery GWAS of general risk tolerance as candidate SNPs for the second stage[cc].

For brevity, we illustrate the steps involved for cigarettes per day (CPD), but analogous procedures apply for each of the other second-stage phenotypes. Our discovery GWAS of general risk tolerance identified 124 lead SNPs. Of these, 52 SNPs were directly available in the CPD summary statistics, whereas 72 were either not available or their GWAS sample sizes were

[cc] Okbay *et al.* used all SNPs with a $P$ value less than $1 \times 10^{-4}$, rather than just the lead SNPs.

too small to meet our inclusion criterion: to ensure the quality of the estimates for the second-stage SNPs, we limit our lookup procedure to second-stage SNPs with GWAS sample sizes of at least one-half of the maximum sample size for that GWAS. For each of these 72 SNPs, we determined whether there exists a suitable "proxy-lead SNP" that satisfies three conditions: (1) the SNP is in high LD ($r^2 > 0.8$) with the risk-tolerance associated SNP; (2) the SNP is available in the summary statistics of both the discovery GWAS of general risk tolerance and the GWAS of CPD; and (3) the SNP has a CPD sample size of at least one-half the maximum sample size in the CPD GWAS. To determine the set of SNPs in high LD with the risk-associated SNP, we employ PLINK[37] with our main reference panel[dd]. A proxy-lead SNP was available for an additional 44 of the 72 SNPs (with $r^2$'s ranging from 0.80 to 1.00, with a mean $r^2 = 0.94$). Whenever more than one proxy was available for a SNP, we chose the proxy with the largest $r^2$. Ties were broken by choosing the closest SNP to the original lead SNP in base pair distance. Our final list of lead and proxy-lead SNPs for CPD therefore contained 52 + 44 = 96 SNPs.

###### *8.2.3 Stage 2, part 1: SNP lookup and search for previously identified significant loci*

We *individually* tested each of the *k* lead and proxy-lead SNPs for experiment-wide significance by examining whether each is significantly associated with the second-stage phenotype, using a Bonferroni-corrected significance level of $0.05/k$ (for CPD, $k = 96$). We refer to the lead and proxy-lead SNP that reached Bonferroni-corrected significance in the second stage as "second-stage hits."

For each second-stage hit, we then identified the set of SNPs in the second-stage hit's locus by identifying the SNPs in a 1,000 kb window around the corresponding first-stage lead SNP and with $r^2 > 0.1$ with that corresponding first-stage lead SNP (we used PLINK[37] and our main reference panel to compute LD). We then checked if any of these SNPs are genome-wide significant in the second-stage summary statistics. This allowed us to determine whether the second-stage hits tag genomic regions that were previously found to be genome-wide significant in the GWAS of the second-stage phenotype.

###### *8.2.4 Stage 2, part 2: Testing the set of lead and proxy-lead SNPs for enrichment and sign concordance*

For each second-stage phenotype, we performed a non-parametric Mann-Whitney test[67] of joint enrichment to test whether the set of lead and proxy-lead SNPs have a *P* value distribution that is significantly different from the *P* value distribution of a randomly-chosen, matched set of "comparison SNPs" in the second-stage summary statistics. Because we expect the second-stage phenotypes to be highly polygenic, with many SNPs having weak but true associations, it would have been inappropriate to test the null hypothesis that the *P* value distribution of the lead and proxy-lead SNPs is uniform. For each lead and proxy-lead SNP, the comparison SNPs are randomly selected from among the set of SNPs that have a minor allele frequency within one percentage point of the lead or proxy-lead SNP[ee]. As in Okbay *et al.* (2016)[18], we generated 1,000 comparison SNPs for each of the *k* lead or proxy-lead SNPs, and compared the *P* value distribution for this group of $k \times 1000$ SNPs with that of the *k* lead or proxy-lead SNPs.

We also conducted a sign test to assess whether the lead and proxy-lead SNPs have effects in the predicted (or concordant) direction in the second stage. For cigarettes per day (CPD), lifetime

[dd] Throughout this section, we employ our main reference panel to compute the LD between SNPs.

[ee] The matched SNPs are drawn with replacement from the set of SNPs in the second-stage summary statistics that excludes the *Y* lead or proxy-lead SNPs.

cannabis use, self-employment status, ADHD, bipolar disorder, and schizophrenia (for which a higher phenotype value corresponds to more risk taking, and which we estimated to be positively genetically correlated with general risk tolerance) we classified a SNP as having an effect "in the predicted direction" if the sign of its effect is concordant with that for general risk tolerance. For age of smoking onset and smoking cessation (for which a higher phenotype value corresponds to less risk taking, and which we estimated to be negatively genetically correlated with general risk tolerance) we classified a SNP as having an effect "in the predicted direction" if the sign of its effect is discordant with that for general risk tolerance. For example, a SNP that increases general risk tolerance and decreases the age of smoking onset has an effect in the predicted direction. For ADHD, bipolar disorder, schizophrenia, and height, we did not predict the direction of the SNPs' effects; we simply conducted a sign test to establish whether SNPs' effects are more likely to be sign concordant with those for general risk tolerance than expected by chance.

#### 8.3 Results from proxy-phenotype enrichment analyses

Q-Q plots for the lead and proxy-lead SNPs in each second-stage phenotype are shown in **Extended Data Fig. 8.1**. These plots show results from the sign test mentioned above and highlight any second-stage hits. Further details on each of the lead and proxy-lead SNPs are reported in **Supplementary Table 8.1**, which reports summary statistics from the first and second stages for the lead SNPs associated with general risk tolerance and their proxies.

##### *8.3.1 Do risk-tolerance-associated SNPs predict smoking behaviors?*

For CPD, 52 lead SNPs are directly available in the second-stage summary statistics, and we identified 44 proxy-lead SNPs. Out of 96 lead or proxy-lead SNPs, 51 (53.1%) have signs in the predicted direction ($P = 0.31$). There are no second-stage hits. Moreover, the Mann-Whitney test of joint enrichment fails to reject the null hypothesis that the $P$ values of the lead and proxy-lead SNPs are drawn from the same $P$ value distribution as a set of randomly-selected SNPs ($P = 0.88$).

For age of smoking onset, 52 lead SNPs are directly available in the second-stage summary statistics, and we identified 45 proxy-lead SNPs. Of a total of 97 lead or proxy-lead SNPs, 54 (55.7%) have signs in the predicted direction ($P = 0.15$). There were again no second-stage hits, and the Mann-Whitney test of joint enrichment fails to reject the null ($P = 0.48$).

For smoking cessation, 53 lead SNPs are directly available in the second-stage data, and we identified 44 proxy-lead SNPs. Of a total of 97 lead or proxy-lead SNPs, 61 (62.9%) have signs in the predicted direction ($P = 0.007$). There were no second-stage hits. The Mann-Whitney test of joint enrichment fails to reject the null ($P = 0.11$).

##### *8.3.2 Do risk-tolerance-associated SNPs predict lifetime cannabis use?*

For lifetime cannabis use, 117 lead SNPs are directly available in the second-stage data, with no additional proxy-lead SNPs. Of these lead SNPs, 76 out of 117 (65.0%) have signs in the predicted direction ($P = 8 \times 10^{-4}$). One SNP, rs993137, reaches Bonferroni-corrected significance ($P = 1.7 \times 10^{-4}$, before Bonferroni correction) and is thus a second-stage hit, and the sign of the effect of this SNP is in the predicted direction. This SNP does not tag any previously identified genome-wide hit for lifetime cannabis use, and is thus a novel association. The Mann-Whitney test of joint enrichment fails to reject the null hypothesis ($P = 0.13$).

Interestingly, rs993137 falls within the *CADM2* gene, which was found to be significantly associated with lifetime cannabis use in a gene-based test by Stringer *et al.* (2016)[78]. Stringer *et al.* also note that *CADM2* has previously been associated with body mass index (BMI), processing speed, and autism disorders; these phenotypes themselves have been previously associated with cannabis use[98–100]. It is noteworthy that *CADM2* contains lead SNPs for all our primary and supplementary GWAS. The NHGRI-EBI GWAS Catalog database[49] reports previous associations with age at menarche, cognitive function, and educational attainment, among other phenotypes, as well as those mentioned by Stringer *et al.* (2016).

###### *8.3.3 Do risk-tolerance-associated SNPs predict self-employment?*

For self-employment, 56 lead SNPs are directly available in the second-stage data, along with 43 proxy-lead SNPs. Of a total of 99 lead or proxy-lead SNPs, 64 (64.6%) have signs in the predicted direction ($P$ = 0.002). There is one second-stage hit, SNP rs7387531 ($P = 1.1 \times 10^{-4}$, before Bonferroni correction), and the sign of its effect is in the predicted direction.[ff,gg] To our knowledge, no robust association has previously been reported between a genetic variant and self-employment; thus, if the association with rs7387531 is robust, this would be the first genetic variant to be found to be significantly associated with self-employment.

Lastly, the Mann-Whitney test of joint enrichment fails to reject the null hypothesis ($P$ = 0.49). The significant SNP rs7387531 is located within a candidate inversion on chromosome 8 (~7.89 to 11.79 Mb), and that candidate inversion contains lead SNPs for our GWAS of general risk tolerance, adventurousness, automobile speeding propensity, ever smoker, and number of sexual partners. The NHGRI-EBI GWAS Catalog database[49] reports SNP associations within the breakpoints of the candidate inversion with many other phenotypes, among them are neuroticism, extraversion, schizophrenia, and chronotype.

###### *8.3.4 Do risk-tolerance-associated SNPs predict ADHD?*

For ADHD, 116 lead SNPs are directly available in the second-stage data, with one additional proxy-lead SNP. Of a total of 117 lead or proxy-lead SNPs, 74 (63.2%) SNPs have concordant signs ($P$ = 0.003). There are four second-stage hits (rs10905461, $P = 5.0 \times 10^{-5}$ before Bonferroni correction; rs3764002, $P = 2.4 \times 10^{-4}$; rs7783012, $P = 4.4 \times 10^{-6}$; and rs786250, $P = 2.6 \times 10^{-5}$) and all of these have concordant signs. These SNPs are all in new loci: none of them tag any previously identified genome-wide associations for ADHD. The Mann-Whitney test of joint enrichment rejects the null hypothesis of no-enrichment ($P$ = 0.008).

None of the four Bonferroni-significant SNPs are located within any of the long-range LD regions and candidate inversions in which we found lead SNPs for all or most of our primary and supplementary GWAS (**Supplementary Information section 3.2**).

We also note that ADHD is significantly associated with our polygenic score of general risk tolerance (**Supplementary Information section 10**) and, as with all second-stage traits we

---

[ff] In an *ex post* analysis, we looked up rs7387531 in the summary statistics of the replication GWAS of self-employment from van der Loos *et al.* (2013)[79]. rs7387531's association with self-employment did not replicate ($P$ = 0.061 on a two-sided test, but with the wrong sign). However, this replication attempt was severely underpowered: the replication sample was small, comprising only the STR cohort ($n$ = 3,271). Further, rs7387531 had an $R^2$ of ~0.004% in the discovery GWAS of general risk tolerance; due to the winner's curse, the true $R^2$ is likely to be smaller than that, and the $R^2$ of rs7387531 on self-employment could be even smaller.

[gg] We assume that higher general risk tolerance leads to a higher probability of being self-employed. For the opposite view, see ref.[246].

analyze here (except height), it is also significantly genetically correlated with general risk tolerance.

###### *8.3.5 Do risk-tolerance-associated SNPs predict bipolar disorder?*

For bipolar disorder, 47 lead SNPs are directly available in the second-stage data, with 49 additional proxy-lead SNPs. Of a total of 96 lead or proxy-lead SNPs, 64 (66.7%) SNPs have concordant signs ($P$ = $7.1 \times 10^{-4}$). There are no second-stage hits. The Mann-Whitney test of joint enrichment fails to reject the null hypothesis of no-enrichment ($P$ = 0.87).

###### *8.3.6 Do risk-tolerance-associated SNPs predict schizophrenia?*

For schizophrenia, 122 lead SNPs are directly available in the second-stage data, with no additional proxy-lead SNPs. Of a total of 122 lead or proxy-lead SNPs, 85 (70.0%) SNPs have concordant signs ($P$ = $8.3 \times 10^{-6}$). There are sixteen second-stage hits, and 13 of these have concordant signs. Four of these SNPs do not tag any loci that were previously identified in published GWAS on schizophrenia (rs13327339, $P$ = $4.8 \times 10^{-5}$ before Bonferroni correction; rs1374197, $P$ = $4.8 \times 10^{-5}$; rs2357023, $P$ = $4.0 \times 10^{-4}$; and rs3764002, $P$ = $3.8 \times 10^{-4}$); two of these four SNPs (rs1374197 and rs2357023) have concordant signs. rs3764002 was also a second-stage hit for ADHD.

Four of the 16 Bonferroni-significant SNPs are located within long-range LD regions or a candidate inversion in which we found lead SNPs in all or most of our primary and supplementary GWAS (**Supplementary Information section 3.2**). The SNP rs3849046 is located within a long-range LD region on chromosome 5 (~135.4 to 138.4 Mb) in which we found two lead SNPs for general risk tolerance, but no lead SNPs for any other main GWAS. The SNP rs1417998 is located in the long-range LD region on chromosome 6 (~25.3 to 33.4 Mb) that covers the HLA-complex, and that region contains lead SNPs for all our GWAS except drinks per week (for which it contains a suggestive association). The SNPs rs624244 and rs1531518 are both located within a candidate inversion on chromosome 18 (~49.1 to 55.5 Mb), and that region is noteworthy because it contains lead SNPs for all our main GWAS.

The Mann-Whitney test of joint enrichment rejects the null hypothesis of no-enrichment ($P$ = $3.0 \times 10^{-9}$) relative to randomly selected SNPs. Although schizophrenia has been shown to be very polygenic[101], the Mann-Whitney test rejects the hypothesis that the observed enrichment is due to polygenic inflation of test statistics over the entire genome. However, it is possible that the overlap between risk tolerance and schizophrenia is explained by enrichment of broad classes of SNPs (e.g., functionally important or conserved regions), rather than specific shared pathways. This is because the Mann-Whitney null distribution is only matched on minor allele frequency; however, the null sample could be further matched based on other attributes (e.g., LD score, functional annotations) to examine whether the joint enrichment is accounted for by these attributes.

These 16 second-stage hits for schizophrenia, together with the strong enrichment of the general-risk-tolerance lead SNPs in the schizophrenia GWAS as well as the high sign concordance of their effects on schizophrenia, suggest that part of the genetic signal for schizophrenia and general risk tolerance may be concentrated in the same genomic regions. We also note that five of our general-risk-tolerance lead SNPs are located in loci that had been found by previous GWAS to be associated with schizophrenia (as we detail in **Supplementary Information section 3.3**).

##### *8.3.7 Do risk-tolerance-associated SNPs predict height?*

For height, 57 lead SNPs are available in the data, along with 42 proxy-lead SNPs. Out of these 99 lead or proxy-lead SNPs, 53 (53.5%) have concordant signs ($P = 0.27$), and eight are second-stage hits. Four of these eight SNPs have concordant sign, and all tag loci with previously identified genome-wide associations. The Mann-Whitney test of joint enrichment rejects the null hypothesis that the second-stage $P$ values are drawn from a null distribution ($P = 1.4 \times 10^{-5}$) and reveals much enrichment (as can be seen from **Extended Data Fig. 8.1**).

These result for height partially contradicting our initial framing of height as a negative control. Such enrichment in the absence of genetic correlation between two traits could be due to a number of reasons[102], including an enrichment of all polygenic traits for certain regions in the genome (e.g., functional or evolutionary conserved regions). As discussed above, the Mann-Whitney null distribution does not account for this form of regional polygenic enrichment. We do not further explore this here, but we note that this may be an interesting question for future research.

### 9 Multi-trait Analysis of GWAS (MTAG)

#### 9.1 Introduction

Because many phenotypes are genetically correlated[24,70], information contained in the GWAS of different but related phenotypes can be used to increase detection and the predictive power of our analysis. We used Multi-trait Analysis of GWAS (MTAG)[103] to increase the precision of the estimates of our GWAS of self-reported general risk tolerance and to improve our ability to detect lead SNPs associated with general risk tolerance. MTAG offers several advantages over lookup-based methods, such as the proxy-phenotype method (**Supplementary Information section 8**); MTAG allows for more than two phenotypes, it increases detection power for all included phenotypes, and it works even in the presence of sample overlap across the various GWAS.

With MTAG, we leveraged the additional information contained in the GWAS summary statistics of six phenotypes related to risk tolerance or risky behavior. The six phenotypes are the five supplementary GWAS phenotypes adventurousness, automobile speeding propensity, drinks per week, ever smoker, and number of sexual partners, as well as lifetime cannabis use[78,hh,ii]. We selected these six phenotypes because they each plausibly capture a different dimension of risk-taking behavior, and they all are significantly genetically correlated with general risk tolerance (**Supplementary Table 7.1**, Panels **A1** and **A3**). Additionally, we conducted an additional MTAG analysis that also included self-employment[79] among those phenotypes, but ultimately decided not to include self-employment in our baseline analysis, as we explain below.

We also used the summary statistics of this MTAG analysis to construct polygenic scores of general risk tolerance and evaluate their power in predicting a suite of phenotypes available in three validation cohorts (**Supplementary Information section 10**).

#### 9.2 Methods

We used the summary statistics from the meta-analysis of the discovery and replication GWAS of general risk tolerance and of the six aforementioned phenotypes[jj] as input for our MTAG analysis.

MTAG builds on the assumption that the correlation in the effect size of a SNP across phenotypes is the same for all SNPs. This assumption is strong and often violated; however, Turley *et al.* (2017)[103] analytically show that, as long as the SNPs have a non-null association with all phenotypes, MTAG is still a consistent estimator with a lower mean squared error than a

[hh] The summary statistics for most of these phenotypes come from GWAS that were conducted in samples that included the UKB and 23andMe cohorts. This should not bias the results because, as mentioned above, MTAG works well in the presence of sample overlap across the various GWAS.

[ii] We do not use the summary statistics from our sixth supplementary GWAS phenotype, the first PC of the risky behaviors, because that would have been redundant given that we already use the summary statistics of the four risky behaviors.

[jj] As usual, we applied genomic control using the intercepts of LD Score regressions to adjust the summary statistics used as input for our MTAG analysis. As the MTAG summary statistics are invariant to any scaling performed on the summary statistics used as inputs for the analysis, whether genomic control has been applied to the input summary statistics should not affect the results. We did not apply genomic control using the intercepts of LD Score regressions to adjust the summary statistics outputted by the MTAG analysis, because the adjustment is already built into the MTAG estimates.

corresponding GWAS. A problem might arise in MTAG for SNPs that have no effect on one phenotype but a sizeable effect on another. This might be the case for SNPs in the proximity of genes implicated in a biological process that is specific to one of the phenotypes, but unlikely to be implicated for general risk aversion. Such examples might include nicotine or cannabis receptors, or alcohol metabolism. We therefore excluded from this analysis all SNPs within 1Mb of the genes *CHRNA5* and *CHRNB3* (nicotinic receptors), *CNR1* and *CNR2* (cannabinoid receptors), and *ADH1B* (Alcohol Dehydrogenase)[kk].

We imposed a MAF filter of 0.01 and a sample size filter $N \geq \frac{2}{3} \times P_{90}(N)$ to the SNPs for all datasets, where $P_{90}(N)$ denotes the ninth decile of the sample size of a dataset. MTAG automatically limits the analysis to the 5,869,552 SNPs present in all data sets.

We also considered adding self-employment[79] to the seven phenotypes in the MTAG analysis, but doing so would have limited the number of shared SNPs to 2,232,479. Below, we also report (in parentheses) the results of the MTAG analysis that also included self-employment (and that therefore included a total of eight phenotypes).

We ran MTAG (July 13, 2017 release) and then clumped the resulting MTAG summary statistics for the general-risk-tolerance phenotype using our main reference panel[33] and PLINK 1.9, with the same thresholds we used for the main GWAS analysis (**Supplementary Information section 2.8**). These thresholds included a primary *P* value threshold ($5\times10^{-8}$), a secondary *P* value threshold ($1\times10^{-4}$), an $r^2$ threshold (0.1), and a SNP window defined in kilobases (1,000,000 kb)[ll].

To assess whether the lead SNPs from the MTAG analysis of general risk tolerance are in loci that have not been identified by previous GWAS of risk tolerance, we repeated the steps we had followed to assess the novelty of the lead SNPs from our discovery GWAS of general risk tolerance (described in **Supplementary Information section 2.10**).

#### 9.3 Results

##### *9.3.1 MTAG summary statistics*

Leveraging the information present in the seven (eight with self-employment) selected phenotypes, MTAG increases the number of independent SNPs reaching genome-wide significance for general risk tolerance from 124 to 312 (225 with self-employment). **Supplementary Table 9.1** reports the 312 lead SNPs. 127 of these are in loci that overlap the 124 loci already identified in our discovery GWAS of general risk tolerance. The top SNP is located in the locus in the *CADM2* gene on chromosome 3 that has previously been identified by Day *et al.*[50] and that has also been associated in a concurrent study on general risk tolerance by

[kk] It is worth noting that SNPs close to the *ADH1B* gene were top-hits in recent GWAS of alcohol consumption (Clark *et al.* (2017)[247] and **Supplementary Table 3.2**). By contrast, SNPs in the proximity of the cannabis receptor genes were not associated with life-time cannabis use at the genome-wide significant level in the GWAS of lifetime cannabis use whose summary statistics we obtained[78]. SNPs close to the nicotine receptors genes have been found to be significantly associated with daily cigarettes smoked but not with ever smoker (TAG *et al.* (2010)[29] and **Supplementary Table 3.2**), although this could be due to the limited statistical power of those GWAS. For precautionary reasons, we still drop those SNPs from the MTAG analysis. None of these regions contain top-hits for the single-trait GWAS of general risk tolerance, and therefore their exclusion should have little bearing on our analysis.

[ll] As noted before, we used a very wide SNP window of 1,000,000 kb, which effectively makes the $r^2$ and *P* value thresholds the only binding parameters for the PLINK clumping algorithm.

Strawbridge *et al.*[51]. A total of 185 of the 312 lead SNPs are novel genome-wide significant associations with general risk tolerance that have not been identified by our discovery GWAS, by Day *et al.*, or by Strawbridge *et al.*

MTAG increases the mean $\chi^2$ for general risk tolerance from 1.81 to 2.21 (1.89 to 2.23 with self-employment). To achieve a similar increase in $\chi^2$, the sample size for the GWAS would have to be increased from 975,353 to 1,452,014 (975,353 to 1,346,482 with self-employment). This increase in signal can be seen by comparing the Q-Q plots for the MTAG analysis with the seven phenotypes and for the meta-analysis of the discovery and replication GWAS of general risk tolerance (**Extended Data Fig. 9.1a**).

**Extended Data Fig. 9.1b** displays the Manhattan plots for the MTAG analysis of general risk tolerance (top panel) and for the discovery GWAS of general risk tolerance (bottom panel), revealing strong similarities. Indeed, the genetic correlation between the summary statistics from the MTAG analysis and from the meta-analysis of the discovery and replication GWAS of general risk tolerance is very close to unity ($\hat{r}_g$= 0.959, *SE* = 0.0025 without self-employment; $\hat{r}_g$= 0.975, *SE* = 0.0016 with self-employment).

##### *9.3.2 Robustness tests*

MTAG can lead to inflated results if the summary statistics for some of the phenotypes come from highly-powered GWAS but have relatively low genetic correlations with the phenotype of interest. Here, this might be the case for the ever smoker phenotype, since the summary statistics of that phenotype have a mean $\chi^2$ statistics of 2.006 but a genetic correlation of only 0.28 with general risk tolerance. However, we repeated the above MTAG analysis without the ever smoker phenotype (and without self-employment), and we still found 299 independent SNPs that reach genome-wide significance, 296 of which are either the exact same SNPs or within loci of the original MTAG analysis. The remaining three SNPs have *P* values close to our *P* value threshold of $5\times10^{-8}$.

##### *9.3.3 Predictive power of general-risk-tolerance polygenic score constructed with the MTAG summary statistics*

We constructed a polygenic score using the summary statistics from the MTAG analysis that used the seven selected phenotypes. This score allows us to measure how the increase in signal translates into increased out-of-sample predictive power for several measures of risk tolerance, personality traits, and risky behaviors in the Add Health and the HRS cohorts. The construction of the polygenic score with the LDpred method[104] (with the Gaussian mixture weight 0.3), the definition of the predicted phenotypes, and the results of these analyses are described in detail in **Supplementary Information section 10**. Comparison of the predictive power of the polygenic scores constructed using the MTAG and GWAS summary statistics are reported in **Extended Data Figs. 10.1-10.2** and in **Supplementary Tables 10.1-10.4**. As expected, the predictive power of the polygenic scores constructed using the MTAG summary statistics is on average slightly higher than that of the scores constructed using only the GWAS summary statistics.

### 10 Predictive power of general-risk-tolerance polygenic score

To assess the out-of-sample predictive power of the genetic variants associated with self-reported general risk tolerance, and to gauge the potential for leveraging these associations in empirical research in the behavioral sciences, we constructed various polygenic scores (PGS) and used them to predict several risk-related phenotypes, personality traits, and real-world measures of risky behaviors.

In this section, we first describe in detail the phenotypes we predicted, which fall in three main domains: measures of risk tolerance (including general risk tolerance), personality traits, and risky behaviors. To be able to analyze several phenotypes in all of these domains, we used six different datasets with contain rich phenotypic information: the Add Health, HRS, NTR, STR[mm], UKB-siblings[nn], and Zurich cohorts.

Next, we delineate and motivate the methodology that we followed to construct the polygenic scores used for prediction. We constructed three polygenic scores in total. Our first two polygenic scores were constructed with the LDpred[104] method, which accounts for the linkage disequilibrium (LD) between SNPs. The first was constructed using the summary statistics from the meta-analysis of the discovery and replication GWAS of general risk tolerance; the second was constructed using the MTAG summary statistics (for details, see **Supplementary Information section 9**). Our third polygenic score was constructed with the classical method, which simply weights SNPs by their GWAS effect size[105,106]. Due to data access limitations, the 23andMe cohort could not be included in the meta-analysis whose summary statistics we used to construct the polygenic scores in the NTR, STR, and Zurich cohorts. The polygenic score using the MTAG summary statistics was only constructed for the Add Health, HRS, and UKB-siblings cohorts.

Across our analyses, we find that the polygenic scores' predictive power is within the range expected according to theory[107,108], when we take into account the SNP heritability of general risk tolerance and cross-cohort heterogeneity. In addition, our polygenic scores are predictive of both general risk tolerance and alternative measures of risk tolerance, such as financial and income gamble risk tolerance. Furthermore, the scores are predictive of a wide variety of other phenotypes: from personality traits such as openness to experience and behavioral inhibition, to real-world economic behaviors such as being an entrepreneur or owning a business, to risky health behaviors such as drinking, smoking, or drug use, to precautionary behaviors such as having life and health insurance. These results are robust to controlling for educational attainment, cognitive performance, and personality traits. Finally, for some of the predicted phenotypes, the predictive power of our polygenic scores is comparable to, or even exceeds, the predictive power of cognitive performance or educational attainment.

[mm] The STR1 and STR2 cohorts were merged for the prediction analysis. See Appendix Section 10.5.3 for details.

[nn] As mentioned in **Supplementary Information section 4**, the UKB-siblings cohort was defined in the same way as the full UKB cohort, but only includes individuals with at least one full sibling in the UKB.

#### 10.1 Phenotype definition

##### *10.1.1 Risk-tolerance phenotypes*

We predict both general-risk-tolerance measures and several alternative measures of risk tolerance. These alternative measures have been developed to elicit risk tolerance in specific domains, such as financial investment decisions or career choices. Three such alternative measures are available in the cohorts we analyze: "financial risk tolerance," "income gamble risk tolerance," and "lottery-elicited risk tolerance." These alternative measures have been shown to correlate with non-hypothetical risky behaviors such as smoking and drinking, being self-employed, or holding a risky investment portfolio[1–5]. As a negative control, we also predicted height in the STR cohort.

###### *(1) General risk tolerance*

The general-risk-tolerance phenotypes we used are comparable to the measures used in the discovery and replication GWAS of general risk tolerance (these measures are described in **Supplementary Information section 1.2**).

In the STR cohort, the following question adapted from the German Socioeconomic Panel[4] was asked to the respondents: *"How do you see yourself: Are you generally a person who is fully prepared to take risks or do you try to avoid taking risks? Please tick a box on the scale, where the value 1 means 'unwilling to take risks' and the value 10 means 'fully prepared to take risks'."* The sample size for general risk tolerance in the STR cohort is 8,012, and the mean response is 4.5 (**Supplementary Table 10.1**).

In the Add Health cohort, the latest wave asks respondents who were 24-34 years old the following question: *"How much do you agree with each statement about you as you generally are now, not as you wish to be in the future?: I like to take risks."* Likert-scale response options include: *[1] strongly agree; [2] agree; [3] neither agree nor disagree; [4] disagree; [5] strongly disagree*. We reverse coded this variable, so that individuals more likely to take risks were coded with a "5" rather than a "1," and vice versa. In total, we have data available for 4,749 European respondents for this variable, and the mean response is 3.0 (**Supplementary Table 10.1**).

In the UKB-siblings cohort we have information on self-reported general risk tolerance. All of the phenotypes predicted in the UKB-siblings cohort are the same as the UKB measures used in our primary and supplementary GWAS. For a detailed description of these phenotypes, see **Supplementary Information section 1.2**.

###### *(2) Financial risk tolerance*

Financial risk tolerance was measured with the following survey question in the STR cohort[10]:

*"Are you a person that is fully prepared to take financial risk or do you try to avoid taking financial risk? Please tick a box on the scale, where the value 1 means: 'not at all willing to take risks' and the value 10 means: 'very willing to take risks'."*

The sample size for financial risk tolerance in the STR cohort is 8,038, with a mean response of 3.5 (**Supplementary Table 10.1**).

##### *(3) Income gamble risk tolerance*

The income gamble risk tolerance phenotype is available in the STR and HRS cohorts. In the STR cohort the income gamble risk question is[109]:

*"Imagine the following hypothetical situation. You are the sole provider for your household, and you have the choice between two equally good jobs:*

*Job A will with certainty give you SEK 25,000 per month after taxes for the rest of your life.*

*Job B will give you a 50-50 chance of SEK 50,000 per month after taxes for the rest of your life, and a 50-50 chance of SEK X per month after taxes for the rest of your life.*

*Which job do you choose?"*

The question was asked three times, and the amount X consecutively takes the values 20,000, 22,000 and 17,000. At the time, one US dollar was worth approximately seven SEK. A respondent receives one point for each time the risky Job B is chosen, so that a value of 0 indicates the lowest risk tolerance, and a value of 3 indicates the highest risk tolerance.

The sample size for income gamble risk tolerance in the STR cohort is 7,577, with a mean response of 2.0 (**Supplementary Table 10.1**).

In the HRS cohort, the income gamble risk question was asked in several waves. Here, we summarize how we created the income gamble risk tolerance variable in the HRS cohort. We provide further details in the Appendix at the end of this section.

In the first wave the income gamble risk question is phrased as follows:

*"Suppose that you are the only income earner in the family, and you have a good job guaranteed to give you your current (family) income every year for life. You are given the opportunity to take a new and equally good job, with a 50-50 chance it will double your (family) income and a 50-50 chance that it will cut your (family) income by a third. Would you take the new job?"*

If the answer is "yes" to the first question, then the respondent is asked this follow-up question:

*"Suppose the chances were 50-50 that it would double your (family) income, and 50-50 that it would cut it in half. Would you still take the new job?"*

If the answer is "no" to the first question, then the respondent is asked this follow-up question:

*"Suppose the chances were 50-50 that it would double your (family) income and 50-50 that it would cut it by 20 percent. Would you then take the new job?"*

In the following waves, the question wording is slightly modified, albeit keeping the same overall structure (see the Appendix at the end of this section for more details). In order to merge the information from different waves, we first coded each variable so that higher values imply higher risk tolerance, and then we took the average of the residuals from a separate OLS regression of each wave's phenotype on an intercept and birth year, birth year squared, birth year cubed, sex, as well as three interaction terms between sex and the three birth year variables.

The sample size for income gamble risk tolerance is 7,302 in the HRS cohort, with a mean answer of 1.8 (**Supplementary Table 10.1**).

##### *(4) Lottery-elicited risk tolerance*

The lottery-elicited risk tolerance phenotype is only available in the Zurich cohort. It was elicited using a method called Multiple Price List (MPL)[4,25,110], which is commonly used in studies on risk tolerance based on incentivized gambles or lotteries. This phenotype was measured in an incentivized computerized experiment, in which respondents were asked lottery questions presented in three separate screens, corresponding to three tables. Each screen shows a table with 20 rows, on which respondents are required to make 20 binary decisions between a fixed lottery (option A) and a certain outcome (option B), which changes from row to row.

In each table, option A offers the possibility to "Win X CHF with a probability of 50% and win Y CHF with a probability of 50%." The following values for X and Y are used for the three tables: Table 1: (20, 10) CHF; Table 2: (20, 0) CHF; and Table 3: (50, 20) CHF. (The expected values for option A are thus 15, 10, and 35 for the three tables). Option B displays the guaranteed win as the interval [X, Y] divided into 20 equal steps presented in descending order on each screen. (Between 10 and 19.5 CHF on table 1, between 0 and 20 CHF on table 2, and between 20 and 48.5 CHF on table 3.) Respondents chose between either option A or B for each of the 20 rows and the software enforced consistency so that respondents could only switch once from option B to option A on each screen. We calculated a screen's certainty equivalent as the average of the two option B wins just above and below the respondents' switching point. Individual i's lottery-elicited risk tolerance measure was then calculated as the median of the calculated relative risk premia (*rrp*) of the three lottery questions:

$$measure_i = median(rrp_{i,j}) = median\left(\frac{(B_{i,j} - E[A_j])}{|E[A_j]|}\right), \tag{8}$$

where $E[A_j]$ is the expected value of option A on screen $j$ and $B_{i,j}$ is the certainty equivalent for individual $i$ on screen $j$. (For the subsamples from Munich and Innsbruck, currencies were converted to EURO according to purchasing power.)

The sample size of the lottery-elicited risk tolerance is 2,610 in the Zurich cohort.

##### *10.1.2 Personality traits*

Personality traits have been defined as "the relatively enduring patterns of thoughts, feelings, and behaviors that reflect the tendency to respond in certain ways under certain circumstances"[111]. Therefore general risk tolerance can be viewed as one of the many traits characterizing an individual's personality, and is certainly very much related to established personality measures such as behavioral inhibition (the tendency to have a restrained or fearful response to unfamiliar events), openness to experience (the degree of intellectual curiosity, creativity, and preference for novelty and variety), or sensation-seeking (the tendency to search for experiences and feelings that are varied, novel, complex, and intense). Indeed several studies at the intersection of economics and personality psychology document the phenotypic correlation between measures of personality traits and general risk tolerance[2–4,112–117]. Motivated by these findings, we investigate the predictive power of our genetic measure of risk tolerance for various phenotypic measures of personality traits.

A large range of measurements of personality traits is available in our datasets, from the widely used five-factor model of personality (the Big Five), to more specific measures like behavioral inhibition and locus of control. Below we provide a detailed description of our personality phenotypes, and we present the summary statistics in **Supplementary Table 10.2**.

##### *(1) Behavioral inhibition*

Behavioral inhibition is defined as a behavioral style characterized by a restrained or fearful response to unfamiliar events, both social and non-social, and has been shown to be moderately stable over time[118]. The 16-item Adult Measure of Behavioral Inhibition (AMBI) battery is available in the STR cohort ($n$ = 7,608) cohort. An example question is: “Do you feel awkward when you are approached by someone new?” Each question is answered on a three-point scale. Summing the responses leads to a variable taking values in the interval [0, 32]. We reverse coded the variable so that a higher score implies a higher behavioral *disinhibition*, since we expect behavioral disinhibition to be positively correlated with general risk tolerance, as has been found in previous studies[2].

##### *(2) Locus of control*

Locus of control is a personality trait defined on a spectrum ranging from internal control, where an individual feels that the outcomes of events are contingent on his or her own behavior and attributes, to external control, where an individual believes that external forces outside of personal control determines the outcome of events. The 12-item locus of control scale is available in the STR cohort for 6,777 individuals. It is coded such that higher numbers indicate an internal locus of control. An example question is: “Becoming a success is a matter of hard work; luck has little or nothing to do with it”[119]. We expect internal locus of control to be positively correlated with general risk tolerance, and this is what has been found in previous studies[2].

##### *(3) Big Five personality traits*

Several decades of research in personality psychology beginning in the 1970s have led to a widely shared taxonomy of personality traits, known as the Big Five[120–122]. Identified via factor analysis of common language descriptors of personality, these five distinct and independent personality characteristics are openness to experience, conscientiousness, extraversion, agreeableness, and neuroticism. Such traits have been measured both in the HRS and in the NTR cohorts.

In the HRS cohort, the Big Five personality traits were measured in four waves between 2006 and 2012, with 26 items in 2006-2008 and 31 in 2010-2012. The original 26 items in 2006 and 2008 were obtained from the MIDUS survey[123]. Extraversion, agreeableness, and conscientiousness were measured with five items, openness to experience with seven, and neuroticism with four. In 2010 and 2012, five items from the International Personality Item Pool[124] were added for conscientiousness. As is common in the literature, the responses to each item were elicited on a four-point scale[121]. For each trait, scores were constructed by taking the mean of all items in the respective category after recoding items to make their direction consistent, and then averaged across waves and standardized to have zero mean and unit variance. A score was set to missing if more than half of the items in that category had missing values. The resulting sample sizes are 8,253 for openness, 8,268 for conscientiousness, 8,271 for extraversion, 8,271 for agreeableness, and 8,264 for neuroticism. Given the limited evidence on the sign of the phenotypic correlation between general risk tolerance and the Big Five traits, we do not have an *ex ante* expectation about the signs of their associations with our polygenic score of general risk tolerance.

In the NTR cohort, the NEO Five-Factor Inventory (NEO-FFI) was sent to adult participants in three waves in 2003, 2009 and 2013. Each of the five personality traits is measured with 12 items, and the responses to each of these were given on a five-point Likert scale. Scores were standardized to have zero mean and unit variance in each wave and then averaged across waves, yielding a sample size of 8,526.

###### *(4) Sensation seeking*

Sensation seeking is defined by the search for experiences and feelings that are "varied, novel, complex and intense," and by the readiness to "take physical, social, legal, and financial risks for the sake of such experiences"[125].

In the NTR cohort, sensation seeking was assessed using a short version of the Zuckerman sensation seeking scale (SSS)[126–128]. The scale consists of 12 items that measure four subdomains of sensation seeking. Thrill and adventure seeking ($n$ = 8,649), defined as the desire to engage in extreme sports and risky activities; experience seeking ($n$ = 8,185), defined as the desire for experiences that engage the mind and senses such as art, travel, food or dress; disinhibition ($n$ = 8,618), defined as the desire to find release through interactions with other disinhibited people, wild parties and sexual disinhibition; and boredom susceptibility ($n$ = 8,621), defined as the intolerance to boredom. A combined total score is then constructed as the total sum of all subscales. The SSS was measured at six time points between 1991 and 2007. Scores were standardized to have zero mean and unit variance in each wave and then averaged across waves.

###### *(5) Attention-deficit hyperactivity disorder*

We also examined the predictive power of our score for attention-deficit hyperactivity disorder (ADHD). ADHD is not a personality trait, but it is nonetheless of interest because it has been shown to be phenotypically correlated with risk-taking behaviors, such as starting one's own business[129–131]. ADHD is a chronic condition of persistent behavioral problems that often begins in childhood and persists into adulthood. The fourth edition of the Diagnostic and Statistical Manual of Mental Disorders (DSM-IV) defines ADHD as "a persistent pattern of inattention and/or hyperactivity-impulsivity that is more frequently displayed and more severe than is typically observed in individuals at a comparable level of development."

In the NTR cohort, symptoms of ADHD were assessed by the Conners ADHD index[132,133] and were measured at three occasions in 2004, 2007 and 2013. The scores were standardized to have zero mean and unit variance in each wave and averaged across waves, yielding a total of 8,457 respondents.

##### *10.1.3 Risky behaviors*

Building on previous studies that demonstrate the predictive power of risk tolerance for various behaviors[1–5,28], we consider a wide range of phenotypes associated with lifestyle risks. We are interested in two categories of real-world behaviors: *risky* and *precautionary*. The first category focuses on behaviors that increase an individual's exposure to unpredictable outcomes and includes smoking, drinking, unhealthy food consumption, financial investment, and career choices. The second category focuses on behaviors that aim to prevent or curtail potential future negative outcomes, and includes investing in health or life insurance as well as preventative health care measures such as taking medications and getting regular medical screenings.

Below we provide a detailed description of our real-world phenotypes. **Supplementary Table 10.3** presents summary statistics. For a more detailed description of measures drawn from the STR cohort, see Beauchamp *et al*. (2015)[2].

##### *(1) Smoking*

Smoking is associated with increased risks of heart disease, stroke, lung cancer (and other types of cancers), and other chronic lung diseases[134]. Smoking has also been shown to be phenotypically correlated with risk tolerance[1–5].

In the HRS cohort, we constructed a binary variable for "ever smoker" that equals "1" if an individual respondent reported ever having been a tobacco smoker in at least one wave. 57% of the 8,617 HRS respondents are coded as ever smokers.

In the STR cohort, respondents are coded as "ever smokers" if they indicated they currently smoke regularly, used to smoke regularly, currently smoke on and off, or used to smoke on and off. 53% of the 13,921 respondents in the STR cohort are coded as smokers.

In the Add Health cohort, respondents are coded as "ever smokers" if they reported having ever smoked regularly (i.e., at least one cigarette every day for 30 days) in Wave 4, the latest wave of the Add Health data. 52.6% of 4,775 European ancestry respondents were coded as smokers. We also constructed a continuous variable for age of smoking initiation among those who were coded as smokers, again in Wave 4. The mean age of smoking initiation in the Add Health cohort is 16.9, and the variable is defined for 2,493 European respondents.

In the UKB-siblings cohort we have information on whether the respondent has ever been a smoker. For a detailed description of the phenotype used, see **Supplementary Information section 1.2**.

##### *(2) Drinking*

Similar to smoking, drinking is a risky behavior known to have negative health consequences[135] and has also been correlated phenotypically with risk tolerance[1,2]. The Add Health, HRS, and STR datasets include a rich set of questions on drinking habits.

In the HRS cohort, we calculated the number of alcoholic drinks per week reported by each respondent in each wave, took the average across waves, and normalized the resulting measure in a variable with a mean of approximately zero and a variance of approximately one[oo]. Importantly, in waves 1 and 2, respondents were only able to choose among 5 categories: 0

[oo] The normalized variable is the inverse normal cumulative distribution function (CDF) of the respondents' percentile ranks. Given the discrete nature of the underlying drinking reports, the inverse normal CDF has some mass points and does not have a mean of exactly zero and a variance of exactly one. This is true of all the other phenotypes that have been normalized.

drinks per week, 1 or fewer drinks per week, 1-2 drinks per week, 3-4 drinks per week, or 5 or more drinks per week. We coded these as 0, 1, 1.5, 3.5, and 5 drinks per week for each wave, respectively. From wave 3 onward, respondents were first asked how many days per week they had *any* alcoholic drinks, and then were asked how many drinks they had on average on each of those days. For these waves, we simply multiplied each of these two responses to calculate drinks per week. This information was available for 8,652 respondents.

Similarly, in Add Health we calculated the number of drinks per week for 3,712 European respondents in Wave 4. Respondents were asked both how many days they consumed alcohol in the last 12 months *and* how many alcoholic drinks they consumed each time they drank in the last 12 months. We multiplied these responses together and divided by 52 to arrive at our final drinks per week variable. The average number of drinks per week in Add Health was 5.6.

In the UKB-siblings cohort we have information on the number of drinks per week. For a detailed description of the phenotype used see **Supplementary Information section 1.2**. Notice that drinks per week was normalized in the HRS but not in the Add Health and UKB-siblings cohorts, where it is simply reported as a count.

In the STR cohort, we constructed a binary variable equal to "1" if respondents reported having alcoholic drinks more than twice in the past month. Specifically, a first question asks respondents if they drank strong beer, wine, or liquor more than twice in the past month. If respondents answer in the negative, a follow-up question asks about their frequency of drinking for each type alcohol separately, and "twice per month" was one of eight response categories. Overall, we classified individuals as alcohol consumers if they answered the first question in the affirmative or if their answers to the follow-up questions indicated that they usually drink beer, wine, or liquor at least twice a month. 82% of the 12,551 respondents in the STR cohort were coded as such.

The STR dataset contains additional information regarding excessive drinking over the life course, which might better capture risky drinking behavior. We constructed a dummy variable equal to "1" for respondents who reported any of the following: 1) ever having a period in their life when they drank too much; 2) ever having a period in their life when someone else objected to their drinking; or 3) ever having a period in their life when they chose drinking instead of working, spending time on hobbies, or spending time with family or friends. 8% of the 13,269 STR respondents were coded as excessive drinkers.

##### *(3) Cannabis consumption*

Similar to cigarettes and alcohol, cannabis abuse can lead to negative health[136] and social consequences[137], especially during adolescence and early adulthood. Furthermore in 2008, at the time of reporting, recreational use of marijuana was illegal in the United States, and therefore its consumption carried the additional risk of breaking the law.[pp]

[pp] Starting with Oregon in 1973, the possession of marijuana was decriminalized by some states, but remained illegal across the United States. Starting with California in 1996, some states legalized the use of marijuana for medical purposes. Recreational use was legalized only starting with Colorado in 2012. While the negative health and social consequences were still prevalent through the sample, the legal consequences of cannabis consumption varied depending on the residence of the respondents, which is unknown to us.

Using data from Wave 4 of Add Health, we constructed a binary variable for "ever cannabis" that equals "1" if an individual respondent reported having ever used cannabis. 60.0% of the 4,742 European respondents had a value of "1" for this variable.

##### *(4) Unhealthy food consumption*

Consumption of fried food is a common risky health behavior that has been linked to cardiovascular disease and other negative health consequences[138]. Questions about fried food consumption were asked to a subset of 3,078 respondents in the HRS cohort who participated in the 2013 Healthcare and Nutrition Mailout. We looked at propensity to eat fried food either at home or as takeout. In reporting how much fried food they consumed, respondents could answer "Never," "Less than once per week," "1-3 times a week," "4-6 times a week," or "Daily." We normalized the measures separately for home and for takeout, resulting in two measures with means of approximately zero and variances of approximately one. Our measure of overall propensity to eat fried food is the sum of these two normalized measures.

##### *(5) Number of sexual partners*

As a final measure of risky health behavior, we consider the total number of sexual partners, which is a risk factor for contracting sexually transmitted infections (STIs)[139,140]. In Wave 4 of Add Health, respondents were asked with how many female and male partners they had ever engaged in any type of sexual activity. We summed the values for female and male partners. The mean number of sexual partners for the 4,603 European respondents was 13.5.

In the UKB-siblings cohort we have information on the number of sexual partners. For a detailed description of the phenotype used, see **Supplementary Information section 1.2.** Notice that number of sexual partners was normalized in the UKB-siblings cohort but not in the Add Health cohort, where it is simply reported as a count.

##### *(6) Automobile speeding propensity*

In the UKB-siblings cohort we have information on automobile speeding propensity. For a detailed description of the phenotype used, see **Supplementary Information section 1.2**.

##### *(7) First PC of risky behaviors*

Following the approach laid out in **Supplementary Information section 1.2**, we computed the first principal component (PC) of our four supplementary risky behaviors (automobile speeding propensity, drinks per week, ever smoker, and number of sexual partners) in the full UKB sample, and then extracted the individuals comprising the UKB-siblings cohort.

##### *(8) Financial market participation*

We consider two measures of exposure to financial risk, both of which have been correlated to phenotypic measures of risk tolerance[1,2,4,28]. The first is financial market participation. This phenotype captures what economists refer to as the "extensive margin" of financial risk-taking behavior (i.e., does one participate in the financial market or not).

In the HRS cohort, we coded a dummy variable that equals "1" if a respondent ever reports a strictly positive "net value of stocks, mutual funds, and investment trusts" across all available waves. Using this measure, 68% of 8,652 HRS respondents participated in financial markets.

Similarly, in the STR cohort we constructed a variable coded "1" if respondents reported a positive value of either stock or bond holdings, using information from the single wave in this cohort when assets were reported. Overall, 30% of the 3,285 STR respondents were coded as participating in the financial market.

***(9) Equity share and share of financial assets***

Our second measure of exposure to financial risk considers the risk profile of respondents' total wealth. We use the share of respondents' total wealth held in equity (i.e., stocks) and similar risky financial assets. This phenotype captures what economists refer to as the "intensive margin" of financial risk-taking behavior (i.e., how much of a person's wealth is exposed to financial market fluctuations).

In the HRS cohort, we calculated for each wave the ratio of "net value of stocks, mutual funds, and investment trusts" over "total wealth including secondary residence." For each respondent, we then averaged this measure across all available waves. This measure roughly reflects the amount of total wealth an individual holds in risky financial assets. We note that mutual funds and investment trusts might include equities, as well as other risky assets such as bonds and derivatives. We refer to this resulting variable as the "share of financial assets." In total, we dropped 50 observations reporting negative equity holdings and two observations for which the share of financial asset in a wave exceeds unity. The mean value of the share of financial assets variable across the 8,599 respondents is approximately 8%.

In the STR cohort, we instead calculate equity share as the ratio of the value of assets held in stocks to the sum of the value of assets held in property, stocks, bonds, bank, boat, and other assets such as jewelry, antiquities, and art. These data are only available for a subset of the respondents, because the associated questions were removed from the survey in the later waves (in an effort to reduce survey length). The 3,170 respondents in STR cohort held on average 3.5% of their total wealth in stocks.

***(10) Career choices***

While some careers are relatively secure and provide a steady stream of income, others are riskier and have greater income fluctuations. Prominent examples of the latter are owning a business or becoming an entrepreneur.

In order to capture this domain of risk taking, which has been correlated to phenotypic measures of risk preferences[2,4,5,28], we considered two separate measures in the STR cohort. First, we constructed a binary variable equal to "1" if respondents reported ever running their own business. 25% of the 7,979 individuals in STR cohort were coded as having ever run their own business.

As a second measure, we consider a self-report of being an entrepreneur, defined as someone who "commercializes a new innovation or idea … [and who] has, or plans to have, a number of employees and strives to expand the business," as opposed to a "self-employed person [who] owns and runs his/her own company, for instance a restaurant or a law firm, where he/she works

[and] … normally does not strive to expand over a certain limit and has 0 or a few employees." This question was asked only of respondents who had reported ever running their own business. We constructed a binary variable equal to "1" if respondents reported being entrepreneurs, and "0" if the respondents reported being self-employed persons or if they reported never having run their own businesses. Based on this measure, 5% of the 7,981 individuals in STR cohort were coded as entrepreneurs.

##### *(11) Preventative healthcare and healthcare utilization*

Focusing now on precautionary behaviors, we first consider choices regarding preventive healthcare.

We constructed an index of preventative health care and health care utilization by combining several available measures for 8,648 respondents in the HRS cohort. Specifically we considered whether respondents had visited a doctor in the last two years, whether respondents had visited a dentist in the last two years, and whether respondents had recently undergone a number of specific procedures or screenings. These included cholesterol screening, flu shot, breast exam (for females), mammogram (for females), pap smear (for females), and prostate exam (for males). We took the simple sum of dummy indicators for having undergone or participated in any of these eight healthcare measures. Since not all of these items apply to both sexes, we normalized these sums separately for men and women, and then combined the measures, obtaining an index with a mean of approximately zero and a variance of approximately one.

##### *(12) Health insurance*

In order to limit the risk associated with an unexpected negative health event, individuals can enroll in health insurance. Prior to the implementation of The Patient Protection and Affordable Care Act (popularly known as "Obamacare" or "Healthcare reform") in the US, this form of risk prevention was an individual choice (rather than required by law) for most individuals. In the HRS cohort, we measure whether individuals choose to have health insurance. As is the case for our other phenotypes, risk tolerance has been shown to predict whether a person has health insurance[1].

According to national and state eligibility criteria, some HRS respondents in certain waves were automatically granted health insurance (i.e., Medicaid or Medicare recipients and CHAMPUS/VA healthcare recipients), and therefore these observations are not used in the construction of this measure for these respondents. For the remaining observations, we calculated whether each respondent-wave observation responded affirmatively to questions about having each of several forms of health insurance (e.g., non-Medicare and non-CHAMPUS/VA federal health insurance, health insurance from current or previous employers, health insurance from spouse's current or previous employers, other health insurance). For each respondent, we then took the average of non-missing observations across all waves until (and including) the 2011 wave[qq]. Our resulting measure indicates the percentage of waves during which a respondent had healthcare, among waves during which it was an optional choice. On average, the 8,454 respondents in the HRS cohort were covered by health insurance 95% of the time.

---

[qq] The individual mandate of the Affordable Care Act of 2010 was not yet in effect during 2011, the final wave from which we draw these data.

*(13) Life insurance*

Finally, individuals purchase life insurance to mitigate the negative consequence of their deaths for their families and loved ones. Phenotypic risk tolerance has been shown to predict whether a person has life insurance[1].

We measured whether respondents are consistently covered by life insurance by calculating the percentage of waves during which HRS respondents reported having life insurance. On average, the 8,652 respondents in the HRS cohort report being covered by life insurance more than two-thirds (71%) of the time.

#### 10.2 Methods

##### *10.2.1 Polygenic scores*

We constructed three polygenic scores. We constructed our first polygenic score with the LDpred[104] method, using summary statistics from our GWAS of general risk tolerance; we also constructed our second score with the LDpred method, but using the summary statistics of general risk tolerance outputted by the MTAG analysis (**Supplementary Information section 9**); and we constructed our third score with the classical method, using summary statistics from our GWAS of general risk tolerance. We will refer to these scores as LDpred-GWAS, LDpred-MTAG, and Classical-GWAS scores respectively.

Due to data access limitations and to avoid overfitting, different cohorts were included in the GWAS whose summary statistics were used to construct the scores. In the Add Health and HRS cohorts, the summary statistics from the meta-analysis of the discovery and replication GWAS of general risk tolerance were used to construct the scores, with no cohort excluded ($n$ = 975,353). For the analyses in the NTR and Zurich cohorts, the scores were constructed with the summary statistics from a meta-analysis that excluded the 23andMe cohort (due to data access limitations, the sample size of the resulting meta-analysis was $n$ = 466,571). For the analysis in the STR cohort, we excluded the 23andMe cohort (due to data access limitations) as well as the STR cohort itself (to avoid overfitting[141]) from the meta-analysis whose summary statistics we used to construct the scores (the sample size of the resulting meta-analysis is $n$ = 458,558)[rr]. And for the analysis in the UKB-siblings cohort, to avoid overfitting[141] we reran our discovery GWAS after excluding all individuals in the UKB-siblings cohort from the UKB cohort (the sample size of the resulting meta-analysis was $n$ = 937,353).

The prediction analyses with our LDpred-MTAG score was only conducted in the Add Health, HRS, and UKB-siblings cohorts, since these were the only three cohorts for which 23andMe data could be used, due to data access limitations.

Both the LDpred and the classical polygenic scores are calculated as the weighted sum of the $M$ individual genotypes:

(9)
$$\hat{S}_i = \sum_{j=1}^{M} \hat{\beta}_j \, g_{ij} \, ,$$

[rr] We also note that, since the HRS, NTR, and Zurich cohorts do not contain data on general risk tolerance, none of these cohorts was included in any meta-analysis of general risk tolerance.

where $\hat{S}_i$ denotes the polygenic score of individual $i$, $\hat{\beta}_j$ is the estimated additive effect size of the effect-coded allele at SNP $j$, and $g_{ij}$ is the genotype of individual $i$ at SNP $j$ (coded as having 0, 1, or 2 instances of the effect-coded allele)[105].

For the LDpred scores, the estimated additive effect sizes (the $\hat{\beta}_j$'s) are the estimates from the summary statistics adjusted for LD between the SNPs. To calculate LDpred scores, an LD reference file and a validation reference file must be provided. For HRS, STR, UKB-siblings, and the Zurich cohorts we used the 1000 Genomes-imputed data (Phase 1, Version 3) of the HRS[ss] cohort as the reference sample. For the Add Health cohort, we used the HRC (Haplotype Reference Consortium) Genomes-imputed data (Version 1.1) of Add Health (n=4,775) as the reference sample. For the NTR cohort, the reference sample used for the construction of the LDpred scores consists of all five European populations from the 1000 Genomes dataset: Utah Residents (CEPH) with Northern and Western European Ancestry, Finnish, British, Iberian, and Toscani individuals ($n = 381$).

The LDpred method relies on a Gaussian mixture weight that corresponds to the assumed fraction of SNPs that are causal. We used the software called LDpred[104] to generate a score for each of the following mixture weights: 1, 0.3, 0.1, 0.03, 0.01, 0.003, 0.001, 0.0003, and 0.0001, using the individuals' best guess data and using only SNPs in the HapMap consortium phase 3 release[142,143] with a MAF > 0.01 and an imputation quality of more than 0.7. We report only the analyses with the LDpred scores constructed with the Gaussian mixture weight 0.3, as these scores tended to perform better both across cohorts and predicted phenotypes.

For the classical polygenic scores, the estimated additive effect size $\hat{\beta}_j$ for SNP $j$ is the GWAS estimate for SNP $j$. We used the software PLINK[37] to produce the classical scores, using the individuals' best guess data; as with the LDpred method, we only use SNPs in the HapMap consortium phase 3 release[142] with a MAF > 0.01 and an imputation quality of more than 0.7 (the prediction results are thus comparable across the two methods)[tt].

We expect our LDpred-GWAS score to have greater predictive power than our Classical-GWAS score, since the LDpred method takes into account the non-independence between SNPs. Further, since the MTAG summary statistics leverage the additional information contained in the GWAS of genetically correlated phenotypes, we expect our LDpred-MTAG score to have the greatest predictive power.

---

[ss] The HRS genotype data used as a reference sample for the STR and the Zurich cohorts was restricted according to the following quality control criteria. We removed 13,973 SNPs that have been flagged as having incorrect annotations from the HRS cohort[248]; we restricted the reference file to SNPs with imputation quality greater than 0.7, MAF greater than 0.01, SNP call rate greater than 95%; and we removed individuals with a call rate less than 95%, as well as related individuals and individuals not of North-Western European ancestry. The resulting sample contained 7,302 individuals.
For the HRS and UKB-siblings cohort, we restricted the reference file to HapMap3 SNPs with MAF greater than 0.01 and SNP call rates greater than 98%; we also removed individuals with a call rate less than 98%, related individuals, and individuals not of European ancestry. The resulting sample contained 8,353 individuals.

[tt] The classical score is often calculated with an LD-pruning and *P* value thresholding procedure. The LD-pruning is meant to achieve independence between the predictive set of SNPs, and the *P* value threshold excludes SNPs that are not estimated to be significantly associated with the phenotype. This procedure often discards information that could increase the predictive power if it were properly accounted for[104]. Here, we do not use an LD-pruning and *P* value thresholding procedure to drop SNPs prior to constructing the scores.

##### *10.2.2 Main prediction exercise*

We ran two separate Ordinary Least Squares (OLS) regressions for each phenotype, cohort, and score. The first regression includes only our baseline controls: sex, birth year, birth-year squared, birth-year cubed, as well as the interactions between sex and the three birth-year variables, and the first ten principal components (PCs) of the cohort-specific genetic relatedness matrix; in the NTR and STR cohorts, dummy variables that indicate the different genotype platforms were also included. The second regression is identical to the first one, except that it also includes the polygenic score. Since the scores have mean zero and unit variance, the estimated coefficients on the score represent the change in real-word outcomes associated with a one-standard-deviation increase in the polygenic score of general risk tolerance. Our measures of interest are the regression coefficients on the scores and the incremental $R^2$ (or pseudo-$R^2$) of the scores, defined as the difference between the $R^2$ (or pseudo-$R^2$) of the two regressions. The 95% confidence intervals for the incremental $R^2$ estimates are calculated with the bootstrap percentile method, with 1,000 bootstrap samples[144,145].

##### *10.2.3 Robustness to inclusion of additional control variables*

Are these polygenic scores of general risk tolerance potentially useful for the empirical researcher in the behavioral sciences? Since many social-science datasets contain rich information about the individual characteristics of the participants, it is important to understand whether the polygenic scores for general risk tolerance are still predictive of risky behaviors even after controlling for relevant variables that capture important individual characteristics. For this reason, when analyzing real-world risky behaviors, we also estimate the incremental $R^2$ (or pseudo-$R^2$) of the scores while controlling for cognitive performance, personality traits, and educational attainment in the baseline regression.

These traits were selected as additional controls because they could plausibly be important determinants of risky and precautionary behaviors, and because many of these traits are phenotypically correlated with risk tolerance[2]. This robustness exercise also makes it easier to compare our prediction results with the existing literature, which usually controls for some of these personal characteristics[6,110,146,147].

Furthermore, we also calculated the incremental $R^2$ for cognitive performance, for personality traits, and for educational attainment, defined as the difference between the $R^2$ of a regression including only our baseline controls, and another regression including the baseline control and the traits of interest. These three incremental $R^2$ values provide the applied researcher with interesting benchmark comparisons, and enable us to understand whether the predictive power of our polygenic score is comparable to the predictive power of other important individual characteristics.

For the Add Health cohort, we control only for educational attainment and a measure of verbal cognition, since information on personality traits is available only for a small subsample of individuals. The measure of verbal cognition is a modified version of the Peabody Picture Vocabulary Test which was collected in the first wave of Add Health, when participants were 12 to 20 years old. In this test, respondents have to select the illustration that best fits the meaning of the word that an interviewer has just read aloud. This computer-adapted test consisted of eighty-seven items, and scores were age-standardized.

For the HRS cohort, the cognitive performance measure is the average of measures from waves 2-10. In each of these waves, the cognitive performance measure is the sum of a total word recall summary score (based on immediate and delayed word recall tasks) and a mental status summary score (based on counting, naming, and vocabulary tasks)[uu]. The Big Five factors (defined above) were used as measures of personality traits[vv], and years of education as a measure of educational attainment.

For the STR cohort, the standardized score on a military conscription test was used as a measure of cognitive performance, behavioral inhibition and locus of control were used as measures of personality traits, and years of education was used as a measure of educational attainment. The cognitive performance measure was merged using conscription data provided by the Military Archives of Sweden. Men were required by law to participate in military conscription around the age of 18. We use the stanine scores of four subtests of logical, verbal, spatial, and technical ability. Following Rietveld *et al.* (2014)[80], we use the first principal component of these four stanine scores as the measure of cognitive performance. No measure of cognitive performance is available for females.

Finally, for the UKB-siblings cohort we control for educational attainment, neuroticism, and cognition. Our measure of neuroticism follows Okbay *et al.* (2016)[18]: it is constructed as a respondent's score on a 12-item version of the Eysenck Personality Inventory Neuroticism scale. Individuals must answer at least 10 out of 12 binary-response questions to be included, and questions that remain missing will be coded as 0. To obtain a measure of cognition, we used a test designed to measure fluid intelligence. The test consists of thirteen logic and reasoning questions and has a two-minute time limit. Each respondent took the test up to four times. We used the mean of the standardized score across the occasions on which the respondent took the test as our measure of cognition.

##### *10.2.4 Methodology for binary and censored phenotypes*

For the analysis of some real-world risky behaviors, we used specific statistical methods to model more accurately the distribution of the dependent variable. For binary outcomes, such as engaging in smoking, drinking, or participating in financial markets, we estimated Probit models. Instead of the estimated coefficient, which has no clear interpretation in a Probit model, we report the average marginal effect of the polygenic score. We report this as $\frac{\partial \Pr(y=1|PGS,X)}{\partial PGS} = \hat{\beta}_{PGS} \cdot \hat{E}\left[\phi(PGS \cdot \hat{\beta}_{PGS} + X\hat{\beta})\right]$, where $\hat{\beta}_{PGS}$ is the estimated coefficient on the polygenic score, $\phi$ is the Gaussian probability density function, $\hat{E}[\cdot]$ takes the sample average, $PGS$ is the polygenic score, and $X$ contains the control variables. In the absence of a meaningful $R^2$ measure for the Probit model, we used the McFadden pseudo-$R^2$, $\left(\frac{l_0 - l_M}{l_0}\right)$, where $l_0$ is the log-likelihood of a model with only a constant, and $l_M$ is the log-likelihood of the full model. Our measure of the incremental pseudo-$R^2$ is thus $\frac{l_X - l_{X,PGS}}{l_0}$, which indicates the difference between the log-likelihood of the model controlling for baseline controls and the score $(l_{X,PGS})$ and the log-likelihood of the model controlling only for the baseline controls $(l_X)$, scaled by the log-likelihood of a model with only a constant $(l_0)$.

[uu] In Wave 2, these measures are present only for a subset of respondents (the AHEAD cohort).
[vv] See above for a detailed description of the personality variables in the HRS cohort.

For equity share and share of financial assets phenotypes, we estimate Tobit models instead of OLS regressions, to account for the point mass at zero (due to many respondents in our dataset not holding any stocks)[ww]. The same measure of incremental pseudo-$R^2$, $\left(\frac{l_X - l_{X,PGS}}{l_0}\right)$, is reported[xx].

The 95% confidence intervals for all of these estimates are calculated with the bootstrap percentile method, with 1,000 bootstrap samples[144,145].

#### 10.3 Results

Results are presented in **Extended Data Figs. 10.1-10.2** and **Supplementary Tables 10.1-10.4**. For each predicted phenotype, cohort, and score, these tables report summary statistics for the regression sample, the estimated incremental $R^2$ of the score, and the sign and $P$ value of the estimated regression coefficient on the polygenic score.

Overall, we find that our preferred polygenic score explains 1.01% to 1.78% of the variation in general risk tolerance, up to 1.4% of the variation in several personality traits, and up to 1.94% of variation in real-world behaviors. As we discuss in **Supplementary Information section 10.4**, our incremental $R^2$ estimates from the prediction of general risk tolerance fall within the range we expect based on theory[107,108].

Below, we focus our discussion on the results for the LDpred-GWAS score. As mentioned above, the results displayed for the LDpred scores are based on a Gaussian mixture weight of 0.3. We chose this weight because the corresponding scores consistently performed well across cohorts and phenotypes in our analyses. The results for the LDpred-MTAG and Classical-GWAS scores are generally similar, although the predictive power of the LDpred-MTAG score tends to be slightly higher and that of the Classical-GWAS score tends to be slightly lower.

##### *10.3.1 Risk tolerance*

In the UKB, the incremental $R^2$ of the LDpred-GWAS score is 1.62% (CI: 1.37% - 1.90%). In the Add Health cohort we estimate a much smaller predictive power, with incremental $R^2$ of 1.01% (CI: 0.57% - 1.62%). In the STR[yy] cohort we estimate an incremental $R^2$ of 1.05% (CI: 0.64% - 1.50%). Using the summary statistics from the MTAG analysis improves our predictive power: the incremental $R^2$ of the LDpred-MTAG score is 1.78% in the UKB-siblings cohort and 1.07% in the Add Health cohort.

---

[ww] The Tobit model is a maximum likelihood estimator proposed by James Tobin[249] that assumes a linear relationship between regressors $X$, a normally distributed error term $\varepsilon$, and a continuous latent variable $y^*$. The observed dependent variable is $y = y^*$ whenever $y^* > 0$ but is $y = 0$ otherwise. This model is used to account for censoring and a point mass at zero, as is observed for our measures of equity share and of share of financial assets.

[xx] Because the Tobit model has a continuous likelihood, the McFadden pseudo-$R^2$ can sometimes be smaller than zero or greater than 1, which is nonsensical for an $R^2$ measure. Whenever that happened, such as in the case of share of financial assets in the HRS cohort when cognitive performance or educational attainment are controlled for, we reported "N.A". in the table of results.

[yy] As mentioned above, due to data access limitations, the summary statistics used to construct the score in the STR cohort were based on a meta-analysis that excluded the 23andMe cohort, and therefore had a much smaller GWAS sample size ($n_{GWAS}$ of 466,571).

The estimated regression coefficients for all scores have the expected sign and are all highly statistically significant (with $P$ values < 0.0005), indicating that the scores significantly predict general risk tolerance out of sample.

##### *(1) Alternative measures of risk tolerance*

The predictive power of our polygenic score for alternative measures of risk tolerance is also significant, but limited in magnitude.

For the financial risk tolerance phenotype, the incremental $R^2$ of the LDpred-GWAS score in the STR cohort is 0.44%. For the income gamble risk tolerance phenotype, the incremental $R^2$ is 0.24% in the HRS cohort, and 0.34% in the STR cohort. When predicting either financial risk tolerance or income gamble risk tolerance, the coefficients on the LDpred-GWAS scores are always highly statistically significant.

By contrast, we find that our polygenic scores of general risk tolerance do not predict the lottery-based measure of risk tolerance in the Zurich cohort. For this measure, the incremental $R^2$ is 0.09% and the coefficient on the score is not distinguishable from zero ($P$ = 0.12).

The lack of predictive power of our polygenic score for lottery-based risk tolerance is surprising, as this phenotype is considered the workhorse of risk preference measurement in economic theory[110]. It is important to note, however, that this null result does not necessarily mean that our polygenic score fails to capture variation in economically relevant risk preferences, given its predictive power for risky economic behaviors such as entrepreneurship and financial risk taking. Indeed, the predictive power of lab-based measures for real world behavior has itself come under scrutiny[148,149], and lottery-based measures of risk preferences usually have lower predictive power than self-reported measures[4,6,146,147]. Further, recent research in both economics and psychometrics has found modest levels of correlations between lottery-elicited measures of risk tolerance and the general-risk-tolerance phenotype used in our discovery and replication GWAS[4,6]. Our results thus add to the small but growing body of evidence that lottery-based and self-reported measures of risk tolerance may indeed capture different aspects of decision-making over uncertainty.

##### *10.3.2 Personality traits*

Our results provide interesting evidence regarding the complex and interlacing structure of personality traits and their relationship to general risk tolerance. Genetic predisposition for risk tolerance is predictive of openness to experience, sensation seeking, extraversion, behavioral inhibition, and to a lower extent ADHD. Not surprisingly, these domains are the ones that have been theoretically and phenotypically connected to an individual's behavior in new, unexpected, and uncertain circumstances. Furthermore, these results corroborate our findings of positive genetic correlations between general risk tolerance and openness, extraversion, and ADHD (**Supplementary Information section 7.3**).

Looking at the Big Five personality traits, our LDpred-GWAS score is predictive of openness to experience (with incremental $R^2$ estimates of 1.45% and 0.42% in the HRS and NTR[zz] cohorts, respectively), extraversion (0.94% in HRS and 0.17% in NTR), and to a lower extent

[zz] As mentioned above, the scores in the NTR cohort were also based on a meta-analysis that excluded the 23andMe cohort.

agreeableness (0.09% and 0.11%). The estimated coefficients on the LDpred-GWAS scores for these three phenotypes always have a $P$ value ≤ 0.4%. In contrast, conscientiousness was never predicted by our polygenic score, and neuroticism was significantly predicted in the HRS cohort (incremental $R^2$ of 0.31%), but not in the NTR cohort.

The results described above are broadly consistent with the estimated sign and magnitude of the genetic correlations between general risk tolerance and extraversion, openness, and neuroticism (**Supplementary Information section 7.3**).

Finally, our LDpred-GWAS score was predictive of sensation seeking (with incremental $R^2$ estimates of 1.13% for the composite sensation seeking phenotype, and ranging from 0.33% to 0.75% for the four sensation seeking subscales), behavioral inhibition (0.50%), and ADHD (0.50%). Furthermore, the estimates of the coefficients on the scores are all highly significant and positive, as expected. No score is predictive of locus of control. Using the summary statistics from the MTAG analysis does not improve our predictive power substantially.

###### *10.3.3 Risky behaviors*

###### *(1) Main prediction results*

**Supplementary Table 10.3A** reports the predictive power of the scores for risky behaviors, while **Supplementary Table 10.3B** reports the predictive power of the scores for precautionary behaviors.

Overall across all our validation cohorts, the predictive power of the LDpred-GWAS scores is low but positive across a wide range of risky and precautionary behaviors, with incremental $R^2$ estimates ranging from 0.02% to 1.36%. Although most incremental $R^2$ estimates are low, the estimated coefficients on the LDpred-GWAS score have the expected sign for 22 of the 25 regressions we ran ($P = 8\times 10^{-5}$), and are always in the expected direction whenever the estimated coefficient is significant at the 5% level (16 of the 25 measures, $P = 2 \times 10^{-15}$)[aaa].

Our LDpred-GWAS score are predictive of economic behaviors such as being an entrepreneur (1.36%) or owning a business (0.57%), as well as health behaviors such as number of sexual partners (0.65% and 0.79% in Add Health and UKB-siblings), automobile speeding propensity (0.59%), ever using cannabis (0.37%), smoking (incremental $R^2$ ranging from 0.10% to 0.25% depending on cohort and phenotype), drinking (incremental $R^2$ ranging from 0.02% to 0.19%, depending on cohort and phenotype), and the first PC of risky behaviors (0.96%). For precautionary behaviors, both life and health insurance coverage are significantly predicted by our LDpred-GWAS scores, with an incremental $R^2$ of 0.17% and 0.10%, respectively. In contrast, our scores are not significantly predictive of financial market participation, equity share or share of financial asset, age of smoking initiation, fried food consumption, and preventative health care utilization.

Several results stand out. The estimated coefficient imply that a one-standard-deviation increase in the polygenic score is associated with a 1.6 percentage points higher probability of being an entrepreneur, 3.4 percentage points higher probability of having owned a business, 2.2 additional

[aaa] These $P$ values refer to the probability of finding this many concordant signs or significant coefficients; they are derived from one-tailed binomial tests (the first coming from 25 coin flips with probability of 0.5; the second from 25 coin flips with probability of 0.05).

lifetime sexual partners, 3.5 percentage points higher probability of ever having smoked marijuana, and a 2 to 3 percentage points higher probability of ever being a smoker.

Using the MTAG summary statistics to construct the LDpred scores in the Add Health, HRS, and UKB-siblings cohorts generally increases our predictive power. For example, the incremental $R^2$ estimates for the first PC of risky behaviors and for number of sexual partners double, to 1.96% and 1.53% respectively. In the case of ever using cannabis, the incremental $R^2$ increases by ~0.40% relative to using the LDpred-GWAS and Classical-GWAS scores.

###### *(2) Robustness to inclusion of additional control variables*

To further assess the potential for using polygenic scores in empirical research, we investigated whether the predictive power of our polygenic scores is robust to including in the regressions additional control variables for cognitive performance, personality traits, and educational attainment. We obtained very similar point estimates but larger confidence intervals than in the regressions that do not include these additional variables. **Supplementary Table 10.4** displays these results. Note that, due to missing observations in the additional control variables, the sample sizes decrease (in particular, the samples for the STR cohort includes males only, as the cognitive performance measure is only available for males). The pattern of results is nonetheless very similar, though the confidence interval of the estimated coefficients and incremental $R^2$ values generally increase, reducing our ability to distinguish them from zero. Above and beyond the measures of personal characteristics, our polygenic scores of risk tolerance are still significantly predictive of being an entrepreneur, owning a business, the first PC of risky behaviors, the number of sexual partners, automobile speeding propensity, ever using marijuana, ever being a smoker, being an excessive drinker and drinks per week (in the STR and UKB-siblings cohorts only), and having health or life insurance. Just as for the results displayed in **Supplementary Table 10.3**, LDpred scores constructed using either the GWAS or the MTAG summary statistics are usually more predictive than Classical-GWAS scores.

Furthermore, the predictive power of our polygenic scores for many risky behaviors is comparable in magnitude to that of cognitive performance, personality, and educational attainment. The estimated 95% confidence intervals of the incremental $R^2$ of our polygenic scores almost always contain the estimated incremental $R^2$ of cognitive performance, educational attainment, or personality traits.

##### *10.3.4 Height as a negative control*

Neither the LDpred-GWAS nor the Classical-GWAS scores significantly predict height in the STR cohort (**Supplementary Table 10.1**). In the UKB-siblings cohort, the incremental $R^2$ are indistinguishable from zero but, given the large sample size, the coefficients of the LDpred-GWAS and LDpred-MTAG scores are statistically different from zero. These observations, combined with the above results, suggest that our scores capture some true polygenic signal for general risk tolerance and that our above results are not artifacts of our estimation procedure or of chance.

#### 10.4 Expected predictive power of general-risk-tolerance polygenic score

As mentioned before, the predictive power of our polygenic scores is within the range expected according to theory, when cross-cohort heterogeneity of the GWAS and predicted phenotypes' SNP heritabilities are taken into account. Daetwyler *et al.*[107] have developed a theoretical formula for the expected predictive power of a polygenic score, when predicting the phenotype whose GWAS summary statistics were used to construct the score in an independent cohort. The formula assumes independent SNPs, equal heritability of the phenotype in the discovery and validation cohorts, and perfect genetic correlation of the phenotype across cohorts. Daetwyler *et al.*[107] show that, under those assumptions, the expected predictive power of a polygenic score is equal to:

$$E(R^2) = h^2 \left( \frac{1}{1 + 1/\lambda h^2} \right)$$

where $h^2$ is the SNP heritability of the phenotype (which is assumed to be the same in the validation cohort and in the GWAS whose summary statistics are used to construct the score), and $\lambda = n_{GWAS}/M$ is the ratio of the sample size in the discovery GWAS ($n_{GWAS}$) and the effective number of SNPs evaluated in the validation cohort ($M$). For the discovery meta-analysis we use in our Add Health ($n_{\text{GWAS}} = 975{,}353$) and UKB-siblings ($n_{\text{GWAS}} = 937{,}353$) analyses, $h^2$ is 0.041 when estimated by LD Score regression (**Supplementary Table 4.1**) and 0.045 when estimated using HESS (**Table 1**); in the STR cohort ($n_{\text{GWAS}}$ = 458,558), $h^2$ is 0.055 when estimated by LD Score regression and 0.063 when estimated by HESS (**Supplementary Table 6.1**)[bbb]; and a reasonable value for $M$ is 60,000, which is the midrange of 50,000 to 70,000 suggested by Wray *et al.* 2013[141].

Given a range of $h^2$ values between 0.041 and 0.045 (0.055 and 0.063 for the STR cohort), this formula suggests an expected predictive power between 1.60% to 1.86% in the UKB-siblings cohort, 1.64% to 1.90% in the Add Health cohort, and between 1.63% and 2.05% in the STR cohort. Our estimated incremental $R^2$'s for general risk tolerance are 1.62% (CI: 1.37% – 1.90%) in the UKB-siblings cohort and ~1.0% in both the Add Health and STR cohorts—slightly less than what we would expect given the Daetwyler formula.

One explanation for the relative underperformance of our polygenic scores is cross-cohort heterogeneity. To account for this, De Vlaming *et al.*[108] generalize the Daetwyler formula by allowing for unequal heritability between the prediction ($h_p^2$) and discovery cohorts ($h_D^2$) and imperfect genetic correlation ($r_g < 1$), while still assuming independent SNPs. With these generalizations, the expected predictive power of a polygenic score becomes:

$$E(R^2) = r_g^2 h_p^2 \left( \frac{1}{1 + 1/\lambda h_D^2} \right).$$

As before, we let $h_p^2$ and $h_D^2$ range between 0.041 and 0.045 (0.055 and 0.063 for the STR cohort). For the UKB-siblings cohort, we estimate a genetic correlation of 0.93 between the

[bbb] These heritability estimates for the STR cohort are those for the UKB cohort only (and not those for the meta-analysis of the UKB and replication cohorts whose summary statistics were used to construct the scores in the STR cohort). These estimates are higher because they are not attenuated by cross-cohort heterogeneity, unlike the heritability estimates for the GWAS whose summary statistics were used in the Add Health and UKB-siblings analyses.

discovery and validation cohorts (using the full UKB GWAS instead of a GWAS of the UKB-siblings subset to increase precision of this estimate). For the Add Health and STR cohorts, where sample sizes are too low to permit direct estimation of the genetic correlation, we consider $r_g$ values between 0.7 and 0.9.

The expected incremental $R^2$ now ranges between 1.51% and 1.61% for the UKB-siblings cohort, 0.80% to 1.54% for the Add Health cohort, and between 0.80% and 1.66% for the STR cohort. The observed predictive power of our score is now consistent with what we would expect in theory. This demonstrates the importance of considering cross-cohort heterogeneity when forming expectations of the performance of polygenic scores.

De Vlaming *et al.*[108] additionally allows for imperfect genetic correlations between individual cohorts within the discovery sample. When we account for this within-discovery sample heterogeneity (using the estimated genetic correlations reported in **Supplementary Information section 7.4.2**, which range from 0.76 to 0.83), our expected $R^2$ ranges become 1.44%% – 1.67% for the UKB-siblings cohort, 0.84% – 1.62% for the Add Health cohort, and 0.80% – 1.67% for the STR cohort. These ranges remain consistent with our observed $R^2$ estimates.[ccc]

Lastly, our polygenic score performs consistently better in the UKB-siblings validation cohort than in the Add Health and STR validation cohorts, given our expected $R^2$ values. This discrepancy could be explained by cross-cohort heterogeneity that we were unable to account for. For example, we have assumed throughout that $h_p^2 = h_D^2$; however, the heritability of risk tolerance may be systematically lower in the Add Health and STR cohorts, where direct heritability estimation was not possible.

Thus, while polygenic scores constructed from our GWAS results can already aid the study of some related phenotypes, future research would benefit from the collection of measures of risk tolerance that are more similar across cohorts.

[ccc] We do not account for imperfect genetic correlations between cohorts within our replication meta-analysis; these cohorts only account for 3.5% of the individuals in our discovery meta-analysis and are unlikely to affect our results.

#### 10.5 Appendix: definition of the income risk gamble variable in the HRS cohort and merger of the STR1 and STR2 cohorts

##### *10.5.1 Income gamble risk tolerance in the HRS cohort*

In the HRS cohort the income gamble risk question was slightly modified after the first wave[109]. In the first wave, respondents were first asked the following question:

*"Suppose that you are the only income earner in the family, and you have a good job guaranteed to give you your current (family) income every year for life. You are given the opportunity to take a new and equally good job, with a 50-50 chance it will double your (family) income and a 50-50 chance that it will cut your (family) income by a third. Would you take the new job?"*

Respondents who answered "yes" to the first question were asked this follow-up question:

*"Suppose the chances were 50-50 that it would double your (family) income, and 50-50 that it would cut it in half. Would you still take the new job?"*

Respondents who answered "no" to the first question were asked this follow-up question:

*"Suppose the chances were 50-50 that it would double your (family) income and 50-50 that it would cut it by 20 percent. Would you then take the new job?"*

In the original variable coding, the variable was coded as 1 if the respondent accepted the new job with the risk of cutting income by half, and a value of 4 if the respondent always chose to stay with the job with a guaranteed income. We reverse coded the original phenotype so that a higher value implies higher risk tolerance.

The income gamble risk tolerance question was not asked in waves 2 and 3. In wave 4 and all subsequent waves, the initial question was:

*"Suppose that you are the only income earner in the family. Your doctor recommends that you move because of allergies, and you have to choose between two possible jobs. The first would guarantee your current total family income for life. The second is possibly better paying, but the income is also less certain. There is a 50-50 chance the second job would double your total lifetime income and a 50-50 chance that it would cut it by a third. Which job would you take -- the first job or the second job?"*

If the first job was chosen as the answer to the initial question, then this follow-up question was asked:

*"Suppose the chances were 50-50 that the second job would double your lifetime income and 50-50 that it would cut it by twenty percent. Would you take the first job or the second job?"*

If the first job was chosen as the answer to the follow-up question, then a second follow-up question was asked:

*"Suppose the chances were 50-50 that the second job would double your lifetime income and 50-50 that it would cut it by 10 percent. Would you take the first job or the second job?"*

If instead the second job was chosen as the answer to the initial question, then a different follow-up question was asked:

*"Suppose the chances were 50-50 that the second job would double your lifetime income, and 50-50 that it would cut it in half. Would you take the first job or the second job?"*

If the first job was chosen as the answer to the follow-up question, then a second follow-up question was asked:

*"Suppose the chances were 50-50 that the second job would double your lifetime income and 50-50 that it would cut it by seventy-five percent. Would you take the first job or the second job?"*

In the original variable coding, the respondent is given a value of 1 if the respondent accepts the second job with the risk of cutting the income by seventy-five percent, and a value of 6 if the respondent always chooses to stay with the job with a guaranteed income. Again, we reverse coded the responses so that a higher value corresponds to a higher risk tolerance. Further, since this survey measure has two additional response categories relative to the measure used in the first wave, we converted this survey measure's responses to the 4-point scale that disregards the second follow-up questions; a value of 5 or 6 corresponds to 4, a value of 4 corresponds to 3, a value of 3 corresponds to 2, and value 1 or 2 corresponds to 1.

We constructed the income gamble risk tolerance phenotype as the average response across waves, as follows. We first computed the residuals for wave $w$ as

$$y_{w,i} = \boldsymbol{X_i}\boldsymbol{\beta_w} + \epsilon_{w,i},$$

$$\hat{\epsilon}_{w,i} = y_{w,i} - \boldsymbol{X_i}\widehat{\boldsymbol{\beta}}_{\boldsymbol{w}} \tag{10}$$

where $y_{w,i}$ is income gamble risk tolerance in wave $w$ for individual $i$ and $\boldsymbol{X_i}$ contains the intercept and the control variables (birth year, birth year squared, birth year cubed, sex, as well as three interaction terms between sex and the three birth year variables). We then calculated the average residual across waves as $\hat{\epsilon}_i = \overline{\hat{\epsilon}_{w,i}}$. To ensure this averaged residual is not associated with any of the control variables in $\boldsymbol{X_i}$ either, we computed residual $p_i$ as

$$\hat{\epsilon}_i = \boldsymbol{X_i}\boldsymbol{\beta} + \eta_i,$$

$$p_i = \hat{\epsilon}_i - \boldsymbol{X_i}\widehat{\boldsymbol{\beta}}, \tag{11}$$

and used this residual $p_i$ is used as the phenotype for the prediction in the HRS cohort. The sample size for income gamble risk tolerance is 7,302 in the HRS cohort (**Supplementary Table 10.1**).

##### *10.5.2 Merging the STR1 and STR2 cohorts*

Since the STR1 and STR2 cohorts have been genotyped with different genotyping arrays (for details, see **Supplementary Table 2.1**), we kept them separate for the GWAS of general risk tolerance (**Supplementary Information section 2**). However, we merged the cohorts for the prediction analyses. First, we constructed and standardized the polygenic scores and calculated the 10 genetic principal components (PCs) separately in the STR1 and STR2 cohorts. We then merged the individual-level data by pooling the polygenic scores into a single variable. Two separate sets of PCs were used, imputing the value 0 for the 10 PCs of STR2 for the individuals in the STR1 cohort, and vice versa. In all the prediction analyses in the merged STR cohort, we included a dummy variable indicating whether the individual belonged to the STR1 or STR2 cohort.

### 11 Testing hypotheses about specific genes and gene sets

Earlier studies have shown that risk tolerance and related individual characteristics are moderately heritable[7,150], thus providing a motivation to investigate the biological and molecular bases of this heritability. Indeed, there is a voluminous literature that links specific biological pathways (i.e. brain regions, neuronal populations, hormones, receptors, and neurotransmitters) to decision making and behavior. Different research designs and proxies for biological pathways have been used, all with specific advantages and disadvantages (see, for example, Nave *et al.* (2015)[151] for a review of studies on the role of oxytocin on trust).

One part of this literature is candidate gene studies that have attempted to leverage insights from psychology and biology to derive hypotheses that could be tested using specific genes as proxies for biological pathways. Although the hypothesis-based approach seems intuitively reasonable, it is now widely accepted that findings from candidate gene studies on human behavior often fail to replicate. The reasons for this include very small effects of common genetic variants on human behavior, small sample sizes in candidate gene studies that led to underpowered statistical tests, publication bias, and insufficient controls for correlations between genotypes and relevant environmental conditions[15,16,18,20,152–155].

We systematically reviewed the prior literature on biological pathways that have been hypothesized to be linked to risk tolerance. Then, we used MAGMA[156] on the summary statistics from the meta-analysis of the discovery and replication GWAS of general risk tolerance ($n$ = 975,353; as usual, we applied genomic control using the intercept of the LD Score regression to these summary statistics prior to the analyses) to re-evaluate the findings of this literature.

In addition, since risk tolerance has been hypothesized to be under recent evolutionary pressure[157–159], we tested whether a set of genes previously identified as having been under evolutionary pressure in Europeans is associated with general risk tolerance.

#### 11.1 Literature review

##### *11.1.1 Search algorithm*

We conducted a comprehensive literature review on biological pathways that may influence risk tolerance. Consistent with our GWAS, we restricted our review to research involving healthy individuals. Thus, we excluded studies that focused on neuropathologies, addictions, or other disorders. However, we included articles using animal models to study risk tolerance because the investigated biological mechanisms may have similar effects on human behavior. Our search algorithm employed a relatively broad definition of risk tolerance and included psychometric as well as behavioral measures (e.g. self-reported risk tolerance, choices between monetary lotteries, or gambling tasks). Risk tolerance among animals was typically measured by observing choices between probabilistic food options or by quantifying risk-assessment behavior during exploration. Following the analysis plan for our GWAS of general risk tolerance, we considered impulsivity and novelty-seeking as conceptually different traits and excluded them from this literature review. We included all proxies of biological pathways that are tested in the literature, including candidate genes, molecules represented by specific receptors, pharmacological interventions that are used to manipulate specific pathways (e.g., nasal oxytocin spray, neurotransmitter receptor blockers, agonists, antagonists), as well as very distal proxies like the 2D:4D digit ratio.

We employed two parallel approaches to translate these criteria into search algorithms: first, we used a "bottom-up" approach and searched for studies investigating the relationship between risk taking and any biological pathway. The "bottom-up" search criterion we used was:

> *("risky behavior" OR "risk-aversion" OR "risk preference" OR "risk-taking" OR "risk-seeking") AND ("GWAS" OR "gene" OR "SNP" OR "allele" OR "genetic" OR "candidate" OR "hormones" OR "neurotransmitter" OR "neuropeptide" OR "biology" OR "biological pathway").*

We also conducted a "top-down" search which specifically cued our search criterion with biological pathways that were mentioned in the prior literature on risk tolerance. This "top-down" search criterion was:

> *("risky behavior" OR "risk-aversion" OR "risk preference" OR "risk-taking" OR "risk-seeking") AND ("catecholamines" OR "monoamines" OR "dopamine" OR "adrenaline" OR "noradrenaline" OR "norepinephrine" OR "glutamate" OR "serotonin" OR "GABA" OR "BDNF" OR "testosterone" OR "acetylcholine" OR "NMDA" OR "AMPA" OR "testosterone" OR "estradiol" OR "progesterone" OR "cortisol" OR "glucocorticoid" OR "vasopressin" OR "oxytocin" OR "ghrelin" OR "leptin").*

We used both of these algorithms in two different search engines; the ISI Web of Knowledge and Google Scholar. Both search engines yielded highly overlapping results.

We screened approximately 1,000 articles and found 132 which matched our criteria, spanning different scientific fields (from management to neuroscience), different models (e.g. human or rodent), different risk measures (from psychometric to behavioral measures), different biological pathways (e.g. monoamines, sex hormones), and different approaches (from genotyping to pharmacological manipulations). We included articles regardless of whether the reported results were statistically significant or not.

##### *11.1.2 Results of the literature review*

The results of our literature review are compiled in **Supplementary Table 11.1**. Overall, the literature on biological mechanisms for risk tolerance is relatively recent: most studies were published after 2003. However, earlier work on related concepts that were excluded by our selection criteria (such as novelty seeking) exists.

Three main biological mechanisms comprising five biological pathways have been tested by this literature. The first of these is the relationship between risk tolerance and sex hormones, especially testosterone and estrogens. Testosterone and estrogens are the two primary steroid sex hormones in humans. They are principally (but not exclusively) synthesized in the ovaries (estrogen) and testicles (testosterone), and are responsible for the development and regulation of the male (testosterone) and female (estrogen) reproductive system and secondary sex characteristics. The link between those hormones and risk taking is generally motivated by (and discussed as corroborating of) the observed behavioral and attitudinal differences between males and females. A large part of the literature testing the relationship between risk tolerance and sex hormones uses the distal 2D:4D digit ratio to approximate prenatal testosterone levels. Some studies use measures of sex hormones that are derived from saliva or blood samples.

The second frequently examined biological mechanism is the relationship with the neurotransmitters dopamine and serotonin. Those two monoamines have been repeatedly implicated in behavior and decision-making: dopamine plays a major role in reward-motivated behavior and in addiction, whereas serotonin is implicated in mood. Studies testing the associations between these monoamine pathways and risk tolerance typically test genetic variants implicated in the synthesis or the signaling pathways of these neurotransmitters (e.g., *DRD4*, *SERT* or *5-HTTLPR*). Alternatively, pharmacological manipulations are used including blockers, agonists, or antagonists of specific dopamine or serotonin receptors.

The third main biological pathway tested by the literature involves cortisol. Cortisol is another steroid hormone belonging to the glucocorticoid class, and is notably produced in response to stress. Cortisol triggers gluconeogenesis (the formation of glucose), and activates anti-stress and anti-inflammatory pathways. Studies testing the associations between the cortisol pathways and risk tolerance typically use a biochemical quantification of cortisol level in salivary or blood samples as an independent measure.

In sum, our literature review highlighted five main biological mechanisms potentially underpinning the risk-tolerance phenotype: the testosterone, estrogen, dopamine, serotonin and cortisol pathways.

#### 11.2 Gene analysis and competitive gene-set analyses with MAGMA

We used MAGMA[156] together with the summary statistics from the meta-analysis of the discovery and replication GWAS of self-reported general risk tolerance to re-evaluate the findings of this literature.

We conducted several types of analyses based on these results. First, for all five mechanisms we constructed gene sets, based on external databases of biological function, and tested these gene sets for association with risk tolerance. Second, for two of those pathways (dopamine and serotonin), there are a significant number of candidate gene studies in humans. These candidate gene studies most commonly test a specific subset of genes and genetic variants (SNPs or structural variants) within the dopamine and serotonin pathways. We therefore built a gene set from 15 of the 17 most commonly tested genes of those two pathways (**Supplementary Table 11.4**) in order to re-evaluate the associations of those candidate genes with risk tolerance. Third, we tested specific SNPs in these 15 genes that have been tested in the prior candidate gene literature. Lastly, we tested whether a set of genes that has been identified as having been under selection among Europeans was enriched for association with general risk tolerance.

MAGMA can perform two types of analyses: a gene analysis and a competitive gene-set analysis. For a given gene, a MAGMA gene analysis tests the joint association of all the SNPs in the gene with the phenotype. This aggregation of the SNPs reduces the number of tests compared to a SNP-based analysis, which results in an increase in statistical power and may lead to the detection of joint effects of multiple weaker SNP associations that would otherwise be missed. A disadvantage of gene analysis is that it is uninformative about the direction of the effect, as it simultaneously tests multiple SNPs that can have opposite effects within a gene.

For a given gene set, a MAGMA competitive gene-set analysis tests whether the genes in the gene set are more strongly associated with general risk tolerance than the other genes in the genome. Gene sets are groups of genes that share certain characteristics (e.g., biological or functional characteristics). Like a gene analysis, a competitive gene-set analysis has increased statistical power relative to a SNP-based analysis, as it involves fewer tests. Moreover, a gene-set analysis allows tests of specific biological hypotheses by defining appropriate gene-sets, and may provide direct insights into the underlying biological pathways or cellular functions of the phenotype. MAGMA uses a regression framework that compares the mean association of the genes in the gene set with the mean association of all other genes in the genome (equivalent to a one-sided two-sample *t*-test). To account for possible LD between genes, MAGMA uses a gene correlation matrix, which was based on the European ancestry samples from the 1000 Genomes project phase 1[77]. Additional covariates are included in the regression to correct for any confounding effects of gene size and gene density (gene size, log gene size, gene density, log gene density).

###### *11.2.1 Competitive gene-set analysis with MAGMA: Testing gene sets related to dopamine, serotonin, testosterone, estrogen, and glucocorticoids*

As mentioned in **Supplementary Information section 11.1**, not all of the biological pathways that were studied in the previous literature on risk tolerance were studied using candidate genes as proxies for these pathways. Furthermore, the candidate genes (e.g. *DRD4*) that were studied are not the only genes that are involved in the respective biological pathways (e.g. dopamine). Thus, to begin with, we constructed and tested gene sets that represent the five major biological pathways that were previously studied in the context of risk tolerance. (We note that we also performed e*x post* MAGMA gene set analyses for GABA and glutamate neurotransmitters. We describe these analyses in **Supplementary Information section 12.5**.)

We selected all gene sets related to the five major biological pathways (dopamine, serotonin, testosterone, estrogen and cortisol) from the Molecular Signature Database (MSigDB, v5.1)[160], and from the gene sets compiled by Hawrylycz *et al.* (*Nature Neuroscience* 2015)[161]. There were 38 gene sets related to the five pathways, from which we removed four duplicate gene sets. The remaining 34 gene sets had much overlap within each biological pathway, and we therefore merged all the gene sets belonging to a given pathway, resulting in five gene sets corresponding to each of the five pathways (dopamine, serotonin, testosterone, estrogen and glucocorticoids (cortisol)). We report both the initial and the merged gene sets in **Supplementary Table 11.2**.

We conducted a competitive gene-set analysis with MAGMA to test each of these five gene sets for association with risk tolerance. We applied a resampling-based *P* value adjustment[162] using 10,000 permutations to correct for multiple testing (as we conducted a test for each of the five gene sets).

None of these five gene sets showed a significant association with general risk tolerance after correction for multiple testing (the lowest corrected *P* value was 0.27, **Supplementary Table 11.3**).

##### *11.2.2 Gene analysis and competitive gene-set analysis with MAGMA: Testing candidate genes from the prior literature*

We used MAGMA to conduct a gene analysis to test the candidate genes identified in our literature review for association with general risk tolerance, employing all of the SNPs in our GWAS summary statistics within the physical location of the genes being tested. 15 of the 17 candidate genes that we identified in the literature review were autosomal, and were thus eligible for testing (since our GWAS of general risk tolerance was restricted to autosomal SNPs). In addition to the gene analysis, we conducted a competitive gene-set analysis with MAGMA to test whether the genes in the gene set comprising these 15 genes are more strongly associated with general risk tolerance than the other genes in the genome.

From the gene analysis, none of the genes showed a significant association with general risk tolerance after Bonferroni correction to correct for multiple testing (the 15 genes are not in LD; **Supplementary Table 11.4**). From the competitive gene-set analysis, the set of 15 candidate genes was also not significantly associated with risk tolerance ($P = 0.55$, **Supplementary Table 11.4**).

Our MAGMA competitive gene-set analysis of the cortisol, dopamine, serotonin, estrogen, and testosterone gene sets has some likely limitations. First, the statistical power of MAGMA competitive gene-set analyses barely increases with increasing sample sizes[163]. Second, important effects of numerous single SNPs in a gene or gene set may be overshadowed by the high average *P* values of other SNPs in the gene or gene set. Third, only specific pathways within the tested gene sets might be relevant, and merging many pathways into major cortisol, dopamine, serotonin, estrogen, and testosterone gene sets decreased our statistical power to discover specific pathways. In **Supplementary Information section 12.7.4**, we further discuss these limitations in the context of our *ex post* MAGMA competitive gene-set analysis of four glutamate and GABA gene sets; that analysis also returned null results, despite the fact that our Gene Network and DEPICT analyses both point to a role for glutamate and GABA neurotransmitters.

Importantly, however—and unlike for glutamate and GABA—none of the bioinformatics analyses we report in **Supplementary Information section 12** point to the cortisol, dopamine, serotonin, estrogen, and testosterone pathways or related genes either. Further, as reported above none of the 15 commonly-tested candidate genes were significant in our MAGMA gene analysis after Bonferroni correction for 15 tests; by contrast, our MAGMA gene analysis of ~18,000 genes identified several glutamate and GABA genes that were significant after Bonferroni correction for ~18,000 tests (**Supplementary Information section 12.2**). Moreover, as we discuss in **Supplementary Information section 11.2.4** just below, none of the SNPs tagging the 15 most commonly-tested autosomal candidate genes within the dopamine and serotonin pathways have non-negligible effects on general risk tolerance. Therefore, while our MAGMA competitive gene-set analysis may suffer from some limitations, other analyses we conducted also fail to point to a role for the cortisol, dopamine, serotonin, estrogen, and testosterone pathways, and instead point to a role for glutamate and GABA neurotransmitters.

##### *11.2.3 Replication of specific SNPs tested in the prior candidate gene literature*

As a complement to the MAGMA gene-based tests, we also tested the individual SNPs that have been tested for association with risk tolerance in the previous literature and that are located within the 15 autosomal genes tested with MAGMA. We used the summary statistics from the meta-analysis of the discovery and replication GWAS of general risk tolerance for these tests. Out of the 15 autosomal genes, three have been tested in the previous literature for association with alleles of an underlying structural variant, and we treat these separately below, where we looked up a total of 18 SNPs available in our GWAS summary statistics for these three genes. The remaining 12 autosomal genes contained 75 SNPs that were individually tested in the previous literature, of which 74 were available in our meta-analysis of the discovery and replication GWAS of general risk tolerance. Thus, our GWAS summary statistics contain the majority of candidate SNPs within these genes that were tested in the previous literature, and we performed Bonferroni-correction for testing 92 candidate SNPs.

Out of the 74 previously tested SNPs within the 12 autosomal genes, two replicated with Bonferroni-corrected $P$ values less than 0.05. The SNPs were rs36339 (in the gene *SLC1802*; Bonferroni-corrected $P$ = 0.004), and rs174699 (in the gene *COMT*; Bonferroni-corrected $P$ = 0.034). The $R^2$'s of rs36339 and rs174699 in the summary statistics of our discovery GWAS of general risk tolerance are 0.00185% and 0.00141%, respectively. By comparison, the smallest $R^2$ among the lead SNPs from our discovery GWAS of general risk tolerance is almost twice as large, at 0.0032% (rs1935571). The effects of rs36339 and rs174699 are considerably smaller than what could be found at the genome-wide significance level with our current discovery sample of 939,908 individuals, and they are several orders of magnitude smaller than what candidate gene studies in the existing literature—with typical sample sizes rarely exceeding a few hundred to a few thousand individuals—could have found. The sample sizes required to have 80% statistical power to detect the effects of rs36339 and rs174699 at the 5% level of significance (without Bonferroni correction) would be 424,260 and 556,654, respectively; the corresponding sample sizes at the genome-wide significance level ($5 \times 10^{-8}$) would be 2,140,573 and 2,808,560, respectively. We therefore found that 10 of the 12 hypothesized autosomal candidate genes harbor no candidate SNPs that replicate, while two (*SLC1802* and *COMT*) each contain a SNP that is suggestively associated with general risk tolerance (with a $P$ value less than 0.05 after Bonferroni correction for 92 tests) but whose effect is very small.

In the previous literature, three of the 15 autosomal candidate genes (*DRD4, DRD5* and *SLC6A4* (the latter is also commonly referred to as *SERT* or *5-HTT*)) have mainly been tested for association with behavioral traits using alleles of structural variants located within these genes. For example, risk tolerance has previously been tested for association with a variable number tandem repeat in the *DRD4* gene referred to as the 7R polymorphism[11]. Our GWAS summary statistics include nine SNPs within the *DRD4* gene. Although we do not know the exact LD between these nine SNPs and the 7R polymorphism, it is very likely that at least some of them tag the 7R polymorphism very well because the *DRD4* gene spans only 3,401 base pairs on chromosome 11, and variants in such close physical proximity tend to be in high LD. The lowest Bonferroni-corrected $P$ value of a SNP in the *DRD4* gene is 0.47 (rs201554946). The gene *DRD5* has mainly been tested for association with a microsatellite that can take many different alleles[164], and no SNPs within *DRD5* were found in our literature review to have been tested directly for association with risk tolerance. *DRD5* spans 2,375 base pairs on chromosome 4, in which we have seven SNPs in our GWAS summary statistics, and none of the SNPs have a Bonferroni-corrected $P$ value less than 1. The gene *SLC6A4* has mainly been tested for association with risk tolerance via a degenerate repeat polymorphism in a region of the gene called *5-HTTPLR*[165], and alleles of this repeat polymorphism are tagged by the SNPs rs2129785, rs11867581, and rs25531[166]. The SNPs rs2129785 and rs11867581 are available in our GWAS results and their Bonferroni-corrected $P$ values are both 1. Based on these results we cannot reject the null hypothesis that none of these three genes is associated with risk tolerance.

**Extended Data Fig. 3.4b** and **Fig. 1c** display local Manhattan plots of the loci around the 15 most commonly tested candidate genes in the prior literature on the genetics of risk tolerance. Each locus comprises all SNPs within 500 kb of the gene's borders that are in LD ($r^2 > 0.1$) with a SNP in the gene. As can be seen in **Extended Data Fig. 3.4b**, only one of the 15 loci (around the gene *DRD2*) contains lead SNPs for any of our seven GWAS, and these SNPs are only genome-wide significant in our GWAS of drinks per week. One SNP in the *SLC6A4 (SERT)* locus is genome-wide significant in our GWAS of adventurousness. However, this SNP (rs112739039) is not a lead SNP in our GWAS of adventurousness because it is located in the locus of another SNP (rs6505239; see **Supplementary Table 3.2**). The nearest gene to rs112739039, *RAB11FIP4,* is located ~1Mb upstream of *SLC6A4*. In our meta-analysis of general risk tolerance, this *SLC6A4* SNP is not genome-wide significant and is included in the locus defined by rs11080149 and nearest gene *OMG*, which is also located ~1Mb upstream of *SLC6A4* (see **Supplementary Table 3.1**).

The local Manhattan plots of the loci around the 15 candidate genes (**Extended Data Fig. 3.4b** and **Fig. 1c**) can be compared to those of the five main long-range LD regions and candidate inversions described in **Supplementary Information section 3.2** (**Extended Data Fig. 3.4**, **a** and **c**, and **Fig. 1**, **a** and **b**). These five regions contain numerous lead SNPs for most or all of our seven main GWAS.

In summary, no candidate genes that have been tested with SNPs in the previous literature, nor the candidate genes tested with alleles of structural variation, contain genome-wide significant SNPs in our discovery GWAS, while two candidate genes each contain a SNP with a suggestive association (i.e., with a Bonferroni-corrected $P$ value smaller than 0.05, after Bonferroni correction for 93 tests). However, a sample size of more than 2 million individuals would be required to have 80% power to replicate these two SNPs at the genome-wide significance level, and a sample size of more than 400,000 would be required to replicate these SNPs at the 5% level of significance. We therefore conclude that these two SNPs are relatively unimportant in terms of their effects on general risk tolerance, and that previous candidate gene studies testing these SNPs for association with behavioral phenotypes were severely underpowered to detect their very small effects.

##### *11.2.4 Competitive gene-set analysis with MAGMA: Testing genes under selection among Europeans*

A number of theorists have posited that attitudes toward risk have been under evolutionary pressure[157–159]. In particular, if decisions involving risk have to be made frequently and if decision mistakes have a severe impact on fitness, theoretical models suggest that selective pressure will favor risk attitudes that maximize fitness in a given environment[158]. Consequently, changes in the environment can induce selective pressure for different levels of risk tolerance. It is obvious that the environment in which people live and make choices has changed dramatically over the past few millennia, with many fitness-related risk factors declining in importance, and new ones arising (e.g. due to urbanization, technological progress, and increasing populations). Hence, it is conceivable that general risk tolerance has been under selective pressure in our recent evolutionary past.

To test if risk tolerance was subject to recent selective pressure among people of European descent, we constructed a gene-set based on 17 genes that were highlighted as having been under recent selective pressure among Europeans by the 1000 Genomes phase 3 analyses[167] (see **Supplementary Table 11.5**). We performed a competitive gene-set analysis with MAGMA and with the summary statistics from the meta-analysis of our discovery and replication GWAS of general risk tolerance, to test whether the genes in this gene-set are more strongly associated with general risk tolerance than other genes.

The gene-set based on the genes that have been highlighted as having been under recent selective pressure among Europeans showed no association with general risk tolerance ($P$ value = 0.90, **Supplementary Table 11.5**). Thus, this test provides no evidence suggesting that risk tolerance has been subject to recent selective pressure. However, this absence of evidence is not evidence of absence: our test may have failed to detect evolutionary pressure for several reasons, including limited statistical power. Further, we only tested if a set of genes that have been highlighted as having been under evolutionary pressure is associated with general risk tolerance; future research could also test whether genes that are associated with risk tolerance have been under evolutionary pressure.

#### 11.3 Discussion

In summary, we find no evidence of enrichment for the main pathways and genes that had previously been hypothesized to relate to risk tolerance. We also note that none of the bioinformatics analyses we report in **Supplementary Information section 12** point to these pathways or genes either (we note, however, that some brain regions identified in analyses we report below are areas where dopamine and serotonin play important roles.) By contrast, some those bioinformatics analysis point to a role for the glutamate and GABA neurotransmitters.

Our null replication results are consistent with the poor replication record that candidate gene studies in social-science genomics have typically had[15,16,18,20,152–155]. Given that the sample size of our GWAS is several orders of magnitudes larger than any previous candidate gene study on risk tolerance, our results put a credible, tight upper bound on the effect sizes of the genes that were tested, and allow us to rule out the possibility that these genes have particularly large effect sizes compared to typical genetic variants.

However, our results do not imply that the main biological pathways we identified in our literature review are irrelevant for risk tolerance, as variations further downstream in these pathways (e.g. damaged glands, pharmacological interventions) may have much stronger effects on risk tolerance than the genes that were tested.

### 12 Biological annotation

This section reports biological annotation of our GWAS results. Throughout, we focus on our primary phenotype, self-reported general risk tolerance, because it has by far the largest GWAS sample size and because the other main phenotypes analyzed in this paper are genetically correlated with it. The goal of these analyses is to gain biological insight (1) into the genome-wide biological correlates of the general-risk-tolerance phenotype (with a suite of bioinformatic analyses that use GWAS summary statistics for large sets of SNPs across the genome), and (2) about the lead loci from our discovery GWAS of general risk tolerance (by looking up the functional status of the SNPs in these loci). For all analyses in this section, we use the summary statistics from the meta-analysis of the discovery and replication GWAS of general risk tolerance.

In **Supplementary Information section 12.1**, we use partitioned LD Score regression[74] to test for polygenic enrichment in specific genomic regions, such as gene transcription start sites, evolutionarily conserved regions, and sections of the genome epigenetically modified in certain tissues.

In **Supplementary Information section 12.2**, we use MAGMA[156] to conduct hypothesis-free analyses to identify specific genes that are associated with general risk tolerance. We also use a co-expression database to gain insight into the functions of the significant MAGMA genes.

In **Supplementary Information section 12.3**, we perform transcriptome-wide analysis with Summary-based Mendelian Randomization (SMR), which leverages genome-wide data to discover genes whose expression significantly associates with general risk tolerance. In addition, the "HEIDI" test feature of SMR is able to discard expression quantitative trait loci (eQTL) associations that are caused merely by linkage, thereby discarding genes that are spuriously associated with general risk tolerance.

In **Supplementary Information section 12.4**, we use DEPICT[168] to prioritize tissues, gene sets, and genes that are implicated by our GWAS results. Like MAGMA, DEPICT is a gene-based tool. However, in contrast to MAGMA, DEPICT uses reconstituted gene sets that are based on co-expression patterns to prioritize genes, gene sets, and pathways.

In **Supplementary Information section 12.5**, to follow up on the results from other biological annotation analyses, we conduct an *ex post* MAGMA gene set analysis[156] to test for enrichment of GABA and glutamate pathways in general risk tolerance.

In **Supplementary Information section 12.6**, we report a number of analyses that annotate our general-risk-tolerance lead loci, to gain insights into the biological systems they may affect. First, we check whether our lead loci contain protein-altering variants (i.e., genetic variants which result in a structural difference in the protein structure encoded by a gene, collectively known as 'nonsynonymous' variants). Second, we look up the lead loci in an expression quantitative trait loci (eQTL) database that contains associations between genetic variants and gene expression levels.

Finally, in **Supplementary Information section 12.7**, we highlight the most important results of these analyses and summarize the conclusions we derive from them.

#### 12.1 Functional partitioning of heritability with stratified LD Score regression

##### *12.1.1 Background and methods*

In this section, we discuss the results of our stratified LD Score regression analyses[74]. (In addition to ref.[74], the Supplementary Information of Okbay *et al.*[18] also contains a detailed description of the methodology.) With these analyses, we break down ("partition") the SNP-based heritability of general risk tolerance across SNPs with various functional genomic annotations. We used the GWAS summary statistics from the meta-analysis of the discovery and replication GWAS of general risk tolerance (after applying genomic control using the intercept of the LD Score regression, as usual) for these analyses.

Stratified LD Score regression is based on the relationship

(12)
$$E\left[\chi_j^2\right] = N\sum_{c=1}^{C} \tau_c \ell(j,c) + Na + 1,$$

where $\chi_j^2 = N\hat{\beta}_j^2$ is the GWAS chi-square statistic for SNP $j$, $N$ is the GWAS sample size, $c$ indexes the functional categories (which do not have to be disjoint), $\ell(j,c)$ is the stratified LD Score of SNP $j$ with respect to functional category $c$, $\tau_c$ is the average contribution to heritability of a SNP due to its membership in category $c$, and $a$ is a term that measures the contribution of confounding biases such as cryptic relatedness and population stratification.

Finucane *et al.*[74] present derivations of this equation and show how estimates of $\tau_c$ that result from estimating the implied regression can be used to obtain estimates of the heritability ascribable to the various functional categories. Enrichment is then defined as the fraction of the total heritability captured by the functional annotation category divided by the fraction of SNPs in that category.

To partition the SNP-based heritability of risk tolerance using the results of our GWAS meta-analysis, we followed the procedure described by Finucane *et al.*[74] and applied in two recent papers by Okbay *et al.*[16,18]. That is, we used the stratified LD scores calculated from the European-ancestry samples in the 1000 Genomes Project (1000G) (accessed on March 14, 2016), but in the regressions themselves included only the chi-square statistics of the ~1 million HapMap3 SNPs with minor allele frequency (MAF) > 0.05. This decision was motivated by the facts that HapMap3 SNPs are a comprehensive set of common genetic markers that can be imputed with high accuracy across different groups and genotyping platforms[143,169]; that the LD scores of SNPs with low MAFs can introduce too much statistical noise when used in the analysis; and that the per-SNP heritability may substantially differ for common and rare SNPs.

We performed two distinct analyses. We first estimated the stratified LD Score regression for the functional genomic regions of the "baseline" model. The functional categories in the baseline model consist of one category containing all SNPs, 24 categories corresponding to 24 main functional annotations of interest (which include, for example, evolutionarily conserved regions in mammals and regions epigenetically regulated to be accessible to gene transcription), categories corresponding to 500 bp windows around regions belonging to each of these 24 annotations (to correct for spurious associations driven by variants located near the physical borders of the functional genomic regions), and categories corresponding to 100-bp windows around ChIP-seq peaks (regions that are DNase hypersensitive, also known as "open chromatin," or associated with the histone marks H3K4me1, H3K4me3, H3K9ac, or H3K27ac). The full baseline model contains 53 predictors (including the predictor for the category containing all SNPs).

Second, to gain tissue-level resolution, we used the tissue-level annotations provided by Finucane *et al.* These consist of 220 cell-type-specific annotations for four histone marks that are subsequently grouped into 10 broad tissue type annotations (Adrenal/Pancreas, Central Nervous System, Cardiovascular, Connective/Bone, Gastrointestinal, Immune/Hematopoietic, Kidney, Liver, Skeletal Muscle, and Other). Histone marks are posttranslational modifications of histones that alter their interaction with the DNA wound around them. For example, the acetylation of lysine (an amino acid present in the histone protein) may facilitate gene expression by reducing the electrical attraction between DNA and the histone, making the DNA strand more receptive for transcription (and thus, gene expression). While certain histone marks are associated with increased DNA transcription, other histone marks' effects on DNA transcription depend on the combination of the chemical form and the location of the modification. For instance, mono-methylation of histone H3K9 (denoted H3K9me1) is associated with activation of gene expression, while tri-methylation of H3K9 (denoted H3K9me3) is associated with repression of gene expression[170]. The association between SNPs and histone modifications differs per tissue and developmental stage, and has been mapped extensively by the RoadMap Epigenomics project[171]. For instance, SNPs important for eye function are likely to be "tightly bound" to histone proteins in the liver, where they do not need to be expressed, but "loosely" bound to histones in the eye, where their expression is crucial. Here, we used LDSC partitioning of heritability to test if risk tolerance SNPs are enriched in (or near) tissue-specific histone marks, which would suggest that those tissues might be of biological importance for general risk tolerance.

The functional annotation categories we used here are associated with the histone marks H3K4me1, H3K4me3, H3K9ac and H3K27ac, which are all known to increase transcription rate[172]. However, we note that the probability of any gene being transcribed and thus expressed further depends on the presence of other epigenetic modifications, such as epigenetic modifications of the DNA sequence itself (for example, methylation of CpG-islands). This means that the four histone modifications we study here do not deterministically increase gene expression rates, but rather do so in a probabilistic manner.

We added each of the 10 tissue annotations to the baseline model (resulting in 10 separate regressions, each with 54 predictors), and assessed the magnitude and statistical significance of the observed enrichment. To benchmark these tissue-level results, we compared them to the corresponding estimates we obtained using the summary statistics of the most recent GWAS of height[83] (http://www.broadinstitute.org/collaboration/giant/index.php).

To correct for multiple hypothesis testing, we applied a Bonferroni correction for 52 two-sided tests in the baseline model (i.e., for 52 annotations[ddd]), and for 10 two-sided tests in the tissue type models (i.e., for 10 tissue types).

[ddd] The baseline model has 53 predictors, but we do not adjust for the predictor for the category containing all SNPs.

##### 12.1.2 Results: The "baseline" model

The results for the baseline model are shown in **Supplementary Table 12.1**. The baseline annotations "Conserved" and "H3K9ac peaks" are the most enriched, with enrichment estimates that remain significant after Bonferroni correction for 52 tests. The "Conserved" category shows the strongest enrichment (∼15.8-fold) and accounts for 41% (*SE* = 5.0%) of the heritability of general risk tolerance; the "H3K9ac peaks" category shows a 5.5-fold enrichment and accounts for 21% (*SE* = 3.7%) of the heritability of general risk tolerance.

##### 12.1.3 Results: Tissue types

The results of the tissue-level analyses are reported in **Extended Data Fig. 12.3a**, and **Supplementary Table 12.2**. For general risk tolerance, the enrichment of Central Nervous System is strongest (∼2.94-fold, $P = 5.90 \times 10^{-20}$), followed by Adrenal/Pancreas (~2.47-fold, $P = 1.41 \times 10^{-7}$). The enrichment estimates for Immune/Hematopoietic, Cardiovascular, Liver, and Skeletal Muscle are also significant, and also survive Bonferroni correction. Our estimates imply that SNPs bearing the Central Nervous System and Adrenal/Pancreas annotations account for 43.8% (*SE* = 2.8%) and 23.1% (*SE* = 2.5%) of the heritability of general risk tolerance, respectively; the corresponding figure for Immune/Hematopoietic is 37.6% (*SE* = 3.6%). These estimates are all highly significant and survive Bonferroni correction for ten tests.

However, the enrichment statistics are potentially misleading because of possible confounding. For example, many SNPs bear multiple annotations. It is thus of interest to examine the $\tau_c$ estimates of each tissue (i.e., the coefficients from the stratified LD Score regression). The $\tau_c$ for a given tissue type and phenotype corresponds to the effect of a one-unit increase in a SNP's stratified tissue-specific LD score on the expected square of the SNP's GWAS estimate from the phenotype's GWAS (where the SNP genotype has been standardized), after controlling for the annotations from the baseline model. It is also an estimate of the expected increase in the phenotypic variance accounted for by a SNP due to the SNP's being in the given tissue category, controlling for the annotations from the baseline model. SNPs that bear a tissue annotation with a large and positive $\tau_c$ will tend to account for a larger share of a phenotype's heritability.

**Extended Data Fig. 12.3a** shows, for general risk tolerance and height, the ratio of the $\tau_c$ estimates over the LD Score estimates of phenotypic heritability[eee,fff]. For each phenotype, we normalized the $\tau_c$'s and their standard errors by the LD Score heritability[ggg] to increase comparability across phenotypes. Here, only Central Nervous System ($z = 8.81$, $P = 6.04 \times 10^{-19}$) and Immune/Hematopoietic ($z = 3.19$, $P = 1.42 \times 10^{-3}$) have positive coefficients surviving Bonferroni correction (note that these analyses excluded SNPs in the MHC region on chromosome 6). We also verified that the coefficient for Immune/Hematopoietic remained positive and highly significant ($z = 4.71$, $P = 2.47 \times 10^{-6}$) even after controlling for Central Nervous System (in addition to the baseline annotations) in the LD Score partitioned regression.

---

[eee] Based on the LD score framework, $h^2_{LDSC,y} = \sum_c M_c \tau_{c,y}$, where $y$ denotes the phenotype and $M_c$ is the number of SNPs with annotation $c$ among the SNPs used to calculate the LD scores. Thus, normalizing the $\tau_c$'s by $h^2_{LDSC,y}$ is equivalent to normalizing the $\tau_c$'s by a weighted sum of the $\tau_c$'s, where the weights are given by the number of SNPs with the different annotations.

[fff] The confidence intervals were obtained with the delta method, assuming zero covariance between the $\tau_c$'s and the phenotypes' LD score heritabilities.

[ggg] Each phenotype's LD score heritability was obtained from the phenotype's LD score regression [42].

Thus, the positive coefficient for Immune/Hematopoietic is not due to possible overlap between the Immune/Hematopoietic and Central Nervous System annotations.

In these respects, general risk tolerance differs notably from a physical trait such as height, for which Connective/Bone has the largest positive *z*-score ($z$ = 6.08, $P = 1.20 \times 10^{-9}$) and Central Nervous System is the only tissue-level annotation with a *negative* coefficient[74,hhh]. Finally, we note that the coefficients for Cardiovascular, Gastrointestinal, Liver, and Skeletal Muscle (which were significantly enriched) have *negative* $\tau_c$ coefficients for general risk tolerance. This means that the GWAS signal is actually negatively enriched for histone marks in these tissues, after taking the baseline annotations into account.

##### 12.1.4 Discussion

The results of our baseline annotation model are similar to those reported by Finucane *et al.*[74] for a set of nine phenotypes and to those reported by Okbay *et al.*[16,18] for educational attainment, subjective well-being, depressive symptoms, and neuroticism. That is, evolutionary conserved regions (i.e. conserved in mammals[173]) bear the most significant enrichment, followed by annotations relating to modification of gene expression. This is in line with a large body of literature that suggests that complex traits are likely to mainly be affected by genetic variants involved in the modification of gene expression, rather than by direct coding variants[174]. In particular, the significant enrichment of "H3K9ac histone marks" points to the involvement of active gene promoters in general risk tolerance[175].

The evolutionarily conserved category denotes regions of the genome that have been conserved in mammals throughout evolution. The evolutionary conservation of a genomic region can be studied through assessment of the "mutation rate" of such a region. That is, some regions of the genome accumulate base-pair substitutions more slowly than predicted by a model of selective neutrality[173], which implies that mutations in such regions tend to have deleterious effects that decrease evolutionary fitness, and are therefore subject to natural selection.

As for the tissue enrichments, we obtained significant $\tau_c$ estimates for Central Nervous System and Immune/Hematopoietic cell types. As explained above, this implies that risk-tolerance associated SNPs that are in or near histone marks in these tissues contribute relatively more to the per-SNP heritability of risk tolerance than most other SNPs. Because risk tolerance is a psychological trait, central nervous system enrichment is not surprising. More interesting is the current lack of significance of the $\tau_c$ estimates for Adrenal/Pancreas. This suggests the highly significant enrichment estimate for Adrenal/Pancreas is due to overlap with SNPs bearing other annotations rather than to the partial effect of bearing the Adrenal/Pancreas annotation, and may possibly question the role for the stress-response system in risk tolerance[176–178]. We note, however, that the lack of significance of the $\tau_c$ estimates for Adrenal/Pancreas could also be due to lack of statistical power or to imprecisions or imperfections in the SNP annotations. The stress-response system is driven by the hormones cortisol and (nor)epinephrine (also known by the name (nor)adrenaline), which are all produced in the adrenals.

---

[hhh] We caution that a detailed quantitative comparison of the results for the height and the general-risk-tolerance GWAS might be misleading, because the sample size of the height GWAS is smaller than that of the risk-tolerance GWAS, and because the heritability of height is substantially higher than the heritability of general risk tolerance. Nonetheless, our results suggest that different tissue types tend to be relatively more enriched for the two phenotypes.

To the best of our knowledge, we are the first to find significant LDSC tissue enrichment of the immune system in a psychological or neuropsychiatric trait (although enrichment of *specific* immune pathways and/or immune tissue enhancer regions has been implicated in schizophrenia[69,179], bipolar disorder[180] and major depression[181]). We note that involvement of the immune system is not corroborated by our other analyses (aside from the large number of genes significant in the MHC region across out MAGMA, SMR and lookup analyses), and it is unclear how to interpret this finding.

#### 12.2 Identifying genes associated with general risk tolerance with MAGMA

To identify genes that are significantly associated with general risk tolerance, we conducted a gene analysis with MAGMA[156] (defined in **Supplementary Information section 11.2**) for each of ~18,000 genes, in a hypothesis-free manner. We then used the Gene Network[182] co-expression database to gain insight into the functions of the significant MAGMA genes.

##### *12.2.1 Gene analysis with MAGMA: Testing ~18,000 protein-coding genes*

The gene-based analysis was performed with MAGMA[156] using the summary statistics from our meta-analysis of the discovery and replication GWAS of general risk tolerance (after applying genomic control using the intercept of the LD Score regression).

All the SNPs from these summary statistics were annotated to genes based on NCBI 37.3.13 gene definitions. Only SNPs located between the transcription start and stop sites of a gene were annotated to that gene. A per-gene test statistic is calculated as the mean of the GWAS $-\log_{10}(P)$ values for all the SNPs between the transcription start and stop sites of a gene; MAGMA then calculates a $P$ value for the resulting gene test statistic, using a procedure that takes into account the non-independence of the SNPs within the gene due to LD[183]. We used our main reference panel (described in **Supplementary Information section 2.4**) as reference data to estimate LD. Bonferroni correction was applied to account for multiple testing, counting each gene as an independent test.

The SNP-to-gene annotation yielded 18,224 protein-coding genes containing at least one SNP present in the current GWAS. After Bonferroni correction for 18,224 tests, 285 genes showed a significant association with general risk tolerance (**Supplementary Table 12.3**). We will henceforth refer to these as the "MAGMA genes." The top ten MAGMA genes are: *CADM2* (chr. 3: 85 Mb, $P = 1.08 \times 10^{-50}$; all the $P$ values reported in this paragraph are Bonferroni-corrected $P$ values), *MSRA* (chr. 8: 9 Mb, $P = 1.61 \times 10^{-28}$), *XKR6* (chr. 8: 10 Mb, $P = 3.54 \times 10^{-23}$), *FOXP2* (chr.7: 113 Mb, $P = 9.94 \times 10^{-19}$), *MFHAS1* (chr. 8: 8 Mb, $P = 2.76 \times 10^{-18}$), *RP1L1* (chr. 8: 10 Mb, $P = 2.04 \times 10^{-15}$), *FOXP1* (chr. 3: 71 Mb, $P = 2.23 \times 10^{-13}$), *RBFOX1* (chr. 16: 5 Mb, $P = 1.99 \times 10^{-12}$), *ARNTL* (chr. 11: 13 Mb, $P = 2.27 \times 10^{-11}$), and *ERI1* (chr. 8: 9 Mb, $P = 7.49 \times 10^{-11}$).

The results show that several long-range LD or inversion regions of the genome contain a disproportionate number of MAGMA genes (which is not surprising, since MAGMA does not correct for LD between genes in its gene-based test). For instance, the Human Leukocyte Antigen (HLA) region (and its surrounding area) carries a relatively large number of significant genes: there are 29 significant genes in the chr.6: 25-32 Mb region, of which 11 are zinc finger genes, six are histone protein genes, and three are olfactory function genes (none of the *HLA*-

genes themselves are significant, however). The large candidate inversion region on chromosome 8 (chr.8: 8 to 12 Mb, previously associated with neuroticism and depressive symptoms[18]) and its surrounding area carry 21 significant genes, 13 of which are among the top 30 most significant MAGMA genes, and of which *MSRA* is the most significant by far (Bonferroni-corrected $P = 1.61 \times 10^{-28}$). A long-range LD region on chromosome five (chr.5: 135-138 Mb) contains 10 significant genes, of which *ETF1* is the most significant by far (Bonferroni-corrected $P = 7.46 \times 10^{-7}$). Finally, we note that several of these genes (*MSRA, ERI1, XKR6, and ETF1*) were also significant in the SMR analyses, although only in the analysis of blood *cis*-eQTLs, and not in the analysis with GTEx data on *cis*-eQTLs from brain regions (lack of replication with the brain eQTL data may be because these data are based on relatively small samples and statistical power was consequently limited).

##### *12.2.2 Using co-expression to predict gene function*

We used a co-expression database called Gene Network[182] (accessed 6 September 2017) to gain further insight into the functions of the MAGMA genes. The advantage of Gene Network lies in its ability to assign functions to a gene, even if that gene has not been annotated with a function in an actual empirical experiment (as described below). This is an important advantage because many human genes have not yet been studied in depth and hence have poorly understood functions. Therefore, looking up only the genes with empirically established functions in databases is in most cases not fully informative.

The gene functions used by Gene Network are derived from expert-curated sets of genes known as “gene sets” or “pathways.” The experts involved in the curation of such gene sets continuously review the empirical literature of a gene, and assign a gene its functions based on empirical results from laboratory experiments. Gene Network then assigns genes “membership” to these expert-curated gene sets by establishing “guilt-by-association” based on their co-expression with genes that are known members of such gene sets. A total of 14,461 functionally defined gene sets served as input, taken from standard databases such as Gene Ontology[184] (GO), Reactome[185,186], and Kyoto Encyclopedia of Genes and Genomes[187] (KEGG). These co-expression patterns also serve as input for DEPICT[168] (**Supplementary Information section 12.4**). All members of the functionally defined gene sets or pathways share a particular biological function, which can range from very specific and bottom-up functions (e.g., “clathrin binding”) to more coarsely defined and top-down functions (e.g., “hormone activity”). The biological function can also refer to the cellular site where the gene performs its function (e.g., “neuromuscular junction”).

We briefly describe the Gene Network method here, noting that in-depth documentation of the method can be found in Fehrmann *et al.*[182] and Pers *et al.*[168]. The Supplementary Information of Okbay *et al.*[18] also contains a detailed description. Gene Network uses information from a total of 77,840 publicly available human, mouse, and rat Affymetrix microarrays from the Gene Expression Omnibus. These microarrays measure expression levels of at least 19,997 human genes (or, in the case of mouse and rat experiments, their mouse- or rat-orthologs). Gene co-expression levels were measured in Pearson correlation coefficients. The resulting correlation matrix was broken down into principal components (PCs), which gave rise to a total of 2,206 reliable “transcriptional components” (TCs).

Each PC (i.e. transcriptional component) captures a shared pattern of gene expression across experiments, implying shared biology of the genes that load highly on the PC. The PCs can capture subtle patterns of co-expression that potentially reflect shared biological pathways among genes, which would typically be overshadowed by strong global transcriptomic effects, and thus missed in a straightforward correlation analysis.

Subsequently, the authors of this method tested whether the pattern of co-expression captured by the PC significantly differentiated members of the gene set from all other genes, by testing the difference between the mean PC loading of the gene set versus the mean loading of all genes *not* in the gene set, using a Welch $t$-test for unequal variances. Each row of the resultant $14{,}461 \times 2{,}206$ matrix of $t$-statistics (each element corresponding to a gene set and PC) was then correlated with the PC loadings of the individual genes, to test whether the expression patterns of each gene and gene set were correlated. Finally, the thresholds for declaring a correlation between a gene's PC loadings and a gene set's $t$-statistics as statistically significant were chosen to satisfy False Discovery Rate (FDR) $< 0.05$. Thus, each gene's membership to each gene set was tested, allowing significant associations to be queried from the Gene Network database.

From this database, we first recorded for each MAGMA gene all statistically significant results (i.e., all gene sets that are significantly correlated with the genes' PC loadings), derived from gene sets and pathways from the Gene Ontology[184] (http://amigo.geneontology.org/amigo) domains Biological Process, Cellular Compartment, and Molecular Function. We also recorded the results listed under the Reactome[185] (http://www.reactome.org/) pathways and under the Kyoto Encyclopedia of Genes and Genomes[187] (KEGG, http://www.genome.jp/kegg/) pathways. In each case, we report (in **Supplementary Table 12.4**) the five most frequently occurring gene sets across the MAGMA genes.

Gene Network also reports organs, tissues, and cell types in which the gene under investigation is significantly and specifically expressed (compared to other tissues). For each MAGMA gene, we recorded all these organs, tissues and cell types where the area under the receiver operating characteristic curve (AUC), with respect to the discriminating power of measured gene expression, exceeded 0.80 according to Gene Network. Gene Network derived this AUC from the difference between the samples of the focal tissue and all other tissues in the distribution of the queried gene's expression level. These differences were determined by text-mining the descriptions provided by experimenters who uploaded the expression data to the Gene Expression Omnibus (GEO). Note that the tissue/cell type labels taken from the Medical Subject Headings (MeSH) database can refer to different levels of a hierarchy and are therefore not necessarily mutually exclusive.

We note that the method has three potential inherent drawbacks. First, only genes with reliable gene expression data (i.e., which have survived quality control) are included in the database. Therefore, the co-expression matrix might exclude genes that may be important but that have not yielded reliable expression data. For instance, *FTO* (a gene implicated in obesity) is a notable gene not included in the database.

Second, the method does not assign functions to genes that have unique patterns of gene expression but that may well play a role in known biological processes.

The third potential drawback of our querying method is that some of the significant MAGMA genes may only capture signal from other, nearby MAGMA genes, and not have any independent causal effects of their own. In this analysis, we conduct the search for each of the MAGMA genes and sum their predicted gene functions as if they were independent. This potential dependence among genes must, however, be taken into account when interpreting the predicted gene function counts.

For each of the three Gene Ontology (GO) domains, the Reactome pathways, the KEGG pathways, and the significant organs, tissues, and cell types, **Supplementary Table 12.4** lists the five most frequently occurring predicted gene functions, cellular components, and cell-type/tissues for the 266 MAGMA genes that were present in the Gene Network database. The most frequently occurring terms overwhelmingly point to neural function and anatomy. To a lesser degree, functions related to gene expression are also highlighted.

The top five tissues are all brain tissues: prefrontal cortex, which is predicted for 88 genes, frontal lobe (81 genes), visual cortex (81 genes), parietal lobe (80 genes), and putamen (which is part of the basal ganglia; 80 genes). The most interesting of these tissues is likely the prefrontal cortex, which is crucial for inhibition of emotion and planning of behavior[188], and is one of the last brain regions to mature during pubertal development[189].

The top terms for GO biological process and GO molecular function are “glutamate signaling pathway” (12 genes) and “glutamate receptor activity” (15 genes), respectively, while the top occurring terms for GO cellular component refer to dendritic (22 genes) and synaptic (22 genes) cell components. GO biological process also infers several functions related to modification of gene expression, namely “chromatin modification and organization” (11 genes) and “histone modification” (9 genes). The top terms for Reactome are “neuronal system” (20 genes), with the remainder of top terms referring to synaptic and neurotransmitter functions. Four out of five terms for KEGG are neural, with “calcium signaling pathway” (20 genes) as the top term. The only non-neural annotation for KEGG is “spliceosome” (18 genes).

#### 12.3 Identifying genes associated with general risk tolerance with SMR

##### *12.3.1 Background*

The rationale for the Summary-based Mendelian Randomization (SMR) test, which is one of several methods that test for an association between gene expression and trait variation (e.g., refs.[190–192]), is that GWAS associations for complex traits and common diseases are enriched in non-coding regulatory regions of the genome (e.g., ref.[69]), suggesting that many causal variants influence trait variation through regulation of gene expression. There is also evidence that causal variants often do not target the gene closest to the association signal (e.g., ref.[193]).

SMR differs from our eQTL lookups of top GWAS SNPs (**Supplementary Information section 12.6.3**) in several important ways. First, it differs in its starting point: instead of testing *all top GWAS SNPs* for association with a probe, it tests *all gene expression probes* with a significant SNP association (i.e., all significant *cis*-eQTLs in a certain tissue dataset) for association with the phenotype. To this end, it uses SNPs as instrumental variables for gene expression levels to estimate the association between gene expression levels and the phenotype (as described in further detail below).

In this sense, SMR is more powerful than the eQTL lookups, where the number of tests is informed by the stringent GWAS $P$ value threshold (as only genome-wide significant SNPs are put forward for eQTL testing). In SMR, the number of tests is informed by the number of significant eQTLs, which runs in the thousands. In this way, SMR can in principle uncover many more gene-phenotype associations than the eQTL lookups.

In addition, the "HEIDI" test feature of SMR is able to discard eQTL associations that are caused by mere linkage, thereby discarding genes that are "spuriously" associated with the phenotype (as described in further detail below). This feature distinguishes SMR from other transcriptome-wide analysis methods, and from the eQTL lookups, which may uncover genes that are merely associated with the phenotype due to linkage. We still perform the eQTL lookups, however, as they can give us some insight into the functional annotation of our GWAS top hits.

##### *12.3.2 Methods*

Summary statistic-based Mendelian Randomization (SMR)[192] is a method for estimating the effect of an exposure (*x*), such as gene expression, on a phenotype of interest (*y*), using summary-level data on estimated SNP effects (*z*, the instrumental variable) from large-scale GWAS and eQTL studies. In a Mendelian Randomization (MR) framework, the effect of *x* on *y* ($b_{xy}$) is:

$$\hat{b}_{xy} = \hat{b}_{zy}/\hat{\beta}_{zx},$$

where $\hat{b}_{zy}$ is the estimated SNP effect from a GWAS for trait $y$ and $\hat{\beta}_{zx}$ is the estimated SNP effect on the expression level of gene *x* from an independent eQTL study.

The sampling variance of $\hat{b}_{xy}$ is given by:

$$var(\hat{b}_{xy}) \approx \frac{b_{zy}^2}{\beta_{zx}^2}\left[\frac{var(\hat{\beta}_{zx})}{\beta_{zx}^2} + \frac{var(\hat{b}_{zy})}{b_{zy}^2} - \frac{2cov(\hat{\beta}_{zx}, \hat{b}_{zy})}{\beta_{zx}b_{zy}}\right],$$

where $cov(\hat{\beta}_{zx}, \hat{b}_{zy})$ is assumed to be zero if $\beta_{zx}$ and $b_{zy}$ are estimated in independent cohorts. If $\hat{\beta}_{zx}$ and $\hat{b}_{zy}$ are used in place of $\beta_{zx}$ and $b_{zy}$ (which are unknown), then the SMR test statistic, which is approximately $\chi^2$, takes the form:

$$T_{SMR} = \hat{b}_{xy}^2/var(\hat{b}_{xy}) \approx \frac{z_{zy}^2 z_{zx}^2}{z_{zy}^2 + z_{zx}^2},$$

where $z_{zy}$ and $z_{zx}$ are the estimated *z* statistics from the GWAS and eQTL studies, respectively.

A significant SMR test result could be due to causality (i.e., if the effect of *z* on *y* is via *x*), pleiotropy (i.e., if *z* has pleiotropic effects on *x* and *y*) or linkage (i.e., if $z_1$ affects *x* and $z_2$ affects *y*, and $z_1$ and $z_2$ are in linkage disequilibrium). Like all MR methods, SMR is unable to distinguish between causality and pleiotropy on the basis of a single genetic instrumental variable (i.e., the top *cis*-eQTL as opposed to multiple *cis* and *trans*-eQTLs), and so we do not distinguish between pleiotropy and causality in our analyses.

On the other hand, it is possible, under some assumptions, to distinguish pleiotropy (or causality) from linkage using the HEIDI (Heterogeneity In Dependent Instruments) test developed by Zhu *et al.*[192]. The rationale for the HEIDI test is that if the same causal variant has a pleiotropic effect on gene expression and the trait, then the estimate of $b_{xy}$ should be identical for any SNP in LD with the causal variant. Therefore, a test for linkage is equivalent to testing for heterogeneity in the estimates of $b_{xy}$ for the top *cis*-eQTL and any other significant SNPs in the *cis*-eQTL region.

If there are $m$ such variants (excluding the top *cis*-eQTL), and if we define $d_i = b_{xy(i)} - b_{xy(top)}$ and $\hat{\mathbf{d}} = \{\hat{d}_1, \hat{d}_2, \dots \hat{d}_m\}$, then the HEIDI test against the null hypothesis **d** = 0 takes the form:

$$T_{HEIDI} = \sum_{i}^{m} z_{d(i)}^2,$$

where $z_{d(i)}$ equals $\hat{d}_i / \sqrt{var(\hat{d}_i)}$. SNPs with $r^2 > 0.9$ with the top *cis*-eQTL are excluded from the set of $m$ SNPs in the HEIDI test, because statistical power to distinguish between linkage and pleiotropy is limited if the causal variants have close to perfect LD. The test also excludes SNPs with eQTL $P$ values > $1.6 \times 10^{-3}$, due to the need to remove weak instrumental variables.

For a complete description of the SMR and HEIDI tests, see Zhu *et al.*[192].

##### *12.3.3 Data*

SMR requires summary statistics from large-scale GWAS and eQTL studies and genotype data from an ancestry-matched reference sample for estimating linkage disequilibrium. We used estimates of SNP effects ($b_{zy}$) from the meta-analysis of our discovery and replication GWAS of general risk tolerance, and we used summary data on *cis*-eQTLs ($b_{zx}$) from the eQTLgen Consortium's meta-analysis of 14,115 whole blood and peripheral blood samples (in prep.). We excluded probes in the MHC (26.1 to 33.8Mb on chromosome 6 based on hg19) due to the complexity of linkage disequilibrium in this genomic region, and we excluded SNPs with MAF < 0.01 and INFO score < 0.3. Analyses were restricted to probes with at least one *cis*-eQTL (defined as +/-1Mb from the middle of the probe) with $P < 5 \times 10^{-8}$, because Mendelian randomization assumes that the instrumental variable has a strong effect on the exposure. After filtering we had access to summary data on 12,751 probes and 10,209,777 (1000 Genomes imputed) SNPs. We used Bonferroni correction to account for multiple testing, resulting in a genome-wide significance threshold for the SMR test of $P < 3.9 \times 10^{-6}$ (= 0.05/12,751).

We based our primary analyses on eQTLs identified in blood (in the eQTLgen data) because the available sample size is at least an order of magnitude larger than for any other human tissue, and so this strategy maximizes statistical power for genes with shared genetic control of regulation across tissues. However, genetic control of regulation of some genes is tissue-specific, and so for each of the 21 probes passing both the SMR and HEIDI tests in the eQTLgen analysis, we also performed SMR analyses using estimates of SNP effects on gene expression in the human brain. We used summary eQTL data from the Genotype Tissue Expression Consortium (GTEx) for 11 brain regions (anterior cingulate cortex, caudate basal ganglia, cerebellar hemisphere, cerebellum, cortex, frontal cortex, hippocampus, hypothalamus, nucleus accumbens basal ganglia, pituitary, putamen basal ganglia) with sample size exceeding 70 (GTEx release v6p). We mapped Illumina expression probes in eQTLgen to GTEx transcripts using Ensembl gene IDs. We applied a relaxed eQTL $P$ value threshold for inclusion of transcripts in the analysis of brain eQTLs ($P < 2.5 \times 10^{-3}$), due to the small number of transcripts tested per tissue, and we used a correspondingly lower significance threshold for the SMR test ($P < 2.16 \times 10^{-4}$, based on correction for testing of 21 probes in each of 11 brain regions).

We used our main reference panel (**Supplementary Information section 2.4)** for LD estimation. All analyses used hg19 coordinates.

##### *12.3.4 Results and discussion*

We used SMR to test for association between gene expression in blood and general risk tolerance using summary data from our GWAS meta-analysis and blood *cis*-eQTLs from the eQTLgen consortium. We identified 42 genes (tagged by 46 probes) at experiment-wide significance ($P < 3.9 \times 10^{-6}$), of which 19 (tagged by 21 probes) passed the HEIDI test, indicating we could not reject the null hypothesis of a single causal variant with pleiotropic effects on both risk tolerance and gene expression in blood (**Supplementary Table 12.5**). Thirteen of the 19 significant genes (68%) were in reported genome-wide significant loci for general risk tolerance, whereas the remaining six (*CTNNA1*, *SIL1*, *ZCCHC7*, *HNRNPK*, *HINFP*, *LYZ*) were "novel" discoveries, insomuch as no reported general-risk-tolerance GWAS locus was located within 0.5Mb. These latter discoveries point to genetic loci that are likely to be identified with genome-wide significance in larger GWAS of general risk tolerance. Several genes passing the SMR and HEIDI tests, including the top ranked gene *MSRA* ($P_{\text{SMR}} = 1.8 \times 10^{-13}$, $P_{\text{HEIDI}} = 8.8 \times 10^{-2}$), *ERI1* ($P_{\text{SMR}} = 1.1 \times 10^{-7}$, $P_{\text{HEIDI}} = 8.5 \times 10^{-2}$) and *XKR6* ($P_{\text{SMR}} = 4.3 \times 10^{-10}$, $P_{\text{HEIDI}} = 0.59$), were also significantly associated with general risk tolerance in the MAGMA analysis (**Supplementary Information section 12.2**).

For each of the 21 probes passing both the SMR and HEIDI tests in the eQTLgen blood analysis, we repeated the SMR analysis for general risk tolerance using summary data on *cis*-eQTLs from the 11 GTEx v6p brain regions (**Supplementary Table 12.5**). Three genes passed both the SMR and HEIDI tests in one or more brain regions: *CTNNA1* (chromosome 5, 138.11Mb) was significant in caudate basal ganglia, cerebellum and cerebellar hemisphere; *CENPV* (chromosome 17, 16.25Mb) was significant in putamen basal ganglia, nucleus accumbens basal ganglia, caudate basal ganglia, anterior cingulate cortex (BA24), hippocampus and hypothalamus; and *ZSWIM7* (chromosome 17, 15.89Mb) was significant in cerebellum, cerebellar hemisphere, cortex, anterior cingulate cortex (BA24) and frontal cortex (BA9). These brain regions overlap with those identified by the DEPICT tissue prioritization analysis (see **Supplementary Information section 12.4**). For *CTNNA1* and *CENPV* (**Extended Data Fig. 12.2**), the risk-tolerance increasing allele was associated with decreased gene expression in blood and each of the respective brain regions. For *ZSWIM7*, the risk-tolerance increasing allele was associated with increased gene expression in blood, cerebellum and cerebellar hemisphere, but the results were inconsistent for cortex, anterior cingulate cortex and frontal cortex (i.e., the risk-tolerance increasing allele was associated with decreased expression). These findings suggest that *CTNNA1* and *CENPV* are putatively functional genes for risk-taking behavior but that more investigation is required to confirm the association with *ZSWIM7*.

According to the GTEx portal website, *CENPV* is highly expressed in many tissues, but especially in the brain's cerebellum, while *CTNNA1* is not particularly strongly expressed in brain tissue (compared to the other tissues). The function of *CENPV* has not been studied frequently, but two studies have discovered functions in cell division[194] and migration[195]. *CTNNA1,* on the other hand, has been studied widely for its role in tumor suppression (e.g. ref.[196]), and mutations in this gene are known to lead to retinal pigment dystrophy, causing macular disease[197]. Both *CTNNA1* and *CENPV* were significantly associated in the MAGMA gene based test, but neither were prioritized by DEPICT at FDR > 0.20. Therefore, the putative roles *CENPV* and *CTNNA1* may play in general risk tolerance remain unclear.

#### 12.4 Prioritization of tissues, gene sets, and genes using DEPICT

##### *12.4.1 Background and methods*

DEPICT (Data-driven Expression Prioritized Integration for Complex Traits) is a computational tool that uses a set of GWAS summary statistics as input and allows enrichment analysis of tissues and gene sets and prioritization of genes in the associated loci (see ref.[168] for details).

For this work, we used DEPICT release 194[iii]. DEPICT was run using the summary statistics of the meta-analysis of the discovery and replication GWAS of general risk tolerance (after applying genomic control using the intercept of the LD Score regression). Only SNPs with GWAS *P* values less than $10^{-5}$ were used as input, and DEPICT-defined loci were defined by clumping these SNPs (PLINK clumping parameters: --clump-p1 1e-5 --clump-r2 0.1 --clump-kb 500, using 1000 Genomes Project pilot phase data as reference). Locus boundaries were then defined using a LD $r^2$ threshold of 0.5, and overlapping loci were merged, yielding 464 autosomal loci comprising 1,060 genes. DEPICT was run using default settings: 500 permutations for bias adjustment; 50 replications for false discovery rate estimation; normalized expression data from 77,840 Affymetrix microarrays for gene set reconstitution; 10,968 reconstituted gene sets for gene set enrichment analysis; and testing 209 tissue/cell types assembled from Medical Subject Headings (MeSH) annotations from 37,427 Affymetrix U133 Plus 2.0 Array samples for enrichment in tissue/cell type expression[168,182]. Furthermore, gene expression data from 37 RNA-sequenced tissues from the Genotype-Tissue Expression (GTEx) Consortium[58] were used for tissue enrichment analysis.

##### *12.4.2 Results: Prioritized tissues, gene sets, and genes*

We begin by reporting the results of the DEPICT tissue enrichment analysis, which we conducted using GTEx RNA-sequencing gene expression data as well as using microarray data from 209 tissues[168,182]. The DEPICT tissue enrichment analysis identifies tissues in which genes near general-risk-tolerance-associated SNPs are significantly overexpressed relative to genes in random sets of loci matched by gene density.

Using GTEx RNA-sequencing gene expression data, we identified eight significantly enriched tissues (FDR < 0.01), which all represent different areas of the brain (**Fig. 3b** and **Supplementary Table 12.6**). The top three prioritized tissues were cortical brain regions, namely “Frontal Cortex (BA9),” “Anterior Cingulate Cortex (BA24)” and “Cortex.” The cortical regions BA24 and BA9 are respectively located in the ventral anterior cingulate area and in the dorsolateral and medial prefrontal cortex[198,199]. These two regions had initially been respectively associated with memory and emotion processing, though more recent work stresses their role in executive functions such as “executive control of behavior,” “inferential reasoning,” and “learning and decision making”[200–204]. The DEPICT tissue prioritization analysis with the GTEx data also points to subcortical regions, including notably the nucleus accumbens, caudate, and putamen, which form the basal ganglia[205], as well as the amygdala. The basal ganglia and amygdala are known for their respective role in motor control (and its impairment in Parkinson’s disease), and in the processing of negative emotions, such as fear. However, recent evidence also

[iii] DEPICT release 194 handles GWAS data imputed with 1000 Genomes Project reference data and was downloaded in February 2016 from:
https://data.broadinstitute.org/mpg/depict/depict_download/bundles/DEPICT_v1_rel194.tar.gz.

supports a role for the basal ganglia and the amygdala in reward processing and decision-making[206–209].

Using microarray data from 209 tissues, we replicated the brain enrichment and identified overall 19 significantly enriched tissues (FDR < 0.01) which are all part of the central nervous system (CNS) (**Extended Data Fig. 12.3b** and **Supplementary Table 12.7**). Note that we also observed an enrichment for the retina, which is part of the CNS but in DEPICT is categorized as "Sense Organs." This analysis additionally showed expression in deeper brain structures, such as the rhombencephalon and metencephalon. The metencephalon, also known as the midbrain, is the major location of dopamine neurons[210] ; we note, however, that this exact location, known as substantia nigra, was not prioritized by DEPICT in the analysis of GTEx tissues (FDR > 0.20). Moreover, the results seem to have little specificity toward the reward and decision-making system, as they also show expression in other CNS regions with diverse functions (e.g., the visual cortex).

Next, we report the results of the DEPICT gene set enrichment and gene prioritization analyses. DEPICT "reconstituted gene sets" are predefined gene sets taken from several bioinformatic databases (including Gene Ontology, Reactome, and Kyoto Encyclopedia of Genes and Genomes) that have been reconstituted using co-expression data to represent the weight of evidence of a given gene belonging to the gene set—as opposed to the all-or-nothing representation for the predefined gene sets (see Pers *et al.*, 2015[168], for details**)**. Each gene's membership score with respect to a given reconstituted gene set is simply the *z*-statistic of the correlation between the gene's vector of transcriptional components (TC) loadings and the binary gene set's *t*-statistics with respect to the TCs—the same *z*-statistics used for testing the statistical significance of the Gene Network predictions. We identified 93 reconstituted gene sets that are significantly enriched (FDR < 0.01) for genes found among the general-risk-tolerance-associated loci (**Supplementary Table 12.8**). We used the Affinity Propagation tool[211] to cluster related reconstituted gene sets into a network diagram (**Fig. 3a**)[jjj]. The most strongly enriched gene sets highlight pathways related to the CNS. These include gene sets for regulation of neuronal projections (most notably synaptic and dendritic development) and synaptic transmission. Several of the most significant pathways relate to synaptic transmission activity, in particular GABA and glutamate neurotransmitter signaling. We were able to prioritize at least one gene at 106 of 464 loci defined by DEPICT (23% of the loci) at FDR < 0.01. In total DEPICT prioritized 122 genes **(Supplementary Table 12.9)** across those 106 loci.

The most significantly enriched reconstituted gene set is "dendrite development." The presence of dendrites distinguishes neurons from all other cell types; these unique structures bestow the ability to receive signals of high spatiotemporal resolution. We mention just one aspect of dendrite development implicated by our results: the coordination or mutual influence required to juxtapose the axonic and dendritic sides of the developing synapse. In certain hippocampal pyramidal neurons, recognition molecules of the NGL/NTNG family produce a compartmentalization of the dendritic tree by ensuring the innervation of distinct portions of the tree by axons from correspondingly distinct sources. *NTGN1* and *NTNG2* encode ligands for receptors encoded by *LRRC4C* (also called *NGL1*) and *LRRC4* (also called *NGL2*) respectively (*NTGN1* is not prioritized by DEPICT, but *NTNG2* and *LRRC4* are prioritized at FDR < 0.01 and *LRRC4C* is prioritized at FDR < 0.05). *LRRC4C* and *LRRC4* are members of the leucine-rich

[jjj] The script to produce the network diagram can be downloaded from https://github.com/perslab/DEPICT.

repeat family, whose overarching function may be the exploitation of both family size and alternative splicing to create unique molecular fingerprints enabling precision wiring between neurons[212,213]. Incoming axons bearing NTGN1 and NTNG2 respectively somehow find their corresponding receptors (LRRC4C, LRRC4) and thereby innervate non-overlapping compartments on distal and proximal dendrites[214]. These leucine-rich repeats are synaptic cell adhesion molecules (CAMs, such as *CADM2*) that transmit signals to the dendritic interior upon trans-synaptic binding, and thus their compartmental distribution may bestow varying information-processing properties along the length of a dendritic tree.

Our results implicate a number of different neurotransmitter receptors embedded in the membrane of the mature dendrite. Glutamate is the most abundantly employed excitatory neurotransmitter in the brain. When glutamate released from the axon of a transmitting presynaptic neuron binds to a fast-acting AMPA-type glutamate receptor embedded in the dendrite of the receiving postsynaptic neuron, the channel coupled to the receptor momentarily opens by a lock-and-key mechanism and permits a massive influx of $Na^+$ ions. This is the primary mechanism leading to rapid depolarization of the receiving neuron, which can lead in turn to the neuron firing its own signal along its axon. Our DEPICT-prioritized genes include *GRIA1*, which encodes the most common subunit appearing in the AMPA-type glutamate receptor.

Two genes encoding subunits of NMDA-type glutamate receptors are among our DEPICT-prioritized genes (*GRIN2A*, *GRIN2B*). NMDA-type glutamate receptors act like coincidence detectors: they only open when glutamate has landed on the receptor and the postsynaptic membrane potential has been sufficiently depolarized by the opening of other excitatory channels to remove the $Mg^{2+}$ block. The emerging picture is of a receptor whose task is not to respond immediately to any given input impinging on the dendrite but rather to admit an agent of change into the dendrite in response to a temporal coincidence of synapse-specific input and membrane depolarization.

So far we have been discussing excitatory neurotransmission—that is, a signal from one neuron to the next that pushes the receiving neuron closer to its own firing threshold. Now we turn to inhibitory neurotransmission, which has the effect of pushing the receiving neuron further away from its firing threshold. The most important source of inhibition in the cortex is mediated by receptors for the neurotransmitter GABA. They produce inhibition in many cases by generating hyperpolarizing changes in membrane potential that counteract the depolarizing changes caused by Na+ influx through opened glutamate receptors. $GABA_A$ receptors mediate the effects of alcohol intoxication and drugs such as benzodiazepines and barbiturates, which have anxiolytic effects

A GABAergic neuron typically establishes outgoing connections only with neurons in its local vicinity. A glutamatergic pyramidal neuron, in contrast, often sends its axon to neurons that are far away (e.g., the opposite hemisphere). When this local connectivity of GABAergic neurons is emphasized, they are called interneurons. Because it is not possible to construct an information-processing device from only inhibitory neurons, we can reasonably say that they play some supporting or modulatory role.

$GABA_A$ receptor subunits are grouped into subfamilies known respectively as $\alpha$, $\beta$, $\gamma$, $\delta$, $\varepsilon$, $\theta$, $\varphi$, and $\rho$[215], the first three of which are represented in our DEPICT-prioritized genes (*GABRA2*, *GABRA4*, *GABRB1*, *GABRG3*). A complete receptor is composed of five subunits; the combinatorial number of possible pentamers is quite impressive, even if we put aside alternative splicing. Some subunits of $GABA_A$ receptors contain regulatory sites for phosphorylation and domains that interact with proteins such as the product of *GPHN* involved in receptor trafficking and localization at synapses[216].

The importance of both excitatory and inhibitory neurotransmission is affirmed by the significant enrichment of gene sets such as "glutamate receptor activity," "abnormal miniature excitatory postsynaptic currents," and "$GABA_A$ receptor activity." Note that the cluster named after the latter gene set contains the second-most significant gene set in the entire analysis (**Fig. 3a**).

Four out of the top five prioritized genes encode proteins with transcriptional regulatory activity, some known to play a role in brain development. The strongest prioritized gene, *FOXG1*, encodes a transcription factor known to cause the congenital variant of Rett-syndrome[217], a severe neurodevelopmental disease. The second strongest prioritized gene, *ASH1L*, encodes a histone lysine N-methyltransferase involved in chromatin modification and has been implicated in autism and neurodevelopmental disorders[218]. The third strongest prioritized gene, *FAM190A* (alias *CCSER1*), is a relatively unknown gene with no previous reported associations to brain-related phenotypes. The fourth and fifth strongest prioritized genes, *FOXO6* and *NEUROD*, both encode transcription factors and have been implicated in Alzheimer's Disease and regulation of memory consolidation, respectively[219,220].

#### 12.5 Competitive gene-set analysis with MAGMA: Testing gene sets related to GABA and glutamate neurotransmitters

##### *12.5.1 Background*

Since both the DEPICT pathway analysis and the Gene Network analysis of MAGMA genes both pointed to glutamate and GABA neurotransmitters, we decided to also perform an *ex post* MAGMA competitive gene-set analysis of relevant glutamate and GABA gene sets. This allowed us to directly compare the enrichment of these gene sets to that of the gene sets associated with dopamine, serotonin, testosterone, estrogen, and cortisol, which we also tested in a MAGMA gene set analysis described, as described in **Supplementary Information section 11.2**.

Similar to what we did in **Supplementary Information section 11.2**, we looked up all relevant glutamate and GABA gene sets in MSigDB v6.1[kkk] and merged the resulting sets into one glutamate and one GABA superset. In addition, we tested the GABA and glutamate gene sets which were significant (FDR < 0.05) in DEPICT (see **Supplementary Information section 12.4**), although we updated the genes in those sets according to the gene sets in MSigDB v6.1.

Thus, in total, we conducted a MAGMA gene set analysis to test four glutamate and GABA gene sets (Panel **B** in **Supplementary Table 11.2**):

1) All GABA sets significant in DEPICT (FDR < 0.05, six gene sets) or with more than 10 counts in the Gene Network analysis of MAGMA genes (one gene set), containing 68 unique genes

[kkk] See http://software.broadinstitute.org/cancer/software/gsea/wiki/index.php/MSigDB_v6.1_Release_Notes for details on this release

2) All glutamate sets significant in DEPICT (FDR < 0.05, 12 gene sets) or with more than 10 counts in the Gene Network analysis of MAGMA genes (zero gene sets), containing 159 unique genes
3) All relevant GABA sets in MSigDB v6.1 (13 gene sets), containing 134 unique genes
4) All relevant glutamate sets in MSigDB v6.1 (27 gene sets), containing 276 unique genes

We tested all four sets together, and MAGMA computed the family-wise error correction accordingly.

###### *12.5.2 Results and discussion*

None of the four gene sets were significant (lower Bonferroni-corrected $P$ = 0.299, Panel B in **Supplementary Table 11.3**). We are unsure why this might be the case but, as we further discuss this in **Supplementary Information section 12.7.4**, our MAGMA competitive gene-set analysis of the four glutamate and GABA gene sets has some likely limitations. First, the statistical power of a MAGMA gene set analysis fails to increase substantially with increasing sample sizes[163]. Second, important effects of single SNPs in a gene set may be overshadowed by the high average $P$ values of other SNPs in the gene or gene set. Third, it might be that only specific glutamate and/or GABA genes or pathways might be relevant for general risk tolerance, and that our merging of those pathways into major glutamate and GABA sets decreased our statistical power for discovery of relevant effects. (In the MAGMA gene-based analysis, only 12 of the 354 unique GABA/glutamate genes included in the four gene sets tested here were significant at Bonferroni-corrected $P$ < 0.05.)

#### 12.6 Lookup of protein-altering variants and *cis*-eQTLs

In this section, we conducted several analyses that annotate the 132 lead SNPs from the meta-analysis of the discovery and replication GWAS of general risk tolerance, to gain insights about the biological systems they may affect. (For consistency with the rest of the bioannotation analyses, we performed these analyses with the 132 lead SNPs of the meta-analysis combining the discovery and replication GWAS of general risk tolerance, and not with the 124 lead SNPs from the discovery GWAS.[III])

[III] Because the lead SNPs of the discovery GWAS are not a perfect subset of the lead SNPs of the meta-analysis of the discovery and replication GWAS, we investigated the overlap of the identified loci. The discovery GWAS identified 124 lead SNPs, and the meta-analysis of the discovery and replication GWAS identified 132 lead SNPs; 97 of these are lead SNPs in both. We investigated if the non-overlapping lead SNPs of each GWAS are in LD with a lead SNP of the other (defined here as $r^2$ > 0.1). The discovery GWAS identified seven lead SNPs that are not located in the locus of a lead SNP of the meta-analysis of the discovery and replication GWAS, and the meta-analysis of the discovery and replication GWAS identified 15 lead SNPs that are not located in the locus of a lead SNP of the discovery GWAS. (The list of these seven and 15 SNPs is available upon request.) These seven and 15 lead SNPs are all highly significant ($P$ < 5E–7) in the other GWAS. In conclusion, approximately 117 loci overlap across the two GWAS, and the 15 additionally identified loci in the meta-analysis of the discovery and replication GWAS are all highly significant in the discovery GWAS. The slightly larger number of lead SNPs in the meta-analysis of the discovery and replication GWAS was to be expected because of the increased statistical power due to the inclusion of the replication GWAS.

Specifically, we examined the overlap between these lead SNPs and various SNPs and genes that have been identified or annotated in previous GWAS and functional genomic databases. First, we checked whether our lead loci contain protein-altering variants. Second, we looked up the lead loci in an expression quantitative trait loci (eQTL) database.

##### *12.6.1 SNPs in LD with lead SNPs*

Since the genome is characterized by widespread linkage disequilibrium (LD), it is important to examine the overlap with SNPs that are in LD with our lead SNPs (as opposed to only considering our lead SNPs). Moreover, some data sources do not contain some of our lead SNPs but contain SNPs in strong LD with them, and a lead SNP does not actually have to be the causal SNP in the locus: it may just be the most often genotyped or most accurately imputed SNP (i.e., the SNP measured with the least measurement error and therefore yielding a lower *P* value); it may tag an unmeasured causal SNP in the LD block; or it may simply be the most significant SNP in the block due to sampling variation. For these reasons, it is crucial to also consider the LD partners of the lead SNP. We obtained a list of SNPs in strong LD with our lead SNPs using the PLINK –r2 command in our main reference panel (described in **Supplementary Information section 2.4**). Here we used an LD cutoff of $r^2 > 0.6$ and a distance cutoff of 250kb from the lead SNP. We refer to these SNPs as the "LD partners" of our lead SNPs. In total, the clumping procedure designated 132 "lead SNPs," which tagged a total of 8,199 LD partners.

##### *12.6.2 LD with protein-altering variants*

Using the Haploreg v4.1 database[221] (download date 18 May 2016, http://www.broadinstitute.org/mammals/haploreg/haploreg.php), we ascertained the protein-altering status of our lead SNPs and their LD partners. More precisely, we ascertained whether our lead SNPs and their LD partners are annotated as "missense," "nonsense," "splice site donor/acceptor," or "frameshift" in the 1000 Genomes Phase 3 European population. The Haploreg database uses annotations from dbSNP[55] to determine functional status. We examined these annotations because protein-altering variants have a higher *a priori* probability of being causally relevant: they can result in stronger phenotypic differences between individuals compared to non-protein-altering variants. Most of the genome is non-coding, which underlines the importance of the detection of protein-altering variants.

The results can be found in **Supplementary Table 12.10**. We found that there were eight protein-altering SNPs among the lead SNPs' LD partners, including two of the lead SNPs themselves. All protein-altering variants were relatively common (with $0.11 < MAF < 0.50$) missense variants (meaning that they result in an amino acid change). The two missense lead SNPs are in genes *WSCD2* (on chromosome 12) and *NF1* (on chromosome 17). Little is known about *WSCD2,* but it is highly expressed in the brain and has predicted functions in long term memory and calcium signaling, according to Gene Network[182]. *NF1* is involved in the genetic disorder "neurofibromatosis," which is characterized by widespread fibromatous tumor formation and cognitive disability[222]. Other missense variants in LD with our lead SNPs were found in *FREM1* and *BHLHE22. FREM1* encodes a protein that may play a role in craniofacial and renal development, while *BHLHE22* is thought to be involved in the differentiation of sensory neurons. The remaining four missense variants in LD with our lead SNPs were in the HLA-region, of which three were in the olfactory receptor gene *OR2J2,* and one in histone gene *HIST1H1T.*

##### *12.6.3 eQTL lookup*

The publicly available results from the Genotype-tissue expression project (GTEx, www.gtexportal.org, version v6p)[58] allow us to look up significant associations between our lead SNP and their LD partners and *cis*-gene expression in distinct human tissues. The GTEx team obtained postmortem samples from human donors and used RNA sequencing to measure RNA levels in distinct tissues from those samples. The GTEx team also genotyped the donors with SNP microarrays. The lookup of the risk-tolerance lead SNPs and their LD partners was performed on the publicly accessible summary statistics (downloaded 13 December 2016, version v6p) of the significant gene-*cis*-SNP pairs from GTEx. We only considered the set of significant *cis*-eQTLs (i.e., gene-*cis*-SNP pairs), where the *cis*-window is defined as a 1MB window around the gene's transcription start site. In addition, when a gene transcript was a significant eQTL for more than one of our query SNPs, we only report the eQTL statistics (i.e., effect size, *P* value) for the SNP in highest LD with the lead SNP (which might be the lead SNP itself). That way, we only report each significant eQTL gene once in each tissue (even though the gene may have appeared several times for the tissue, for several lead SNPs and LD partners). In cases where more than one lead SNP was an eQTL for the same gene, we report the eQTL for only one of them.

To identify statistically significant *cis*-eQTLs, the GTEx consortium computes a gene-specific nominal *P* value threshold for each gene, with a permutation method as described in more detail in ref.[223]. The eQTLs we report here are all statistically significant, in the sense that the nominal *P* value of the slope of the regression of the gene's expression value on the SNP was smaller than the gene-specific nominal *P* value threshold. This regression also included several technical covariates, such as the top principal components of the genetic relatedness matrix.[mmm]

We only recorded findings for the following pre-selected tissues, which we suspected could be of biological relevance for risk tolerance: (1) all 11 brain tissues[nnn] available in the GTEx data; (2) tibial nerve tissue; (3) thyroid tissue; (4) gonadal (i.e. testis and ovary) tissue; and (5) adrenal gland tissue. In addition, we queried (6) the "whole blood" tissue for eQTLs, as this tissue had the highest sample size. Since expression profiles in whole blood and the brain are moderately correlated, we might uncover potentially interesting eQTLs by using information from whole blood[224,225].

The donor samples sizes for the brain tissues range from $n$ = 72 to 103, and sample sizes for the other five tissues range from $n$ = 126 to 338. We included tibial nerve tissue (a peripheral nerve located in the lower leg) because the eQTL results for this tissue are based on a relatively large sample size (compared to the brain tissues) of $n$ = 256. Since central and peripheral nervous tissues are anatomically similar, their expression profiles are likely to be comparable. Thyroid tissue is included because thyroid hormone is essential for healthy brain development, and might be associated with mood and cognition[226,227]. Testis and ovary tissue are included, as these organs produce sex hormones (e.g., testosterone, estrogen) that might be involved in risk tolerance, as detailed in our literature review (**Supplementary Information section 11.1**). Adrenal tissue is included because the adrenals produce hormones associated with the general stress response (i.e., (nor)epinephrine and cortisol), which might be related to risk tolerance, as

[mmm] For additional details, see http://www.gtexportal.org/home/documentationPage#staticTextAnalysisMethods and ref.[223].

[nnn] These include pituitary tissue (which GTEx Portal does not list among brain tissues).

detailed in our literature review (**Supplementary Information section 11.1**). Note that the presence of eQTLs in these tissues *does not* imply statistical enrichment of the GWAS signal in these tissues; it merely shows that our lead SNPs are associated with gene expression in these tissues (which might simply be a result of pleiotropy, and which thus might not be of causal relevance to general risk tolerance).

**Supplementary Table 12.11** reports the results. We find eQTL overlaps in every tissue (including all individual brain tissues) we queried. In total, we find 188 unique gene transcripts for 59 different lead SNPs. 82 of these gene transcripts only had Ensembl gene IDs and no gene symbols (i.e. letter symbol gene names such as *"HLA"* which is an abbreviation of Human Leukocyte Antigen, according to the HUGO gene nomenclature committee), most likely because they do not have named gene products (i.e., proteins). Out of the 188 unique gene transcripts, 47 lie in or near the human MHC (i.e., Major Histocompatibility Complex, also known as Human Leukocyte Antigen (HLA)) region (~chr. 6: 26-32 Mb), of which five are *HLA*-genes.

Finally, we note that there was one eQTL for a GABA gene (*GABRG1,* or $GABA_A$ receptor gamma1 subunit), in which the general-risk-tolerance-increasing allele decreased expression of *GABRG1* in tibial nerve tissue. We found no eQTLs for glutamate genes in any of the queried tissues.

#### 12.7 Summary of main findings from the biological annotation analyses

The overall goal of the analyses described in this section was to gain insight into the biology of general risk tolerance. We employed two general approaches. First, we leveraged data at the genome-wide level to prioritize potentially relevant tissues and genomic annotations (using LDSC partitioning of heritability) and genes (using MAGMA gene-based analysis, SMR transcriptome-wide analysis, and DEPICT gene prioritization analyses). Second, we focused on the loci around the general-risk-tolerance lead SNPs (in the lookups of protein-altering variants and GTEx eQTLs) to gain insight into the biological functions of our lead SNPs. Here, we summarize the results of those analyses.

##### *12.7.1 Enrichment of SNPs annotated to the central nervous system*

First, we tested enrichment of the GWAS signal in 10 different tissues/cell-types using LDSC partitioning of heritability. After controlling for tissue overlap and correcting for multiple testing, we found significant positive involvement for the central nervous system and the immune system. Involvement of the central nervous system was expected (since general risk tolerance is a behavioral trait), but enrichment of immune cell types might be surprising. To the best of our knowledge, immune enrichment in LDSC partitioning analyses has not previously been reported for a behavioral or psychiatric trait, although we note again that enrichment of *specific* immune pathways and/or immune tissue enhancer regions has been implicated in schizophrenia[69,179], bipolar disorder[180] and major depression[181]).

We note that enrichment of immune tissues or function did not come forward in any of the other analyses. Further research is therefore needed to corroborate a role for immune processes in general risk tolerance. Finally, we note that several additional tissues, such as liver and adrenal/pancreatic tissues, showed statistically significant enrichment, but that our estimates of their partial effects on enrichment (i.e., their $\tau_c$ coefficients) were not significant, thereby suggesting that their enrichment may have been primarily driven by other overlapping annotations. We therefore cannot claim a role for the adrenal system in general risk tolerance. However, this non-finding might be a result of the combination of adrenal with pancreas tissue, and the possible non-specificity of histone marks to these particular tissues. More fine-grained analyses are therefore needed to corroborate or refute a role for the adrenal system in modulating risk tolerance.

##### *12.7.2 Enrichment of SNPs near genes that are highly expressed in the frontal and prefrontal cortex*

While the LDSC partitioning results pointed to the central nervous system (which technically includes the brain, brainstem and spinal cord), results from the other analyses allowed us to gain deeper insight into which particular brain regions might be of relevance to general risk tolerance. Although the overlap between the genes prioritized by DEPICT and the significant MAGMA genes was not very high (only 71 out of 285 genes of the MAGMA genes were among the 215 genes prioritized at FDR < 0.05 by the DEPICT gene prioritization analysis), both the Gene Network functional analysis of the MAGMA genes and the DEPICT tissue enrichment analysis with the GTEx tissue-specific gene-expression data separately point to enrichment of the frontal cortex, and in particular the prefrontal cortex. The frontal cortex is a large and functionally and anatomically complex brain area; it is the largest neocortical region in humans and primates, and receives and integrates neuronal input from both cortical and subcortical regions[188]. The posterior dorsal area of the frontal cortex contains the motor cortex, which is responsible for planning and directing bodily movement; damage to this area can result in perturbed motor function or paralysis. The prefrontal cortex (the most anterior part of the brain), on the other hand, is involved in planning, directing, and inhibiting emotion, behavior and cognition (broadly known as "executive function"). Damage to the prefrontal cortex has been well-documented due to the widespread practice of prefrontal lobotomy (a relatively simple surgical procedure) in 20$^{th}$ century America and with textbook patient cases of accidental lesion such as Phineas Gage. Damage to the prefrontal cortex can result in severe impairment marked by behavioral disinhibition and emotional deregulation, cognitive disability, impaired decision making, a tendency to display context-inappropriate behavior, and inability to plan future action[188]. The frontal cortex is also one of the last brain areas to mature during development (with full maturation reached around age 25)[189], which has been postulated to underlie the social and behavioral disinhibition and increased display of risk-taking behaviors that characterizes puberty and adolescence[228–230].

At a finer resolution, the DEPICT tissue enrichment analyses with the GTEx tissue-specific gene-expression data show that genes near general-risk-tolerance-associated SNPs are highly expressed in the prefrontal cortex (BA24 and B9) and the basal ganglia (nucleus accumbens, caudate, putamen), compared to other organs and tissues. The DEPICT tissue enrichment analysis using the microarray data confirmed these CNS regions' enrichment, and additionally showed expression in deeper brain structures like the midbrain, among other tissues. The Gene Network tissue enrichment analyses also point to enrichment of the prefrontal cortex, and of the putamen, which is part of the basal ganglia. Finally, two of the three genes that passed both the SMR and HEIDI tests, *CTNNA1* and *CENPV*, were significant in parts of the basal ganglia (*CTNNA1* was significant in the caudate and *CENPV* was significant in the putamen, nucleus accumbens, and caudate).

##### *12.7.3 A possible convergence with results from the neuroscientific literature*

Interestingly, the prefrontal cortex, the basal ganglia and the midbrain are the major components of the *cortical-basal ganglia circuit*, which includes and connects the reward system, the motor system, and the cognitive and behavioral control systems. This whole network of brain regions has been shown to be critically involved in learning, motivation, behavioral control, and decision-making, notably under risk and uncertainty, in both humans and non-human primates[210,231]. Decision making under risk requires several cognitive processes (e.g. such as reward processing, fear, or higher executive control) underpinned by different neural systems which are coherently connected and integrated in current models of the *cortical-basal ganglia circuit*. In this complex circuit, the processing of reward and decision-making is presumed to be largely underpinned by dopamine[207,232]. Dopaminergic neurons are mostly located in the midbrain (substantia nigra and ventral tegmental area) where they project to the basal ganglia and the prefrontal cortex, and appear to encode signals about past and future rewards, used to guide behavior[207,232].

There is therefore a remarkable concordance between the enriched CNS regions highlighted by the DEPICT analyses and the circuit in charge of reward processing, behavioral control, and decision-making defined by decades of behavioral and physiological studies in human and non-human primates[210]. It is however difficult to claim that this circuit is *specifically* tagged by the DEPICT tissue enrichment analysis with the GTEx data, given that the GTEx data only cover a fraction of all brain regions, mostly restricted to brain regions overlapping with the cortical-basal ganglia circuit. Although these results, in line with a large body of neuroscientific studies, suggest a role of the cortical-basal ganglia circuit in the risk-tolerance phenotype, their inferential value should therefore be considered with caution.

##### *12.7.4 A likely role for glutamate and GABA neurotransmitters in the modulation of risk tolerance*

In addition to neuroanatomy, much speculation has revolved around the role of specific hormones and neurotransmitter systems in modulating general risk tolerance. Our review of the literature attempting to link risk tolerance to biological pathways (**Supplementary Information section 11**) identified five main biological pathways that have been tested by this literature: the steroid hormone cortisol, the monoamines dopamine and serotonin, and the steroid sex hormones estrogen and testosterone. Consistent with the lack of enrichment reported in **Supplementary Information section 11** for genes associated with these five pathways, neither the Gene Network analysis of the MAGMA genes nor the DEPICT gene set prioritization analysis point to these pathways. (As mentioned above, however, we note that our results are consistent with a role of the cortical-basal ganglia circuit in modulating risk tolerance, and that dopamine is presumed to play an important role in that circuit.)

On the other hand, both the Gene Network functional analysis of the MAGMA genes and the DEPICT gene set prioritization analysis point to a role for glutamate and GABA neurotransmitters, which are the main excitatory and inhibitory neurotransmitters in the brain, respectively[233]. They are colloquially referred to as the “workhorses” of the brain, as they are the brain’s principal neurotransmitters. At the time of writing, no other published large-scale GWAS of cognition, personality, or neuropsychiatric phenotypes has pointed to clear roles for both glutamate and GABA, although we note that glutamate neurotransmission has been implicated in recent GWAS of schizophrenia – by recent analyses of common SNPs[69] and rare and *de novo* mutations[234,235] – and of major depression[181]. That our results, unlike those of other well-powered GWAS, point to both these neurotransmitters suggests that the relative balance between excitatory and inhibitory neurotransmission may be a relatively strong contributor to variation in risk tolerance across individuals.

However, we do note that none of the four glutamate and GABA gene sets we tested in an *ex post* MAGMA competitive gene-set analysis were significant (**Supplementary Information section 12.5**). We are unsure why this might be the case, but we note that this analysis had some likely limitations. First, its statistical power barely increases with increasing sample sizes[163]. Second, a MAGMA competitive gene-set analysis might not be the optimal approach for discovery of subtle but phenotypically important genetic effects driven by regulatory regions, as this analysis effectively calculates a weighted average of GWAS $-\log_{10}(P)$ values across each gene in the gene set (where the average is weighted to correct for LD between SNPs). Hence, important regulatory effects of single SNPs may be overshadowed by the high average $P$ values of other SNPs in the gene sets. A third possible explanation is that only specific glutamate and/or GABA genes or pathways might be relevant in general risk tolerance, and that our merging of those pathways into major glutamate and GABA supersets decreased our statistical power for discovery of relevant effects. (In the MAGMA gene-based analysis, only 12 of the 354 unique GABA/glutamate genes included in the four gene sets tested here were significant at Bonferroni-corrected $P < 0.05$.) Hence, the lack of significance of the four glutamate and GABA gene sets we tested in the *ex post* MAGMA competitive gene-set analysis does not nullify the evidence from our Gene Network and the DEPICT analyses that glutamatergic and GABAergic neurotransmission contributes to variation in risk tolerance across individuals.

###### *12.7.5 A possible link between the immune system and risk tolerance*

We now briefly discuss the LDSC immune tissue enrichment. We noted that, to the best of our knowledge, general risk tolerance is the first behavioral trait to show immune tissue enrichment in LDSC partitioned analysis according to tissue type, and that this enrichment was independent of SNPs in the MHC region. We also noted that none of our other analyses show immune enrichment—the Andersson[236] "FANTOM5" enhancers (which are indicative of immune activity[74]) were not a significant genomic annotation in the LDSC partitioned analyses according to genomic annotation, and the DEPICT and Gene Network analyses of MAGMA genes also did not show immune pathways (although it is noteworthy that these bioannotation analyses excluded the MHC region). However, we do find GWAS hits in the MHC region, and several genes in this region came up in the MAGMA and GTEx lookup analyses. That is, 29 of the 285 genes significant in MAGMA and 47 of the 188 GTEx eQTL lookups were in (or around the borders of) the MHC region (while SMR excluded SNPs in the MHC region). While a role for the immune system in neuropsychiatric traits and related behavior is feasible given previous enrichment of immune pathways for psychiatric traits[69,180,181,235], we are unsure how to interpret a role for the immune system in general risk tolerance, given that our results do not unequivocally point in this direction. Researchers have previously theorized a role for a "behavioral immune system"[237,238] in human behavior. This is seen to be complementary to the "real" immune system—that is, humans are said to employ risk averse behaviors to minimize their risk of infectious disease. An extension of this theory could be that individuals who are genetically prone to "weaker" immune systems might be more risk averse than others, but this is extremely speculative. Before accepting such conclusions, we encourage future bioannotation work to uncover which specific immune pathways might be involved in general risk tolerance.

##### *12.7.6 The* **CADM2** *gene and risk tolerance*

Finally, we briefly discuss the top locus from our discovery GWAS of general risk tolerance. The top lead SNP that marks this locus is located in the intronic region of *CADM2*, and is our most significant general-risk-tolerance lead SNP by far, with a *P* value that is fifteen orders of magnitude smaller than the second lead SNP's *P* value ($P = 2.1 \times 10^{-40}$ vs. $P = 7.6 \times 10^{-25}$). The *CADM2* locus was previously identified by a study using the relatively small sample size of the first release of UK Biobank data[50], and all of our six supplementary GWAS phenotypes identified lead SNPs within or near *CADM2* (**Supplementary Information section 3**). *CADM2* was by far the most significant MAGMA gene, and was also prioritized by DEPICT at FDR < 0.01. However, we do note that only *CADM2*'s antisense RNA partner *CADM2-AS1* was a significant cis-eQTL gene in our GTEx lookup tissues, while *CADM2* failed the HEIDI-test for heterogeneity in the SMR analysis. This suggests that *CADM2* may not be the causal gene in the locus, or that its effect on general risk tolerance is not mediated by *CADM2* expression. However, we do stress that the brain eQTL and SMR analyses are strongly underpowered due to the relatively small samples sizes for brain tissues, and we can thus not properly study *CADM2* expression in relevant brain tissues.

According to GTEx, *CADM2* is overexpressed in the brain, and in particular in the frontal cortex. *CADM2* is a large gene (spanning more than 1 Mb) located in a long-range LD region, and has been associated with a myriad of behavioral phenotypes in previous GWAS (**Supplementary Information section 3**). *CADM2* encodes a member of the synaptic cell adhesion molecule family. Molecules in this family have been found to be crucial for synapse formation[56] and plasticity[57], and have been associated with autism[239] and impaired social behavior[240]. However, *CADM2* itself has mainly been studied for its potential role in tumor progression[241]; its specific role in the brain is unclear. We therefore propose *CADM2* as an interesting candidate gene for future follow up work.

### 13 Additional information

#### 13.1 Author contributions

Jonathan Beauchamp, Philipp Koellinger, and Daniel Benjamin designed and oversaw the study. Jonathan Beauchamp closely supervised the analyses and led the writing of the manuscript.

Richard Karlsson Linnér was the lead analyst, responsible for quality control, the meta-analyses, summarizing the overlap across the results of the various GWAS, and the SNP heritability analyses. Edward Kong conducted the population stratification, replication, and proxy-phenotype analyses.

Robbee Wedow led the genetic correlation analyses, and Mark Fontana contributed to those analyses. Pietro Biroli led the polygenic score prediction analyses, and Richard Karlsson Linnér, Edward Kong, Robbee Wedow, Abdel Abdellaoui, Ronald de Vlaming, Mark Fontana, and Michel G Nivard, contributed to those analyses. Pietro Biroli and Christian Zünd conducted the MTAG analyses.

The bioinformatics analyses were led by Fleur Meddens. Analysts who assisted Fleur include: Jacob Gratten and Maciej Trzaskowski (SMR), Anke Hammerschlag (MAGMA), Gerardus Meddens (Gene Network) and Pascal Timshel (DEPICT). Maël Lebreton conducted the review of the literature attempting to link risk tolerance to biological pathways.

Richard Karlsson Linnér prepared the majority of figures; Edward Kong and Stephen Tino also prepared some figures. Aysu Okbay, Cornelius A Rietveld, and Stephen Tino helped with several additional analyses.

David Cesarini, Jacob Gratten, and James Lee provided helpful advice and feedback on various aspects of the study design.

All authors contributed to and critically reviewed the manuscript. Pietro Biroli, Richard Karlsson Linnér, Edward Kong, Fleur Meddens, and Robbee Wedow made especially major contributions to the writing and editing.

#### 13.2 Cohort-level contributions

| Cohort | Author | Study design & mgmt. | Data collec-tion | Geno-typing | Geno-type prep. | Pheno-type prep. | Data analysis |
|---|---|---|---|---|---|---|---|
| **23andMe** | Pierre Fontanillas | | | | | | X |
| **23andMe** | Aaron Kleinman | | | | | | X |
| **23andMe** | Adam Auton | X | | | | | |
| **23andMe** | David A. Hinds | X | | | | | |
| **Add Health** | Jason D Boardman | | X | | | | |
| **Add Health** | Kathleen Mullan Harris | | X | | | | |
| **Add Health** | Matthew B McQueen | | X | | | | |
| **Army STARRS** | Murray B Stein | X | X | | | | |
| **Army STARRS** | Chia-Yen Chen | | | X | X | | X |
| **Army STARRS** | Sonia Jain | | | | | X | |
| **Army STARRS** | Robert J Ursano | X | X | | | | |
| **BASE-II** | Lars Bertram | X | X | X | X | | |
| **BASE-II** | Christina M Lill | | X | X | | | X |
| **BASE-II** | Peter Eibich | | X | | | X | X |
| **BASE-II** | Gert G Wagner | | X | | | X | |
| **HRS** | Ronald de Vlaming | | | | | X | X |
| **NFBC1966** | Ville Karhunen | | | | | X | X |
| **NFBC1966** | Andrew Conlin | X | X | | | X | |
| **NFBC1966** | Minna Männikkö | X | X | | | X | |
| **NFBC1966** | Rauli Svento | X | X | | | X | |
| **NTR** | Michel G Nivard | | | | | X | X |
| **NTR** | Abdel Abdellaoui | | | | X | | X |
| **NTR** | Conor V Dolan | X | | | | | |
| **NTR** | Dorret I Boomsma | X | X | | X | | |
| **RS** | Frank J van Rooij | | X | | | X | X |
| **RS** | Albert Hofman | X | | | | | |
| **RS** | Mohammad A Ikram | X | X | | | | |
| **RS** | Andre G Uitterlinden | | | X | X | | |
| **RS** | Henning Tiemeier | X | X | | | X | |
| **RS** | Patrick JF Groenen | X | | | | | X |
| **RS** | A Roy Thurik | X | | | | | X |
| **RS** | Cornelius A Rietveld | | | | X | X | X |
| **STR** | Cornelius A Rietveld | | | | X | X | X |
| **STR** | Patrik KE Magnusson | X | X | X | X | | |
| **STR** | Robert Karlsson | | | X | X | | |
| **STR** | Magnus Johannesson | X | X | | | X | |
| **STR** | David Cesarini | X | X | | | X | |
| **UKB** | Mark Alan Fontana | | | | X | X | X |

| | | | | | | | |
|---|---|---|---|---|---|---|---|
| **UKB** | Richard Karlsson Linnér | | | | X | X | X |
| **UKB** | Robbee Wedow | | | | | X | |
| **UKB** | Cornelius A Rietveld | | | | X | X | X |
| **UKHLS** | Yanchun Bao | | | | X | | X |
| **UKHLS** | Melissa C Smart | | | | X | X | |
| **UKHLS** | Jon White | | | X | | | |
| **UKHLS** | Meena Kumari | X | | | | | |
| **Viking** | Paul RHJ Timmers | | | | | | X |
| **Viking** | Peter K Joshi | | | | | | X |
| **Viking** | David W Clark | | | | | X | X |
| **Viking** | James F Wilson | X | X | | | | |
| **ZMI** | Ernst Fehr | X | X | | | X | |
| **ZMI** | Daniel Schunk | X | X | X | | X | |
| **ZMI** | Arcadi Navarro | X | | | X | | |
| **ZMI** | Laura Buzdugan | | | | | X | X |
| **ZMI** | Urs Fischbacher | | X | | | | |
| **ZMI** | Gregor Hasler | X | | | | | |
| **ZMI** | Gerard Muntané | | | | X | | X |
| **ZMI** | Klaus M Schmidt | | X | | | | |
| **ZMI** | Matthias Sutter | | X | | | | |
| **ZMI** | Carlos Morcillo-Suarez | | | | | | X |

#### 13.3 Additional acknowledgments

##### *13.3.1 Individual acknowledgements*

Tõnu Esko is supported by the Estonian Research Council Grant IUT20-60 and PUT1660, EU H2020 grant 692145, and European Union through the European Regional Development Fund (Project No. 2014-2020.4.01.15-0012) GENTRANSMED.

Juan R Gonzalez is supported by Spanish Ministry of Economy and Competitiveness (MTM2015-68140-R).

Jacob Gratten was supported by a Level 2 Career Development Fellowship from the Australian National Health and Medical Research Council (1127440), and by National Health and Medical Research Council (NHMRC) grants 1087889 and 1103418.

James MacKillop's contributions were supported by the Peter Boris Chair in Addictions Research.

Ronald C Kessler, in the past three years, received support for his epidemiological studies from Sanofi Aventis; was a consultant for Johnson & Johnson Wellness and Prevention, Sage Pharmaceuticals, Shire, Takeda; and served on an advisory board for the Johnson & Johnson Services Inc. Lake Nona Life Project. Kessler is a co-owner of DataStat, Inc., a market research firm that carries out healthcare research.

Philipp D Koellinger acknowledges financial support from the European Research Council (consolidator grant 647648).

Michel G Nivard is supported by Royal Netherlands Academy of Science Professor Award (PAH/6635) to Dorret I. Boomsma. Michel G Nivard is funded by the BBMRI-NL, a research infrastructure financed by the Netherlands Organization for Scientific Research (NWO project 184.021.007).

Tune H Pers is supported by the Novo Nordisk Foundation and the Lundbeck Foundation.

Danielle Posthuma is supported by the Netherlands Organization for Scientific Research: NWO Brain & Cognition 433-09-228. Statistical analyses were carried out on the Genetic Cluster Computer (http://www.geneticcluster.org) hosted by SURFsara and financially supported by the Netherlands Organization for Scientific Research (NWO 480-05-003, PI: Posthuma) along with a supplement from the Dutch Brain Foundation and the VU University Amsterdam.

Niels Rietveld gratefully acknowledges funding from the Netherlands Organization for Scientific Research (NWO Veni grant 016.165.004).

Robbee Wedow was generously supported by the National Science Foundation’s Graduate Research Fellowship Program (DGE 1144083). Any opinion, findings, and conclusions or recommendations expressed in this material are those of the authors(s) and do not necessarily reflect the views of the National Science Foundation.

Jian Yang is supported by Sylvia & Charles Viertel Charitable Foundation; National Health and Medical Research Council (NHMRC) 1107258 and 1113400.

##### *13.3.2 Extended acknowledgements*

**23andMe, Inc.**

We gratefully acknowledge the contributions of members of 23andMe's Research Team, whose names are listed below:

Michelle Agee, Babak Alipanahi, Adam Auton, Robert K. Bell, Katarzyna Bryc, Sarah L. Elson, Pierre Fontanillas, Nicholas A. Furlotte, David A. Hinds, Karen E. Huber, Aaron Kleinman, Nadia K. Litterman, Jennifer C. McCreight, Matthew H. McIntyre, Joanna L. Mountain, Carrie A.M. Northover, Steven J. Pitts, J. Fah Sathirapongsasuti, Olga V. Sazonova, Janie F. Shelton, Suyash Shringarpure, Chao Tian, Joyce Y. Tung, Vladimir Vacic, and Catherine H. Wilson.

##### *13.3.3 Cohort acknowledgements*

**23andMe, Inc.** – We would like to thank the research participants and employees of 23andMe for making this work possible. 23andMe research participants provided informed consent and participated in the research online, under a protocol approved by the AAHRPP-accredited institutional review board, Ethical and Independent Review Services (E&I Review). Participant data are shared according to community standards that have been developed to protect against breaches of privacy. Currently, these standards allow for the sharing of summary statistics for at most 10,000 SNPs. The full set of summary statistics can be made available to qualified investigators who enter into an agreement with 23andMe that protects participant confidentiality. Interested investigators should for more information.

**Add Health** – The National Longitudinal Study of Adolescent to Adult Health (Add Health) is supported by grant P01 HD031921 to Kathleen Mullan Harris from the Eunice Kennedy Shriver National Institute of Child Health and Human Development (NICHD), with cooperative funding from 23 other federal agencies and foundations. Add Health GWAS data were funded by NICHD grants to Harris (R01 HD073342) and to Harris, Boardman, and McQueen (R01 HD060726). For information about access to the data from this study, contact.

**Army STARRS** – The Army Study to Assess Risk and Resilience in Servicemembers (Army STARRS) acknowledges grant U01MH087981. Data access requests may be sent to.

**BASE-II (Berlin Ageing Study II)** – BASE -II has been funded by the German Federal Ministry of Education and Research (BMBF) and has been formally divided into four subprojects: "Psychology & Project Coordination and Database" (Max Planck Institute for Human Development [MPIB], grant number 16SV5837), "Survey Methods and Social Science" (German Institute for Economic Research and Socioeconomic Panel [SOEP/DIW], grant number 16 SV5537), Medicine and Geriatrics (Charité – Universitätsmedizin, Berlin [Charité], grant number 16SV5536K), and "Molecular Genetics" (Max Planck Institute for Molecular Genetics [MPIMG], now coordinated from University of Lübeck [ULBC], grant number 16SV5538). External scientists can apply to the Steering Committee of BASE -II for data access. Although the data are available for other parties are scientific data and not personal contact data, the scientific data are subject to a security level as if they were personal data to ensure that the BASE -II Steering Committee sufficiently protects the large volume of data collected from each

BASE -II participant. All existing variables are documented in a handbook. See BASE-II project website for more details: https://www.base2.mpg.de/en).

**HRS (Health and Retirement Study)** – HRS is supported by the National Institute on Aging (NIA U01AG009740). The genotyping was funded separately by the National Institute on Aging (RC2 AG036495, RC4 AG039029). Genotyping was conducted by the NIH Center for Inherited Disease Research (CIDR) at Johns Hopkins University. Genotyping quality control and final preparation of the data were performed by the Genetics Coordinating Center at the University of Washington. Genotype data can be accessed via the database of Genotypes and Phenotypes (dbGaP, http://www.ncbi.nlm.nih.gov/gap, accession number phs000428.v1.p1). Researchers who wish to link genetic data with other HRS measures that are not in dbGaP, such as educational attainment, must apply for access from HRS. See the HRS website (http://hrsonline.isr.umich.edu/gwas) for details.

**NFBC1966** – We thank the late professor Paula Rantakallio (launch of NFBC1966), the participants in the NFBC1966 46y study and the NFBC Project Center. The NFBC1966 has received financial support from University of Oulu Grant no. 24000692, Oulu University Hospital Grant no. 24301140, ERDF European Regional Development Fund Grant no. 539/2010 A31592. Andrew Conlin thanks OP Group Research Foundation and Finnish Cultural Foundation North Ostrobothnia Regional Fund for scholarships.

**NTR (Netherlands Twin Register)** – We are grateful to all NTR twins, and their family members who participate in the NTR research. Abdel Abdellaoui was supported by the National Institutes of Health (NIH, R37 AG033590-08). Michel G Nivard is supported by Royal Netherlands Academy of Science Professor Award (PAH/6635) to Dorret I Boomsma and by BBMRI-NL, a research infrastructure financed by the Netherlands organization for Scientific Research (NWO project 184.021.007). The NTR gratefully acknowledges support from “Genetic influences on stability and change in psychopathology from childhood to young adulthood” (ZonMW 912-10-020) and “Genetics of Mental Illness” (ERC-230374). The Netherlands Organization for Scientific Research (NWO) and MagW/ZonMW grants Middelgroot-911-09-032, NWO Groot (480-15-001/674): Netherlands Twin Registry Repository; Biobanking and Biomolecular Resources Research Infrastructure (BBMRI–NL,184.021.007), VU University’s Institute for Health and Care Research (EMGO+) and Neuroscience Campus Amsterdam (NCA). Part of the genotyping was funded by the Genetic Association Information Network (GAIN) of the Foundation for the National Institutes of Health, Rutgers University Cell and DNA Repository (NIMH U24 MH068457-06), the Avera Institute, Sioux Falls, South Dakota (USA) and the National Institutes of Health (NIH R01 HD042157-01A1, MH081802, Grand Opportunity grants 1RC2 MH089951 and 1RC2 MH089995). Data access requests can be sent to & (please make sure you mention the project and the collaborators Prof. Dorret Boomsma, Prof. Conor Dolan, Dr. Abdel Abdellaoui, and Dr. Michel Nivard).

**RS (Rotterdam Study)** – The generation and management of GWAS genotype data for the Rotterdam Study is supported by the Netherlands Organisation of Scientific Research NWO Investments (nr. 175.010.2005.011, 911-03-012). This study is funded by the Research Institute for Diseases in the Elderly (014-93-015; RIDE2), the Netherlands Genomics Initiative (NGI)/Netherlands Organisation for Scientific Research (NWO) project nr. 050-060-810. We thank Pascal Arp, Mila Jhamai, Marijn Verkerk, Lizbeth Herrera and Marjolein Peters for their help in creating the GWAS database, and Karol Estrada and Maksim V. Struchalin for their support in creation and analysis of imputed data. The Rotterdam Study is funded by Erasmus Medical Center and Erasmus University, Rotterdam, Netherlands Organization for the Health Research and Development (ZonMw), the Research Institute for Diseases in the Elderly (RIDE), the Ministry of Education, Culture and Science, the Ministry for Health, Welfare and Sports, the European Commission (DG XII), and the Municipality of Rotterdam. The authors are grateful to the study participants, the staff of the Rotterdam Study and the participating general practitioners and pharmacists.

**STR (Swedish Twin Registry) –** The Jan Wallander and Tom Hedelius Foundation (P2015-0001:1), the Ragnar Söderberg Foundation (E9/11), the Swedish Research Council (421-2013-1061). SALTY constitutes a sub-cohort of the Swedish Twin Registry which receives financial support by Karolinska Institutet. Researchers interested in using STR data must obtain approval from the Swedish Ethical Review Board and from the Steering Committee of the Swedish Twin Registry. Researchers using the data are required to follow the terms of an Assistance Agreement containing a number of clauses designed to ensure protection of privacy and compliance with relevant laws. For further information, contact Patrik Magnusson.

**UKB (UK Biobank) –** This research has been conducted using the UK Biobank Resource. Informed consent was obtained from UK Biobank subjects.

**UKHLS (The UK Household Longitudinal Study) –** These data are from Understanding Society: The UK Household Longitudinal Study (UKHLS), which is led by the Institute for Social and Economic Research at the University of Essex and funded by the Economic and Social Research Council. The data were collected by NatCen and the genome-wide scan data were analysed by the Wellcome Trust Sanger Institute. Information on how to access the data can be found on the Understanding Society website https://www.understandingsociety.ac.uk/ or METADAC website http://www.metadac.ac.uk/

**Viking** - The Viking Health Study – Shetland (VIKING) was supported by the MRC Human Genetics Unit quinquennial programme grant "QTL in Health and Disease." DNA extractions were performed at the Wellcome Trust Clinical Research Facility in Edinburgh. We would like to acknowledge the invaluable contributions of the research nurses in Shetland, the administrative team in Edinburgh and the people of Shetland. Data access requests may be sent to.

**ZMI (Zurich-Munich-Innsbruck cohort)** – EF gratefully acknowledges support from the European Research Council grant number 295642 (Foundations of Economic Preferences). DS gratefully acknowledges support by the German National Science Foundation (DFG) under the grant SCHU 2828/2-1. AN gratefully acknowledges support by the Ministerio de Ciencia e Innovación, Spain (BFU2012-38236 and BFU2015-68649 to AN) by the Spanish National Institute of Bioinfomatics of the Instituto de Salud Carlos III (PT13/0001/0026) and by FEDER (Fondo Europeo de Desarrollo Regional)/FSE (Fondo Social Europeo). GH gratefully acknowledges support by the University of Bern. KS gratefully acknowledges support by the Excellence Initiative of the German government. Support in data collection and preparation by Daniela Glätzle-Rützler, Simon Czermak, Christina Strassmair, René Cyranek, Angel Carreño and Xavier Farre and Adrian Bruhin is gratefully acknowledged.

### 14 References

1. Barsky, R. B., Juster, F. T., Kimball, M. S. & Shapiro, M. D. Preference parameters and behavioral heterogeneity: An experimental approach in the Health and Retirement Study. *Q. J. Econ.* **112,** 537–579 (1997).
2. Beauchamp, J. P., Cesarini, D. & Johannesson, M. The psychometric and empirical properties of measures of risk preferences. *J. Risk Uncertain.* **54,** 203–237 (2017).
3. Becker, A., Deckers, T., Dohmen, T., Falk, A. & Kosse, F. The relationship between economic preferences and psychological personality measures. *Annu. Rev. Econom.* **4,** 453–478 (2012).
4. Dohmen, T. *et al.* Individual risk attitudes: Measurement, determinants, and behavioral consequences. *J. Eur. Econ. Assoc.* **9,** 522–550 (2011).
5. Falk, A., Dohmen, T., Falk, A. & Huffman, D. The Nature and Predictive Power of Preferences: Global Evidence. *IZA Discussion Paper Series* (2015).
6. Frey, R., Pedroni, A., Mata, R., Rieskamp, J. & Hertwig, R. Risk preference shares the psychometric structure of major psychological traits. *Sci. Adv.* **3,** e1701381 (2017).
7. Cesarini, D., Dawes, C. T., Johannesson, M., Lichtenstein, P. & Wallace, B. Genetic variation in preferences for giving and risk taking. *Q. J. Econ.* **124,** 809–842 (2009).
8. Zyphur, M. J., Narayanan, J., Arvey, R. D. & Alexander, G. J. The genetics of economic risk preferences. *J. Behav. Decis. Mak.* **22,** 367–377 (2009).
9. Harden, K. P. *et al.* Beyond dual systems: a genetically-informed, latent factor model of behavioral and self-report measures related to adolescent risk-taking. *Dev. Cogn. Neurosci.* **25,** 221–234 (2017).
10. Benjamin, D. J. *et al.* The genetic architecture of economic and political preferences. *Proc. Natl. Acad. Sci.* **109,** 8026–8031 (2012).
11. Dreber, A. *et al.* The 7R polymorphism in the dopamine receptor D4 gene (DRD4) is associated with financial risk taking in men. *Evol. Hum. Behav.* **30,** 85–92 (2009).
12. Carpenter, J. P., Garcia, J. R. & Lum, J. K. Dopamine receptor genes predict risk preferences, time preferences, and related economic choices. *J. Risk Uncertain.* **42,** 233–261 (2011).
13. Frydman, C., Camerer, C., Bossaerts, P. & Rangel, A. MAOA-L carriers are better at making optimal financial decisions under risk. *Proc. R. Soc. B Biol. Sci.* **278,** 2053–2059 (2010).
14. He, Q. *et al.* Serotonin transporter gene-linked polymorphic region (5-HTTLPR) influences decision making under ambiguity and risk in a large Chinese sample. *Neuropharmacology* **59,** 518–26 (2010).
15. Chabris, C. F., Lee, J. J., Cesarini, D., Benjamin, D. J. & Laibson, D. I. The fourth law of behavior genetics. *Curr. Dir. Psychol. Sci.* **24,** 304–312 (2015).
16. Okbay, A. *et al.* Genome-wide association study identifies 74 loci associated with educational attainment. *Nature* **533,** 539–542 (2016).

17. Harrati, A. Characterizing the genetic influences on risk aversion. *Biodemography Soc. Biol.* **60,** 185–198 (2014).

18. Okbay, A. *et al.* Genetic variants associated with subjective well-being, depressive symptoms, and neuroticism identified through genome-wide analyses. *Nat. Genet.* **48,** 624–633 (2016).

19. Gelman, A. & Carlin, J. Beyond power calculations: Assessing type S (sign) and type M (magnitude) errors. *Perspect. Psychol. Sci.* **9,** 641–651 (2014).

20. Hewitt, J. K. Editorial policy on candidate gene association and candidate gene-by-environment interaction studies of complex traits. *Behav. Genet.* **42,** 1–2 (2012).

21. Ioannidis, J. P. A. Why most published research findings are false. *PLoS Med.* **2,** e124 (2005).

22. Byrnes, J. P., Miller, D. C. & Schafer, W. D. Gender differences in risk taking: a meta-analysis. *Psychol. Bull.* **125,** 367–383 (1999).

23. Croson, R. & Gneezy, U. Gender Differences in Preferences. *J. Econ. Lit.* **47,** 448–474 (2009).

24. Bulik-Sullivan, B. K. *et al.* An atlas of genetic correlations across human diseases and traits. *Nat. Genet.* **47,** 1236–1241 (2015).

25. Holt, C. A. & Laury, S. K. Risk aversion and incentive effects. *Am. Econ. Rev.* **92,** 1644–1655 (2002).

26. Anderson, L. R. & Mellor, J. M. Are risk preferences stable? Comparing an experimental measure with a validated survey-based measure. *J. Risk Uncertain.* **39,** 137–160 (2009).

27. Levenson, M. R. Risk taking and personality. *J. Pers. Soc. Psychol.* **58,** 1073–80 (1990).

28. Weber, M., Weber, E. U. & Nosic, A. Who takes risks when and why: Determinants of changes in investor risk taking. *Rev. Financ.* **17,** 847–883 (2013).

29. The Tobacco and Genetics Consortium. Genome-wide meta-analyses identify multiple loci associated with smoking behavior. *Nat. Genet.* **42,** 441–7 (2010).

30. SSGAC. Pre-registered Analysis Plan - GWAS Risk tolerance. *Open Science Framework* (2016). Available at: https://osf.io/cjx9m/.

31. Marchini, J. L. *et al.* Genotype imputation and genetic association studies of UK Biobank. UK Biobank (2015).

32. Bycroft, C. *et al.* Genome-wide genetic data on ~500,000 UK Biobank participants. Pre-print at *bioRxiv* (2017). doi:http://dx.doi.org/10.1101/166298

33. McCarthy, S. *et al.* A reference panel of 64,976 haplotypes for genotype imputation. *Nat. Genet.* **48,** 1279–1283 (2016).

34. Loh, P.-R. *et al.* Efficient Bayesian mixed-model analysis increases association power in large cohorts. *Nat. Genet.* **47,** 284–290 (2015).

35. Tan, A., Abecasis, G. R. & Kang, H. M. Unified representation of genetic variants. *Bioinformatics* **31,** 2202–2204 (2015).

36. Huang, J. *et al.* Improved imputation of low-frequency and rare variants using the UK10K haplotype reference panel. *Nat. Commun.* **6,** 8111 (2015).

37. Purcell, S. *et al.* PLINK: A tool set for whole-genome association and population-based linkage analyses. *Am. J. Hum. Genet.* **81,** 559–575 (2007).

38. Wain, L. V. *et al.* Novel insights into the genetics of smoking behaviour, lung function, and chronic obstructive pulmonary disease (UK BiLEVE): A genetic association study in UK Biobank. *Lancet Respir. Med.* **3,** 769–781 (2015).

39. UK Biobank. Genotyping and Quality Control of UK Biobank, a Large-Scale, Extensively Phenotyped Prospective Resource: Information for Researchers. Interim Data Release. (2015).

40. Winkler, T. W. *et al.* Quality control and conduct of genome-wide association meta-analyses. *Nat. Protoc.* **9,** 1192–1212 (2014).

41. Willer, C. J., Li, Y. & Abecasis, G. R. METAL: Fast and efficient meta-analysis of genomewide association scans. *Bioinformatics* **26,** 2190–2191 (2010).

42. Bulik-Sullivan, B. K. *et al.* LD Score regression distinguishes confounding from polygenicity in genome-wide association studies. *Nat. Genet.* **47,** 291–295 (2015).

43. Rietveld, C. A. *et al.* GWAS of 126,559 individuals identifies genetic variants associated with educational attainment. *Science* **340,** 1467–1471 (2013).

44. Price, A. L. *et al.* Long-range LD can confound genome scans in admixed populations. *Am. J. Hum. Genet.* **83,** 132–139 (2008).

45. Cáceres, A. & González, J. R. Following the footprints of polymorphic inversions on SNP data: from detection to association tests. *Nucleic Acids Res.* **43,** e53 (2015).

46. Cáceres, A., Sindi, S. S., Raphael, B. J., Cáceres, M. & González, J. R. Identification of polymorphic inversions from genotypes. *BMC Bioinformatics* **13,** 28 (2012).

47. Ma, J. & Amos, C. I. Investigation of inversion polymorphisms in the human genome using principal components analysis. *PLoS One* **7,** e40224 (2012).

48. Sudmant, P. H. *et al.* An integrated map of structural variation in 2,504 human genomes. *Nature* **526,** 75–81 (2015).

49. Welter, D. *et al.* The NHGRI GWAS Catalog, a curated resource of SNP-trait associations. *Nucleic Acids Res.* **42,** D1001-1006 (2014).

50. Day, F. R. *et al.* Physical and neurobehavioral determinants of reproductive onset and success. *Nat. Genet.* **48,** 617–623 (2016).

51. Strawbridge, R. J. *et al.* Genome-wide analysis of risk-taking behaviour and cross-disorder genetic correlations in 116 255 individuals from the UK Biobank cohort. Pre-print at *bioRxiv* (2017). doi:http://dx.doi.org/10.1101/177014

52. Boutwell, B. *et al.* Replication and characterization of CADM2 and MSRA genes on human behavior. *Heliyon* **3,** e00349 (2017).

53. Demontis, D. *et al.* Discovery of the first genome-wide significant risk loci for ADHD. Pre-print at *bioRxiv* (2017). doi:10.1101/145581

54. The Autism Spectrum Disorders Working Group of The Psychiatric Genomics Consortium. Meta-analysis of GWAS of over 16,000 individuals with autism spectrum disorder highlights a novel locus at 10q24.32 and a significant overlap with schizophrenia. *Mol. Autism* **8,** 21 (2017).

55. Sherry, S. T. *et al.* dbSNP: the NCBI database of genetic variation. *Nucleic Acids Res.* **29,** 308–11 (2001).

56. Biederer, T. *et al.* SynCAM, a synaptic adhesion molecule that drives synapse assembly. *Science* **297,** 1525–31 (2002).

57. Robbins, E. M. *et al.* SynCAM 1 Adhesion Dynamically Regulates Synapse Number and Impacts Plasticity and Learning. *Neuron* **68,** 894–906 (2010).

58. Lonsdale, J. *et al.* The Genotype-Tissue Expression (GTEx) project. *Nat. Genet.* **45,** 580–5 (2013).

59. Amiel, J. *et al.* Mutations in TCF4, Encoding a Class I Basic Helix-Loop-Helix Transcription Factor, Are Responsible for Pitt-Hopkins Syndrome, a Severe Epileptic Encephalopathy Associated with Autonomic Dysfunction. *Am. J. Hum. Genet.* **80,** 988–993 (2007).

60. Zweier, C. *et al.* Haploinsufficiency of TCF4 causes syndromal mental retardation with intermittent hyperventilation (Pitt-Hopkins syndrome). *Am. J. Hum. Genet.* **80,** 994–1001 (2007).

61. De Pontual, L. *et al.* Mutational, functional, and expression studies of the TCF4 gene in Pitt-Hopkins syndrome. *Hum. Mutat.* **30,** 669–676 (2009).

62. Blake, D. J. *et al.* TCF4, Schizophrenia, and Pitt-Hopkins Syndrome. *Schizophr. Bull.* **36,** 443–447 (2010).

63. Trowsdale, J. & Knight, J. C. Major Histocompatibility Complex Genomics and Human Disease. *Annu. Rev. Genomics Hum. Genet.* **14,** 301–323 (2013).

64. Murphy, K. & Weaver, C. *Janeway's Immunobiology*. (Garland Science, Taylor & Francis Group, LLC, 2017).

65. O'Brien, R. & Wong, P. Amyloid Precursor Protein Processing and Alzheimer's Disease. *Annu. Rev. Neurosci.* **34,** 185–204 (2011).

66. Berisa, T. & Pickrell, J. K. Approximately independent linkage disequilibrium blocks in human populations. *Bioinformatics* **32,** 283–285 (2016).

67. Nachar, N. The Mann-Whitney U: A test for assessing whether two independent samples come from the same distribution. *Quant. Methods Psychol.* **4,** 13–20 (2008).

68. Yang, J. *et al.* Genomic inflation factors under polygenic inheritance. *Eur. J. Hum. Genet.* **19,** 807–812 (2011).

69. Ripke, S. *et al.* Biological insights from 108 schizophrenia-associated genetic loci. *Nature* **511,** 421–427 (2014).

70. Pickrell, J. K. *et al.* Detection and interpretation of shared genetic influences on 40 human traits. *Nat. Genet.* **48,** 709–718 (2015).

71. Thorgeirsson, T. E. *et al.* Sequence variants at CHRNB3-CHRNB6 and CYP2A6 affect smoking behavior. *Nat. Genet.* **42,** 448–453 (2011).

72. Rose, R. J., Broms, U., Korhonen, T., Dick, D. & Kaprio, J. Genetics of Smoking Behavior. In Y. Kim (Ed.), *Handbook of Behavior Genetics* (pp. 411–432). New York: Springer.

73. Price, A. L. *et al.* Principal components analysis corrects for stratification in genome-wide association studies. *Nat. Genet.* **38,** 904–909 (2006).

74. Finucane, H. K. *et al.* Partitioning heritability by functional category using GWAS summary statistics. *Nat. Genet.* **47,** 1228–1235 (2015).

75. Yang, J., Lee, S. H., Goddard, M. E. & Visscher, P. M. GCTA: a tool for genome-wide complex trait analysis. *Am. J. Hum. Genet.* **88,** 76–82 (2011).

76. Shi, H., Kichaev, G. & Pasaniuc, B. Contrasting the genetic architecture of 30 complex traits from summary association data. *Am. J. Hum. Genet.* **99,** 139–153 (2016).

77. The 1000 Genomes Project Consortium *et al.* An integrated map of genetic variation from 1,092 human genomes. *Nature* **491,** 56–65 (2012).

78. Stringer, S. *et al.* Genome-wide association study of lifetime cannabis use based on a large meta-analytic sample of 32,330 subjects from the International Cannabis Consortium. *Transl. Psychiatry* **6,** e769 (2016).

79. van der Loos, M. J. H. M. *et al.* The Molecular Genetic Architecture of Self-Employment. *PLoS One* **8,** e60542 (2013).

80. Rietveld, C. A. *et al.* Common genetic variants associated with cognitive performance identified using the proxy-phenotype method. *Proc. Natl. Acad. Sci. U. S. A.* **111,** 13790–13794 (2014).

81. Hibar, D. P. *et al.* Common genetic variants influence human subcortical brain structures. *Nature* **520,** 224–229 (2015).

82. Locke, A. E. *et al.* Genetic studies of body mass index yield new insights for obesity biology. *Nature* **518,** 197–206 (2015).

83. Wood, A. R. *et al.* Defining the role of common variation in the genomic and biological architecture of adult human height. *Nat. Genet.* **46,** 1173–1186 (2014).

84. Alpérovitch, A. *et al.* Meta-analysis of 74,046 individuals identifies 11 new susceptibility loci for Alzheimer's disease. *Nat. Genet.* **45,** 1452–1458 (2013).

85. Otowa, T. *et al.* Meta-analysis of genome-wide association studies of anxiety disorders. *Mol. Psychiatry* **21,** 1391–1399 (2016).

86. Robinson, E. *et al.* Genetic risk for autism spectrum disorders and neuropsychiatric variation in the general population. *Nat. Genet.* **48,** 552–555 (2016).

87. Sklar, P. *et al.* Large-scale genome-wide association analysis of bipolar disorder identifies a new susceptibility locus near ODZ4. *Nat Genet* **43,** 977–983 (2011).

88. Lo, M.-T. *et al.* Genome-wide analyses for personality traits identify six genomic loci and show correlations with psychiatric disorders. *Nat. Genet.* **49,** 152–156 (2016).

89. Allport, G. W. & Odbert, H. S. *Trait-names: a psycho-lexical study*. *Psychological monographs* **47,** (Psychological Review Company, 1936).

90. Borghans, L., Duckworth, A. L., Heckman, J. J. & Weel, B. Ter. The economics and psychology of personality traits. *J. Hum. Resour.* **43,** 972–1059 (2008).

91. Heckman, J. J. & Kautz, T. Hard evidence on soft skills. *Labour Econ.* **19,** 451–464 (2012).

92. Joshi, P. K. *et al.* Genome-wide meta-analysis associates HLA-DQA1/DRB1 and LPA and lifestyle factors with human longevity. *Nat. Commun.* **8,** 910 (2017).

93. Benjamin, D. J., Brown, S. A. & Shapiro, J. M. Who is 'behavioral'? Cognitive ability and anomalous preferences. *J. Eur. Econ. Assoc.* **11,** 1231–1255 (2013).

94. Caspi, A. *et al.* The p Factor. *Clin. Psychol. Sci.* **2,** 119–137 (2014).

95. Weber, E. U., Blais, A. E. & Betz, N. E. A Domain-specific Risk-attitude Scale: Measuring Risk Perceptions and Risk Behaviors. *J. Behav. Decis. Mak. J. Behav. Dec. Mak.* **15,** 263–290 (2002).

96. Hanoch, Y., Johnson, J. G. & Wilke, A. Domain Specificity in Experimental Measures and Participant Recruitment: An Application to Risk-Taking Behavior. *Psychol. Sci.* **17,** 300–304 (2006).

97. Einav, B. L., Finkelstein, A., Pascu, I. & Cullen, M. R. How General Are Risk Preferences? Choices under Uncertainty in Different Domains. *Am. Econ. Rev.* **102,** 2606–2638 (2016).

98. Hayatbakhsh, M. R. *et al.* Cannabis Use and Obesity and Young Adults. *Am. J. Drug Alcohol Abuse* **36,** 350–356 (2010).

99. Kelleher, L. M., Stough, C., Sergejew, A. A. & Rolfe, T. The effects of cannabis on information-processing speed. *Addict. Behav.* **29,** 1213–1219 (2004).

100. De Alwis, D. *et al.* ADHD symptoms, autistic traits, and substance use and misuse in adult Australian twins. *J. Stud. Alcohol Drugs* **75,** 211–21 (2014).

101. Loh, P.-R. *et al.* Contrasting genetic architectures of schizophrenia and other complex diseases using fast variance-components analysis. *Nat. Genet.* **47,** 1385–1392 (2015).

102. Bansal, V. *et al.* GWAS results for educational attainment aid in identifying genetic heterogeneity of schizophrenia. Pre-print at *bioRxiv* (2017). doi:10.1101/114405

103. Turley, P. *et al.* MTAG: Multi-Trait Analysis of GWAS. *bioRxiv* (2017). doi:https://doi.org/10.1101/118810

104. Vilhjálmsson, B. J. *et al.* Modeling linkage disequilibrium increases accuracy of polygenicrisk scores. *Am. J. Hum. Genet.* **97,** 576–592 (2015).

105. Dudbridge, F. Power and predictive accuracy of polygenic risk scores. *PLoS Genet.* **9,** e1003348 (2013).

106. Purcell, S. M. *et al.* Common polygenic variation contributes to risk of schizophrenia and bipolar disorder. *Nature* **460,** 748–752 (2009).

107. Daetwyler, H. D., Villanueva, B. & Woolliams, J. A. Accuracy of predicting the genetic

risk of disease using a genome-wide approach. *PLoS One* **3,** e3395 (2008).

108. de Vlaming, R. *et al.* Meta-GWAS Accuracy and Power (MetaGAP) Calculator Shows that Hiding Heritability Is Partially Due to Imperfect Genetic Correlations across Studies. *PLoS Genet.* **13,** e1006495 (2017).

109. Sonnega, A. *et al.* Cohort profile: the Health and Retirement Study (HRS). *Int. J. Epidemiol.* **43,** 576–585 (2014).

110. Charness, G., Gneezy, U. & Imas, A. Experimental methods: Eliciting risk preferences. *J. Econ. Behav. Organ.* **87,** 43–51 (2013).

111. Roberts, B. W. Back to the future: Personality and Assessment and personality development. *J. Res. Pers.* **43,** 137–145 (2009).

112. Wong, A. & Carducci, B. Do sensation seeking, control orientation, ambiguity, and dishonesty traits affect financial risk tolerance? *Manag. Financ.* **42,** 34–41 (2016).

113. Horvath, P. & Zuckerman, M. Sensation seeking, risk appraisal, and risky behavior. *Pers. Individ. Dif.* **14,** 41–52 (1993).

114. Lauriola, M. & Levin, I. P. Personality traits and risky decision-making in a controlled experimental task: an exploratory study. *Pers. Individ. Dif.* **31,** 215–226 (2001).

115. Filbeck, G., Hatfield, P. & Horvath, P. Risk Aversion and Personality Type. *J. Behav. Financ.* **6,** 170–180 (2005).

116. Zaleskiewicz, T. Beyond risk seeking and risk aversion: personality and the dual nature of economic risk taking. *Eur. J. Pers.* **15,** S105--S122 (2001).

117. Almlund, M., Duckworth, A. L., Heckman, J. & Kautz, T. Personality Psychology and Economics. in *Handbook of the Economics of Education* 1–181 (Elsevier, 2011). doi:10.1016/B978-0-444-53444-6.00001-8

118. Gladstone, G. & Parker, G. Measuring a behaviorally inhibited temperament style: Development and initial validation of new self-report measures. *Psychiatry Res.* **135,** 133–143 (2005).

119. Rotter, J. B. Generalized expectancies for internal versus external control of reinforcement. *Psychol. Monogr. Gen. Appl.* **80,** 1–28 (1966).

120. John, O. P. The "Big Five" factor taxonomy: Dimensions of personality in the natural language and in questionnaires. In L. A. Pervin (Ed.), *Handbook of personality: Theory and research* (pp. 66-100). New York: Guilford Press.

121. Costa, P. T. & Widiger, T. A. The 5-factor model of personality and its relevance to personality-disorders. *J. Pers. Disord.* **6,** 343–359 (1992).

122. Digman, J. M. Personality Structure: Emergence of the Five-Factor Model. *Annu. Rev. Psychol.* **41,** 417–440 (1990).

123. Brim, O. G. *et al.* National Survey of Midlife Development in the United States (MIDUS), 1995-1996. (2003). doi:10.3886/ICPSR02760

124. Lachman, M. E. & Weaver, S. L. The Midlife Development Inventory (MIDI) personality scales: Scale construction and scoring. Technical report (1997).

125. Zuckerman, M. Sensation Seeking. in *Handbook of Individual Differences in Social behavior* 455–465 (The Guildford Press, 2009).

126. Zuckerman, M. *Sensation Seeking: Beyond the Optimal Level of Arousal*. (Erlbaum, 1974).

127. Zuckerman, M. *Behavioral expressions and biosocial bases of sensation seeking*. (Cambridge University Press, 1994).

128. Stoel, R. D., De Geus, E. J. C. & Boomsma, D. I. Genetic Analysis of Sensation Seeking with an Extended Twin Design. *Behav. Genet.* **36,** 229–237 (2006).

129. Thurik, R., Khedhaouria, A., Torrès, O. & Verheul, I. ADHD Symptoms and Entrepreneurial Orientation of Small Firm Owners. *Appl. Psychol.* **65,** 568–586 (2016).

130. Verheul, I. *et al.* The association between attention-deficit/hyperactivity (ADHD) symptoms and self-employment. *Eur. J. Epidemiol.* **31,** 793–801 (2016).

131. Verheul, I. *et al.* ADHD-like behavior and entrepreneurial intentions. *Small Bus. Econ.* **45,** 85–101 (2015).

132. Boomsma, D. I. *et al.* Genetic Epidemiology of Attention Deficit Hyperactivity Disorder (ADHD Index) in Adults. *PLoS One* **5,** e10621 (2010).

133. Conners, C., Erhardt, D. & Sparrow, E. *Conners' Adult ADHD Rating Scales (CAARS) technical manual*. (Multi-Health Systems, Inc., 1999).

134. Centers for Disease Control and Prevention (US), National Center for Chronic Disease Prevention and Health Promotion (US) & Office on Smoking and Health (US). *How Tobacco Smoke Causes Disease: The Biology and Behavioral Basis for Smoking-Attributable Disease: A Report of the Surgeon General*. *Public Health* (Centers for Disease Control and Prevention (US), 2010).

135. World Health Organisation. *Global status report on alcohol and health 2014*. (WHO Publications, 2014). doi:/entity/substance_abuse/publications/global_alcohol_report/en/index.html

136. Volkow, N. D., Baler, R. D., Compton, W. M. & Weiss, S. R. B. Adverse Health Effects of Marijuana Use. *N. Engl. J. Med.* **370,** 2219–2227 (2014).

137. Marie, O. & Zölitz, U. 'High' Achievers? Cannabis Access and Academic Performance. *Rev. Econ. Stud.* **84,** 1210–1237 (2017).

138. Cahill, L. E. *et al.* Fried-food consumption and risk of type 2 diabetes and coronary artery disease: a prospective study in 2 cohorts of US women and men. *Am. J. Clin. Nutr.* **100,** 667–675 (2014).

139. Valois, R. F., Oeltmann, J. E. & Waller, J. Relationship between number of sexual intercourse partners and selected health risk behaviors among public high school adolescents. *J. Adolesc. Heal.* **25,** 328–335 (1999).

140. Ashenhurst, J. R., Wilhite, E. R., Harden, K. P. & Fromme, K. Number of Sexual Partners and Relationship Status Are Associated With Unprotected Sex Across Emerging Adulthood. *Arch. Sex. Behav.* **46,** 419–432 (2017).

141. Wray, N. R. *et al.* Pitfalls of predicting complex traits from SNPs. *Nat. Rev. Genet.* **14,**

507–15 (2013).

142. Buchanan, C. C., Torstenson, E. S., Bush, W. S. & Ritchie, M. D. A comparison of cataloged variation between International HapMap Consortium and 1000 Genomes Project data. *J. Am. Med. Informatics Assoc.* **19,** 289–294 (2012).

143. Altshuler, D. M., Gibbs, R. A. & Peltonen, L. Integrating common and rare genetic variation in diverse human populations. *Nature* **467,** 52–58 (2010).

144. Canty, A. & Ripley, B. boot: Bootstrap R (S-Plus) Functions. *R Packag. version 1.3-17* (2015).

145. Davison, A. C. & Hinkley, D. V. *Bootstrap Methods and Their Application*. (Cambridge University Press, 1997).

146. Rustichini, A., DeYoung, C. G., Anderson, J. E. & Burks, S. V. Toward the integration of personality theory and decision theory in explaining economic behavior: An experimental investigation. *J. Behav. Exp. Econ.* **64,** 122–137 (2016).

147. Pedroni, A. *et al.* The risk elicitation puzzle. *Nat. Hum. Behav.* **1,** 803–809 (2017).

148. Levitt, S. D. & List, J. A. What Do Laboratory Experiments Measuring Social Preferences Reveal About the Real World? *J. Econ. Perspect.* **21,** 153–174 (2007).

149. Levitt, S. D. & List, J. A. Viewpoint: On the generalizability of lab behaviour to the field. *Can. J. Econ.* **40,** 347–370 (2007).

150. Polderman, T. J. C. *et al.* Meta-analysis of the heritability of human traits based on fifty years of twin studies. *Nat. Genet.* **47,** 702–709 (2015).

151. Nave, G., Camerer, C. & McCullough, M. Does oxytocin increase trust in humans? A critical review of research. *Perspect. Psychol. Sci.* **10,** 772–789 (2015).

152. Beauchamp, J. P. *et al.* Molecular genetics and economics. *J. Econ. Perspect.* **25,** 57–82 (2011).

153. Rietveld, C. A. *et al.* Replicability and robustness of GWAS for behavioral traits. *Psychol. Sci.* **25,** 1975–1986 (2014).

154. Benjamin, D. J. *et al.* The promises and pitfalls of genoeconomics. *Annu. Rev. Econom.* **4,** 627–662 (2012).

155. Okbay, A. & Rietveld, C. A. On improving the credibility of candidate gene studies: A review of candidate gene studies published in Emotion. *Emotion* **15,** 531–7 (2015).

156. de Leeuw, C. A., Mooij, J. M., Heskes, T. & Posthuma, D. MAGMA: Generalized Gene-Set Analysis of GWAS Data. *PLoS Comput. Biol.* **11,** 1–19 (2015).

157. Hintze, A., Olson, R. S., Adami, C. & Hertwig, R. Risk sensitivity as an evolutionary adaptation. *Sci. Rep.* **5,** 8242 (2015).

158. Netzer, N. Evolution of time preferences and attitudes toward risk. *Am. Econ. Rev.* **99,** 937–955 (2009).

159. Robson, A. J. The Evolution of Attitudes to Risk: Lottery Tickets and Relative Wealth. *Games Econ. Behav.* **14,** 190–207 (1996).

160. Subramanian, A. *et al.* Gene set enrichment analysis: a knowledge-based approach for interpreting genome-wide expression profiles. *Proc. Natl. Acad. Sci. U. S. A.* **102,** 15545–50 (2005).

161. Hawrylycz, M. *et al.* Canonical genetic signatures of the adult human brain. *Nat. Neurosci.* **18,** 1832–1844 (2015).

162. Westfall, P. H. & Young, S. S. *Resampling-Based Multiple Testing: Examples and Methods for p-Value Adjustment*. (Wiley, 1993).

163. de Leeuw, C. A., Neale, B. M., Heskes, T. & Posthuma, D. The statistical properties of gene-set analysis. *Nat. Rev. Genet.* **17,** 353–364 (2016).

164. Sherrington, R. *et al.* Cloning of the human dopamine D5 receptor gene and identification of a highly polymorphic microsatellite for the DRD5 locus that shows tight linkage to the chromosome 4p reference marker RAF1P1. *Genomics* **18,** 423–425 (1993).

165. Nakamura, M., Ueno, S., Sano, a & Tanabe, H. The human serotonin transporter gene linked polymorphism (5-HTTLPR) shows ten novel allelic variants. *Mol. Psychiatry* **5,** 32–38 (2000).

166. Peyrot, W. J. *et al.* Strong effects of environmental factors on prevalence and course of major depressive disorder are not moderated by 5-HTTLPR polymorphisms in a large Dutch sample. *J. Affect. Disord.* **146,** 91–99 (2013).

167. Auton, A. *et al.* A global reference for human genetic variation. *Nature* **526,** 68–74 (2015).

168. Pers, T. H. *et al.* Biological interpretation of genome-wide association studies using predicted gene functions. *Nat. Commun.* **6,** 5890 (2015).

169. Auton, A. *et al.* A global reference for human genetic variation. *Nature* **526,** 68–74 (2015).

170. Barski, A. *et al.* High-Resolution Profiling of Histone Methylations in the Human Genome. *Cell* **129,** 823–837 (2007).

171. Consortium, R. E. *et al.* Integrative analysis of 111 reference human epigenomes. *Nature* **518,** 317–330 (2015).

172. Zhou, V. W., Goren, A. & Bernstein, B. E. Charting histone modifications and the functional organization of mammalian genomes. *Nat. Rev. Genet.* **12,** 7–18 (2011).

173. Lindblad-Toh, K. *et al.* A high-resolution map of human evolutionary constraint using 29 mammals. *Nature* **478,** 476–482 (2011).

174. Raychaudhuri, S. Mapping rare and common causal alleles for complex human diseases. *Cell* **147,** 57–69 (2011).

175. Trynka, G. *et al.* Chromatin marks identify critical cell types for fine mapping complex trait variants. *Nat. Genet.* **45,** 124–130 (2013).

176. Kandasamy, N. *et al.* Cortisol shifts financial risk preferences. *Proc. Natl. Acad. Sci. U. S. A.* **111,** 3608–13 (2014).

177. Cueva, C. *et al.* Cortisol and testosterone increase financial risk taking and may

destabilize markets. *Sci. Rep.* **5,** 11206 (2015).

178. Mehta, P. H., Welker, K. M., Zilioli, S. & Carré, J. M. Testosterone and cortisol jointly modulate risk-taking. *Psychoneuroendocrinology* **56,** 88–99 (2015).

179. Sekar, A. *et al.* Schizophrenia risk from complex variation of complement component 4. *Nature* **530,** 177–183 (2016).

180. Network and Pathway Analysis Subgroup of the Psychiatric Genomics Consortium. Psychiatric genome-wide association study analyses implicate neuronal, immune and histone pathways. *Nat. Neurosci.* **18,** 199–209 (2015).

181. Wray, N. R. *et al.* Genome-wide association analyses identify 44 risk variants and refine the genetic architecture of major depression. *bioRxiv* (2017). doi:https://doi.org/10.1101/167577

182. Fehrmann, R. S. N. *et al.* Gene expression analysis identifies global gene dosage sensitivity in cancer. *Nat. Genet.* **47,** 115–125 (2015).

183. Brown, M. A method for combining non-independent, one-sided tests of significance. *Biometrics* **31,** 987–992 (1975).

184. The Gene Ontology Consortium. Gene ontology: Tool for the identification of biology. *Nat. Genet.* **25,** 25–29 (2000).

185. Croft, D. *et al.* Reactome: A database of reactions, pathways and biological processes. *Nucleic Acids Res.* **39,** 691–697 (2011).

186. Fabregat, A. *et al.* The Reactome pathway Knowledgebase. *Nucleic Acids Res.* **44,** D481–D487 (2016).

187. Kanehisa, M., Goto, S., Sato, Y., Furumichi, M. & Tanabe, M. KEGG for integration and interpretation of large-scale molecular data sets. *Nucleic Acids Res.* **40,** 109–114 (2012).

188. Purves, D. *et al. Neuroscience*. (Sinauer Associates, Inc., 2008).

189. Lenroot, R. K. & Giedd, J. N. Brain development in children and adolescents: Insights from anatomical magnetic resonance imaging. *Neurosci. Biobehav. Rev.* **30,** 718–729 (2006).

190. Gamazon, E. R. *et al.* A gene-based association method for mapping traits using reference transcriptome data. *Nat. Genet.* **47,** 1091–1098 (2015).

191. Gusev, A. *et al.* Integrative approaches for large-scale transcriptome-wide association studies. *Nat. Genet.* **48,** 245–52 (2016).

192. Zhu, Z. *et al.* Integration of summary data from GWAS and eQTL studies predicts complex trait gene targets. *Nat. Genet.* **48,** 481–7 (2016).

193. Smemo, S. *et al.* Obesity-associated variants within FTO form long-range functional connections with IRX3. *Nature* **507,** 371–5 (2014).

194. Tadeu, A. M. B. *et al.* CENP-V is required for centromere organization, chromosome alignment and cytokinesis. *EMBO J.* **27,** 2510–2522 (2008).

195. Honda, Z., Suzuki, T. & Honda, H. Identification of CENP-V as a novel microtubule-associating molecule that activates Src family kinases through SH3 domain interaction.

*Genes to Cells* **14,** 1383–1394 (2009).

196. Vermeulen, S. J. *et al.* The αE-catenin gene (CTNNA1) acts as an invasion-suppressor gene in human colon cancer cells. *Oncogene* **18,** 905–915 (1999).

197. Saksens, N. T. M. *et al.* Mutations in CTNNA1 cause butterfly-shaped pigment dystrophy and perturbed retinal pigment epithelium integrity. *Nat. Genet.* **48,** 144–151 (2015).

198. Öngür, D., Ferry, A. T. & Price, J. L. Architectonic subdivision of the human orbital and medial prefrontal cortex. *J. Comp. Neurol.* **460,** 425–449 (2003).

199. Wise, S. P. Forward Frontal Fields: Phylogeny and Fundamental Function. *Trends Neurosci.* **31,** 599–608 (2008).

200. Miller, E. K. & Cohen, J. D. An integrative theory of prefrontal cortex function. *Annu. Rev. Neurosci.* **24,** 167–202 (2001).

201. Petrides, M. Frontal lobes and behaviour. *Curr. Opin. Neurobiol.* **4,** 207–211 (1994).

202. Paus, T. Primate anterior cingulate cortex: Where motor control, drive and cognition interface. *Nat. Rev. Neurosci.* **2,** 417–424 (2001).

203. Botvinick, M. M., Cohen, J. D. & Carter, C. S. Conflict monitoring and anterior cingulate cortex: An update. *Trends in Cognitive Sciences* **8,** 539–546 (2004).

204. Heilbronner, S. R. & Hayden, B. Y. Dorsal Anterior Cingulate Cortex: A Bottom-Up View. *Annu. Rev. Neurosci.* **39,** 149–170 (2016).

205. Parent, A. & Hazrati, L. N. Functional anatomy of the basal ganglia. *Rev. Neurol.* **25 Suppl 2,** S121–S128 (1997).

206. Baxter, M. G. & Murray, E. A. The amygdala and reward. *Nat. Rev. Neurosci.* **3,** 563–573 (2002).

207. Ikemoto, S., Yang, C. & Tan, A. Basal ganglia circuit loops, dopamine and motivation: A review and enquiry. *Behavioural Brain Research* **290,** 17–31 (2015).

208. Nelson, A. B. & Kreitzer, A. C. Reassessing Models of Basal Ganglia Function and Dysfunction. *Annu. Rev. Neurosci.* **37,** 117–135 (2014).

209. Hikosaka, O., Kim, H. F., Yasuda, M. & Yamamoto, S. Basal Ganglia Circuits for Reward Value–Guided Behavior. *Annu. Rev. Neurosci.* **37,** 289–306 (2014).

210. Haber, S. N. & Knutson, B. The reward circuit: linking primate anatomy and human imaging. *Neuropsychopharmacology* **35,** 4–26 (2010).

211. Frey, B. J. & Dueck, D. Clustering by passing messages between data points. *Science* **315,** (2007).

212. de Wit, J., Hong, W., Luo, L. & Ghosh, A. Role of Leucine-Rich Repeat Proteins in the Development and Function of Neural Circuits. *Annu. Rev. Cell Dev. Biol.* **27,** 697–729 (2011).

213. de Wit, J. & Ghosh, A. Specification of synaptic connectivity by cell surface interactions. *Nat. Rev. Neurosci.* **17,** 4–4 (2015).

214. Lefebvre, J. L., Sanes, J. R. & Kay, J. N. Development of Dendritic Form and Function.

*Annu. Rev. Cell Dev. Biol.* **31,** 741–777 (2015).

215. Sigel, E. & Steinmann, M. E. Structure, Function, and Modulation of GABA $_A$ Receptors. *J. Biol. Chem.* **287,** 40224–40231 (2012).

216. Tyagarajan, S. K. & Fritschy, J.-M. Gephyrin: a master regulator of neuronal function? *Nat. Rev. Neurosci.* **15,** 141–56 (2014).

217. Ariani, F. *et al.* FOXG1 is responsible for the congenital variant of Rett syndrome. *Am. J. Hum. Genet.* **83,** 89–93 (2008).

218. Stessman, H. A. F. *et al.* Targeted sequencing identifies 91 neurodevelopmental-disorder risk genes with autism and developmental-disability biases. *Nat. Genet.* **49,** 515–526 (2017).

219. Fowler, K. D. *et al.* Leveraging existing data sets to generate new insights into Alzheimer's disease biology in specific patient subsets. *Sci. Rep.* **5,** 14324 (2015).

220. Salih, D. A. M. *et al.* FoxO6 regulates memory consolidation and synaptic function. *Genes Dev.* **26,** 2780–2801 (2012).

221. Ward, L. D. & Kellis, M. HaploReg: a resource for exploring chromatin states, conservation, and regulatory motif alterations within sets of genetically linked variants. *Nucleic Acids Res.* **40,** D930-4 (2012).

222. Hyman, S. L., Shores, A. & North, K. N. The nature and frequency of cognitive deficits in children with neurofibromatosis type 1. *Neurology* **65,** 1037–44 (2005).

223. Ardlie, K. G. *et al.* The Genotype-Tissue Expression (GTEx) pilot analysis: Multitissue gene regulation in humans. *Science* **348,** 648–660 (2015).

224. McKenzie, M., Henders, A. K., Caracella, A., Wray, N. R. & Powell, J. E. Overlap of expression quantitative trait loci (eQTL) in human brain and blood. *BMC Med. Genomics* **7,** 31 (2014).

225. Sullivan, P. F., Fan, C. & Perou, C. M. Evaluating the comparability of gene expression in blood and brain. *Am. J. Med. Genet. B. Neuropsychiatr. Genet.* **141B,** 261–268 (2006).

226. Bauer, M., Goetz, T., Glenn, T. & Whybrow, P. C. The thyroid-brain interaction in thyroid disorders and mood disorders. *J. Neuroendocrinol.* **20,** 1101–1114 (2008).

227. Bégin, M. E., Langlois, M. F., Lorrain, D. & Cunnane, S. C. Thyroid function and cognition during aging. *Curr. Gerontol. Geriatr. Res.* **2008,** 1–11 (2008).

228. Casey, B. J., Jones, R. M. & Hare, T. A. The adolescent brain. *Ann. N. Y. Acad. Sci.* **1124,** 111–26 (2008).

229. Bunge, S. A., Dudukovic, N. M., Thomason, M. E., Vaidya, C. J. & Gabrieli, J. D. E. Immature frontal lobe contributions to cognitive control in children: evidence from fMRI. *Neuron* **33,** 301–11 (2002).

230. Zhou, X. *et al.* Behavioral response inhibition and maturation of goal representation in prefrontal cortex after puberty. *Proc. Natl. Acad. Sci. United States Am.* **113,** 3353–3358 (2016).

231. Tobler, P. N. & Weber, E. U. Valuation for Risky and Uncertain Choices. in

*Neuroeconomics* 149–172 (Elsevier, 2014). doi:10.1016/B978-0-12-416008-8.00009-7

232. Schultz, W. Getting formal with dopamine and reward. *Neuron* **36,** 241–263 (2002).
233. Petroff, O. A. C. GABA and Glutamate in the Human Brain. *Neurosci.* **8,** 562–573 (2002).
234. Purcell, S. M. *et al.* A polygenic burden of rare disruptive mutations in schizophrenia. *Nature* **506,** 185–190 (2014).
235. Fromer, M. *et al.* De novo mutations in schizophrenia implicate synaptic networks. *Nature* **506,** 179–184 (2014).
236. Andersson, R. *et al.* An atlas of active enhancers across human cell types and tissues. *Nature* **507,** 455–461 (2014).
237. Schaller, M. & Park, J. H. The Behavioral Immune System (and Why It Matters). *Curr. Dir. Psychol. Sci.* **20,** 99–103 (2011).
238. Schaller, M. The behavioural immune system and the psychology of human sociality. *Philos. Trans. R. Soc. Lond. B. Biol. Sci.* **366,** 3418–26 (2011).
239. Fujita, E. *et al.* Autism spectrum disorder is related to endoplasmic reticulum stress induced by mutations in the synaptic cell adhesion molecule, CADM1. *Cell Death Dis.* **1,** e47–e47 (2010).
240. Fujiwara, T. & Kawachi, I. Social Capital and Health. A Study of Adult Twins in the U.S. *Am. J. Prev. Med.* **35,** 139–144 (2008).
241. Huang, M. *et al.* Generation of a monoclonal antibody specific to a new candidate tumor suppressor, cell adhesion molecule 2. *Tumor Biol.* **35,** 7415–7422 (2014).
242. Howie, B. N., Donnelly, P. & Marchini, J. A flexible and accurate genotype imputation method for the next generation of genome-wide association studies. *PLoS Genet.* **5,** e1000529 (2009).
243. Li, Y., Willer, C. J., Ding, J., Scheet, P. & Abecasis, G. R. MaCH: Using sequence and genotype data to estimate haplotypes and unobserved genotypes. *Genet. Epidemiol.* **34,** 816–834 (2010).
244. Hill, A. V *et al.* Common west African HLA antigens are associated with protection from severe malaria. *Nature* **352,** 595–600 (1991).
245. Hill, W. D., Hagenaars, S. P., Marioni, R. E. & Harris, S. E. Molecular genetic contributions to social deprivation and household income in UK Biobank (n=112,151). *Curr. Biol.* **26,** 3083–3089 (2016).
246. Hsieh, C., Parker, S. C. & van Praag, C. M. Risk, balanced skills and entrepreneurship. *Small Bus. Econ.* **48,** 287–302 (2017).
247. Clarke, T.-K. *et al.* Genome-wide association study of alcohol consumption and genetic overlap with other health-related traits in UK Biobank (N=112 117). *Mol. Psychiatry* **22,** 1376–1384 (2017).
248. Nelson, S., Zhao, W., Smith, J. & Faul, J. Health and Retirement Study: Information for dbGaP users on annotation issues in the Illumina HumanOmni2.5-4v1_D manifest. (2014).

249. Tobin, J. Estimation of Relationships for Limited Dependent Variables. *Econometrica* **26,** 24–36 (1958).
