## Extended Data Items for "Genome-wide association analyses of risk tolerance and risky behaviors in over 1 million individuals identify hundreds of loci and shared genetic influences^1^"

[a] Previous title: Genome-wide study identifies 611 loci associated with risk tolerance and risky behaviors.

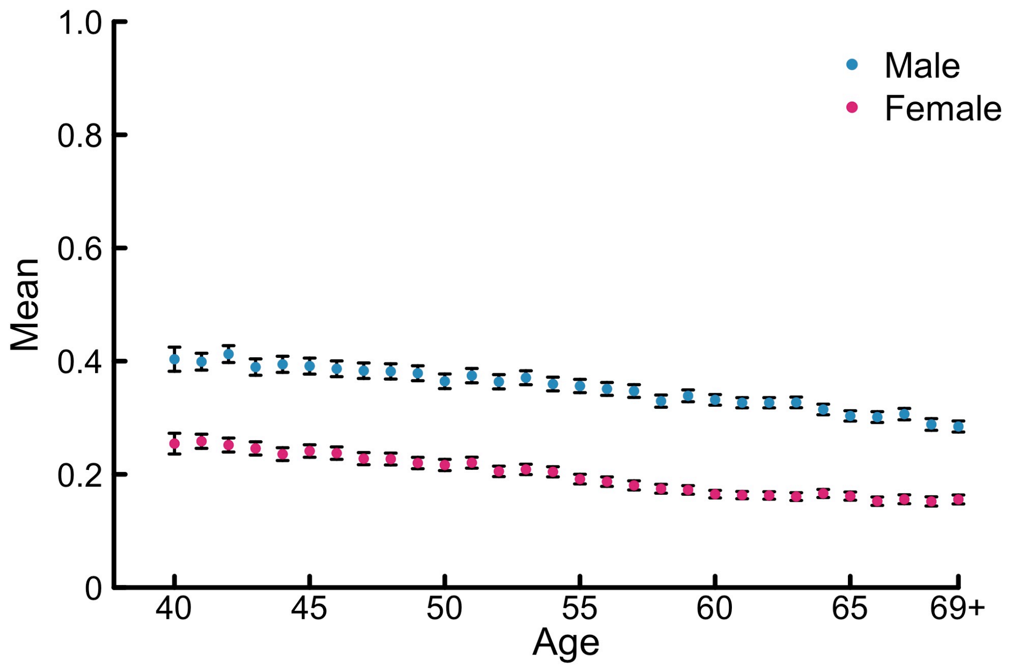


**Extended Data Figure 1.1 | Mean phenotypic general risk tolerance as a function of age, for males and females in the UKB cohort.** The whiskers represent 95% confidence intervals. Individuals aged 69 or older have been grouped together ("69+"), as there were few individuals aged 70 or more.

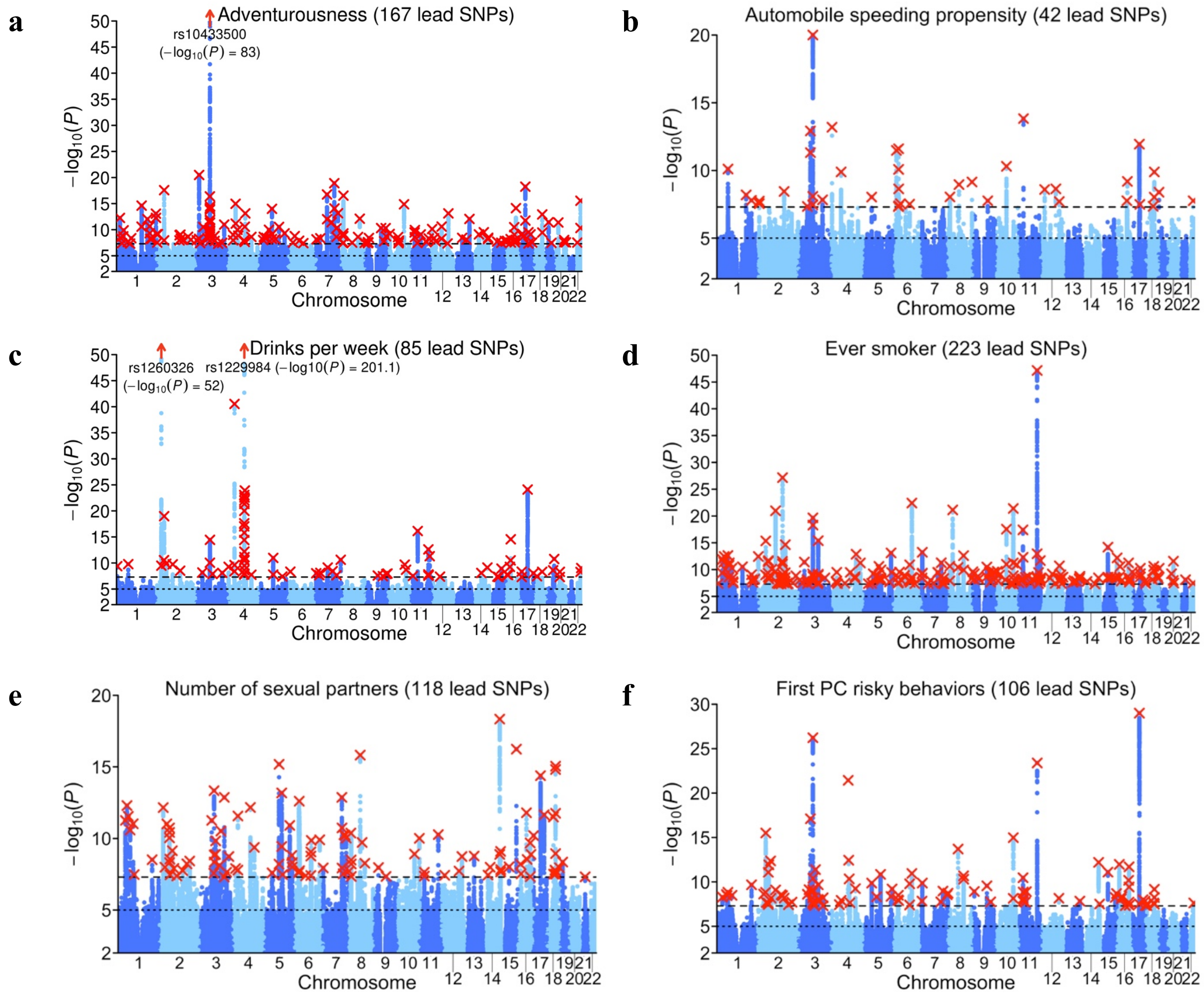


**Extended Data Figure 3.1 | Manhattan plots for the six supplementary GWAS.** Manhattan plots for the GWAS of (**a**) adventurousness, (**b**) automobile speeding propensity, (**c**) drinks per week, (**d**) ever smoker, (**e**) number of sexual partners, and (**f**) the first PC of the risky behaviors. The *x*-axis is chromosomal position, and the *y*-axis is the significance on a $-\log_{10}$ scale. The upper dashed line marks the threshold for genome-wide significance ($P = 5\times10^{-8}$); the lower line marks the threshold for nominal significance ($P = 10^{-5}$). Each approximately independent genome-wide significant association ("lead SNP") is marked by a red ×. Each lead SNP is the SNP with the lowest *P* value within the locus, as defined by our clumping algorithm.

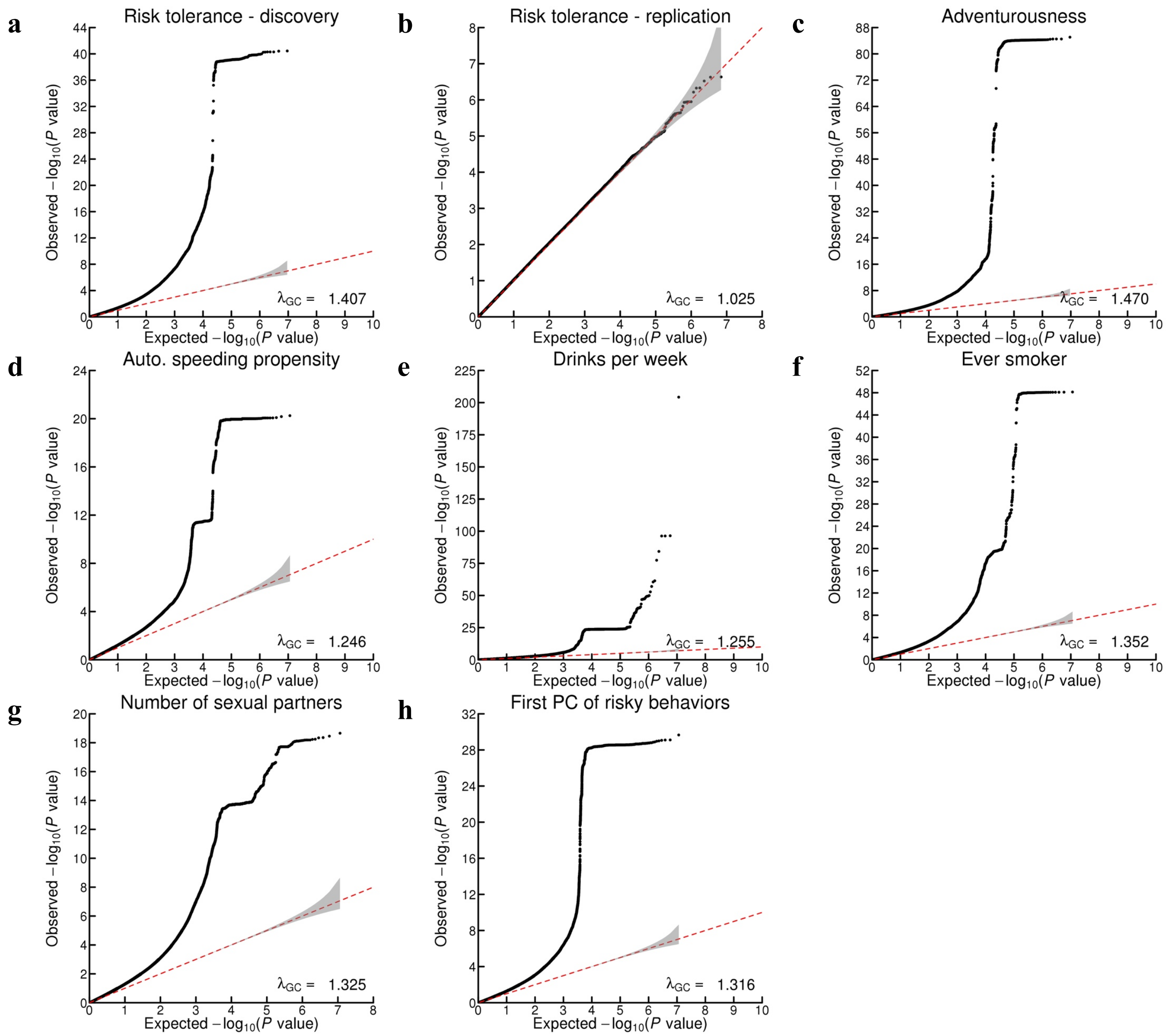


**Extended Data Figure 3.2 | Quantile-quantile plots.** The panels display Q-Q plots for (**a**) the discovery and (**b**) the replication GWAS of general risk tolerance, and for the GWAS of (**c**) adventurousness, (**d**) automobile speeding propensity, (**e**) drinks per week, (**f**) ever smoker, (**g**) number of sexual partners, and (**h**) the first PC of the four risky behaviors, before adjustment of the standard errors. The gray shaded areas in the Q-Q plots represent the 95% confidence intervals under the null hypothesis. Though we report $\lambda_{GC}$, we used the square root of the estimated LD Score intercept to adjust the standard errors of the coefficient estimates in the GWAS, as described in **Supplementary Information section 2.7**.

**a**

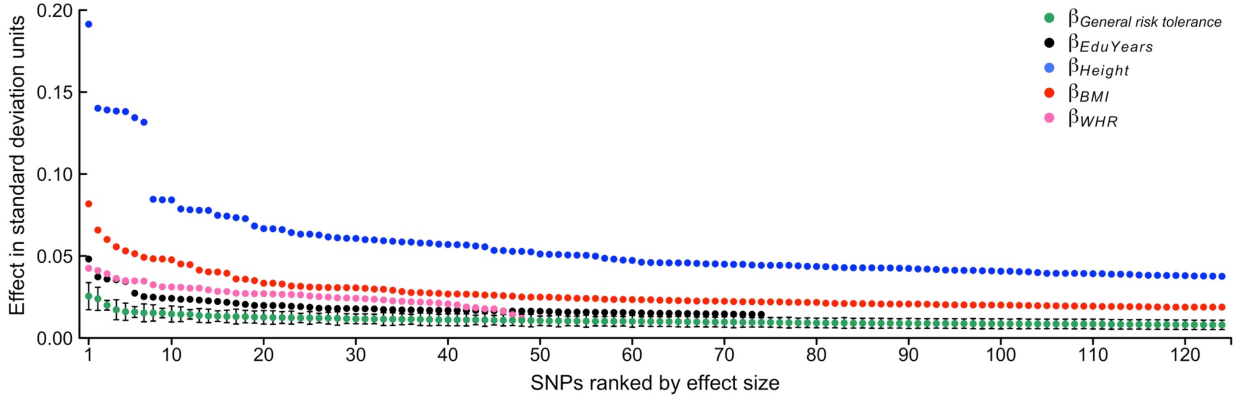


**b**

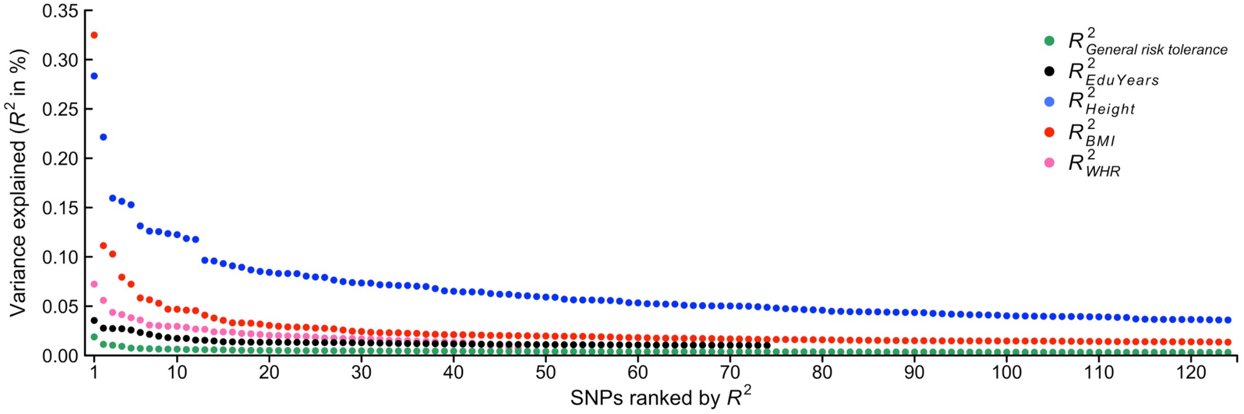


**Extended Data Figure 3.3 | Distribution of effect sizes of the 124 general-risk-tolerance lead SNPs, compared with various phenotypes. a**, Estimated effect sizes (in standard deviations (*SD*) of general risk tolerance per risk-tolerance increasing allele) and 95% confidence intervals from the discovery GWAS of general risk tolerance, with the SNPs ranked by their general-risk-tolerance effect sizes. **b**, variance explained ($R^2$), with the SNPs ranked by their general-risk-tolerance variance explained ($R^2$). The effect sizes are benchmarked against the 124 top associations previously reported for height and for body mass index (BMI), against the 74 top associations previously reported for educational attainment (EduYears), and against the 48 top associations previously reported for waist-to-hip ratio adjusted for BMI (WHR). The effect sizes for height, BMI, and WHR are based on the GIANT consortium's publicly available results for pooled analyses restricted to European-ancestry individuals (https://www.broadinstitute.org/collaboration/giant/index.php/GIANT_consortium); the effect sizes for EduYears are from Okbay *et al.*[1].

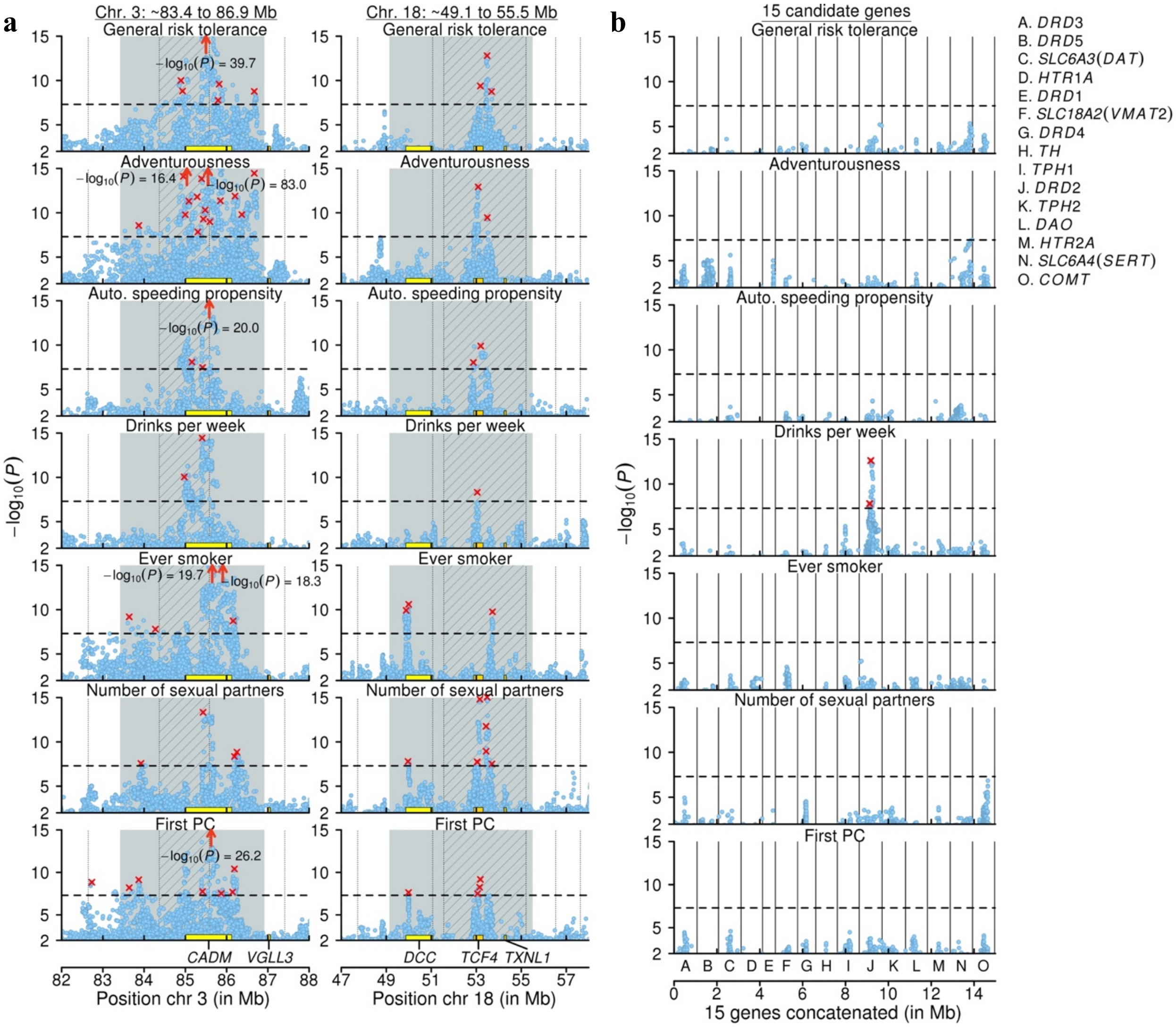
a
Chr. 3: ~83.4 to 86.9 Mb
Chr. 18: ~49.1 to 55.5 Mb
General risk tolerance
Adventurousness
Auto. speeding propensity
Drinks per week
Ever smoker
Number of sexual partners
First PC
−log10(P) = 39.7
−log10(P) = 16.4
−log10(P) = 83.0
−log10(P) = 20.0
−log10(P) = 19.7
−log10(P) = 18.3
−log10(P) = 26.2
CADM
VGLL3
DCC
TCF4
TXNL1
Position chr 3 (in Mb)
Position chr 18 (in Mb)
−log10(P)
b
15 candidate genes
15 genes concatenated (in Mb)
A. DRD3
B. DRD5
C. SLC6A3(DAT)
D. HTR1A
E. DRD1
F. SLC18A2(VMAT2)
G. DRD4
H. TH
I. TPH1
J. DRD2
K. TPH2
L. DAO
M. HTR2A
N. SLC6A4(SERT)
O. COMT

**c**

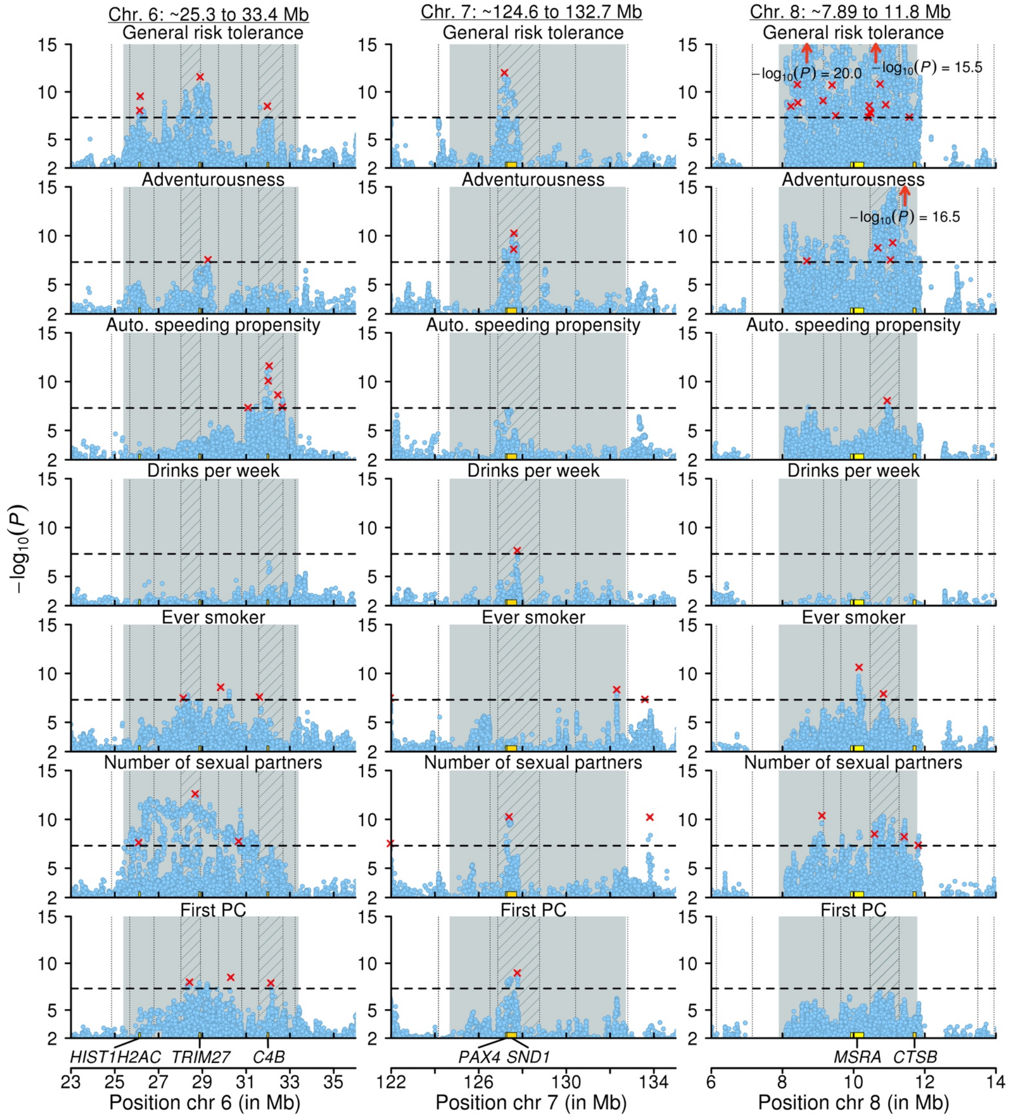


**Extended Data Figure 3.4 | Local Manhattan plots for selected genomic regions.** Each row corresponds to one of our seven GWAS; the $x$-axis is chromosomal position; the $y$-axis is the significance on a $-\log_{10}$ scale; the horizontal dashed line marks the threshold for genome-wide significance ($P = 5\times10^{-8}$); and each approximately independent genome-wide significant association ("lead SNP") is marked by a red ×. **a** and **c**, Plots for two genomic regions that contain lead SNPs for all or most of our seven GWAS. **b**, Plots for the loci around the 15 most commonly tested candidate genes in the prior literature on the genetics of risk tolerance. Each locus comprises all SNPs within 500 kb of the gene's borders that are in LD ($r^2 > 0.1$) with a SNP in the gene. The 15 plots are concatenated and shown together in the panel, divided by the black vertical lines. In panels **a** and **c**, the gray background marks the locations of candidate inversions or long-range LD regions; the gray vertical dotted lines mark the boundaries between the approximately independent LD blocks[2]; and the striped areas denote LD blocks with lead SNPs from all or most of all GWAS. See **Supplementary Information sections 2** and **3** for additional details.

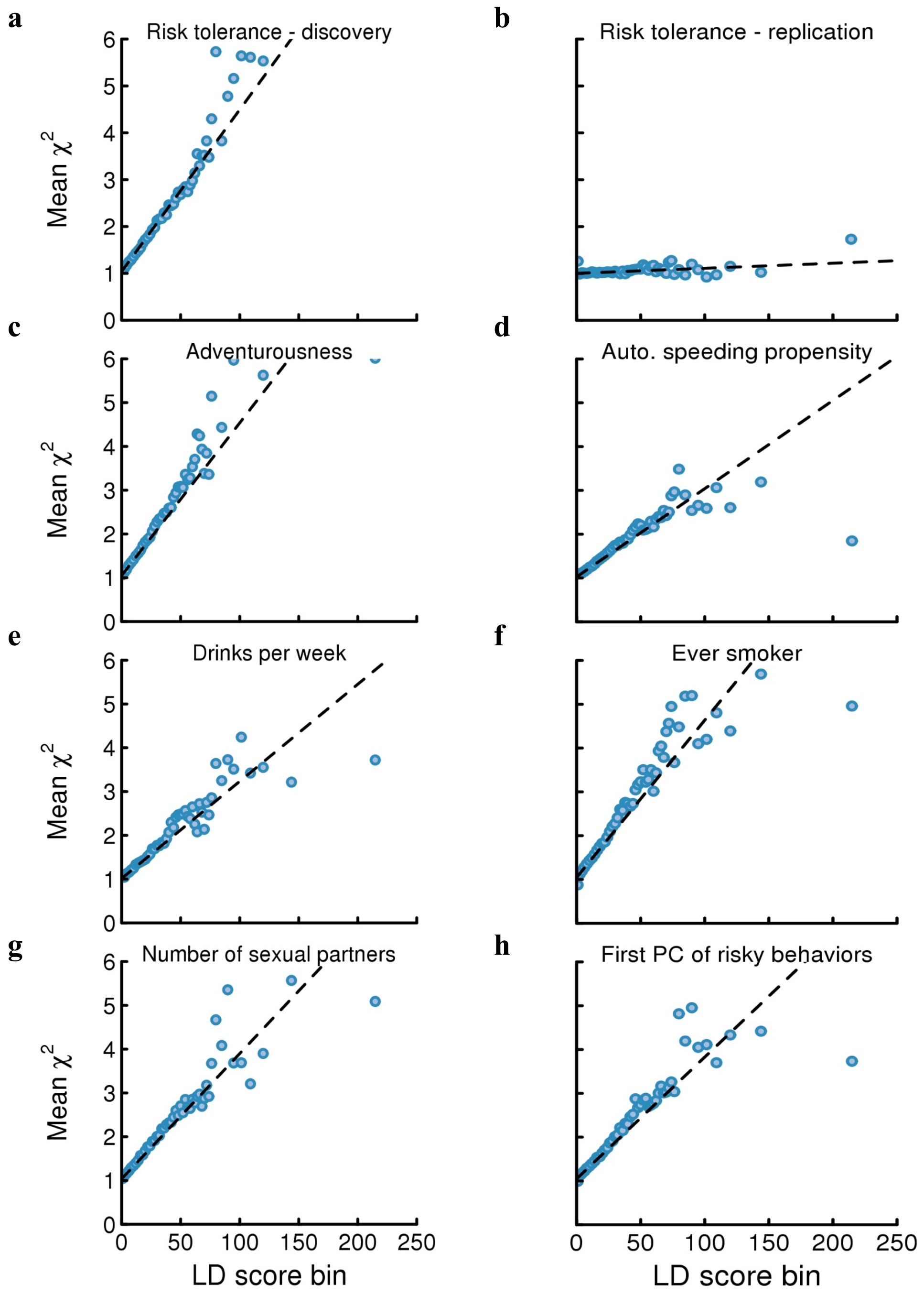


**Extended Data Figure 4.1 | LD Score regression plots.** The plots are based on the summary statistics from (**a**) the discovery and (**b**) the replication GWAS of general risk tolerance, and from the GWAS of (**c**) adventurousness, (**d**) automobile speeding propensity, (**e**) drinks per week, (**f**) ever smoker, (**g**) number of sexual partners, and (**h**) the first PC of the four risky behaviors, before adjustment of the standard errors. Each point represents an LD score bin. The $x$ and $y$ coordinates of the point are the mean LD score and the mean $\chi^2$ statistic of SNPs in that bin. The facts that the intercepts are close to one and that the $\chi^2$ statistics increase linearly with the LD scores for all GWAS suggest that, for all GWAS, the bulk of the inflation in the $\chi^2$ statistics is due to true polygenic signal and not to population stratification.

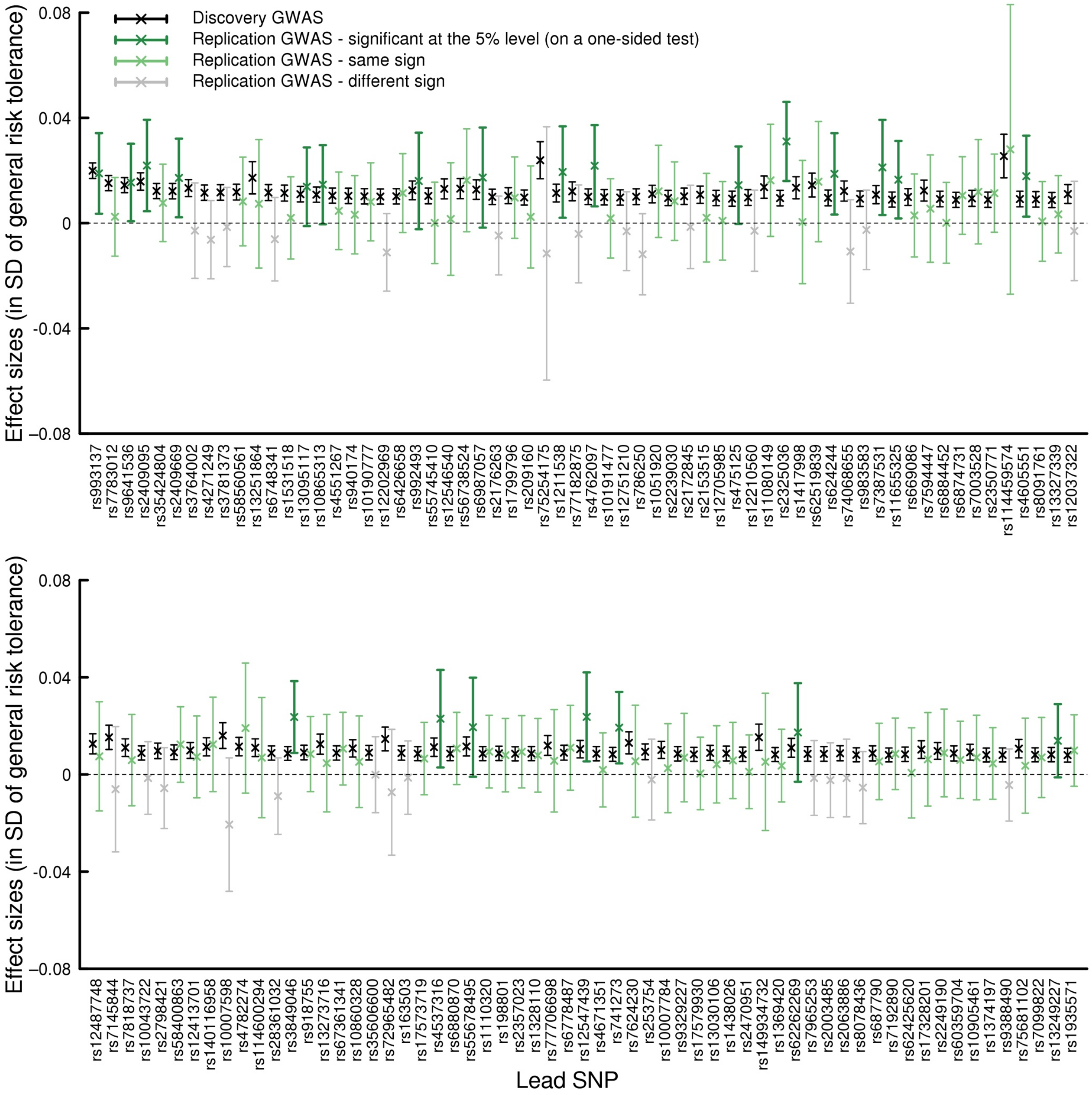


**Extended Data Figure 5.1 | Replication of lead SNPs from the discovery GWAS of general risk tolerance in the replication GWAS.** Estimated effect sizes (in standard deviations (*SD*) of general risk tolerance per risk-tolerance reference allele) and 95% confidence intervals for 122 general-risk-tolerance lead SNPs and 1 proxy-lead SNP, in the discovery and in the replication GWAS of general risk tolerance. (Two lead SNPs were not included in the replication GWAS, and a proxy-lead SNP could only be found for one of them.) The reference allele is the allele associated with higher values of general risk tolerance in the discovery GWAS. SNPs are listed from left to right in descending order of their $R^2$ in the discovery GWAS, with the 62 SNPs with the largest $R^2$'s in the top panel and the remaining 61 SNPs in the bottom panel. Of the 123 lead or proxy-lead SNPs, 94 have the anticipated sign in the replication sample and 23 replicate at the 0.05 significance level (on one-sided tests). See **Supplementary Information section 5** for additional details.

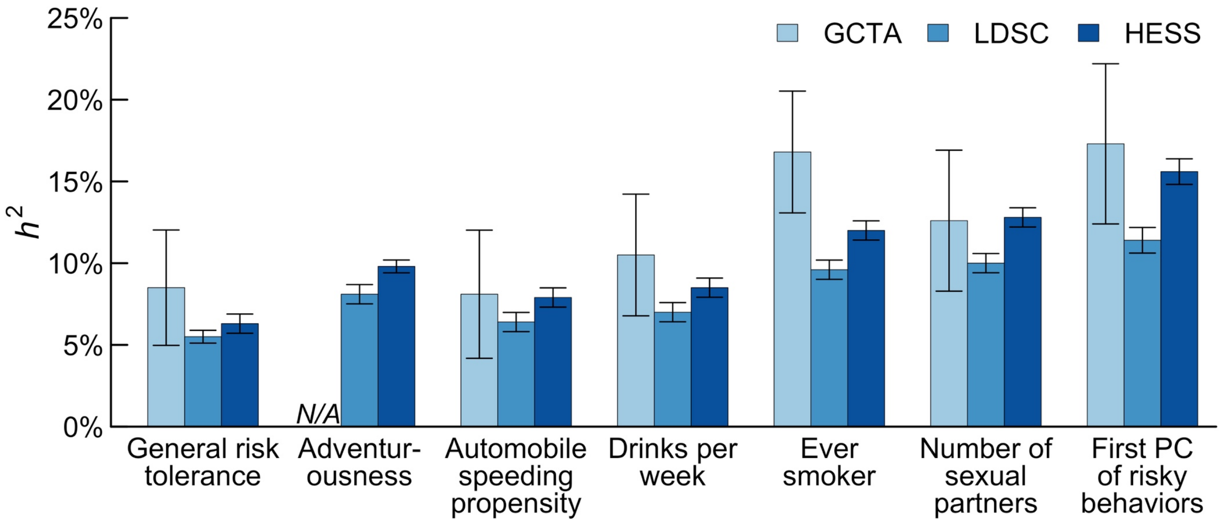


**Extended Data Figure 6.1 | Estimates of the SNP heritability of general risk tolerance and the six supplementary phenotypes.** SNP heritability was estimated with the GCTA, LD Score regression, and Heritability Estimator from Summary Statistics (HESS) methods. GCTA heritability was estimated using a random draw of 30,000 individuals from the UKB GWAS sample, from which we excluded cryptically related individuals, and using all genotyped SNPs with MAF > 0.01 (GCTA SNP heritability was not estimated for adventurousness because this phenotype is not available in the UKB and we did not have access to the individual-level data from 23andMe). For the LD Score and HESS methods, for all phenotypes except adventurousness we used summary statistics from the UKB GWAS only; for the adventurousness phenotype, we used the 23andMe summary statistics. LD Score heritability was estimated using HapMap3 SNPs with MAF > 0.01. HESS heritability was estimated using 1000 Genomes phase 3 SNPs with MAF > 0.05. See **Supplementary Information section 6** and **Supplementary Table 6.1** for additional details.

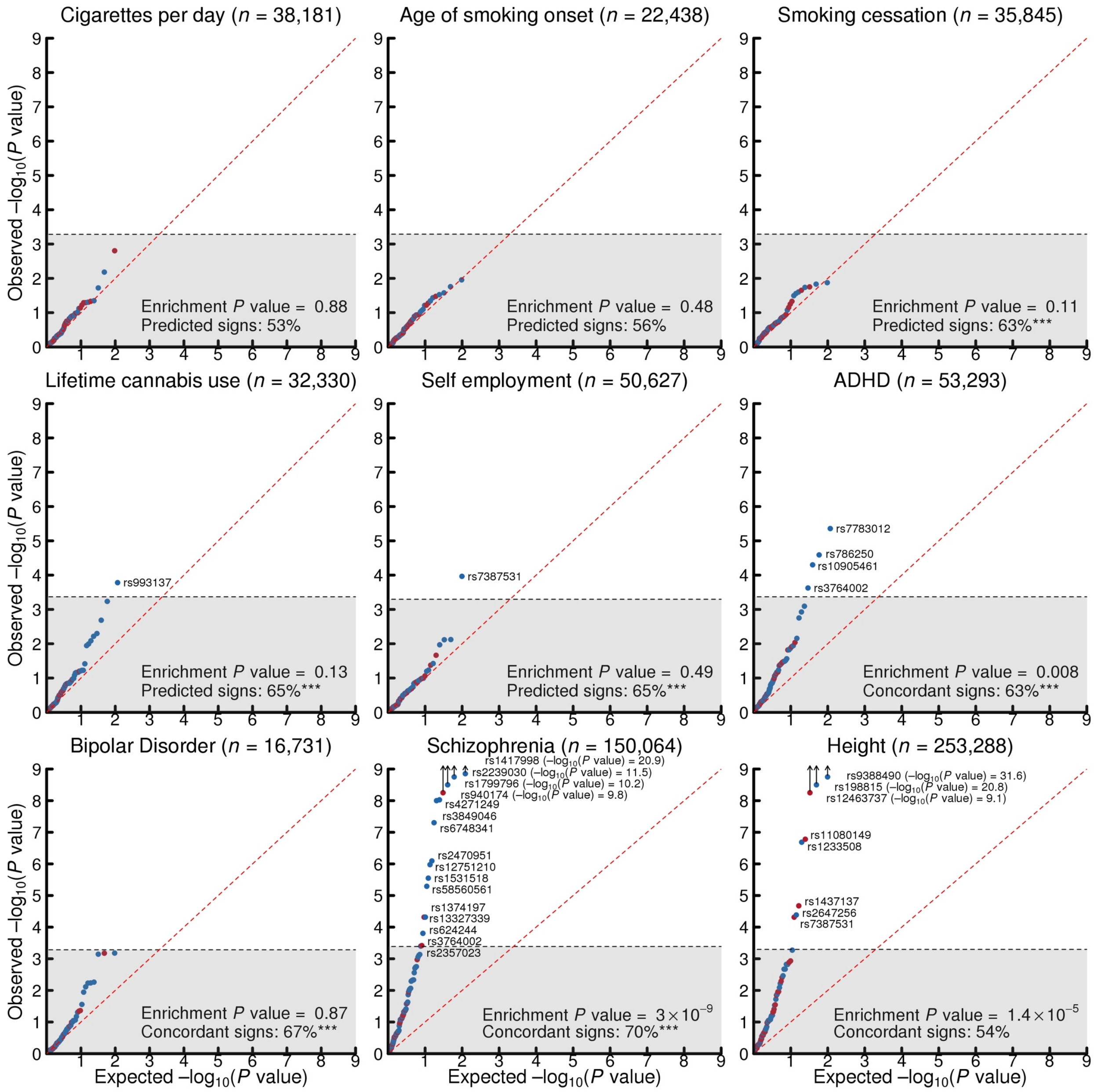


**Extended Data Figure 8.1 | Quantile-quantile plots for the general-risk-tolerance lead SNPs in previous GWAS of other phenotypes.** SNPs with effects in the predicted or concordant direction in the published GWAS are blue, and SNPs with effects in the other direction are red. SNPs outside the grey area pass Bonferroni-corrected significance thresholds that correct for the total number of SNPs we tested for each published GWAS, and are labelled with their rs numbers. Observed and expected $P$ values are on a $-\log_{10}$ scale. For each published GWAS, the enrichment $P$ value corresponds to the Mann-Whitney test of joint enrichment, and the percentage of SNPs with predicted or concordant signs is shown along with stars denoting the $P$ value of the sign test: * $P < 0.10$, ** $P < 0.05$ and *** $P < 0.01$. See **Supplementary Information section 8** for additional details.

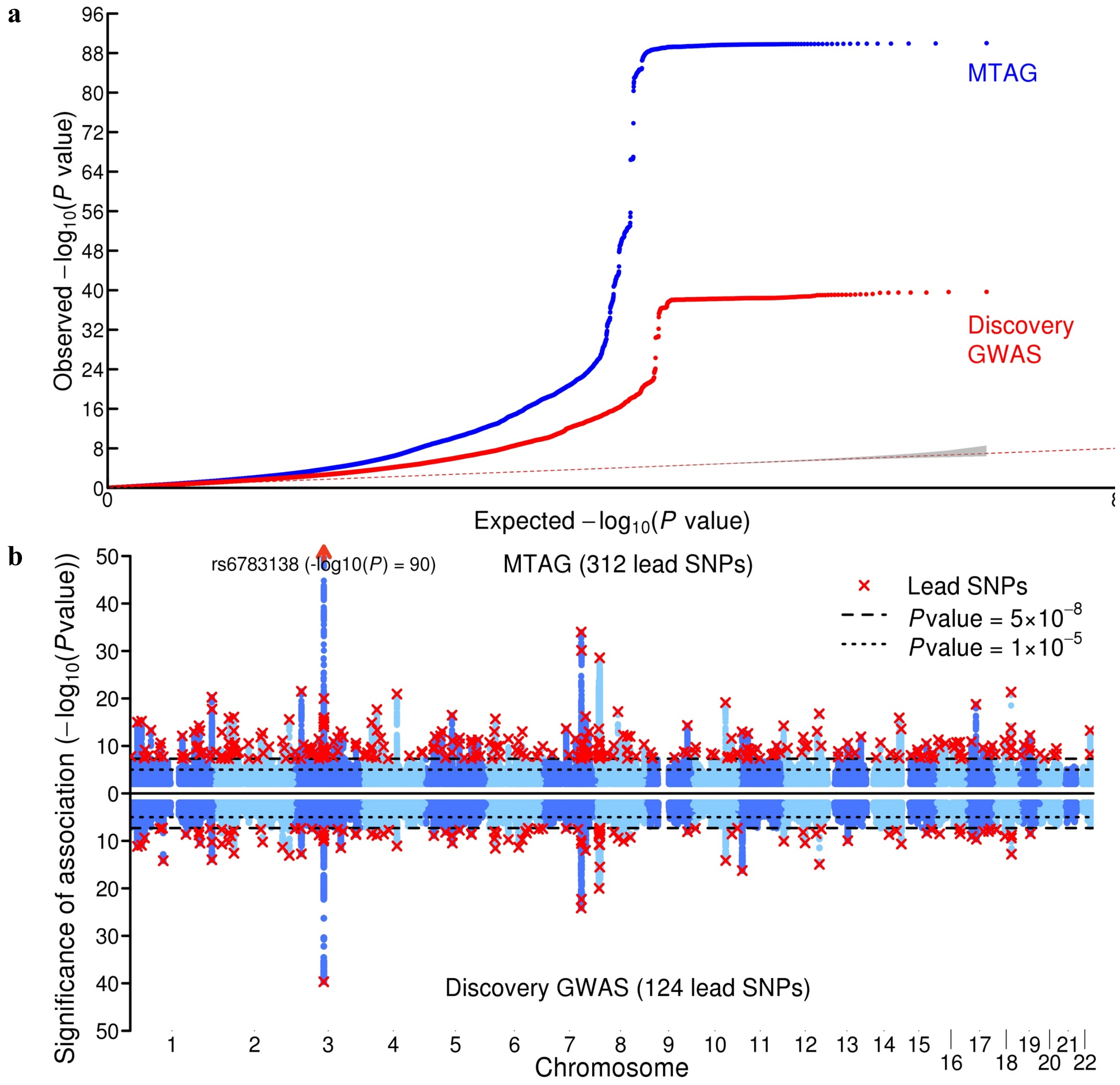


**Extended Data Figure 9.1 | Results from the MTAG analysis of general risk tolerance. a**, Quantile-quantile plots for the MTAG analysis of general risk tolerance (see the **Supplementary Information section 9** for details) and for the discovery GWAS of general risk tolerance. The gray shaded area represents the 95% confidence interval under the null hypothesis. **b**, Manhattan plots for the MTAG analysis of general risk tolerance (top panel) and for the discovery GWAS of general risk tolerance (bottom panel). The *x*-axis is chromosomal position, and the *y*-axis is the significance on a $-\log_{10}$ scale. The long-dashed line marks the threshold for genome-wide significance ($P = 5\times10^{-8}$); the short-dashed line marks the threshold for nominal significance ($P = 10^{-5}$). Each approximately independent genome-wide significant association ("lead SNP") is marked by a red ×.

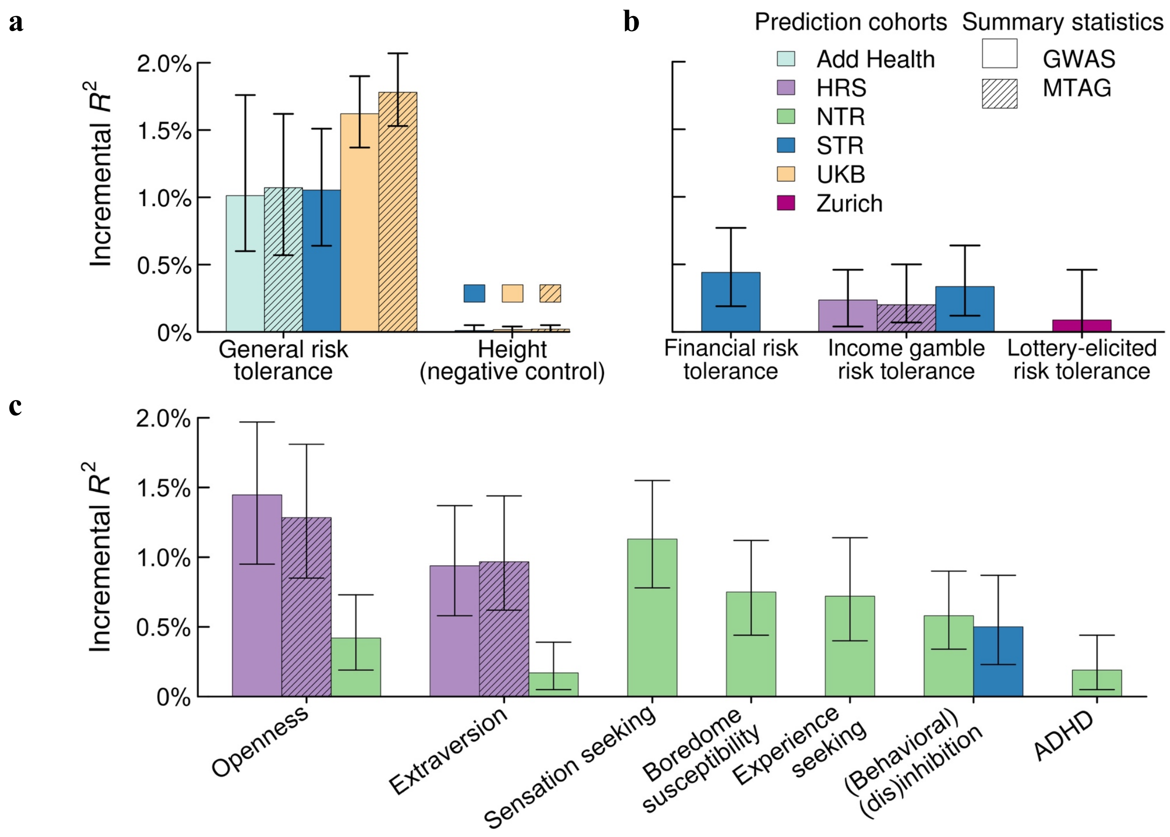


**Extended Data Figure 10.1 | Prediction of measures of risk tolerance and of personality traits with polygenic scores of general risk tolerance.** Incremental $R^2$ is defined as the increase in $R^2$ from adding the score to a regression of the predicted phenotype on controls for sex, age, and the top ten principal components of the genetic relatedness matrix. The scores were constructed using LDpred with our preferred Gaussian mixture weight of 0.3. The validation cohorts are the Add Health, HRS, NTR, STR, UKB-siblings, and Zurich cohorts. For the Add Health and HRS cohorts, scores were constructed using summary statistics from the meta-analysis of the discovery and replication GWAS ($n$ = 975,353) and using summary statistics from the MTAG analysis of general risk tolerance; for the UKB-siblings cohort, scores were constructed in the same way but excluding individuals with at least one full sibling in the UKB from the meta-analysis ($n$ = 937,353); for the other validation cohorts, scores were constructed using summary statistics from meta-analyses that exclude the 23andMe cohort (due to data access limitations) ($n$ = 466,571 for the NTR and Zurich cohorts; for the STR cohort the meta-analysis also excluded the STR cohort, $n$ = 458,558). Results are displayed for the prediction of (**a**) general risk tolerance and height (as a negative control test), (**b**) alternative risk tolerance phenotypes, (**c**) selected personality traits and ADHD. See **Supplementary Information section 10** and **Supplementary Tables 10.1-10.3** for additional details.

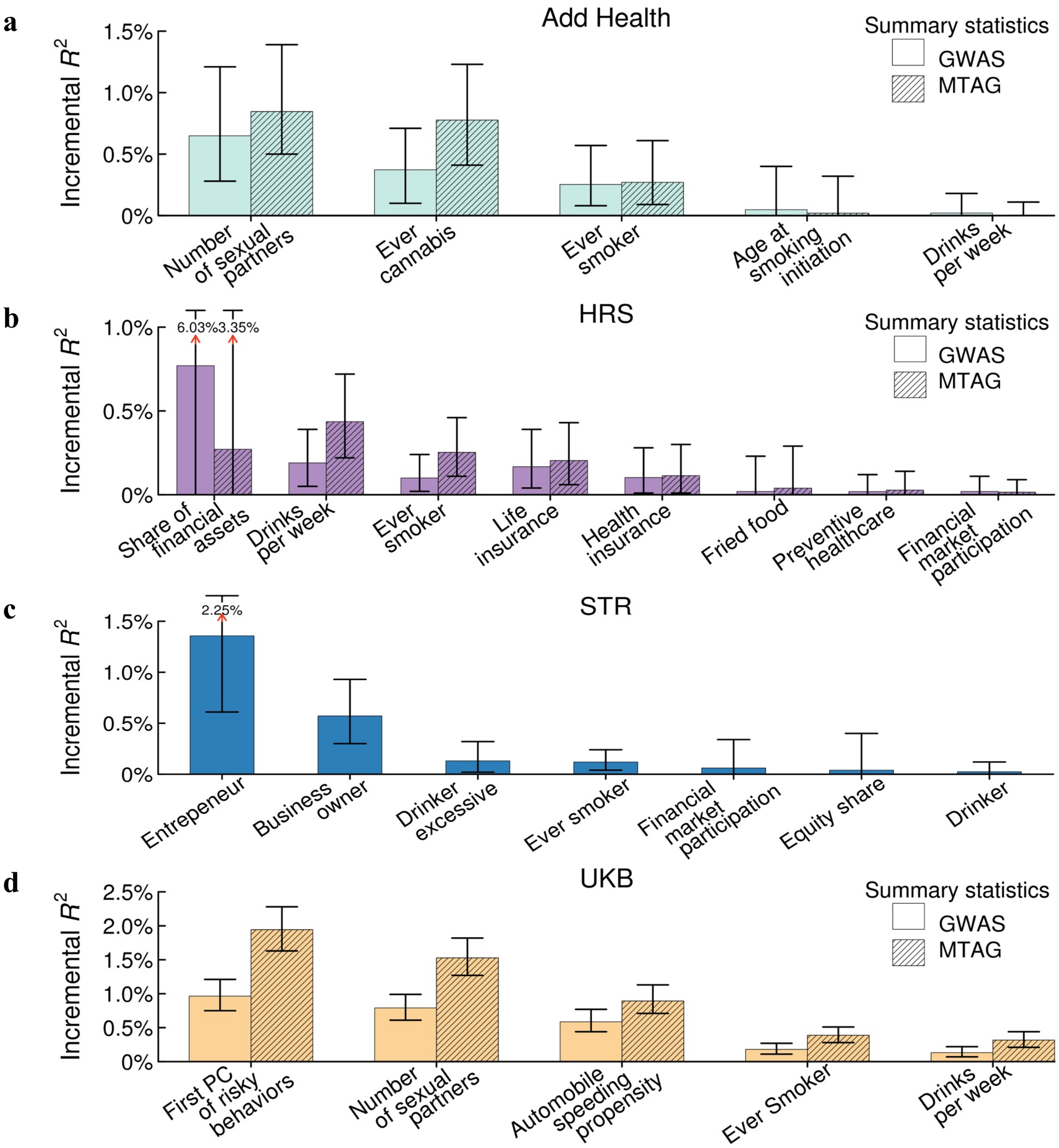


**Extended Data Figure 10.2 | Prediction of risky behaviors with polygenic scores of general risk tolerance.** Incremental $R^2$ is defined as the increase in $R^2$ from adding the score to a regression of the risky behavior on controls for sex, age, and the top ten principal components of the genetic relatedness matrix. The scores were constructed using LDpred with our preferred Gaussian mixture weight of 0.3. Panels **a** to **d** display the results for the Add Health, HRS, STR and UKB-siblings cohorts, respectively. For the Add Health and HRS cohorts, scores were constructed using summary statistics from the meta-analysis of the discovery and replication GWAS ($n$ = 975,353) and from the MTAG analysis of general risk tolerance; for the UKB-siblings cohort, scores were constructed in the same way but excluding individuals with at least one full sibling in the UKB from the meta-analysis ($n$ = 937,353); for the STR cohort, scores were constructed only using summary statistics from a meta-analysis that excludes the 23andMe cohort (due to data access limitations) and the STR cohort ($n$ = 458,558). See **Supplementary Information section 10** and **Supplementary Table 10.3** for additional details.

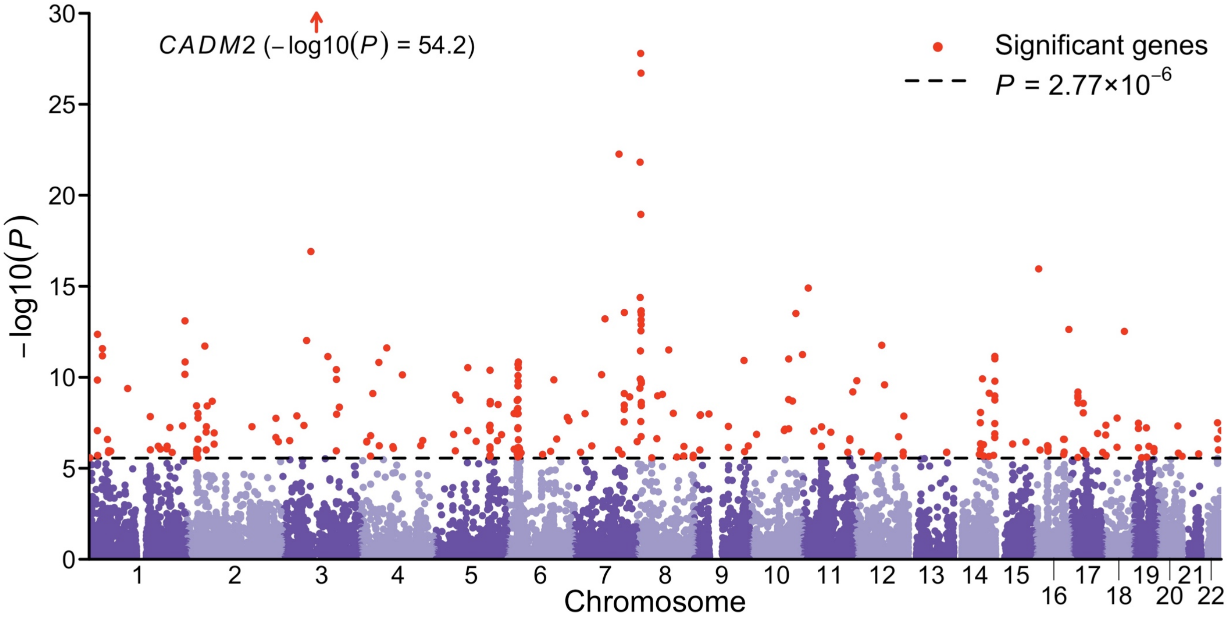


**Extended Data Figure 12.1 | Manhattan plot of the MAGMA gene analysis of general risk tolerance.** The $x$-axis is chromosomal position, and the $y$-axis is the significance of each gene on a $-\log_{10}$ scale. The dashed line marks the Bonferroni-corrected significance threshold (the adjustment is for 18,070 tests, for 18,070 genes; $P = 0.05/18{,}070 \approx 2.77 \times 10^{-6}$). Each of the 285 significant genes are marked with by a red •.

**a**

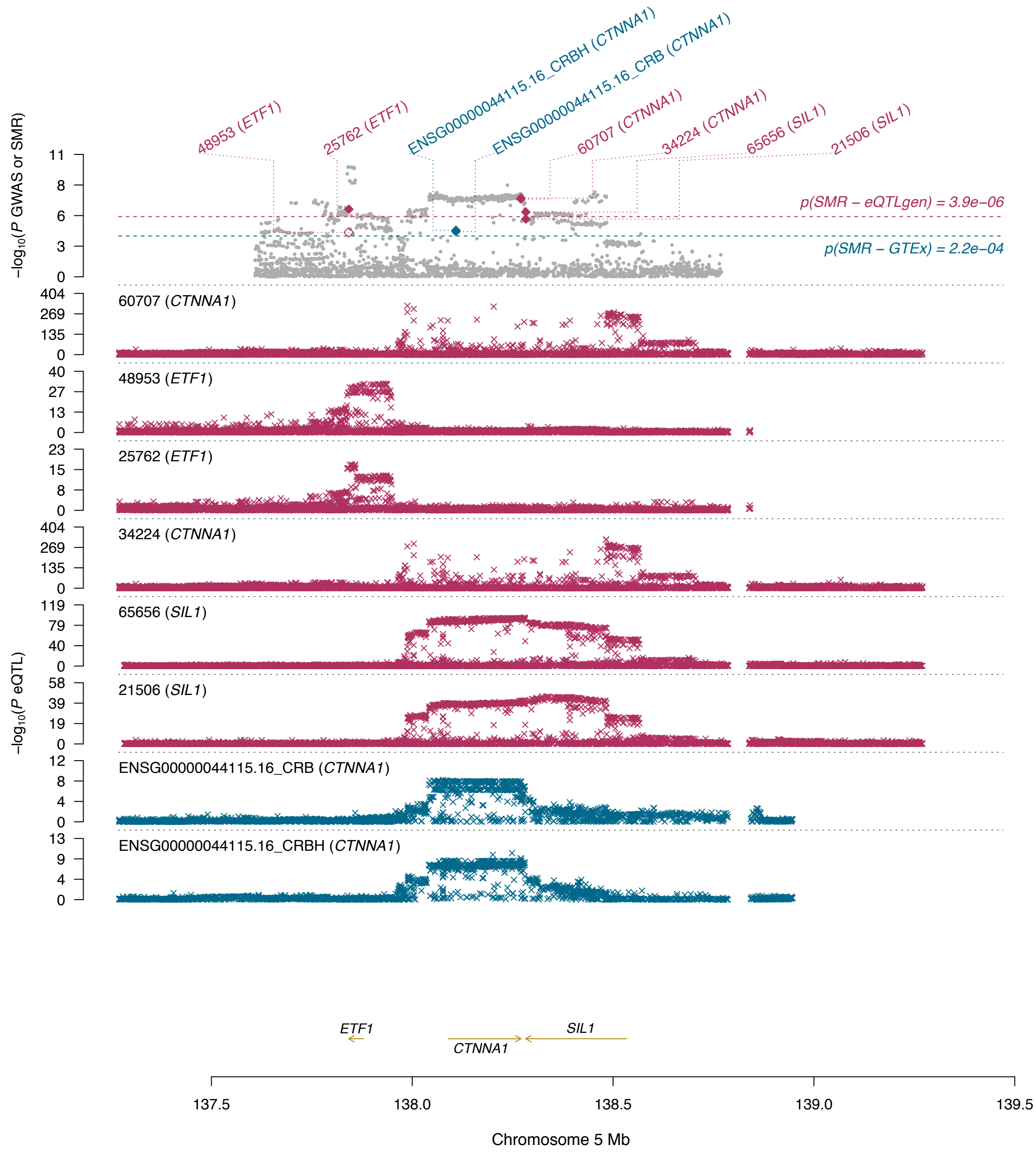

48953 (ETF1)
25762 (ETF1)
ENSG00000044115.16_CRBH (CTNNA1)
ENSG00000044115.16_CRB (CTNNA1)
60707 (CTNNA1)
34224 (CTNNA1)
65656 (SIL1)
21506 (SIL1)
−log10(P GWAS or SMR)
p(SMR − eQTLgen) = 3.9e−06
p(SMR − GTEx) = 2.2e−04
60707 (CTNNA1)
48953 (ETF1)
25762 (ETF1)
34224 (CTNNA1)
65656 (SIL1)
21506 (SIL1)
ENSG00000044115.16_CRB (CTNNA1)
ENSG00000044115.16_CRBH (CTNNA1)
−log10(P eQTL)
ETF1
SIL1
CTNNA1
137.5
138.0
138.5
139.0
139.5
Chromosome 5 Mb

**b**

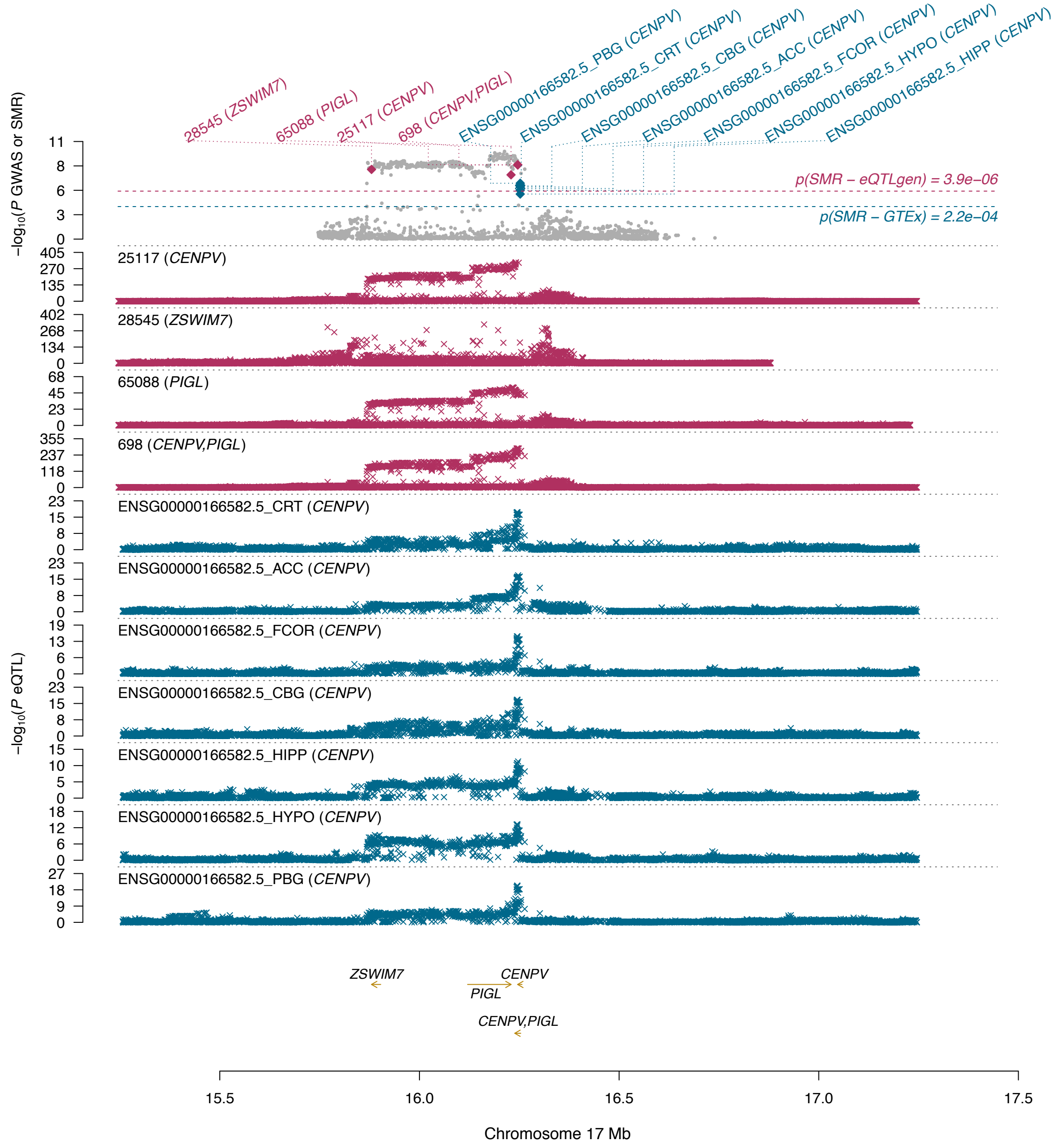

28545 (ZSWIM7)
65088 (PIGL)
25117 (CENPV)
698 (CENPV,PIGL)
ENSG00000166582.5_PBG (CENPV)
ENSG00000166582.5_CRT (CENPV)
ENSG00000166582.5_CBG (CENPV)
ENSG00000166582.5_ACC (CENPV)
ENSG00000166582.5_FCOR (CENPV)
ENSG00000166582.5_HYPO (CENPV)
ENSG00000166582.5_HIPP (CENPV)
−log10(P GWAS or SMR)
p(SMR − eQTLgen) = 3.9e−06
p(SMR − GTEx) = 2.2e−04
−log10(P eQTL)
25117 (CENPV)
28545 (ZSWIM7)
65088 (PIGL)
698 (CENPV,PIGL)
ENSG00000166582.5_CRT (CENPV)
ENSG00000166582.5_ACC (CENPV)
ENSG00000166582.5_FCOR (CENPV)
ENSG00000166582.5_CBG (CENPV)
ENSG00000166582.5_HIPP (CENPV)
ENSG00000166582.5_HYPO (CENPV)
ENSG00000166582.5_PBG (CENPV)
ZSWIM7
CENPV
PIGL
CENPV,PIGL
15.5
16.0
16.5
17.0
17.5
Chromosome 17 Mb

c

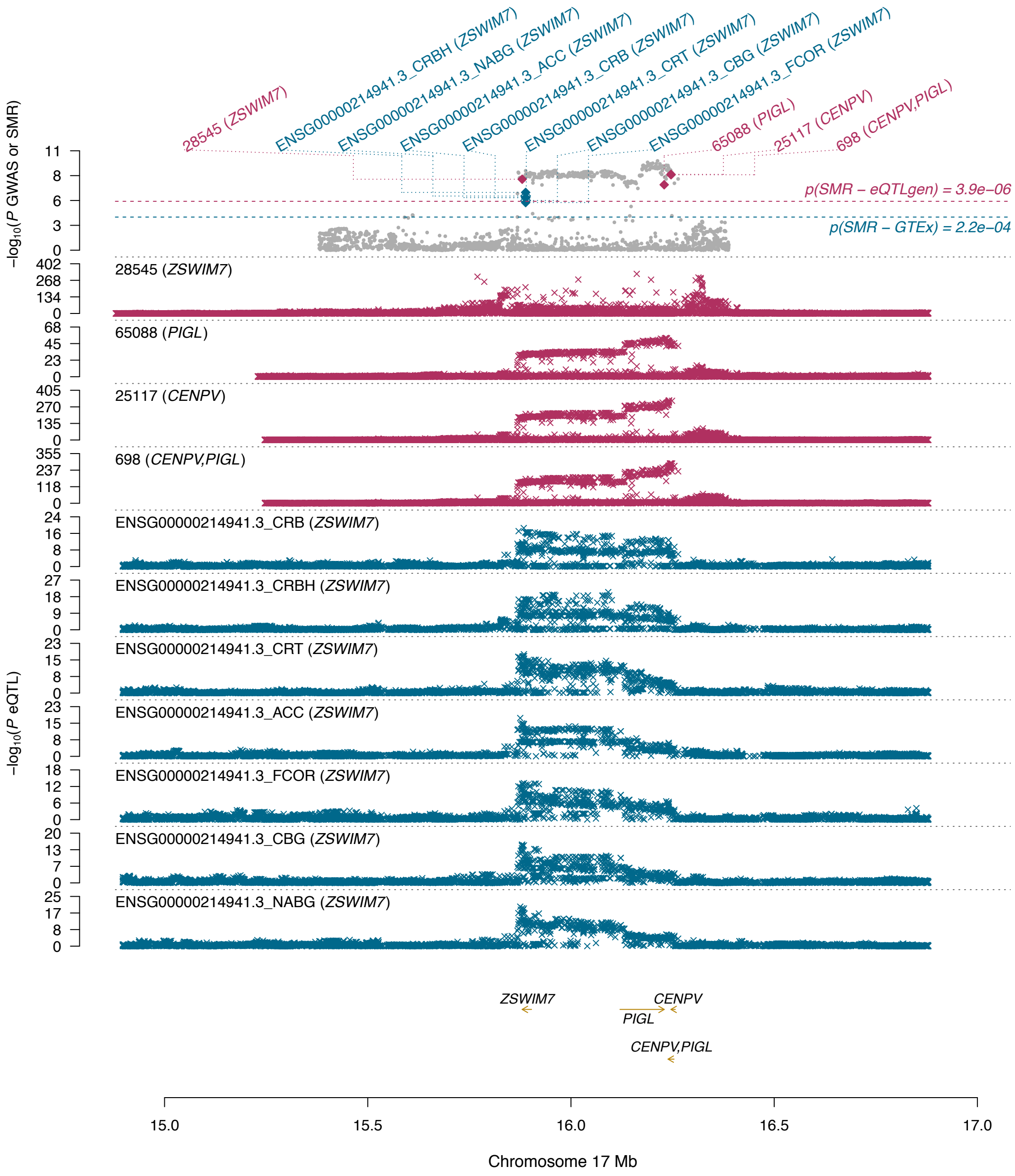


**Extended Data Figure 12.2 | SMR results for general risk tolerance for the genes (a) *CTNNA1,* (b) *CENPV* and (c) *ZSWIM7.*** In each panel, the top plot shows *P* values for SNPs from the GWAS (grey dots) and the SMR test (blue and red diamonds) and the red and blue horizontal dashed lines show the significance threshold for the SMR test in eQTLgen ($P_{\text{SMR-eQTLgen}} = 3.9 \times 10^{-6}$) and GTEx ($P_{\text{SMR-GTEx}} = 2.2 \times 10^{-4}$), respectively. The bottom plots show, in red, eQTL *P* values of SNPs from the eQTLgen study (blood) and, in blue, various GTEx brain regions (PBG: putamen basal ganglia; NABG, nucleus accumbens basal ganglia; CBG, caudate basal ganglia; ACC, anterior cingulate cortex (BA24); HIPP, hippocampus; HYPO, hypothalamus; CRB, cerebellum; CRBH, cerebellar hemisphere; COR, cortex; FCOR, frontal cortex (BA9)).

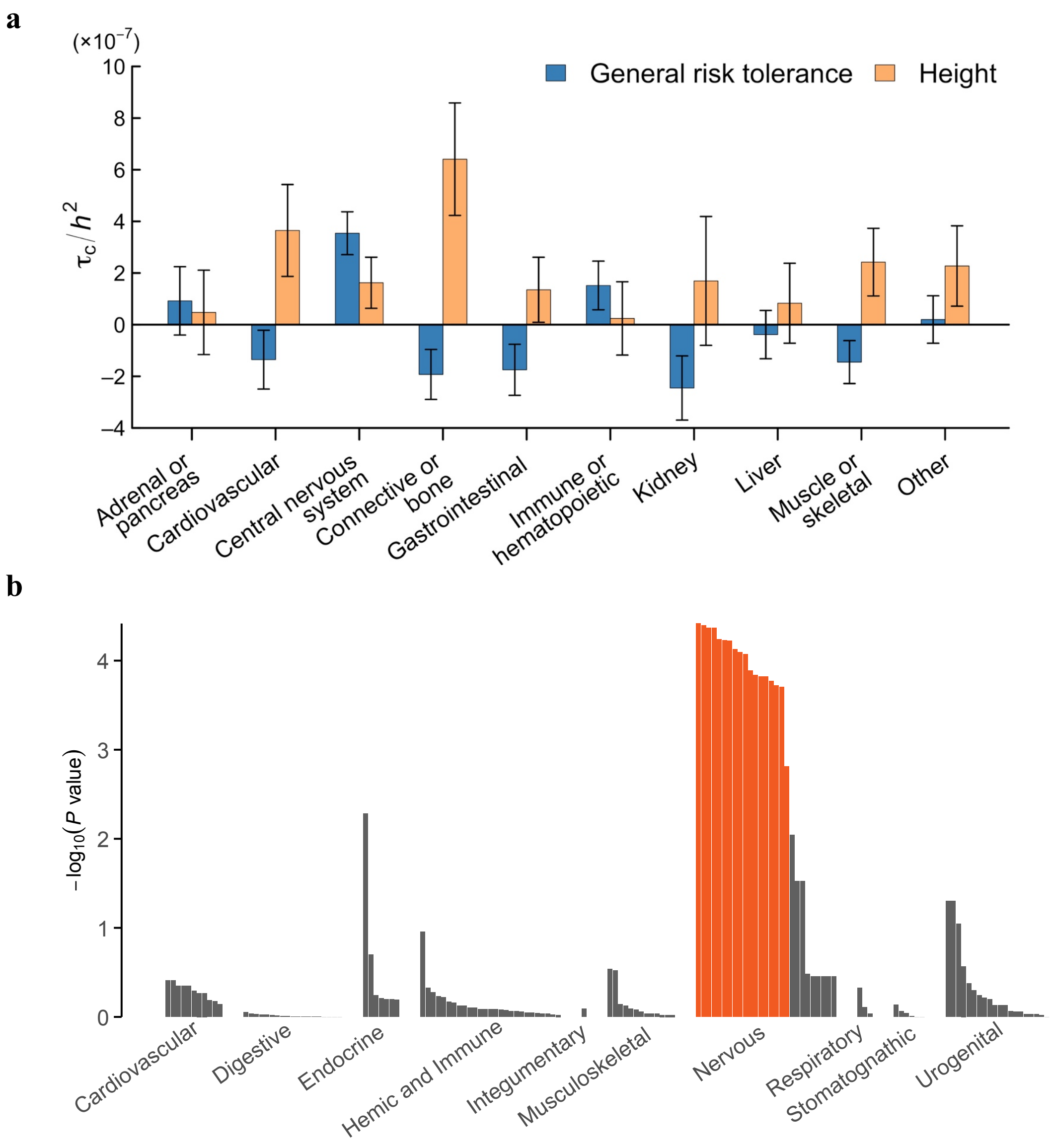


**Extended Data Figure 12.3 | Additional results from selected biological analyses. a**, Functional partitioning of the heritability of general risk tolerance with stratified LD Score regression. The panel shows the expected increase in the phenotypic variance accounted for by a SNP due to the SNP's being in a given category ($\tau_C$), divided by the LD Score heritability of the phenotype ($h^2$). Each estimate of $\tau_C$ comes from a separate stratified LD Score regression, controlling for the 52 functional annotation categories in the baseline model. Error bars represent 95% CIs (not adjusted for multiple testing). To benchmark the estimates, we compare them to those obtained from a recent study of height[3]. **b**, Results of a DEPICT tissue enrichment analysis using microarray-based gene expression data from Fehrmann *et al.*[4] and Pers *et al.*[5]. The panel shows whether the genes overlapping loci associated with general risk tolerance are significantly overexpressed (relative to genes in random sets of loci matched by gene density) in various tissues. Tissues are grouped by physiological system. The orange bars correspond to tissues with significant overexpression (FDR < 0.01). The *y*-axis depicts *P* values on a $-\log_{10}$ scale. See **Supplementary Information section 12** for additional details.

**References:**


1. Okbay, A. *et al.* Genome-wide association study identifies 74 loci associated with educational attainment. *Nature* **533,** 539–542 (2016).
2. Berisa, T. & Pickrell, J. K. Approximately independent linkage disequilibrium blocks in human populations. *Bioinformatics* **32,** 283–285 (2016).
3. Wood, A. R. *et al.* Defining the role of common variation in the genomic and biological architecture of adult human height. *Nat. Genet.* **46,** 1173–1186 (2014).
4. Fehrmann, R. S. N. *et al.* Gene expression analysis identifies global gene dosage sensitivity in cancer. *Nat. Genet.* **47,** 115–125 (2015).
5. Pers, T. H. *et al.* Biological interpretation of genome-wide association studies using predicted gene functions. *Nat. Commun.* **6,** 5890 (2015).
